# The *MUC19* Gene: An Evolutionary History of Recurrent Introgression and Natural Selection

**DOI:** 10.1101/2023.09.25.559202

**Authors:** Fernando A. Villanea, David Peede, Eli J. Kaufman, Valeria Añorve-Garibay, Elizabeth T. Chevy, Viridiana Villa-Islas, Kelsey E. Witt, Roberta Zeloni, Davide Marnetto, Priya Moorjani, Flora Jay, Paul N. Valdmanis, María C. Ávila-Arcos, Emilia Huerta-Sánchez

## Abstract

We study the gene *MUC19*, for which some modern humans carry a *Denisovan-like* haplotype. *MUC19* is a mucin, a glycoprotein that forms gels with various biological functions. We find diagnostic variants for the *Denisovan-like MUC19* haplotype at high frequencies in admixed Latin American individuals, and at highest frequency in 23 ancient Indigenous American individuals, all predating population admixture with Europeans and Africans. We find that the *Denisovan-like MUC19* haplotype is under positive selection and carries a higher copy number of a 30 base-pair variable number tandem repeat, and that copy numbers of this repeat are exceedingly high in American populations. Finally, some Neanderthals carry the *Denisovan-like MUC19* haplotype, and that it was likely introgressed into human populations through Neanderthal introgression rather than Denisovan introgression.

**One-Sentence Summary:** Modern humans and Neanderthals carry a Denisovan variant of the *MUC19* gene, which is under positive selection in populations of Indigenous American ancestry.

## Main Text

Most modern humans of non-African ancestry carry both Neanderthal and Denisovan genomic variants [*1–3*]. While most of these variants are putatively neutral, some archaic variants found in modern humans have been targets of positive natural selection [*4–9*]. Interbreeding with Neanderthals and Denisovans may have thereby facilitated adaptation to the myriad novel environments that modern humans encountered as they populated the globe [*10*]. Indeed, several studies have identified signatures of adaptive introgression in Eurasian and Oceanian populations [*11–20*]. Indigenous American populations, however, present great potential for studying the underlying evolutionary processes of local adaptation [*21*]. In the 25,000 years since the first individuals populated the American continent, these populations would have encountered manifold novel environments, far different from the Beringian steppe, to which their ancestral population was adapted [*22*].

Previous studies identified *MUC19*—a gene involved in immunity—as a candidate for adaptive introgression among populations from the 1000 Genomes Project (1KG). These studies found the region surrounding *MUC19* to harbor several Denisovan variants in Mexicans (MXL) [*23*]; and reported that this region has one of the largest densities of Denisovan alleles in Mexicans [*24*]. *MUC19* was also reported to be under positive selection in North American Indigenous populations using Population Branch Statistic (*PBS*) and integrated Haplotype Scores (*iHS*) methods for detecting positive selection [*25*].

In this study, we confirm and further characterize signatures of both introgression and positive selection at *MUC19* in MXL. We find an archaic haplotype segregating at high frequency in most populations on the American continent, which is also present in two of the late high-coverage Neanderthal genomes—Chagyrskaya and Vindija. MXL individuals harbor Denisovan-specific coding mutations in *MUC19* at high frequencies, and exhibit elevated copy number of a tandem repeat region within *MUC19* compared to other worldwide populations. Our results point to a complex pattern of multiple introgression events, from Denisovans to Neanderthals, and Neanderthals to modern humans, which may have played a unique role in the evolutionary history of Indigenous American populations.

## Results

### Signatures of adaptive introgression at *MUC19* in admixed populations from the Americas

We compiled introgressed tracts that overlap the NCBI RefSeq coordinates for *MUC19* (hg19, Chr12:40787196-40964559) by at least one base pair. Figure 1A shows the density of introgressed tracts for all non-African populations in the region, using introgression maps inferred with `hmmix` [*26*]. All non-African populations harbor introgressed tracts overlapping this region, but at much lower frequencies than the American populations (AMR tract frequency: ∼0.183, non-AMR tract frequency: ∼0.087; Proportions Z-test, *P-value*: 5.011e-14; Fisher’s Exact Test, *P-value*: 2.144e-12; Table S1). Mexicans (MXL)—a population with a large component of Indigenous American genetic ancestry (∼48%; [*27*])—exhibits the highest frequency of the introgressed tracts (0.305; Table S2). Given this, we examined a 742kb window containing the longest introgressed tract found in Mexicans (hg19, Chr12:40272001-41014000; Figure S1). This region contains 135 Denisovan-specific SNPs, classified as such because they are rare or absent in African populations (<1%), present in MXL (>1%), and shared uniquely with the Altai Denisovan. All 135 of these SNPs are sequestered within a core 72kb region (hg19, Chr12:40759001-40831000; shaded gray region in Figure 1A) that has the highest introgressed tract density amongst individuals in the 1KG (see [51]), making both the 742kb and 72kb region outliers for Denisovan-specific SNP density in MXL (742kb region *P-value*: <3.164e-4; 72kb region *P-value*: <3.389e-5; Figure S2; Table S3-S4). In contrast, there are 80 Neanderthal-specific SNPs in MXL found within the larger 742kb region (*P-value*: 0.159; Figure S3; Table S5), with only four located in the 72kb region (*P-value*: 0.263; Figure S3; Table S6).

**Figure 1.**
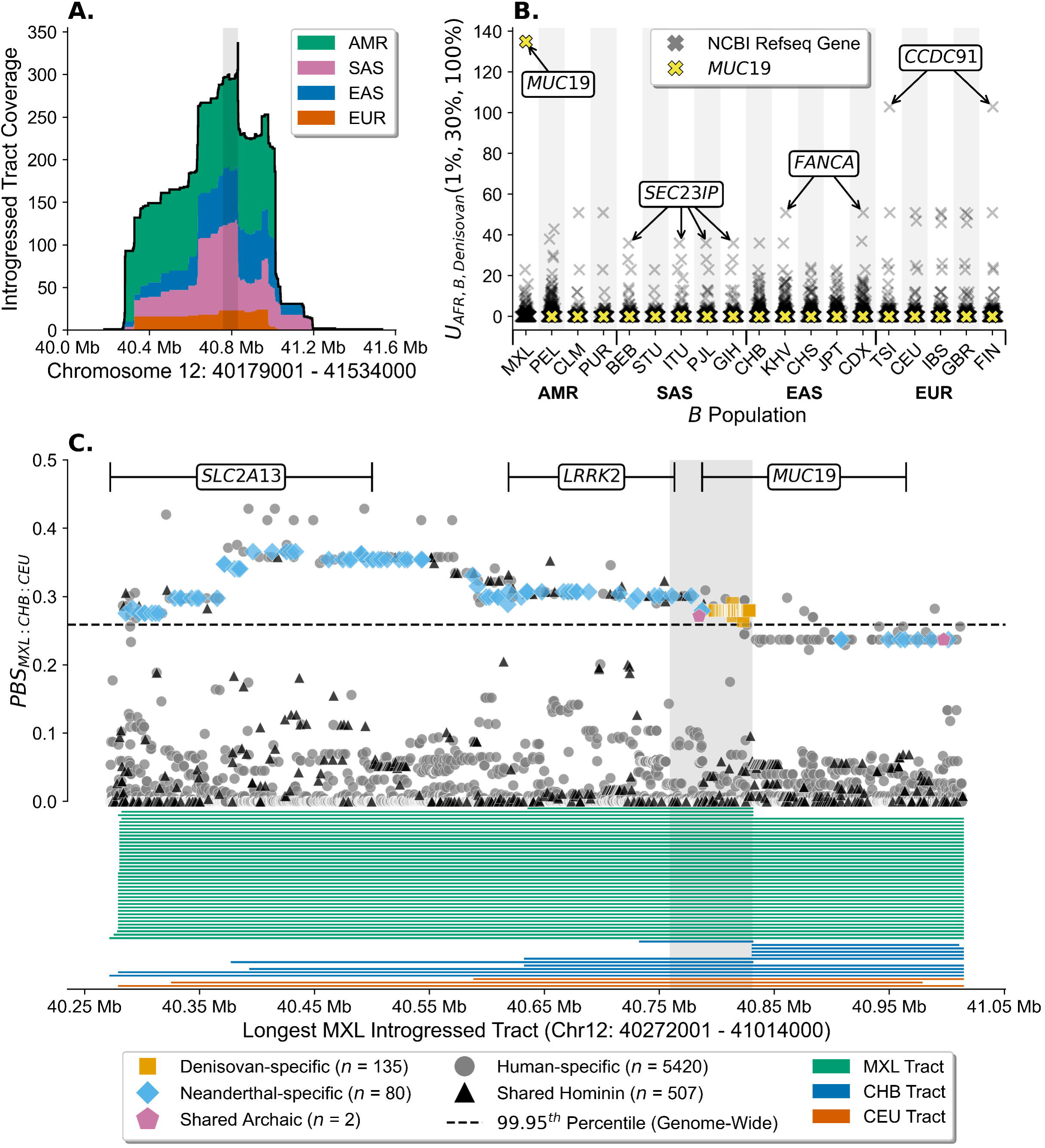
Signals of adaptive introgression at *MUC19*. (**A**) Density of introgressed tracts inferred using `hmmix` that overlap *MUC19* for the 1KG (black outline) and stratified by superpopulation—Admixed Americans (AMR) in bluish green, South Asians (SAS) in reddish purple, East Asians (EAS) in blue, and Europeans (EUR) in vermillion. The gray shaded region corresponds to the focal 72kb region, which is the densest contiguous region of introgressed tracts longer than 40kb. (**B**) *U_AFR,B,Denisovan_(1%, 30%, 100%)* values for each non-African population, stratified by superpopulation, per NCBI Refseq gene (gray X’s), where *MUC19* is denoted as a yellow X. (**C**) Population Branch Statistic (*PBS*) for the Mexican population (MXL) in the 1KG using the Han Chinese (CHB) and Central European (CEU) populations in the 1KG as control populations (*PBS_MXL:CHB:CEU_*) for all SNPs in the 742kb region that corresponds to the longest introgressed tract found in MXL. The orange squares represent Denisovan-specific SNPs, the sky blue diamonds represent Neanderthal-specific SNPs, and the reddish purple pentagons represent shared archaic SNPs—note that all of these archaic SNP partitions are rare or absent in Africa and present in MXL (see [51]). The black triangles represent SNPs present across both modern human populations and the archaics, while the gray circles represent SNPs private to modern humans. The black dashed line represents the 99.95th percentile of *PBS_MXL:CHB:CEU_* scores for all SNPs genome-wide, and the gray shaded region corresponds to the focal 72kb region—the same gray shaded region in panel A. The *MUC19* and *LRRK2* genes are fully encompassed within the 742kb region, while ∼65% of *SLC2A13* overlaps the 742kb region. Below the *PBS_MXL:CHB:CEU_* points are the introgressed tracts for MXL (bluish green), CHB (blue), and CEU (vermillion) sorted from shortest to longest within each population.

To test if natural selection is acting on this region, we computed three statistics; one developed to detect adaptive introgression (*U_A,B,C_(w, x, y)*, *A:* African super population, *B:* non-African populations, *C:* Altai Denisovan; (*w, x, y*) are allele frequency thresholds in *A, B* and *C,* [*24*]), and two for positive selection (*PBS*, and *iHS*). For each gene, we computed *U_AFR,B,Denisovan_(w=1%, x=30%, y=100%)*, which measures the number of Denisovan alleles found in the homozygous state (100%) that are almost absent in Africans (<1%) and reach a frequency of at least 30% in a given non-African population. Figure 1B shows that *MUC19* in MXL is an extreme outlier, as no other gene in any non-African population exhibits such a large value of *U_AFR,B,Denisovan_(1%, 30%, 100%)*. When we compute the same statistic in windows instead of per gene, the *MUC19* region is an outlier only in MXL and is zero for all other non-African populations (*P-value* 72kb region: <3.284e-5; *P-value* 742kb region: <3.139e-4: Figure S4; Table S7-S8). Furthermore, we compared the windowed *U_AFR,B,Denisovan_(w=1%, x=30%, y=100%)* results with their corresponding *Q95_AFR,B,Denisovan_(w=1%, y=100%)* value, which quantifies the 95th percentile of the Denisovan allele frequencies found in a given non-African population *B* for the Denisovan alleles found in the homozygous state (100%), that are almost absent in Africans (<1%), we find that for both the 72kb and 742kb *MUC19* regions that *Q95_AFR,MXL,Denisovan_(w=1%, y=100%)* = ∼30%, which suggests that both the 72kb and 742kb *MUC19* regions exhibit signals consistent with adaptive introgression that are not observed in any other 1KG population (Figure S5-S6; Table S9).

We next computed *PBS_MXL:CHB:CEU_*, where the Han Chinese (CHB) and Central European (CEU) populations were used as control populations, for both the region corresponding to the longest introgressed tract in MXL—742kb—and the 72kb region in *MUC19*, and find that both regions exhibit statistically significant *PBS_MXL:CHB:CEU_* values compared to other 742kb (*PBS_MXL:CHB:CEU_*: 0.066; *P-value*: 0.004) and 72kb (*PBS_MXL:CHB:CEU_*: 0.127; *P-value*: 0.002) windows of the genome respectively (Figure S7; Table S10-S11). We then computed *PBS_MXL:CHB:CEU_* for each SNP in the 742kb region. Figure 1C shows that in MXL there are many SNPs with statistically significant PBS values in that region (417 out of 6144 SNPs), all which present values above the 99.95th percentile of genome-wide *PBS_MXL:CHB:CEU_* values (Benjamini-Hochberg corrected *P-values*: <0.01; see Supplemental Section S1 in [51]). We note that some SNPs have a larger *PBS_MXL:CHB:CEU_* value near the *SLC2A13* gene than within the 72kb *MUC19* region, but this is due to changes in the archaic allele frequency in CHB and CEU, as the introgressed tracts in these populations are more sparse than the introgressed tracts in MXL (see tracts in Figure 1C). When we partition the MXL population into two demes, consisting of individuals with more than 50% and those with less than 50% Indigenous American ancestry genome-wide [*27*], and recompute *PBS,* we find that *PBS* values for archaic variants are elevated among individuals with a higher proportion of Indigenous American ancestry, suggesting that this region was likely targeted by selection before admixture with European and African populations (Figure S8; Table S10-S11).

To exclude the possibility that demographic events such as a founder effect explain the observed signatures of positive selection, we simulated the best fitting demographic parameters inferred for the MXL population [*28*] to obtain the expected null distribution of *PBS* values. We first showed that *PBS* has power to detect adaptive introgression under this demographic model (see Supplemental Section S1 in [51]). We found that demographic forces alone result in lower *PBS* values compared to what is observed at this gene region (see Supplemental Section S1 in [51]), even when we consider a very conservative null model of heterosis. Furthermore, to also consider haplotype-based measures of positive selection, we computed the integrated haplotype score (*iHS*) for every 1KG population using `selscan` [*29*] to provide haplotype-based evidence of natural selection ([51]). Among all 1KG populations, MXL is the only population with an elevated proportion of SNPs with normalized |*iHS*| > 2 in either the 742kb (599 out of 2248 SNPs) or 72kb region (229 out of 425 SNPs; Table S12-S13). In MXL we find that 130 out of the 135 Denisovan-specific SNPs in the 72kb region have normalized |*iHS*| > 2, reflective of positive selection (Figure S9; Table S12-S13, see Supplemental Section S2 in [51]), which supports our previous allele frequency-based tests of natural selection.

### Admixed individuals exhibit an elevated number of variable number tandem repeats at *MUC19*

*MUC19* contains a 30 base pair variable number tandem repeat (VNTR; hg19, Chr12:40876395-40885001; Figure S10), located 45.4kb away from the core 72kb haplotype, but within the larger 742kb introgressed region. To test if individuals who harbor an introgressed tract overlapping the repeat region differ in the number of repeats compared to individuals who do not harbor introgressed tracts, we calculated the number of repeats of the 30bp motif in the 1KG individuals (see [51]; Figure S11; Table S14-S15). For each individual, we first report the average number of repeats between their two chromosomes. The genomes of the four archaic individuals do not harbor a higher copy number of tandem repeats (Altai Denisovan: 296 copies; Altai Neanderthal: 379 copies; Vindija Neanderthal: 268 copies; and Chagyrskaya Neanderthal: 293 copies). Among all individuals from the 1KG, we identified outlier individuals with elevated number of repeats above the 95th percentile (>487 repeats; dashed line in Figure 2). We found that MXL individuals have on average ∼493 repeats and individuals from the admixed American super population have on average ∼417 repeats (Figure 2A; Table S16-S17). In contrast, non-admixed American populations have an average of ∼341 to ∼365 repeats (Figure 2A; Table S16). Out of all the outlier individuals from the 1KG (>487 repeats), a significant proportion of them (∼77%) are from admixed American populations (Proportions Z-test, *P-value*: 3.971e-17; Table S18-S21; Figure S12). Outlier individuals from the Americas also carry a significantly higher copy number of tandem repeats compared to the other outlier individuals from non-admixed American populations (Mann-Whitney U, *P-value*: 5.789e-7; Figure S12; Table S18-S21). In MXL, we find that exactly 50% of individuals exhibit an elevated copy number of tandem repeats (Table S16).

**Figure 2.**
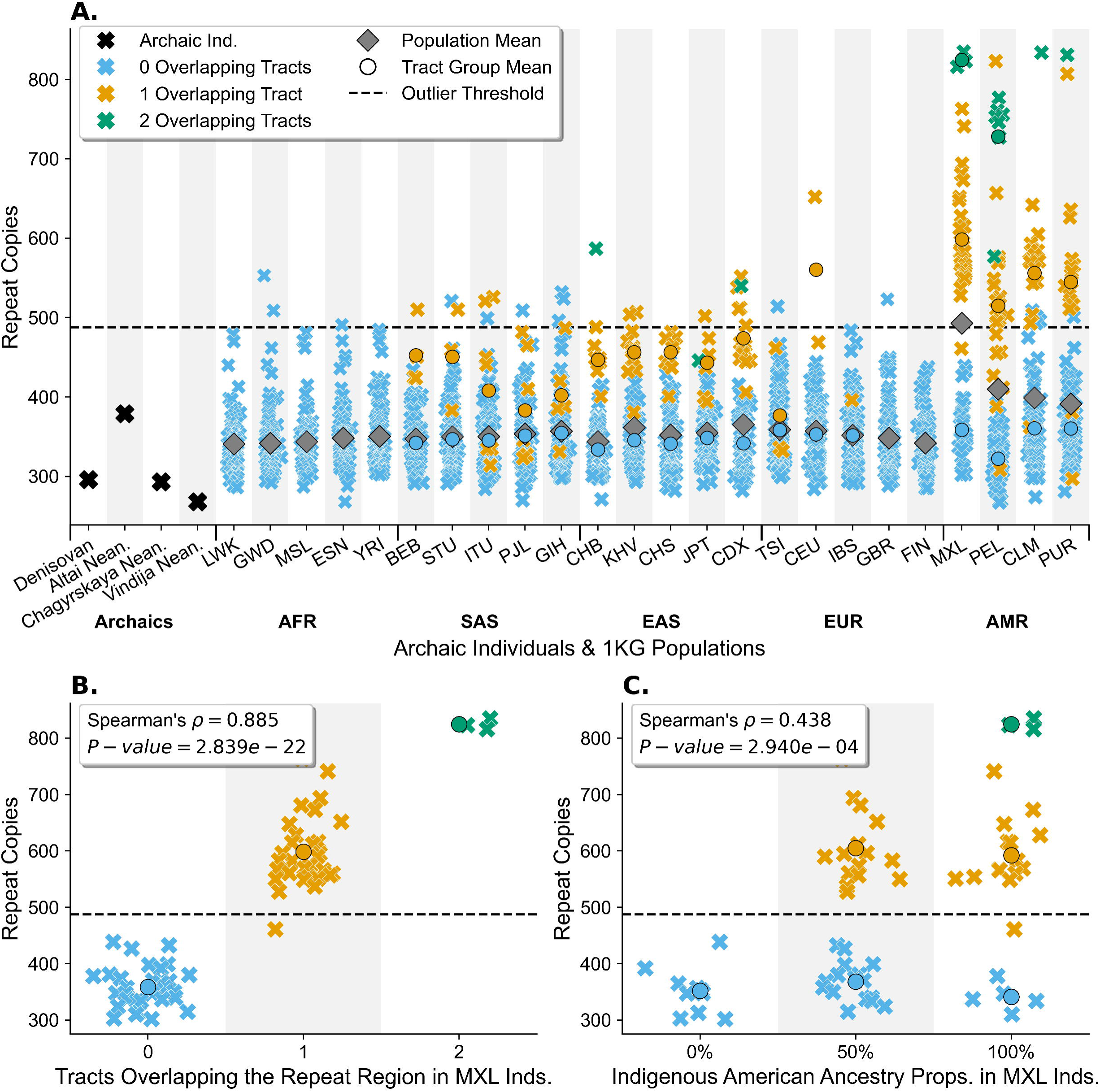
Copy number variation of a 30 base pair variable number tandem repeat motif in the 1KG individuals at *MUC19*. (**A**) Average number of repeat copies between an individual’s two chromosomes for archaic individuals (black X’s), individuals who do not harbor an introgressed tract (sky blue X’s), individuals with one introgressed tract (yellow X’s), and individuals with two introgressed tracts (bluish green X’s) determined by the number of introgressed tracts inferred using `hmmix` overlapping the *MUC19* VNTR, for each population in the 1KG. The mean number of repeat copies stratified by population is denoted by a grey diamond and the average number of repeat copies amongst individuals who carry exactly zero, one, and two introgressed tracts are denoted by sky blue, yellow, and bluish green circles respectively and are stratified by population. The black dashed line denotes the outlier threshold, which corresponds to the 95th percentile of the 1KG repeat copies distribution. Repeat copies appeared similar to the reference human genome (287.5 copies) in the Altai Denisovan (296 copies) and Altai (379 copies), Vindija (268 copies), and Chagyrskaya (293 copies) Neanderthal genomes. (**B**) The relationship between the average number of repeat copies between a MXL individual’s two chromosomes and the number of introgressed tracts overlapping the *MUC19* VNTR region. Note that there is a significant positive correlation between the number of repeat copies and the number of introgressed tracts present in an MXL individual (Spearman’s *ρ*: 0.885; *P-value*: 2.839e-22). (**C**) The relationship between the average number of repeat copies between a MXL individual’s two chromosomes and the proportion of Indigenous American ancestry at the *MUC19* VNTR region. Note that there is a significant positive correlation between the number of repeat copies and the proportion of Indigenous American ancestry in an MXL individual (Spearman’s *ρ*: 0.438; *P-value*: 2.940e-4).

Within individuals exhibiting an outlier number of repeats (>487), a significant proportion (∼86%) have an introgressed tract overlapping the repeat region and these individuals harbor an elevated number of repeats compared to outlying individuals who do not harbor an introgressed tract overlapping the VNTR region (Proportions *Z*-test, *P-value:* 2.127e-29; Mann-Whitney U, *P-value:* 1.398e-06; Figure S13; Table S18-S21). All outlying MXL individuals carry at least one introgressed tract that overlaps with the VNTR region (Figure 2). MXL has more individuals exhibiting an elevated copy number (>487 repeats) than any other 1KG population, and there is a positive correlation between the number of repeats and the number of introgressed tracts that overlap with the VNTR present in a MXL individual (Spearman’s *ρ*: 0.885; *P-value*: 2.839e-22; Figure 2B; Figure S14; Table S22). We find that among MXL individuals, the number of repeats and the Indigenous American ancestry proportion at the repeat region is significantly positively correlated (Spearman’s *ρ*: 0.483; *P-value*: 2.940e-4; Figure 2C; Figure S15, Table S23-S24), while the African (Spearman’s *ρ*: -0.289; *P-value*: 2.072e-2; Figure S15, Table S23-S24) and European (Spearman’s *ρ*: -0.353; *P-value*: 4.191e-3; Figure S15, Table S23-S24) ancestry proportions have a significant negative correlation. Taken together, in MXL, we find that an individual’s VNTR copy number is highly predicted by the number of introgressed tracts that overlap the VNTR. To a lesser extent, the VNTR copy number is also predicted by the Indigenous American ancestry proportion in the repeat region, indicating that individuals with elevated VNTR copy number have higher proportions of Indigenous American ancestry and harbor the introgressed haplotype. Individuals who carry an elevated number of the *MUC19* VNTR are likely to also carry the archaic haplotype, especially in admixed American populations where the archaic haplotype of *MUC19* is found at highest frequencies (Mann-Whitney U, *P-value:* 1.597e-87; Figure S13; Figure 2; Table S18-S21).

Given the difficulties of calling numbers of repeats from short-read data, we examined long-read sequence data from the Human Pangenome Reference Consortium (HPRC) and Human Genome Structural Variant Consortium (HGSVC) [42]. These corroborated our findings (Figure S10; Figure S16), revealing an extra 424 copies of the 30bp *MUC19* tandem repeat exclusively in American samples, arranged in four additional segments of 106 repeats (at 3,171 bp each). This structural variant is exceptionally large; it effectively doubles the size of the ∼12kb coding exon that harbors the tandem repeat (Figures S10).

### Introgression introduced missense variants at *MUC19*

Inspecting the 135 Denisovan-specific SNPs and 4 Neanderthal-specific SNPs in the core 72kb region reveals that some modern humans carry two Denisovan-specific synonymous sites and nine Denisovan-specific non-synonymous sites (Table S25). We quantified the allele frequencies for these nine Denisovan-specific missense variants in present-day populations and in 23 ancient Indigenous American genomes that predate European colonization and the African slave trade (Figure 3A; Table S26-S33). In the admixed American superpopulation, we find that the Denisovan-specific missense mutations are segregating at the highest frequencies (frequency range in AMR,: ∼0.154 - ∼0.157) compared to all other 1KG superpopulations (frequency range in non-AMR,: ∼0 - ∼0.108; Table S27-S28). When we stratify by population instead of by superpopulation, we find the Denisovan-specific missense mutations are segregating at frequencies between ∼0.069 and ∼0.305 amongst admixed American populations, at varying frequencies between ∼0.005 and ∼0.157 throughout European, East Asian, and South Asian populations, and at the highest frequency in MXL where all nine Denisovan-specific missense mutations are segregating at a frequency of ∼0.305 (Figure 3A; Table S29). We find the mean Denisovan-specific missense mutation frequency to be positively correlated with the introgressed tract frequency per population (Pearson’s *ρ*: 0.976; *P-value*: 5.306e-16; Figure S17).

**Figure 3.**
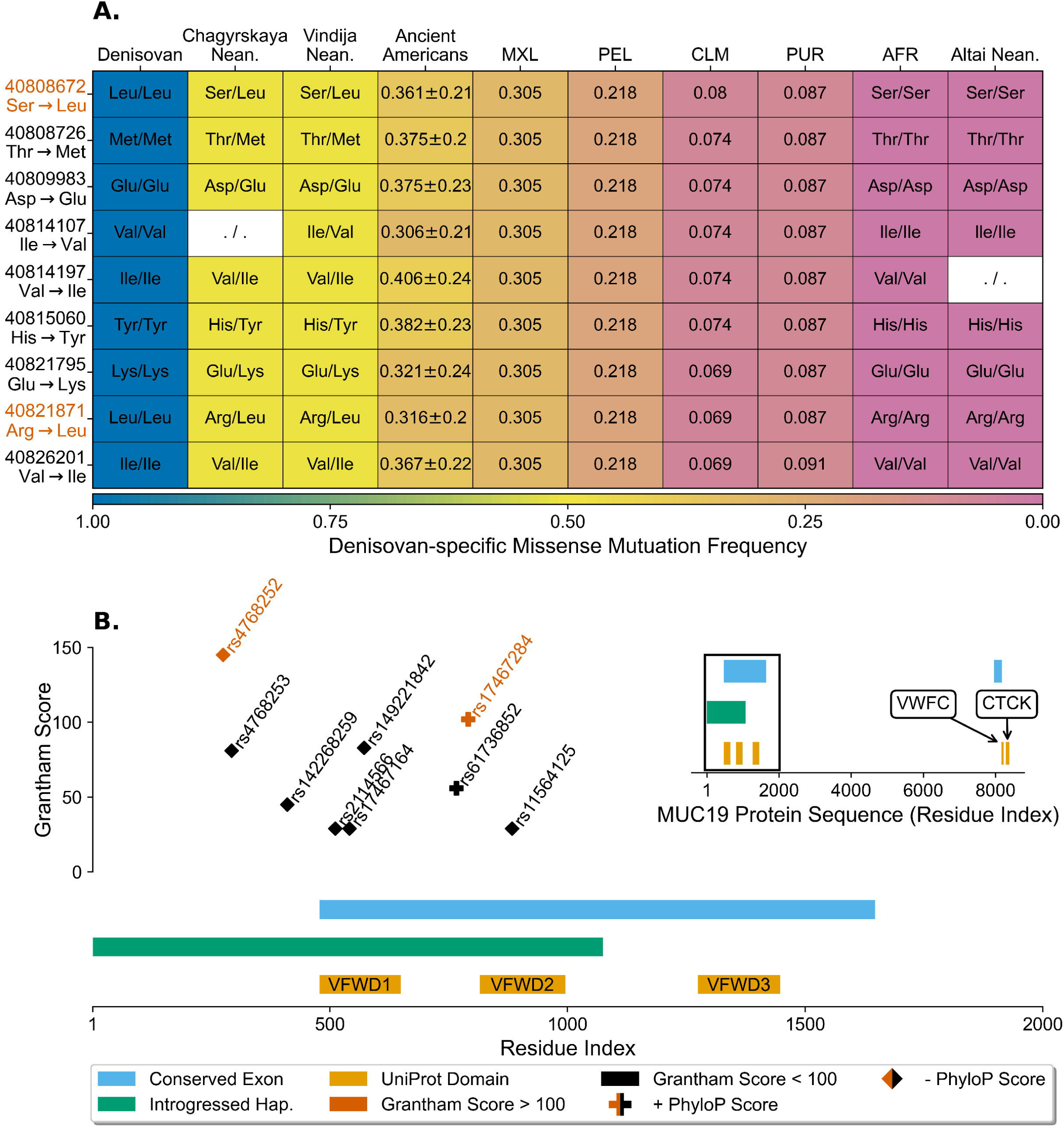
Frequency and protein sequence context of the nine Denisovan-specific missense mutations at the 72kb region in *MUC19*. (**A**) Heatmap depicting the frequency of Denisovan-specific missense mutations (columns) amongst the four archaic individuals (n = 2; per archaic individual), 23 ancient pre-European colonization American individuals (n = 46), the entire African superpopulation in the 1KG (AFR; n = 1008), and admixed American populations in the 1KG—Mexico (MXL; n = 128), Peru (PEL; n = 170), Colombia (CLM; n = 188), Puerto Rico (PUR; n = 208)—where the “n” represents the number of chromosomes in each population. The left hand side of each row denotes one of the nine Denisovan-specific missense mutations where the position and amino acid substitution (hg19 reference amino acid → Denisovan-specific amino acid). The text in each cell represents the Denisovan-specific missense mutation frequency, and for the ancient Americans we also denote the 95% confidence interval. For the archaic individuals, each cell is denoted with the individual’s amino acid genotype and each AFR cell is denoted by the homozygous hg19 reference amino acid genotype. (**B**) Denisovan-specific missense mutations in the context of the MUC19 protein sequence. The first 2000 residues are depicted as the main plot, the full protein sequence is displayed in the smaller subplot. Conserved exons are colored as sky blue and the UniProt domains are colored orange, where the text corresponds to specific UniProt domain identity—Von Willebrand factor (VFW) D domains, VWFC domain, and C-terminal cystine knot-like (CTCK) domain. Each of the nine Denisovan-specific missense mutations are denoted by their rsID, plotted with respect to residue index on the x-axis and their corresponding Grantham score on the y-axis. The color of each Denisovan-specific missense mutation denotes whether the mutation has a Grantham score less than 100 (black) or a Grantham score greater than 100 (vermillion, and the marker denotes whether their respective exon has a negative PhyloP score (diamonds) or a positive PhyloP score (crosses).

We then evaluate the frequency of the nine Denisovan-specific missense mutations in 23 ancient pre-European colonization American individuals, and find that each of the nine Denisovan-specific missense mutations are segregating at higher frequencies than in any admixed American population in the 1KG, but at statistically similar frequencies with respect to MXL (see [51]; Figure 3A; Table S29-S32). These ancient individuals were sampled from a wide geographic and temporal range (Figure S18; Table S26; [51) and do not comprise a single population, yet we detect the presence of the Denisovan-specific missense mutations in sampled individuals from Alaska, Montana, California, Ontario, Central Mexico, Peru, and Patagonia (Table S30). When we quantify the frequency of these mutations in 22 unadmixed Indigenous Americans from the Simons Genome Diversity Project (SGDP), we find that all nine Denisovan-specific missense variants are segregating at a frequency of ∼0.364, which is statistically similar to the ancient American frequencies (see [51]; Table S31-S32), and higher than any admixed American population in the 1KG, albeit at statistically similar frequencies with respect to MXL (Table S31-S32). Given that all nine of the missense mutations are found within a ∼17.5kb region, we quantified the frequency of the Denisovan-specific missense mutation at position Chr12:40808726 in both the ancient individuals and admixed Americans in the 1KG, as this position has genotype information in 20 out of the 23 ancient American individuals (Table S30). We then assessed the relationship between Indigenous American ancestry proportion at the 72kb region, and this Denisovan-specific missense mutation frequency. We find a positive and significant relationship (Pearson’s *ρ*: 0.489; *P-value*: 1.982e-23; Figure S19) between an individual’s Indigenous American Ancestry proportion and their respective Denisovan-specific missense mutation frequency, which suggests that recent admixture in the Americas may have diluted the introgressed ancestry at the 72kb region. We also quantify the frequency of these variants in 44 African individuals from the SGDP, and find all nine Denisovan-specific missense variants at a frequency of ∼0.011, in a single chromosome from a Khomani San individual (Table S33).

To estimate the potential effect of these missense mutations on the MUC19 protein, we relied on Grantham scores [30]. One of the Denisovan-specific missense mutations found at position Chr12:40821871 (rs17467284 in Figure 3B) results in an amino acid change with a Grantham score of 102. This substitution is classified as moderately radical [31] and suggests that the amino acid introduced through introgression is likely to impact the translated protein’s structure or function. This Denisovan-specific missense mutation falls within an exon that is highly conserved across vertebrates (PhyloP score: 5.15, *P-value:* 7.08e-6; Figure 3B) [*32*], indicating that this amino acid residue is likely functionally important, and that the amino acid change introduced by the Denisovan-specific missense mutation may have a significant structural or functional impact. Furthermore, this missense mutation falls between two Von Willebrand factor D domains, which play an important role in the formation of mucin polymers and gel-like matrices [*33*]. Our results suggest that this Denisovan-specific missense mutation is a potential candidate for impacting its translated protein and may affect the polymerization properties of *MUC19* and the viscosity of the mucin matrix.

### Identification of the most likely donor of the introgressed haplotype at *MUC19*

To identify the most likely archaic donor, we investigated the patterns of haplotype divergence at *MUC19* by comparing the modern human haplotypes in the 1KG in the 72kb region (see Methods; shaded region in Figure 1A) to the high-coverage archaic humans. We calculated the sequence divergence—the number of pairwise differences normalized by the effective sequence length— between all haplotypes in the 1KG and the genotypes for the Altai Denisovan and the three high-coverage Neanderthal individuals (Figure S20-S22; Tables S34-S35). Haplotypes from the Americas exhibit a bimodal distribution of sequence divergence for affinities to the Altai Denisovan, which we do not observe for the African haplotypes (Figure 4A), as expected for an introgressed region. When comparing to all four high-coverage archaic genomes at the 72kb region (Figure 4B), there is a clear pattern of sequence divergence for the introgressed haplotypes found in the American super-population of the 1KG (AMR). Interestingly, Figure 4B shows that African haplotypes are closer in sequence divergence to the Altai Neanderthal than to the Altai Denisovan, but the value is not statistically significant (Dataset 1 [52]; [51]). The Altai Neanderthal itself is significantly more distant than expected from the Altai Denisovan (sequence divergence: 0.003782*, P-value:* 0.002, Figure S23, Table S36), and this larger than expected divergence explains why African haplotypes appear closer to the Altai Neanderthal in this region. We corroborate the pattern observed in Figure 4 using PCA to visualize the haplotype structure in this region (Figure S24).

**Figure 4.**
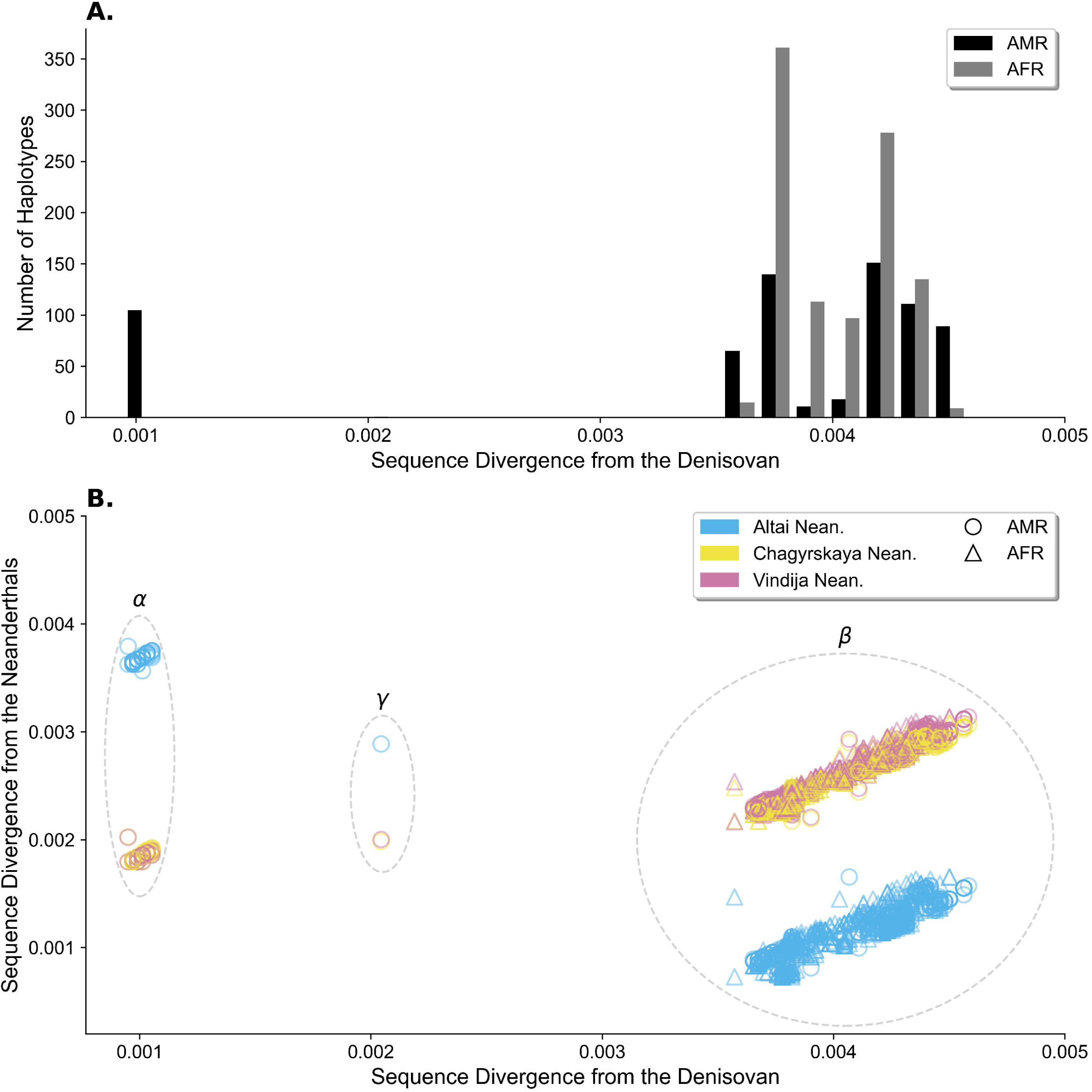
Haplotype divergence at the 72kb region in *MUC19*. (**A**) Distribution of haplotype divergence—number of pairwise differences between a modern human haplotype and an archaic genotype normalized by the effective sequence length—with respect to the Altai Denisovan for all individuals in the Admixed American (AMR, black bars) and African (AFR, gray bars) superpopulations. (**B**) Joint distribution of haplotype divergence from the Altai Denisovan (x-axis) and the Neanderthals (y-axis)—Altai Neanderthal in sky blue, Chagyrskaya Neanderthal in yellow, and Vindija Neanderthal in reddish purple—for all individuals in the AMR (circles) and AFR (triangles) superpopulations. The three grey ellipses (*α*, *β*, and *γ*) represent the three distinct haplotype groups segregating in the 1KG. The *α* ellipse represents the introgressed haplotypes which exhibit a low sequence divergence from the Altai Denisovan, a high sequence divergence from the Altai Neanderthal, and an intermediate sequence divergence—higher compared to the Altai Denisovan but lower compared to the Altai Neanderthal—with respect to the Chagyrskaya and Vindija Neanderthals. The *β* ellipse represents the non-introgresssed haplotypes which exhibit a high sequence divergence from the Altai Denisovan, a low sequence divergence from the Altai Neanderthal, and an intermediate sequence divergence—lower compared to the Altai Denisovan but higher compared to the Altai Neanderthal—with respect to the Chagyrskaya and Vindija Neanderthals. Note that the AMR haplotype within the *γ* ellipse is positioned at intermediate sequence divergence values with respect to the *α* and *β* ellipses, which represents one of seven recombinant haplotypes segregating in the 1KG (see Figure S44 in [51]).

Despite our *U_AFR,MXL,Denisovan_(1%, 30%, 100%)* and archaic SNP density results demonstrating that the introgressed haplotype at the 72kb region shares the most alleles with the Altai Denisovan (Figure 4B), we find that this region is not statistically significantly closer to the Altai Denisovan individual than expected from the genomic background of sequence divergence (sequence divergence: 0.00097*, P-value:* 0.237, Figure S25, Table S37). However, this is not unusual, given that the Altai Denisovan is not genetically closely related to Denisovan introgressed segments in modern humans (see Supplemental Section S5 in [51]), which might suggest that the Denisovan donor population of the 72kb region in *MUC19* is not closely related to the Altai Denisovan individual. Furthermore, the 72kb region is also not statistically significantly closer to Neanderthals than expected from the genomic background of sequence divergence (sequence divergence from the Altai Neanderthal: 0.003648*, P-value:* 0.995; Chagyrskaya Neanderthal: 0.001818, *P-value:* 0.811; Vindija Neanderthal: 0.001816, *P-value:* 0.806; Figure S25, Table S37).

As an additional approach, we used the *D+* statistic to assess which archaic human exhibits the most allele sharing with the introgressed haplotype at the 72kb region in *MUC19* [34, 35]. We performed *D+ (P1, P2; P3, Outgroup)* tests with the following configurations: the Yoruban population (YRI) as *P1*, the focal MXL individual (NA19664) with two copies of the introgressed haplotype with an affinity to the Altai Denisovan as *P2*, and one of the four high-coverage archaic genomes as *P3*; we use the EPO ancestral allele call from the six primate alignment as the Outgroup. We exclusively observe a positive and significant *D+* value (*D+*: 0.743, *P-value*: 1.386e-5; Figure S26; Table S38) when the Altai Denisovan is used as *P3* (the putative donor population). Conversely, when any of the three Neanderthals are used as *P3*, we observe non-significant *D+* values (*P3*: Altai Neanderthal, *D+*: -0.622, *P-value*: 0.999; *P3*: Chagyrskaya Neanderthal, *D+*: 0.175, *P-value*: 0.183; *P3*: Vindija Neanderthal, *D+*: 0.182, *P-value*: 0.174; Figure S26; Table S38). These *D+* suggest that the introgressed haplotype at the 72kb *MUC19* region shares more alleles with the Altai Denisovan, which is not observed with any of the three Neanderthals and provides evidence that the introgressed haplotype found in modern humans is *Denisovan-like*.

When we consider the 742kb region in MXL, we find that it is closest to the Chagyrskaya and Vindija Neanderthals, and significantly closer than expected from the genomic background (sequence divergence from the Chagyrskaya Neanderthal: 0.000661, *P-value*: 0.006; from the Vindija Neanderthal: 0.000656, *P-value*: 0.007; Figure S27-S30; Table S39-41; Dataset 2 [52]; [51). We also tested whether this region is statistically significantly closer to the Altai Denisovan than expected from the genomic background and found that this tract in MXL is also significantly closer than expected to the Altai Denisovan, albeit not as close when compared to the Chagyrskaya and Vindija Neanderthals (sequence divergence from the Altai Denisovan: 0.000806, *P-value*: 0.019; Figure S27-S30; Table S39-S41). We then performed *D+* analyses for the 742kb region with identical configurations as for the 72kb region and observe positive and significant *D+* values when *P3* is Chagyrskaya (*D+*: 0.381, *P-value*: 7.375e-6; Figure S31; Table S42), and Vindija Neanderthals (*D+*: 0.383, *P-value*: 7.505e-6; Figure S31; Table S42), but not when the Altai Neanderthal is *P3* (*D+*: 0.091, *P-value*: 1.442e-1; Figure S31; Table S42). *D+* is, however, significant when the Altai Denisovan is *P3* (*D+*: 0.377, *P-value*: 9.889e-8; Figure S31; Table S42). These *D+* results are consistent with our sequence divergence results, which indicate that the introgressed haplotype at the 742kb *MUC19* region has a high affinity for the Altai Denisovan and the two late Neanderthals, but not the Altai Neanderthal (Figures S20-S31; Tables S34-S42).

Given the high density of Denisovan-specific alleles (Figure S2; Table S4), the sequence divergence, and *D+* results for the 72kb and 742kb region, the most parsimonious explanation is that a Denisovan population could have introduced this haplotype into non-Africans. However, our 742kb results also suggest a Neanderthal population could have introduced the introgressed haplotype. This is further supported by the sequence divergence results at the 72kb region where late Neanderthals exhibit intermediate distance to the introgressed haplotype (Figure 4B), suggesting they harbor some of the Denisovan alleles.

### Neanderthals introduce *Denisovan-like* introgression into non-African modern humans

Based on sequence divergence, the Chagyrskaya and Vindija Neanderthals carry a 742kb haplotype that is most similar to the Altai Neanderthal, with the exception of the 72kb region. To understand why the Chagyrskaya and Vindija Neanderthals exhibit intermediate levels of sequence divergence with the introgressed haplotype present in MXL at the 72kb region in *MUC19* relative to the Altai Denisovan and Altai Neanderthal (see the *α* ellipse in Figure 4B), we computed the number of heterozygous sites for each archaic human. Because the Chagyrskaya and Vindija Neanderthals present intermediate sequence divergences, we expected these two individuals to have more heterozygosity than the Altai Neanderthal. At the 72kb region in *MUC19*, we observe that the Chagyrskaya and Vindija Neanderthals carry an elevated number of heterozygous sites (Chagyrskaya heterozygous sites: 168, *P-value:* 2.307e-4; Vindija heterozygous sites: 171, *P-value:* 3.282e-4; Figure 5A; Figure S32; Table S43) that is higher than those of the Altai Neanderthal (heterozygous sites: 1, *P-value*: 0.679; Figure5A; Figure S32; Table S43) and the Altai Denisovan (heterozygous sites: 6, *P-value*: 0.455; Figure5A; Figure S32; Table S43). The Chagyrskaya and Vindija Neanderthals carry a higher number of heterozygous sites than all African individuals (∼75, *P-value*: 0.424; Figure 5A; Figure S33; Table S44), and have a more similar pattern to non-African individuals carrying exactly one *Denisovan-like* haplotype (∼287, *P-value*: 3.157e-4; yellow X’s in Figure 5A; Figure S33; Table S44). This observation runs opposite to the genome-wide expectation for Neanderthals, as archaic humans have much lower heterozygosity than modern humans (genome-wide heterozygosity is ∼0.00014 - ∼0.00017 for the Neanderthals, ∼0.00019 for the Denisovan, and ∼0.001 for Africans modern humans; Figure S34; Table S45).

**Figure 5.**
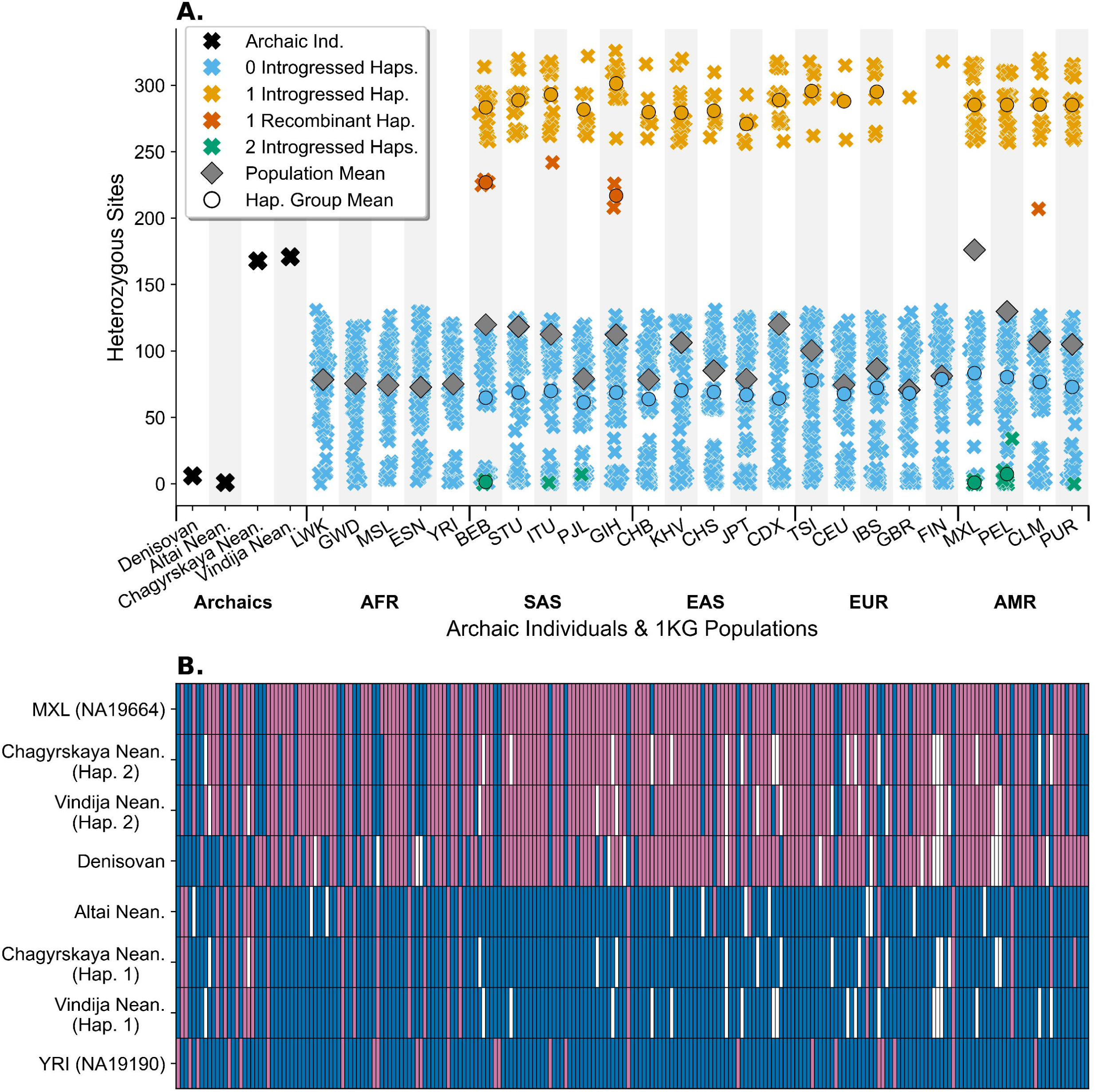
The high levels of heterozygosity in the Chagyrskaya and Vindija Neanderthals are explained by *Denisovan-like* ancestry at the 72kb region in *MUC19*. (**A**) Number of heterozygous sites at the 72kb region in *MUC19* per archaic individual (black X’s), 1KG individuals without the introgressed haplotype (sky blue X’s), 1KG individuals with exactly one copy of the introgressed haplotype (yellow X’s), 1KG individuals with a recombinant introgressed haplotype (vermillion X’s), and 1KG individuals with two copies of the introgressed haplotype (bluish green X’s). The average number of heterozygous sites stratified by population are denoted by the grey diamonds and the average number of heterozygous sites amongst individuals who carry exactly zero, one, and two introgressed haplotypes are denoted by sky blue, yellow, and bluish green circles respectively and are stratified by population. (**B**) Haplotype matrix of the 233 segregating sites (columns) amongst the focal MXL individual (NA19664) with two copies of the introgressed haplotype; the focal YRI individual (NA19190) without the introgressed haplotype; the Altai Denisovan; the Altai Neanderthal; and the two phased haplotypes for the Chagyrskaya and Vindija Neanderthals, respectively. Cells shaded blue denote the hg19 reference allele, cells shaded reddish purple denote the alternative allele, and cells shaded white represent sites that did not pass quality control in the given archaic individual. Note that the focal MXL and YRI individuals are homozygous for every position in the 72kb region in *MUC19* and that the heterozygous sites for the Altai Denisovan and Altai Neanderthal—six and one heterozygous sites respectively—are omitted.

Within modern humans, we find that individuals carrying exactly one *Denisovan-like* haplotype at the 72kb region harbor significantly more heterozygous sites at *MUC19* compared to the rest of their genome (average number of heterozygous sites: ∼287, *P-value*: 3.157e-4; Figure S33; Table S44), which surpasses the number of heterozygous sites at *MUC19* of any African individual (Figure 5A). Individuals carrying two *Denisovan-like* haplotypes harbor significantly fewer heterozygous sites than expected at *MUC19* relative to the rest of their genome (average number of heterozygous sites: ∼4, *P-value*: 6.945e-4; Figure S33; Table S44), while African individuals harbor the expected number of heterozygous sites (average number of heterozygous sites: ∼75, *P-value*: 0.424; Figure S33; Table S44). Given that the Chagyrskaya and Vindija Neanderthals and non-African individuals who harbor one copy of the *Denisovan-like* haplotype exhibit an excess of heterozygous sites at the 72kb region, we hypothesized that the Chagyrskaya and Vindija Neanderthals also harbor one *Denisovan-like* haplotype. This arrangement would explain the elevated number of heterozygous sites and the intermediary sequence divergences with respect to the introgressed haplotype.

To test this hypothesis, we first performed additional tests for gene flow between the archaic individuals using the *D+* statistic within the 72kb *MUC19* region that provided evidence that the Chagyrskaya and Vindija Neanderthals harbor one copy of the *Denisovan-like* haplotype. For these comparisons the Altai Neanderthal is *P1*, either the Chagyrskaya or Vindija Neanderthals are *P2*, and the Altai Denisovan is *P3*, we observe significant and positive *D+* values supporting gene flow between the Denisovan and the Chagyrskaya (*D+:* 0.783; *P-value:* 0.029) and Vindija (*D+:* 0.819; *P-value:* 0.018) Neanderthals (Figure S35; Table S46). To further investigate whether the Chagyrskaya and Vindija Neanderthals harbor one *Denisovan-like* haplotype in the 72kb region, we used `BEAGLE` to phase the 72kb region. As no phasing has been done for archaic humans, we tested the reliability of using the 1KG as a reference panel by constructing a synthetic 72kb region. We sampled one allele from the Altai Neanderthal and one allele from the Altai Denisovan at heterozygous sites in either the Chagyrskaya or Vindija Neanderthals. We found that we could phase the synthetic individual perfectly at this region (see Supplemental Sections S3-S4 in [51]). Encouraged by these results, we phased the Chagyrskaya and Vindija Neanderthals at the 72kb region, and confirmed they carry one haplotype that is similar to the Altai Neanderthal, and one haplotype that is similar to the *Denisovan-like* haplotype in MXL. Relative to the Altai Neanderthal, the Chagyrskaya *Neanderthal-like* haplotype exhibits 3.5 differences, and the Vindija exhibits 4 differences (Figure 5B; Table S47). Relative to the Altai Denisovan, the Chagyrskaya *Denisovan-like* haplotype exhibits 43 differences, and the Vindija haplotype exhibits 41 differences (Figure 5B; Table S47). As expected, the phased *Denisovan-like* haplotype in these two Neanderthals is closest to the *Denisovan-like* haplotype in MXL; the Chagyrskaya exhibits 5 differences, and the Vindija Neanderthal exhibits 4 differences (Figure 5B; Table S48). We show that, in the 72kb region, the introgressed haplotype in MXL is statistically significantly closer to the phased *Denisovan-like* haplotype present in Chagyrskaya and Vindija Neanderthals (sequence divergence from Chagyrskaya Neanderthal haplotype: 0.000104, *P-value:* 0.003; sequence divergence from Vindija Neanderthal haplotype: 0.000083, *P-value:* 0.002; Figure S36; Table S48; Dataset 3 [52]; [51]). Due to the potential introduction of biases when phasing ancient DNA data, to investigate if the Chagyrskaya and Vindija Neanderthals carry a *Denisovan-like* haplotype we developed an approach called Pseudo-Ancestry Painting (*PAP*, see [51]) to assign the two alleles at a heterozygous site to two source individuals. We found that using an MXL (NA19664) and a YRI (NA19190) individual as sources maximizes the number of heterozygous sites in the Chagyrskaya (*PAP* Score: 0.94, *P-value:* 3.683e-4) and Vindija (*PAP* Score: 0.929, *P-value:* 8.679e-05) Neanderthals (Figure S37; Table S49).

In sum, our analyses suggest that some non-Africans carry a mosaic region of archaic ancestry: a small *Denisovan-like* haplotype (72kb) embedded in a larger Neanderthal haplotype (742kb), that was inherited through Neanderthals, who themselves acquired Denisovan ancestry from an earlier introgression event (Figure S38). This is consistent with the literature, where Denisovan introgression into Neanderthals is rather common [*37, 38*]. Thus, we refer to the mosaic haplotype found in modern humans as the archaic haplotype.

## Discussion

The study of adaptive archaic introgression has illuminated candidate genomic regions that affect the health and overall fitness of global populations. In this study, we pinpointed several aspects of the gene *MUC19* that highlight its importance as a candidate to study adaptive introgression: one of the haplotypes that span this gene in modern humans is of archaic origin; modern humans inherited this haplotype from Neanderthals, who in turn inherited it from Denisovans; the haplotype introduced nine missense mutations that are at high frequency in both Indigenous and Admixed American populations; individuals with the archaic haplotype carry a massive coding VNTR expansion relative to the non-archaic haplotype, and their functional differences may help explain how mainland Indigenous Americans adapted to their environments, which remains under-explored. This study adds an example to the growing literature of natural selection acting on archaic alleles at coding sites, or possibly an example of natural selection acting on human VNTRs, a developing research frontier [see, 39].

A larger implication of our findings is that archaic ancestry could have been a useful source of standing genetic variation as the early Indigenous American populations adapted to new environments, with genes like *MUC19* and other mucins possibly mediating important fitness effects [*40*]. The variation in the *MUC19* coding VNTR in global populations dovetails with this idea and adds to a growing body of evidence for the important role of structural variants in human genomics and evolution [*41-42*]. In American populations, particular haplotypes carrying the most extreme copy numbers were selected and are now relatively frequent. This VNTR expansion effectively doubles the functional domain of this mucin, indicating an adaptive role driven by environmental pressures particular to the Americas. However, we cannot know whether the non-synonymous variants or the VNTR is driving natural selection as they are linked in haplotypes, and our evidence for positive selection is tied to SNP variation and not to the VNTR itself.

Another interesting aspect of *MUC19* is the evolutionary history of the introgressed region. Our observation of a 72kb Denisovan haplotype found in Neanderthals and non-African modern humans that is nested within a larger Neanderthal haplotype, suggests that the smaller Denisovan haplotype was first introgressed into Neanderthals, who later admixed with modern humans to introduce the full 742 kb haplotype. While the Altai Neanderthal does not harbor the Denisovan haplotype at the 72kb region, the other two chronologically younger Neanderthals (Chagyrskaya and Vindija) do. We phased these younger Neanderthals (see Supplementary Sections S3-S5 in [51]) and showed that they harbor exactly one Denisovan-like haplotype, which explains why they exhibit an excess of heterozygosity. The *Denisovan-like* haplotype in the younger Neanderthals is also statistically significantly closer to the archaic haplotype present in MXL (Figure S36; Table S48), providing additional evidence that modern humans obtained this haplotype through an interbreeding event with Neanderthals. Despite the introgressed archaic haplotype having an excessive amount of shared alleles with the Altai Denisovan at the 72kb region, the Altai Denisovan harbors several private mutations—14 and 6 mutations in the homozygous and heterozygous state respectively—that are absent across all 287 *Denisovan-like* haplotypes in the 1KG, suggesting that the introgressing Denisovan population may not be closely related to Altai Denisovan (see Supplemental Section S5; [51]). Indeed, the introgressed haplotype in the 72kb region is present at low frequencies in other non-African populations including Papuans—where the genome-wide Denisovan ancestry of Papuans has been estimated to originate from a population of Denisovans that was not closely related to the Altai Denisovan [*33*]. Finding two highly divergent haplotypes maintained in polymorphism in two Neanderthal populations, and finding the archaic haplotype at high frequencies in American populations but not at fixation may point to a balanced polymorphism [*45*]. More generally, the evolutionary history of this region suggests a complex history that involves recurrent introgression and natural selection, and it parallels complex introgression patterns from other regions of the genome [*46–48*].

Finally, we find a single San individual who carries the nine Denisovan missense variants in heterozygous form, uniquely among all African individuals considered here. The sequence divergence between this San haplotype and the archaic MXL haplotype at the 72kb region is high (0.001342), further supporting the origin of the archaic haplotype in non-Africans as introgressed. Khoe-San populations are estimated to have diverged from other African groups 120 thousand years ago [43]. Finding a divergent haplotype in the San is consistent with a previous study [44], as ∼1% of their ancestry can be attributed to lineages diverged from the main human lineage beyond 1 million years ago. We note that this San individual does not harbor an extended number of repeat copies of the VNTR (301 copies), which further supports the importance of the VNTR expansion in the Americas. Furthermore, we cannot determine if this variant found its way into the San through modern admixture of non-African ancestry into Sub-Saharan populations.

Perhaps the largest knowledge gap concerning why the archaic haplotype of *MUC19* would be under positive selection is its underlying function. Mucins are secreted glycoproteins responsible for the gel-like properties and the viscosity of the mucus [*49*]. Mucins are characterized by proline, threonine, and serine (PTS) tandem repeats, which in *MUC19* are structured into 30bp tandem repeats. The massive difference in copy numbers of the 30bp PTS tandem repeat domains carried by individuals harboring the *Human-like* and archaic haplotypes strongly suggests *MUC19* variants differ in function as a consequence of different molecular binding affinities between variants. This is the case in other mucins, such as *MUC7*, where variants carrying different numbers of PTS repeats exhibit different microbe-binding properties [*40*]. If the two variants of *MUC19* also have differential binding properties, this would lend support to why positive selection would increase the frequency of the archaic haplotype in American populations. Yet, there is limited medical literature associating variation in *MUC19* with human fitness. Further experimental validation of how VNTRs and the Denisovan-specific missense mutations affect MUC19 function is necessary to understand the effect the archaic haplotype may exert on the translated *MUC19* protein, and how it modifies its function during the formation of mucin polymers.

Methods developed in evolutionary biology can be useful for identifying candidate variants underlying biological functions. Future functional and evolutionary studies of the *MUC19* region will not only provide insight into specific mechanisms of how variation at this gene confers a selective advantage, but also specific evolutionary events that occurred in the history of humans. Beyond improving our understanding of how archaic variants facilitated adaptation in novel environments, our findings also highlight the importance of studying archaic introgression in understudied populations, such as admixed populations from the Americas [*50*]. Genetic variation in American populations is less well-characterized than other global populations; it is difficult to deconvolve Indigenous ancestries from European, African, and—to a lesser extent—South Asian ancestries, following 500 years of European colonization [*29*]. This knowledge gap is exacerbated by the high cost of performing genomic studies, building infrastructure, and generating scientific capacity in Latin America—but it is a worthwhile investment—as our study shows that leveraging these populations can lead to the identification of exciting candidate loci that can expand our understanding of adaptation from archaic standing variation.

## Acknowledgments

We would like to thank Alyssa Funk for contributing to the development of the *PBS* analysis, Ratchanon Pornmongkolsuk for early visualizations of global frequencies of *MUC19*, and Diego Ortega del Vecchyo and Paolo Provero for their insightful comments and discussion. We would also like to thank the Crawford and Ramachandran laboratories, especially Ria Vinod, Julian Stamp, Chibuikem Nwizu, Cole Williams, and Leah Darwin for their invaluable feedback and support throughout the duration of this project. Part of this research was conducted using computational resources and services at the Center for Computation and Visualization, Brown University.

## Funding

The Leakey Foundation (to FAV).

National Institutes of Health (1R35GM128946-01 to EHS).

Alfred P. Sloan Foundation (to EHS).

Blavatnik Family Graduate Fellowship in Biology and Medicine (to DP).

Brown University Predoctoral Training Program in Biological Data Science (NIH T32 GM128596 to DP and ETC).

National Institutes of Health (R35GM142978 to PM).

Burroughs Wellcome Fund (Career Award at the Scientific Interface to PM).

National Institutes of Health (R01NS122766 to PNV).

Human Frontier Science Program (to EHS, FJ, and MAA).

## Author contributions

Conceptualization: FAV, DP, EHS

Formal analysis: FAV, DP, EJK, VAG, KEW, VVI, RZ, DM, PM, FJ, PNV, MAA, EHS

Supervision: DM, PM, FJ, PNV, MAA, EHS

Writing – original draft: FAV, DP, EHS

Writing – review & editing: EJK, VAG, KEW, VVI, RZ, DM, PM, FJ, PNV, MAA

## Data and materials availability

The 1,000 Genomes Project Phase III, Simons Genome Diversity Project, high-coverage archaic genomes, Human Pangenome Reference Consortium, and Human Genome Structural Variant Consortium datasets are all publicly available. Ancient American genomes are available after signing data agreements from the original publications. All software used in this study is publicly available, and all statistical tests are described in the methods. All the information needed to reproduce the results in this study is described in the methods and supplemental methods. Additionally, the original code and final results can be found at: https://github.com/David-Peede/MUC19; intermediary files used to produce our final results can be found at: https://doi.org/10.5061/dryad.z612jm6pj; and the introgressed tracts, repeat information, phased late Neanderthal haplotypes, and Datasets S1-S4 can be found at: https://doi.org/10.5281/zenodo.15042423.

## Supplementary Materials

Materials and Methods

Supplementary Text

Figs. S1 to S60

Tables S1 to S67

Datasets S1 to S4

References (*52-102*)

