## Supplemental Figures for "The *MUC19* Gene: An Evolutionary History of Recurrent Introgression and Natural Selection"

Science
AAAS

### Supplementary Materials for

#### The MUC19 Gene: An Evolutionary History of Recurrent Introgression and Natural Selection

Authors: Fernando A. Villanea, David Peede, Eli J. Kaufman, Valeria Añorve-Garibay, Elizabeth T. Chevy, Viridiana Villa-Islas, Kelsey E. Witt, Roberta Zeloni, Davide Marnetto, Priya Moorjani, Flora Jay, Paul N. Valdmanis, María C. Ávila-Arcos, Emilia Huerta-Sánchez.

**The PDF file includes:**

Materials and Methods
Supplementary Text
Figs. S1 to S60
Tables S1 to S67
References (54-99)

**Other Supplementary Materials for this manuscript include the following:**

Data S1 to S4

#### Materials and Methods

Data Processing

*Modern Human Data*

Sequence data for the *MUC19* locus were obtained from a publicly available global reference panel, the 1,000 Genomes Project Phase III (1KG), which contains a diverse set of individuals from multiple populations [54]. The autosomal variant sites from the integrated callset VCF files for the 1KG were downloaded from http://ftp.1000genomes.ebi.ac.uk/vol1/ftp/release/20130502 and the local ancestry calls for admixed American individuals [27] were downloaded from https://personal.broadinstitute.org/armartin/tgp_admixture. Data for *MUC19* in the Papuan and present day Indigenous American individuals was obtained from the Simons Genome Diversity Project (SGDP) [55; 56]. The autosomal variant sites VCF files for the SGDP were downloaded from https://sharehost.hms.harvard.edu/genetics/reich_lab/sgdp/phased_data2021. Both the 1KG and SGDP datasets were filtered to remove multi-allelic and structural variant sites. For the *iHS* analyses conducted using the 1KG dataset, we removed additional sites that did not have an ancestral allele call and were no longer bi-allelic after considering the ancestral allele call, where we annotated the dataset using the  ancestral allele calls in fasta format for the hg19 assembly using the Enredo, Pecan, Ortheus (EPO) pipeline, which was download from http://ftp.ensembl.org/pub/release-74/fasta/ancestral_alleles [57; 58]. Given that the modern human genotypes were imputed and only include information for variable sites, any site that was not removed during the filtering process was assumed to be homozygous reference as was done in Huerta-Sanchez et al. [18]. As the ACB and ASW admixed populations have a high proportion of African ancestry, individuals from these populations were removed and not considered in any analysis. Note that the analyses in the section "Copy number polymorphism of a 30bp tandem repeat motif between the *Human-like* and a *archaic* haplotypes" were conducted on the 1KG data aligned to the hg38 reference assembly, while all other analyses were completed using the hg19 reference which was soft masked for repetitive regions. Data aligned to hg38 was downloaded from the Human Pangenome Reference Consortium (HPRC) [59]: https://projects.ensembl.org/hprc/ and the Human Genome Structural Variation Consortium (HGSV) [42]: https://www.internationalgenome.org/data-portal/data-collection/hgsvc2. To ensure that the 135 Denisovan-specific alleles at *MUC19* were not artefacts arising from short-read sequencing technology, we used two sets of high-coverage long-read genomes as an orthogonal dataset to confirm the presence of all 135 alleles (Dataset 4 [52]). The first set consisted of three genomes from the HGSVC (individual-population: HG00864-CDX, HG03009-BEB, NA20847-GIH). The second set comprised an additional three genomes (individual-population: HG01122-CLM, HG02252-PEL, HG02262-PEL) from the 1000 Genomes Project Oxford Nanopore Technologies Sequencing Consortium (1KG-ONT; https://s3.amazonaws.com/1000g-ont/index.html?prefix=ALIGNMENT_AND_ASSEMBLY_DATA/FIRST_100/) [60]. We converted the genomic coordinates of the 135 Denisovan-specific alleles from the hg19 to hg38 assembly using the UCSC liftover tool (https://genome.ucsc.edu/cgi-bin/hgLiftOver) and confirmed that the reference alleles are identical between the two assemblies. Subsequently, we verified the presence of all 135 alleles in the three HGSVC individuals using variants called with the Phased Assembly Variant caller [42]. For the three individuals from the 1KG-ONT dataset, we confirmed the presence of these Denisovan alleles using variant calls generated by both Clair3 and PEPPER-Margin-DeepVariant callers [61]. These six individuals were chosen because each harbors at least one *Denisovan-like* haplotype at the 72kb region.

*Archaic Human Data*

The autosomal all-sites VCF files and the BED files consisting of the suggested general filtering best practices for the four high-coverage archaic genomes were downloaded from https://www.eva.mpg.de/genetics/genome-projects. The autosomal VCF files for the archaic genomes were initially filtered with their respective BED files to exclude regions of the genome that are prone to alignment errors as was done in the original [62; 63]. The initially filtered archaic VCF files were then merged using `BCFtools v1.16`, and after the initial merging the resulting VCF files were filtered to only include sites that were mono-allelic or bi-allelic where at least one archaic had a MQ >= 25 and GQ >= 40—archaics that did not meet this threshold were coded as missing data. Additionally for our *D*+ analysis, we generated another merged archaic dataset by adding the additional requirement that at a given site we could determine the ancestral and derived state as defined by the hg19 ancestral sequence.

*Combined Data*

The autosomal VCF files for each modern human dataset (i.e., 1KG and SGDP) and each filtered archaic genomes were initially merged using `BCFtools v1.16`, after the initial merging the resulting VCF files were filtered to only include sites with mono-allelic or bi-allelic SNPs and where the respective archaic had a MQ >= 25 and GQ >= 40. For *D*+ analyses we generated another combined dataset per archaic, with the additional requirement that the given site must have an ancestral allele call present as defined by the hg19 ancestral sequence. Additionally to assign sites into the different SNP set partitions (see the Archaic SNP Density section) and phasing analyses we merged the 1KG dataset with all four filtered archaic genomes archaics with the same filtering scheme for the merged datasets with the single archaic. We retained sites that had at least one archaic pass the filtering criteria and for any archaic that did not meet the filtering criteria was coded as missing data. Since the modern human genotypes were imputed and only include information for variable sites—unlike the archaic data which contains information for all sites—any site that was originally absent in the 1KG or SGDP datasets but present in archaic data we assumed to be homozygous for the reference allele in the modern humans [18]. After each of the combined datasets were curated, we annotated coding sites for NCBI RefSeq genes [64] using the NCBI RefSeq Select transcripts (downloaded from https://hgdownload.cse.ucsc.edu/goldenpath/hg19/database/ncbiRefSeqSelect.txt.gz) and

SnpEff v5.1 [65]. Lastly, for better computational efficiency when performing analyses all of the final VCF files were converted to Zarr arrays using scikit-allele v1.3.5 [94] (10.5281/zenodo.597309).

*Pre-Contact Indigenous American Genomes*

Genomic data for *MUC19* in ancient individuals was generated by combining high coverage (>1X) pre-European contact genomes from the literature, including nine individuals from California, one from Ontario [66], four from Peru [67] four from Patagonia [68], one from Alaska [69], one from Montana [70], and three from Central Mexico [71]. Sequence reads were downloaded in FASTQ format and aligned to hg19 in BAM format using bwa v7.17 [72]. Reads were then sorted, duplicates were removed, and all non-autosomal chromosomes were removed using SAMtools v1.9 [73]. Using ANGSD v0.92 we further filtered out reads that had a quality score less than 30 and then determined the read depth of the alleles present at the Denisovan-specific coding sites. Then, for each ancient individual we determined the genotype at each of the Denisovan-specific coding sites which we had sequencing information for by first considering any allele that had a read depth of two or greater, and then ensured that site was mono-allelic or bi-allelic for only the hg19 reference and/or Denisovan-specific alternative allele.

Identification of the *MUC19* Introgressed Region

To identify the genomic coordinates of the introgressed region in *MUC19*, we inferred introgressed tracts using hmmix [26] for chromosome 12. Specifically, we inferred introgressed tracts using the -haploid option which generates two sets of inferred tracts per individual—i.e., one for each haploid genome—and only retained inferred archaic tracts that had a posterior probability greater than or equal to 0.8, and that overlapped with at least one base pair of the *MUC19* NCBI RefSeq coordinates for the hg19 assembly (Chr12:40787196-40964559). As we were only interested in the tracts that overlap *MUC19*, if an individual's haplotype had two inferred tracts overlapping *MUC19* we stitched them together following the approach implemented in [74] by taking the union of the two respective tracts—i.e., the minimum of the two start positions and the maximum of the two end positions. We then performed a Proportions *Z*-test to determine if the proportion of introgressed tracts in the admixed American populations is greater than in non-admixed American populations using the statsmodels.stats.proportion.proportions_ztest function implemented in statsmodels v0.13.2 [75], and a Fisher's Exact Test to assess if introgressed tracts are overrepresented in admixed American populations than non-admixed American populations using the scipy.stats.fisher_exact as implemented in scipy v1.7.2 [81]—for both statistical tests African populations were not included among non-admixed American populations and *P-values* less than 0.05 were considered statistically significant. Lastly, we identified two distinct regions, the 742kb region containing the longest archaic tract in any MXL individual (Chr12:40272001-41014000, in NA19725), and a focal 72kb region (Chr12:40759001-40831000) which has the highest density of inferred introgressed tracts amongst non-African populations in the 1KG that is larger than 40kb, which is the length needed to confidently differentiate between introgression and incomplete lineage sorting [18]. It should be noted that in Figure 1A we omitted the 2.613 Mb tract in the PUR individual HG01108 for visual clarity, but the version of this plot including this tract along with population specific plots can be viewed in Figures S39-S43.

Archaic SNP Density

Throughout this study we compute different subsets of archaic specific alleles found in non-African populations. For a SNP to first be considered an archaic allele, we require that an allele must be rare in the African superpopulation (i.e., at a frequency less than 0.01) and at a frequency greater than 0.01 in the respective non-African population as was done in [76]. For the SNP to be considered Denisovan-specific we further required the archaic allele to be fixed in the sequenced Denisovan and not fixed in any of the three high-coverage Neanderthals. Similarly, for a SNP to be considered Neanderthal-specific we further required the archaic allele to be fixed in at least one Neanderthal and not fixed in the sequenced Denisovan. For an SNP to be considered as a shared archaic allele we further required the archaic allele to be fixed in the sequenced Denisovan and in at least one Neanderthal. For the SNPs that were not identified as archaic SNPs—i.e., the union of the Denisovan-specific, Neanderthal-specific, and shared archaic SNPs—we classified alleles segregating in the 1KG and absent in all archaic individuals as Human-specific. Lastly, non-Archaic SNPs that are segregating in at least one archaic and are also segregating in the 1KG are considered to be shared hominin alleles. To determine if non-African populations at the 742kb longest introgressed tract region and at the core 72kb *MUC19* region harbor more archaic SNPs than expected we computed the number of Denisovan-specific and Neanderthal-specific SNPs for each region. To assess if our 742kb and 72kb *MUC19* regions have a higher archaic SNP density than expected, we compared the observed archaic SNP density to a distribution of archaic SNP densities from the genomic background of 742kb and 72kb non-overlapping windows with comparable effective sequence length density—i.e., within one standard deviation from the mean for each windowed effective sequence length distribution. To calculate *P-values,* we determined the proportion of windows from the genomic background with an archaic-specific SNP density greater than or equal to what we observed at the 742kb and 72kb *MUC19* regions, respectively, where a *P-value* less than 0.05 is considered statistically significant.

Positive Selection

*Population Branch Statistic*

We utilized the Population Branch Statistic (*PBS*) to assess if the archaic haplotype has been subjected to positive selection in Admixed American populations. *PBS* uses the logarithmic transformation of pairwise estimates of $F_{ST}$ to measure the branch length in the target population since its divergence from the two control populations [77]. We chose MXL as the target population and CEU and CHB as our control populations. To account for differences in sample sizes between populations we used Hudson's estimator of $F_{ST}$ as it has been shown to not only be a conservative estimator, but also robust to differences in sample sizes [78]. Note that since $F_{ST}$ and *PBS* both represent branch lengths on unrooted trees, in all *PBS* computations we set negative $F_{ST}$ and *PBS* values to zero to be conservative. To assess if there is evidence of positive selection at the 742kb longest introgressed tract region and the 72kb densest introgressed tract region we computed $PBS_{MXL:CHB:CEU}$ at each region and then to compute *P-values* we compared each observed $PBS_{MXL:CHB:CEU}$ value to a distribution of $PBS_{MXL:CHB:CEU}$ values from the genomic background of 742kb and 72kb non-overlapping windows with comparable effective sequence length—i.e., within one standard deviation from the mean for each windowed effective sequence length distribution—and determined the proportion of windows from the genomic background with a per-region $PBS_{MXL:CHB:CEU}$ value greater than or equal to what we observed at the 742kb and 72kb region, respectively, where a *P-value* less than 0.05 is considered statistically significant. Additionally, we computed $PBS_{MXL:CHB:CEU}$ for every SNP in the genome and identified outlier SNPs in the 742kb region by setting the significance threshold at the genome-wide 99.95th $PBS_{MXL:CHB:CEU}$ percentile—for information on how we assessed significance for SNPs within the 742kb region see Supplemental Section S1.

*Integrated Haplotype Score*

To corroborate our signals of selection we additionally performed a haplotype-based test for selection by computing integrated haplotype scores (*iHS*) [79] for all populations in the 1KG. *iHS* measures the decay in linkage disequilibrium from a core SNP due to new mutations and recombination events, as recent positive selection is expected to result in haplotypes that are long, frequent, and have a high haplotype homozygosity in a population. Normalized $|iHS| > 2$ reflect that the haplotype is longer than expected under neutrality—and is commonly considered the threshold for evidence of positive selection at a locus [79; 29]—with extreme positive and negative values indicating that the derived and ancestral haplotypes are unusually long, respectively. To compute *iHS* we used the 1KG dataset with ancestral allele information. Using `selscan v2.0.0` [29] and the recombination maps from [81] we first computed the unstandardized *iHS* for every SNP with a minor allele frequency > 0.05 per 1KG population and then normalized *iHS* by derived frequency bins as described in [79]. As there is no formal way to assess significance for normalized *iHS,* to determine if the focal 742kb longest introgressed tract region and the 72kb densest introgressed tract region showed signals consistent with positive selection we assessed if these regions harbored clusters of extreme *iHS* (i.e., normalized $|iHS| > 2$) [79; 80]. Specifically, for every 1KG population we binned the genome into 742kb and 72kb non-overlapping windows and computed the proportion of SNPs with normalized $|iHS| > 2$ for windows with more than 10

SNPs to build a genome-wide distribution of the proportion of SNPs with extreme *iHS*. We then assessed if the focal 742kb and 72kb regions fall within the top 1% of windows with the highest proprtions of extreme *iHS*. Additionally, we repeated these analyses for all archaic SNPs, which are described in Supplemental Section S2.

*U-Statistics*

To complement our tests for positive selection we also performed explicit tests for adaptive introgression using the $U_{A,B,C}(w, x, y)$ statistic [24]. If a genomic region is adaptively introgressed we would expect there to be many sites within that region where the archaic allele is at high frequency in the non-African population that has experienced adaptive introgression and absent or rare amongst African populations. The $U_{A,B,C}(w, x, y)$ statistic quantifies the number of sites in a given region where the archaic individual (*C)* has an allele frequency of *y%*, that the allele is at a frequency less than *w%* in a control population (*A*), and that the allele is segregating at a frequency greater than *x%* in a target population (*B*). For this study we used $U_{AFR,B,Denisovan}(1\%, 30\%, 100\%)$ which quantifies the number of sites where the Denisovan allele is found in the homozygous state that are segregating at a frequency less than 0.01 in the African super population and is segregating at a frequency greater than 0.3 in a given non-African population (*B*). To assess if the entire *MUC19* gene exhibits signatures of adaptive introgression we computed $U_{AFR,B,Denisovan}(1\%, 30\%, 100\%)$ for every NCBI RefSeq gene that has at least one segregating site amongst the Denisovan and the 1KG, for all non-African populations (*B*). Given the variance in the effective sequence lengths and number of segregating sites amongst NCBI RefSeq genes, we decided that it was not feasible to assess statistical significance, but it should be noted that *MUC19* in MXL is the maximum $U_{AFR,B,Denisovan}(1\%, 30\%, 100\%)$ value for all NCBI RefSeq genes amongst all non-African populations (*B*), as no other gene in any non-African population exhibits such a large value of $U_{AFR,B,Denisovan}(1\%, 30\%, 100\%)$ (see Figure 1B in the main text). In order to assess statistical significance, we computed $U_{AFR,B,Denisovan}(1\%, 30\%, 100\%)$ for all non-African populations (*B*) for both the 742kb longest introgressed tract in MXL region and the focal 72kb region and to test if the observed $U_{AFR,B,Denisovan}(1\%, 30\%, 100\%)$ is larger than expected, we compared each of our observed values to a distribution of $U_{AFR,B,Denisovan}(1\%, 30\%, 100\%)$ from the genomic background of 742kb and 72kb non-overlapping windows with comparable effective sequence length—i.e., within one standard deviation from the mean for each windowed effective sequence length distribution. To calculate *P-values* for each non-African population (*B*)*,* we determined the proportion of windows from the genomic background with an $U_{AFR,B,Denisovan}(1\%, 30\%, 100\%)$ greater than or equal to what we observed at the 742kb and 72kb *MUC19* regions, respectively, where a *P-value* less than 0.05 is considered statistically significant.

*Q-Statistics*

To provide an orthogonal line of evidence for adaptive introgression specific to MXL, we computed the $Q95_{A,B,C}(w, y)$ statistic [24]. Under a scenario of adaptive introgression, one would expect that the introgressed alleles in the recipient population are at high frequencies. The $Q95_{A,B,C}(w, y)$ summarizes the site frequency spectrum in the target population (*B*), conditioned on the allele being present at a frequency of at least *y*% in the archaic population (*C*) and less than *w*% in a control population (*A*), which is accomplished by quantifying the 95th percentile of this conditional site frequency spectrum in the target population (*B*). In this study, we computed the $Q95_{AFR,B,Denisovan}(1\%, 100\%)$ statistic, which measures the 95th percentile of allele frequencies in a given non-African population (*B*) for alleles found in a homozygous state in Denisovans and that are segregating at a frequency less than 1% in the African super population. To provide evidence that there are signals consistent with adaptive introgression exclusive to MXL we computed $Q95_{AFR,B,Denisovan}(1\%, 100\%)$ for all non-African populations (*B*) for both the 742kb longest introgressed tract in MXL region and the focal 72kb region, as well as in non-overlapping windows with comparable effective sequence length—i.e., within one standard deviation from the mean for each windowed effective sequence length distribution. Beyond summarizing how high Denisovan allele frequencies are, we also assessed the number of Denisovan alleles at high frequencies. For this, we analyzed the joint distribution of $Q95_{AFR,B,Denisovan}(1\%, 100\%)$ and $U_{AFR,B,Denisovan}(1\%, 30\%, 100\%)$ statistics for each non-African population (B). We compared the observed values in the 742kb and 72kb *MUC19* regions to the joint distribution of these statistics from the genomic background of non-overlapping windows. Although there is no formal statistical test for significance using this joint distribution, it should be noted that for both the 742kb longest introgressed tract in MXL and the focal 72kb region, we observed $Q95_{AFR,MXL,Denisovan}(1\%, 100\%) = 0.305$ and $U_{AFR,MXL,Denisovan}(1\%, 30\%, 100\%) = 136$. These values are higher than any other non-African population at both focal *MUC19* regions, which is consistent with evidence for adaptive introgression specific to MXL in these regions.

#### Sequence Divergence

To assess the extent of divergence between *MUC19* haplotypes harbored by the various individuals in this study, we calculated sequence divergence which corresponds to number of pairwise differences between chromosomes normalized by the effective sequence length—i.e., the total number of sites that passed quality control. We use the term haplotype to refer to a single chromosome from a phased modern human individual and genotypes to refer to the two chromosomes of an archaic individual that are unphased. For information on assessing sequence divergence using the phased archaic data at the focal 72kb region see Supplemental Section S5.

##### *Identifying the Donor of the Longest Introgressed Tract found in MXL*

To determine the most likely archaic source of the 742kb longest introgressed tract (Chr12:40272001-41014000) found in an MXL individual (i.e., NA19725), inferred using `hmmix` [26] we computed the sequence divergence between each of the NA19725's chromosomes and the genotypes of the four archaic individuals. To identify the archaic donor for the 742kb longest introgressed tract, we compared the observed sequence divergence to a distribution of sequence divergence from the genomic background of 742kb non-overlapping windows with comparable effective sequence length density—i.e., within one standard deviation from the mean for 742kb windows effective sequence length distribution. To calculate *P-values,* we determined the proportion of windows from the genomic background with a sequence divergence less than or equal to what we observed at the 742kb region. After correcting for two multiple comparisons—i.e., one per haplotype—using the Bonferroni correction, a *P-value* less than 0.025 is considered significant. To assess differences in the frequency of the 742kb introgressed haplotype among 1KG populations we then computed the observed sequence divergence and *P-values* for all haplotypes in the 1KG. Given that the 742kb introgressed tract found in NA19725 was closest to the two late Neanderthals (i.e., Chagyrskaya and Vindija Neanderthals) we classified a 1KG haplotype as introgressed at the 742kb region if it was significantly closer than expected to either of the two late Neanderthals. We then performed a Proportions *Z*-test to determine if the proportion of introgressed haplotypes at the 742kb region in the admixed American populations is greater than in non-admixed American populations using the `statsmodels.stats.proportion.proportions_ztest` function implemented in `statsmodels v0.13.2` [75], and a Fisher's Exact Test to assess if introgressed haplotypes at the 742kb region are overrepresented in admixed American populations than non-admixed American populations using the `scipy.stats.fisher_exact` function implemented in `scipy v1.7.2 [81]`—for both statistical tests African populations were not included among non-admixed American populations and *P-values* less than 0.05 were considered statistically significant.

*Modern Human Haplotype-Archaic Human Sequence Divergence at the Focal 72kb Region*

To determine the haplotype identity—i.e., the most likely donor—for the 72kb *MUC19* region in 1KG individuals, we calculated the sequence divergence for all pairwise possibilities between each 1KG haplotype and the genotypes of the four archaic individuals. We then characterized a 1KG haplotype as being *Denisovan-like* at the 72kb region if the sequence divergence to the sequenced Denisovan was less than 0.00144524 (or 70 pairwise differences between a single 1KG chromosome and the two Denisovan chromosomes), corresponding to the lone black bar of the bimodal distribution in Figure 2A and the $\alpha$ ellipse in Figure 2B**.** Seven 1KG haplotypes exhibited intermediary sequence divergence levels with respect to the sequenced Denisovan between 0.002023 and 0.002044 and were determined to be recombinant haplotypes (see Figure S44 and the $\gamma$ ellipse in Figure 2B) and all 1KG haplotypes with a sequence divergence larger than 0.002044 include all African chromosomes and were considered to be *Human-like* haplotypes (see the $\beta$ ellipse in Figure 2B). We then performed a Proportions *Z*-test to determine if the proportion of *Denisovan-like* haplotypes in the admixed American populations is greater than in non-admixed American populations using the `statsmodels.stats.proportion.proportions_ztest` function implemented in `statsmodels v0.13.2` [75], and a Fisher's Exact Test to assess if *Denisovan-like* haplotypes are overrepresented in admixed American populations than non-admixed American populations using the `scipy.stats.fisher_exact` function implemented in `scipy v1.7.2` [81]—for both statistical tests African populations were not included among non-admixed American populations and *P-values* less than 0.05 were considered statistically significant.

*Altai Denisovan Sequence Divergence from the Altai Neanderthal and Africans at the Focal 72kb Region*

To test if African haplotypes in the 1KG project appear to have an affinity to the Altai Neanderthal because they both have an elevated sequence divergence from the Altai Denisovan, we calculated sequence divergence as the average number of pairwise differences between all African haplotypes and the Altai Denisovan normalized by the effective sequence length. To assess if Africans have an elevated sequence divergence from the Altai Denisovan at the 72kb region, we computed *P-values* as the proportion of windows from the genomic background with a sequence divergence greater than or equal to what we observed at the 72kb region. Similarly, we also computed sequence divergence between the Altai Neanderthal and the Altai Denisovan as the average number of pairwise differences between the two archaic genotypes normalized by the effective sequence length. To determine if the sequence divergence between the Altai Neanderthal and Denisovan is elevated at the 72kb region, we calculated *P-values* as the proportion of windows from the genomic background with a sequence divergence greater than or equal to what we observed at the 72kb region. For both Africans and the Altai Neanderthal, we considered a *P-value* less than 0.05 as statistically significant.

#### Site Patterns Tests of Introgression

To further corroborate claims of introgression based on our sequence divergence results we used the *D+* statistic to formally test hypotheses about local introgression [34;35]. The *D+* statistic utilizes observed site patterns from three populations and an outgroup—Newick format: *(((P1, P2), P3), O)*; site pattern format: *(P1's allelic state, P2's allelic state, P3's allelic state, O's allelic state)*—as a proxy for gene tree frequencies, where *P1* represents a population assumed to have not received gene flow from *P3, P2* represents a potential recipient population of introgression from the *P3* donor population, and an outgroup is used to polarize the ancestral states. *D+* specifically utilizes four site patterns: *ABBA, BABA, BAAA,* and *ABAA* (where an *A* denotes the ancestral allele and *B* denotes the derived allele) to test for asymmetries in site pattern frequencies. Under a scenario of no gene flow the *D+* statistic is expected to be zero, a significant and positive *D+* value indicates that *P2* and *P3* share more derived and ancestral alleles than expected, which may be explained by introgression. For all *D+* tests we used the ancestral allele calls from the six primate alignment inferred from EPO pipeline to polarize ancestral states [56-58]. Since the *D+* statistic is normally distributed, to assess if observed *D+* values significantly differed from zero at the 742kb longest introgressed tract region and at the focal 72kb *MUC19* region, for each test we constructed *Z*-distributions of *D*+ values from 742kb and 72kb non-overlapping windows with comparable effective sequence length and computed *P-values* using the `scipy.stats.norm.sf` function implemented in `scipy v1.7.2 [81]` and a *P-value* less than 0.05 is considered statistically significant.

*Patterns of Allele Sharing Between the Introgressed Haplotype in MXL and the Archaics*

To further investigate the signals of introgression at *MUC19* we used *D*+ as a complementary approach to our sequence divergence results. While using all sites to compute sequence divergence uses more information, this extra information may add noise due to new mutations arising in both the recipient and donor population since the introgression event, thus we calculated *D*+ for the 742kb longest introgressed tract region in MXL and the focal 72kb *MUC19* region to further identify the most likely donor of the introgressed haplotype in MXL. To test hypotheses about introgression at each region we calculated *D*+ for all combinations of *P1 = {YRI}, P2 = {NA19664}* who is an MXL individual who harbors two copies of the introgressed haplotype, and *P3 = {Altai Denisovan, Altai Neanderthal, Chagyrskaya Neanderthal, Vindija Neanderthal}* at both focal regions, and compared the observed *D*+ values to the genomic background as previously described above.

*Gene Flow between Denisovans and Late Neanderthals*

To assess the possibility of gene flow between Denisovans and the late Neanderthals at the 72kb *MUC19* region we performed *D*+ tests of introgression among the archaic individuals. To test this hypothesis we computed *D*+ for all combinations of *P1 = {Altai Neanderthal}, P2 = {Chagyrskaya Neanderthal, Vindija Neanderthal},* and *P3 = {Altai Denisovan}* at the focal 72kb region, and compared the observed *D*+ values to the genomic background as previously described above.

#### Heterozygosity in the Archaics and 1KG

To understand if the 72kb *MUC19* region is an outlier for heterozygosity in the four archaic genomes we counted the number of heterozygous sites for the 72kb *MUC19* region and compared the observed number of heterozygous sites to a distribution of heterozygous site counts from the genomic background of 72kb windows with comparable effective sequence length density—i.e., within one standard deviation from the mean of the windowed effective sequence length distribution. To calculate *P-values*, we determined the proportion of windows from the genomic background where the number of heterozygous sites is greater than or equal to what we observed at the 72kb *MUC19* region, where a *P-value* less than 0.05 is considered statistically significant. Additionally, we were interested in understanding if African individuals and individuals carrying the *Denisovan-like* haplotype at the 72kb region are also outliers for heterozygosity in our focal

72kb *MUC19* region. To do so we computed the average number of heterozygous sites amongst all African individuals (n = 504), heterozygous individuals (n = 255) who carry exactly one copy of the *Denisovan-like* haplotype at the 72kb region, and homozygous individuals (n = 16) who carry two copies of the *Denisovan-like* haplotype at the 72kb region, and compared our observed values to distributions of average heterozygous site counts from the genomic background of 72kb windows with comparable effective sequence length density—i.e., within one standard deviation from the mean of the windowed effective sequence length distribution. To calculate *P-values* for the African and heterozygous individuals, we determine the proportion of windows from the genomic background with an average number of heterozygous sites greater than or equal to what is observed at the 72kb *MUC19* region, and to calculate *P-values* for homozygous individuals we determine the proportion of windows from the genomic background with an average number of heterozygous sites less than or equal to what is observed at the 72kb *MUC19* region, where a *P-value* less than 0.05 is considered statistically significant. Lastly, we counted the number of heterozygous sites for all 1KG and archaic individuals across all autosomes to establish our genome-wide expectation for patterns of heterozygosity.

Phasing the Archaic Genomes at *MUC19*

*Benchmarking Phasing with a Synthetic Neanderthal*
To evaluate the feasibility of phasing a late Neanderthal at the core 72kb region, we first generated a Synthetic Neanderthal from sampling one allele from the Altai Denisovan and the other allele from the Altai Neanderthal using the 1KG and all archaics combined dataset. This Synthetic Neanderthal harbors 155 heterozygous sites, representing the intersection of sites where the Altai Denisovan and Altai Neanderthal are fixed for different allelic states and those that are heterozygous in at least one of the late Neanderthals—i.e., the Chagyrskaya and Vindija Neanderthals. We then used `BEAGLE v5.4` [2] to phase this Synthetic Neanderthal at the 72kb region, using 1KG individuals as the reference panel and including the Synthetic Neanderthal, Altai Denisovan, and Altai Neanderthal in the phasing panel. Our analysis demonstrated that the Synthetic Neanderthal could be perfectly phased at the core 72kb region, which provided the motivation to phase each of the late Neanderthals at this region (see Supplemental Section S3 for a detailed discussion).

*Phasing the Late Neanderthals at the 72kb MUC19 region*
Building on our benchmarking results, which demonstrated that we could perfectly phase a Synthetic Neanderthal at the core 72kb region (see Supplemental Section S3), we implemented a two-step approach to phase the late Neanderthals—i.e., the Chagyrskaya and Vindija Neanderthals—at the 72kb region using the combined dataset of 1KG and all archaic genomes. In the first step, we statistically phased the heterozygous sites for each late Neanderthal that overlapped with segregating sites in the 1KG or with sites where the Altai Denisovan and Altai Neanderthal were fixed for different allelic states. To do so, we utilized `BEAGLE v5.4`, with

1KG individuals serving as the reference panel, and included the late Neanderthal, Altai Neanderthal, and Denisovan in the phasing panel [2]. In the second step, we attempted to resolve the remaining heterozygous sites in each late Neanderthal that could not be statistically phased—i.e., the sites that are invariant in the 1KG and did not overlap with a fixed difference between the Altai Denisovan and Altai Neanderthal. For these sites, we used a read-based phasing approach. Using `IGV v2.8.10`, we inspected reads from the BAM files and inferred haplotypes based on reads overlapping adjacent heterozygous sites that had been statistically phased. These inferred read-based haplotypes were then validated by checking their consistency with the phase of adjacent heterozygous sites determined by `BEAGLE v5.4` (see Supplemental Section S4 for a detailed discussion).

Psuedo-Ancestry Painting (*PAP*) in the late Neanderthals

As genome-wide phasing is not currently possible for archaic genomes, we calculated Pseudo-Ancestry Painting (*PAP*) scores in order to assign alleles from a target heterozygous individual to haplotypes with fixed differences from two source individuals. Specifically, let $T = \{het_1, \ldots, het_n\}$ denote the set of all $n$ heterozygous sites (i.e., $het_i$) in a target individual for a given region, and let $S^1 = \{aac_1, \ldots, aac_n\}$ and $S^2 = \{aac_1, \ldots, aac_n\}$ denote the sets of alternative allele counts where $aac_i \in \{-1, 0, 1, 2\}$ (note that $-1$ represents a missing genotype due to not passing quality control in that given individual), at all of the heterozygous sites in $T$ for two source individuals, respectively. The *PAP* score corresponds to the number of heterozygous sites in $T$ that can be explained by the two source individuals, normalized over the total number of heterozygous sites, and is defined as:

$$PAP\ Score = \frac{1}{n}\sum_{i=1}^{n} 1_{\{S_i^1 \neq S_i^2\}},$$

where $1_{\{S_i^1 \neq S_i^2\}}$ is an indicator variable that is defined as:

$$1_{\{S_i^1 \neq S_i^2\}} = 1, \text{ if } S_i^1 = 0 \text{ and } S_i^2 = 2,$$
$$1_{\{S_i^1 \neq S_i^2\}} = 1, \text{ if } S_i^1 = 2 \text{ and } S_i^2 = 0,$$
$$1_{\{S_i^1 \neq S_i^2\}} = 0, \text{ otherwise.}$$

We aimed to explain the excess of heterozygosity in the Chagyrskaya and Vindija Neanderthals targets, by calculating *PAP* scores using a pairing of the Altai Neanderthal and Denisovan as sources, as well as a pairing of an MXL individual (i.e., NA19664) who is homozygous for the *Denisovan-like* haplotype at the 72kb region and an YRI individual (i.e., NA19190) who is homozygous for the *Human-like* haplotype. To ensure that the *PAP* scores are properly behaved we computed additional *PAP* configurations where the Altai Neanderthal and Denisovan are the target individuals and the focal MXL (i.e., NA19664) and YRI (i.e., NA19190) individuals are sources, as a negative control experiment. For each configuration, to assess if *PAP* scores are significantly elevated at the focal 72kb *MUC19* region we compared the observed *PAP* scores, per configuration, to a distribution of *PAP* scores from the genomic background of 72kb non-overlapping windows with comparable effective sequence lengths—i.e., within one standard deviation from the mean of the windowed effective sequence length distribution. To calculate *P-values* per configuration, we determined the proportion of windows from the genomic background with a *PAP* score greater than or equal to what we observed at the 72kb *MUC19* region, where a *P-value* less than 0.05 is considered statistically significant.

#### Copy Number Polymorphism of a 30bp Tandem Repeat Motif Between the *Human-like* and *Archaic* Haplotypes

##### *Repeat Counts from Short-Read Data*

We used the high-coverage 1KG genomes from [82]. We followed previously established methods to estimate repeat length from short-read data [83]. We used the `view` command implemented in `SAMtools v1.9` to extract and count reads from CRAM files for each sample that maps to the repeat region (hg38, Chr12:40482139-40491565). This process was repeated for two non-repetitive control regions of the human genome (hg38, Chr7:5500000-5600000 and Chr12:6490000-6590000) to calculate the average read density for each sample. The fraction of enrichment or depletion of reads in the repeat region, relative to the control regions, was used to estimate the repeat length and average number of repeat copies compared to the reference human genome, which contains 287.5 copies of the 30bp repeat. After estimating the number of repeat copies for each 1KG individual, we determined the number of inferred introgressed tracts overlapping the repeat region (hg19, Chr12:40876395-40885001) in the same manner as described in the *"Identification of the MUC19 Introgressed Region"* Methods subsection. We defined outlier individuals with an elevated number of repeat copies as those having more than 487 repeat copies, corresponding to the 95th percentile of the 1KG repeat copy number distribution at *MUC19*.

To investigate the relationship between copy number variation and introgression at *MUC19*, we first partitioned all 1KG individuals into two subsets, each consisting of two groups for hypothesis testing: 1) admixed American vs non-admixed American individuals; and 2) individuals with at least one introgressed tract overlapping the repeat region vs individuals with no introgressed tracts overlapping the repeat region. For each subset, we calculated the proportion of outlier individuals within each group and performed a Proportions *Z*-test to determine if there was an enrichment of individuals with an elevated number of repeat copies between the groups, using the `statsmodels.stats.proportion.proportions_ztest` function implemented in `statsmodels v0.13.2` [75]. Additionally, for each subset we conducted a Mann-Whitney

*U*-test to assess whether there was a significant difference in the distributions of repeat copies between the two groups, using the `scipy.stats.mannwhitneyu` function implemented in `scipy v1.7.2 [81]`. For both sets of statistical tests, *P-values* less than 0.05 were considered statistically significant.

Following the analysis using all 1KG individuals, we then sought to further investigate the relationship between copy number variation and introgression at *MUC19*, by only considering the outlying individuals. To do so, we partitioned the 118 individuals with an elevated number of repeat copies into two different subsets, both consisting of two groups for hypothesis testing: 1) admixed American vs non-admixed American outlying individuals, and 2) outlying individuals with at least one introgressed tract overlapping the repeat region vs outlying individuals with no introgressed tracts overlapping the repeat region. For each subset, we computed the proportion of individuals within each group with respect to all outlier individuals and again performed a Proportions *Z*-test, using the `statsmodels.stats.proportion.proportions_ztest function` implemented in `statsmodels v0.13.2` [75]. Similarly, we conducted a Mann-Whitney *U*-test to assess differences in the distributions of repeat copies between the two outlier groups, using the `scipy.stats.mannwhitneyu` function implemented in `scipy v1.7.2` [81]. For both sets of statistical tests, *P-values* less than 0.05 were considered statistically significant.

To directly assess the relationship between copy number variation and introgression at *MUC19*, we performed a series of correlation tests to investigate the association between an individual's number of introgressed tracts overlapping the repeat region (i.e., 0, 1, or 2 tracts) and the number of repeat copies. Specifically, we calculated Spearman's correlation coefficient for all 1KG individuals, as well as for each super population and population, using the `scipy.stats.spearmanr` function implemented in `scipy v1.7.2` [81]. Correlation coefficients were not computed for super populations or populations with no introgressed tracts overlapping the repeat region, and *P-values* less than 0.05 were considered statistically significant. Lastly, to directly assess the relationship between copy number variation and admixture in the Americas, we first computed the ancestry proportions in the repeat region for each admixed individual, which was done by intersecting each admixed American individual's local ancestry call BED files with the repeat region (hg19, Chr12:40876395-40885001) using `BEDTools v2.31.0` [84]. For each admixed American population, we then computed Spearman's correlation coefficient between an individual's ancestry proportion (i.e., 0%, 50%, or 100%) per ancestry component (i.e., Indigenous American, European, and African ancestry) and the number of repeat copies, again using the `scipy.stats.spearmanr` function implemented in `scipy v1.7.2 [81]`. After applying a Bonferroni correction to account for three multiple comparisons—i.e., one per ancestry component—*P-values* less than 0.0167 were considered statistically significant.

*Repeat Counts from Long-Read data*

Phased long-read genomes were obtained from the HPRC and HGSVC [59, 42]. A region corresponding to hg38 Chr12:40482543-40491234 was extracted from each of the FASTA files, and FASTA files with variation at those coordinates were extracted from Chr12:40479026-40491234. These regions were then trimmed to match the start and end of hg38 Chr12:40482593-40491199, to match the coordinates of the largest Simple Tandem Repeat from Tandem Repeat Finder [85]. The repeat length was divided by 30 and then averaged between the two haplotypes for each individual to calculate the estimated repeat copies. To ensure that the repeat copies inferred from long-read data was comparable to our inferences from short-read data we performed a least-squares linear regression and corresponding Pearson's correlation coefficient between the estimated repeat copies for individuals with both types of sequencing data available using the `scipy.stats.linregress` function implemented in `scipy v1.7.2 [81]`, where a *P-value* less than 0.05 was considered statistically significant.

<u>Denisovan-specific Coding Mutations</u>

The focal 72kb *MUC19* region contains two Denisovan-specific synonymous mutations and nine Denisovan-specific missense mutations relative to the hg19 reference genome. To estimate the potential impact of these missense mutations, we annotated each mutation with its respective Grantham score. Grantham scores quantify the physicochemical distance between amino acids and are informative about how a mutation alters the protein's functional properties. Following the classification by Li et al. [73], we categorized the effects of each mutation as follows: Grantham score < 50 = conservative, 51–100 = moderately conservative, 101–150 = moderately radical, and > 150 = radical. To assess the conservation of each substituted base, we annotated each coding mutation with the PhyloP score of the reference base (obtained from the UCSC genome browser 100 vertebrates Basewise Conservation track [32]).

Since missense mutations change the amino acid identity, which may potentially affect the protein's function, we first aimed to determine whether admixed American populations are more likely to harbor Denisovan-specific missense mutations than non-admixed American populations. For each of the nine Denisovan-specific missense mutations, we computed the frequency of the mutation within both groups and performed a Proportions *Z*-test using the `statsmodels.stats.proportion.proportions_ztest` function implemented in `statsmodels v0.13.2` [75] to assess if there is an enrichment of these mutations among admixed populations. Additionally, we used Fisher's Exact Test using the `scipy.stats.fisher_exact` function as implemented in `scipy v1.7.2` [81] to determine whether these missense mutations are overrepresented in admixed American populations compared to non-admixed Americans. African individuals were excluded from the non-admixed American population group in both analyses and *P-values* less than 0.05 were considered statistically significant. Next, we evaluated if there is an enrichment of Denisovan-specific missense mutations between three focal groups: MXL individuals from the 1KG, Indigenous American individuals from the SGDP, and ancient Indigenous American individuals. For each of the nine Denisovan-specific missense mutations, we conducted pairwise comparisons between these three groups using the Proportions Z-test and Fisher's Exact Test, following the same approach as outlined above. For comparisons including ancient Indigenous American individuals, we only considered individuals who passed quality control at the respective site, and again, *P-values* less than 0.05 were considered statistically significant. To assess the relationship between the frequency of introgressed tracts overlapping *MUC19* and the frequency of Denisovan-specific missense mutations among 1KG populations, we performed a least-squares linear regression and corresponding Pearson's correlation coefficient between the introgressed tract frequency and mean Denisovan-specific missense mutation frequency among 1KG populations using the `scipy.stats.linregress` function implemented in `scipy v1.7.2 [81]`, where a *P-value* less than 0.05 was considered statistically significant. Lastly, to test if recent admixture in the Americas has diluted the introgressed ancestry at *MUC19* we quantified the relationship between an individual's Indigenous American ancestry proportion at the focal 72kb region and the frequency of a Denisovan-specific missense mutation at position Chr12:40808726. Given that all nine of the Denisovan-specific missense mutations are found within a ~17.5kb region, we used the missense mutation at Chr12:40808726 as this position has genotype information in 20 out of the 23 ancient American individuals (Table S30). For each admixed American individual in the 1KG we computed their respective Indigenous American ancestry proportion for the focal 72kb region in a similar manner as described in "*Repeat Counts from Short-Read Data*" Methods subsection, and since all of the ancient American individuals pre-date the colonization events in the Americas we assumed their Indigenous American ancestry proportion is 100%. We then assessed the relationship between an individual's Indigenous American ancestry proportion and the frequency of the Denisovan-specific missense mutation at position Chr12:40808726 by performing a least-squares linear regression and corresponding Pearson's correlation coefficient using the `scipy.stats.linregress` function implemented in `scipy v1.7.2 [81]`, where a *P-value* less than 0.05 was considered statistically significant.

Main figures data and code:

Data underlying Figures 1-5 is available at: https://github.com/David-Peede/MUC19/blob/main/figure_nbs/data.tar.gz

Python code to replicate Figure 1 is available at:

https://github.com/David-Peede/MUC19/blob/main/figure_nbs/figure_1_v_revisions.ipynb

Python code to replicate Figure 2 is available at:

https://github.com/David-Peede/MUC19/blob/main/figure_nbs/figure_2_v_revisions.ipynb

Python code to replicate Figure 3 is available at:

https://github.com/David-Peede/MUC19/blob/main/figure_nbs/figure_3_v_revisions.ipynb

Python code to replicate Figure 4 is available at:

https://github.com/David-Peede/MUC19/blob/main/figure_nbs/figure_4_v_revisions.ipynb

Python code to replicate Figure 5 is available at:

https://github.com/David-Peede/MUC19/blob/main/figure_nbs/figure_5_v_revisions.ipynb

#### Supplementary Text

##### Section S1: Assessing Significance of *PBS* Values at the Focal Regions

Computing *P-values* for each per-SNP *PBS* value in the 742kb region presents two main challenges: 1) the SNPs in this region are correlated due to linkage disequilibrium (LD), which means that SNPs are not independent from one another; and 2) maintaining sufficient statistical power to correctly reject the null hypothesis despite the large number of multiple comparisons. Because of this, we computed *P-values* to assess per-region *PBS* tests, and demonstrate that both the focal 742kb and 72kb regions *PBS* values are statistically significant compared to the genomic background of non-overlapping windows (Figure S7; Table S10-S11). We believe that per-region tests are the most appropriate for evaluating *PBS* significance, as they account for both of the aforementioned challenges.

*Assessing Significance of Per-Region PBS Values using Different Empirical Distributions*

As previously mentioned, we believe that the per-region tests are the most appropriate for evaluating the significance of *PBS* in both the focal 742kb and 72kb *MUC19* regions. Given that there is substantial evidence to demonstrate that the MXL population has experienced introgression in both focal regions, we sought to re-assess significance using a null distribution derived from introgressed regions in MXL. To do so we relied on the previously published `SPrime` introgression maps for MXL as `SPrime` reports the putative introgressed alleles in MXL and specifies if the allele matches the Altai Denisovan and/or the Altai Neanderthal [2]. We then restrict our analyses only to the sites that `SPrime` denotes as a match with the Altai Denisovan or Altai Neanderthal, and then recomputed our observed $PBS_{MXL:CHB:CEU}$ value for both the focal 742kb and 72kb *MUC19* regions. To assess if our focal regions exhibit elevated $PBS_{MXL:CHB:CEU}$ values we built the a null distribution from the genomic background of 742kb and 72kb non-overlapping windows—of similar effective sequence length—by again computing per-region $PBS_{MXL:CHB:CEU}$ values only for the `SPrime` sites where the allele matches the Altai Denisovan and/or the Altai Neanderthal. To calculate *P-values* for each focal region, we determined the proportion of windows from the genomic background with a per-region $PBS_{MXL:CHB:CEU}$ value greater than or equal to what we observed at the 742kb and 72kb *MUC19* regions, respectively, where a *P-value* less than 0.05 is considered statistically significant. Concordant with our per-region $PBS_{MXL:CHB:CEU}$ results using all sites we find that both the focal 742kb ($PBS_{MXL:CHB:CEU}$ : 0.291, *P-value*: 7.92e-4) and 72kb ($PBS_{MXL:CHB:CEU}$ : 0.280, *P-value*: 0.002) *MUC19* regions are statistically significant when only considering `SPrime` sites where the allele matches the Altai Denisovan and/or the Altai Neanderthal, which further reinforces the signals of positive selection observed in MXL (Figure S45; Table S50).

Throughout this study, when computing *PBS* we have used MXL as our target population and CHB and CEU as our control populations, because 1) our $U_{AFR,B,Denisovan}(1\%, 30\%, 100\%)$ and *iHS* analyses only find signals consistent with introgression and positive selection in MXL and not other AMR populations, and 2) East Asian populations are likely more closely related to the ancestral population that expanded into the Americas, while European populations admixed with the Indigenous American populations during colonization in the Americas. To further understand if the signals of positive selection at *MUC19* are unique to MXL and to assess if our per-region *PBS* results are robust to the choice of control population we recomputed $PBS_{A,B,C}$ for all pairwise combinations of target populations (i.e., *A = {AMR populations}*) and control populations (i.e., *B = {EAS and SAS populations}* and *C = {EUR populations}*). For each unique combination of target and control populations we computed the observed $PBS_{A,B,C}$ value for each focal *MUC19* region (i.e., the 72kb and 742kb regions). To compute *P-values* we compared each observed $PBS_{A,B,C}$ value to a distribution of $PBS_{A,B,C}$ values from the genomic background of 742kb and 72kb non-overlapping windows with comparable effective sequence length and determined the proportion of windows from the genomic background with a per-region $PBS_{A,B,C}$ value greater than or equal to what we observed at the 742kb and 72kb region, respectively, where a *P-value* less than 0.05 is considered statistically significant. Consistent with our $U_{AFR,B,Denisovan}(1\%, 30\%, 100\%)$ and *iHS* analyses we never observe a significant per-region $PBS_{A,B,C}$ value for either the focal 742kb or 72kb *MUC19* regions when the target population is PEL, CLM, or PUR regardless of the two control populations (Tables S51-S52). In stark contrast, we find that when MXL is the target population we always observe a significant per-region $PBS_{A,B,C}$ value for both the focal 742kb and 72kb *MUC19* regions for all permutations of control populations (Tables S51-S52). Taken together these analyses demonstrate that the signals of positive selection at *MUC19* is unique to MXL (i.e., not observed in other AMR populations) and that our per-region *PBS* results are not dependent on using CHB and CEU as the two control populations.

*Empirically Assessing Significance of Per-SNP PBS Values*

To further reiterate, computing *P-values* for each per-SNP PBS value in the 742kb region presents two main challenges: 1) the SNPs in this region are correlated due to LD and consequently SNPs are not independent of one another; and 2) maintaining sufficient statistical power to correctly reject the null hypothesis despite the large number of multiple comparisons. Nevertheless, we first used a non-parametric approach to identify outlier per-SNP *PBS* values at the focal 742kb region. Specifically, we determined the number of SNPs out of the 6144 total SNPs that have a *PBS* value greater than 99.95th percentile of the genome-wide distribution (i.e., *PBS* > ~0.259). We find a total of 417 such SNPs, which includes 208 out of 217 archaic SNPs, all 135 Denisovan-specific SNPs, and 72 out of the 80 Neanderthal-specific SNPs in the 742kb region (Table S53).

As with any outlier approach, our stringent outlier threshold results in 0.05% of SNPs, genome-wide, being labeled as outliers genome-wide. Another way to demonstrate how extreme these per-SNP *PBS* values are is to determine how much these *PBS* values deviate from the genome-wide per-SNP *PBS* distribution. Therefore, for each of the 417 outlier SNPs, we computed the percentile rank, which corresponds to the percentage of the genome-wide distribution of *PBS* values that are less than the given outlier *PBS* value (Figure S46; Table S54). The mean percentile rank of all outlier SNPs is 99.975 (+/- 95% CIs: 0.001), which demonstrates that, on average, the outlier SNPs we identified in the 742kb region exhibit *PBS* values in the top 0.025% of per-SNP *PBS* values genome-wide (Table S54).

Throughout this manuscript we have used the Bonferroni correction to correct for multiple comparisons. This approach ensures that the family-wise error rate—the probability of obtaining at least one false positive among all independent tests—is less than our significance threshold of 0.05. However, it has two main limitations in our context: 1) the SNPs in the 742kb region are correlated due to LD, leading to multiple dependent tests rather than independent ones; and 2) due to the large number of comparisons (i.e., n = 6144 SNPs), this stringent approach reduces the statistical power for detecting true positives. These limitations are demonstrated in Figure S47, where not a single SNP has a Bonferroni-corrected *P-value* less than 0.05; in fact, the smallest Bonferroni-corrected *P-value* we obtain is ~0.122. Furthermore, using the Bonferroni-corrected *P-values* would suggest that all of the SNPs in this region are putatively neutral, implying that the observed *PBS* values are a consequence of neutral demographic processes. If this were indeed true, it would represent an extraordinary case of genetic drift, as it would have resulted in 417 SNPs exhibiting *PBS* values in the top 0.05% of the genome-wide distribution—a truly remarkable feat of genetic drift if true.

These Bonferroni-corrected *P-values* suggest that none of our observed per-SNP *PBS* values in the focal 742kb region are consistent with an evolutionary history of natural selection, which contradicts our per-region *PBS* results and our orthogonal lines of evidence for natural selection using $U_{AFR,MXL,Denisovan}(1\%, 30\%, 100\%)$ and *iHS*. We argue that this contradiction arises from the violation of Bonferroni's assumption of independence and the notoriously conservative nature of this approach. To overcome these limitations, we take inspiration from the GWAS community, which encounters similar hypothesis testing challenges, and instead correct for the False Discovery Rate (FDR). This approach accounts for the expected proportion of false positives among the tests we conclude are statistically significant. Specifically, we employ the Benjamini–Hochberg Procedure (BHP), which accounts for the correlated dependent tests in the focal 742kb region due to LD. We direct readers to Benjamini and Hochberg 1995 for a more thorough statistical explanation. Outlined below is how we implemented the BHP in our study:

Let $m$ represent the number of per-SNP *PBS* values in the focal 742kb region, let $p_i$ represent the uncorrected *P-value* for the $i^{th}$ SNP, and let $bhp_i$ represent the BHP corrected *P-value* for the $i^{th}$ SNP.

1) We set our FDR threshold at 0.01, meaning that if we determine *N* total tests to be significant, we expect $N \times 0.01$ of these tests to be false positives.
2) We compute uncorrected *P-values* (denoted as $p_i$) for each per-SNP *PBS* value in the 742kb region ($m$ = 6144 SNPs) as: $p_i$ = (the number of *PBS* `values in the per-SNP genome-wide distribution ≥ the respective observed` *PBS* value) / (total number of *PBS* values in the per-SNP genome-wide distribution). We then rank these uncorrected *P-values* in ascending order such that $p_1 \leq p_2 \leq \ldots \leq p_m$.
3) From the ranked uncorrected *P-values*, we compute the corrected *P-value* (denoted as $bhp_i$) as $bhp_i = (p_i \times m) / i$. To ensure non-decreasing $bhp_i$, we take the cumulative minimum of the $bhp_i$*'s* from rank 1 to rank $m$.
4) We determine a test as statistically significant if $bhp_i < 0.01$.

Figure S48 shows the BHP-corrected *P-values* for the SNPs in the focal 742kb region. We find that 485 SNPs are statistically significant, including all 417 SNPs above the 99.95th genome-wide percentile (Table S55). These 485 statistically significant SNPs have a mean percentile rank of 99.969 (+/- 95% CIs: 0.002), indicating that these SNPs, on average, have per-SNP *PBS* values in the top ~0.033% of the genome-wide distribution (Table S55). Furthermore, given our stringent FDR threshold of 0.01, we expect that of these 485 SNPs, approximately 4.85 (485 × 0.01 = 4.85) are false positives. This suggests that all of our 417 outlier SNPs are statistically significant, as the expected false positives would be fewer than the difference between the 485 significant SNPs and our 417 more stringently defined outlier SNPs.

*Assessing Significance Using Simulations with the Demographic Model for MXL*

To assess whether the observed *PBS* values could arise due to neutral processes or negative selection, we used `SLiM 4.1` [87] to simulate a region around *MUC19*. Our goal was to test the hypothesis that neutral demographic processes alone or a scenario of heterosis cannot explain the observed signatures of positive selection in MXL individuals. We simulated a genomic region of 742kb (Chr12: 40272001–41014000), which corresponds to the focal region with the longest introgressed tract found in MXL, using the genetic structure from the modern human genome build hg19. We divided the 742kb region into exons and non-coding regions to reflect its genomic structure. Exon ranges were defined using GENCODE v.14 annotations [88], with neutral blocks defined as regions between exons. To best reflect the demographic history of MXL individuals, we simulated under a demographic model with parameters recently inferred for MXL from the 1000 Genomes Project (1KG) [97]. We chose this model because it most accurately captures the complex demographic history of the MXL population from the 1KG.

In particular, Medina-Muñoz et al. [28] inferred previously unknown demographic parameters, including the split time between East Asian and ancestral Indigenous American populations, using 50 genomes of unadmixed Indigenous individuals from Mexico within a four-population model (i.e., Africans, Europeans, East Asians, and Indigenous Americans). To infer parameters of the recent admixture history in the MXL population (e.g., timing and admixture proportions), they used the ancestry tract length distribution obtained from their local ancestry inference, and the `TRACTS` [89] method to infer the timing of admixture events. Subsequently, they created a demographic model for the MXL population by combining the four-population model and the parameter estimates of recent admixture history. Using simulations, they demonstrated that this model could reproduce patterns of allele frequencies and ancestry tract length distributions observed in the MXL population. To incorporate archaic variation, we extended the Medina-Muñoz et al. [28] model to include an introgression event from Neanderthals into the ancestral population of Europeans, East Asians, and Indigenous Americans, as well as an introgression event from Denisovans into the ancestral populations of East Asians and Indigenous Americans. Our extended demographic model consists of 11 populations and the following changes from the original model without archaic introgression:

1) An ancestral population at equilibrium splits into two subpopulations: Archaic Humans 20,225 generations ago and Anatomically Modern Humans (AMH) 16,671 generations ago.
2) 15,090 generations ago, the Archaic population splits into the Denisovan (DEN) and Neanderthal (NEA) populations.
3) 3,046 generations ago, the AMH population splits into two subpopulations: Africans (AFR) and Out of Africa Bottleneck (OOA).
4) 2,069 generations ago, a pulse of gene flow (lasting one generation) occurs from the Neanderthal (NEA) to Out of Africa Bottleneck (OOA) populations with an introgression probability of 5% [91].
5) 1,764 generations ago, the Out of Africa Bottleneck (OOA) population splits into the European (EUR) and East Asian (EAS) populations.
6) 1,552 generations ago, a pulse of gene flow (lasting one generation) occurs from the Denisovan (DEN) to East Asian (EAS) populations with an introgression probability of 5% [96].

7) 1,123 generations ago, the Indigenous American population (MXB) splits from the East Asian (EAS) population.
8) 16 generations ago, the admixed population (MXL) splits from the Indigenous American population (MXB) population consisting of 75.08% Indigenous American ancestry and 24.92% European ancestry.
9) 13 generations ago, a single pulse of admixture occurs from the African (AFR) and East Asian (EAS) populations to the admixed population (MXL) population with admixture proportions of 11.18% and 0.74%, for African and East Asian ancestry, respectively.

We acknowledge that the proportion of introgression used (5% for both Neanderthals and Denisovans) is higher than typically inferred. We chose these values to ensure a sufficient number of simulated replicates with archaic alleles segregating in the present modern human populations. All parameters are described in Supplementary Table S56, and the model is illustrated in Supplementary Figure S49.

We first simulated two different scenarios to have two null expectations to test for positive selection. In one scenario (neutral), we assume all mutations are neutral. In the second scenario (negative), we assume that mutations are deleterious when mutations occur in exons and neutral otherwise. We assume deleterious mutations are recessive to simulate heterosis, which has previously been suggested to mimic signatures of adaptive introgression [9, 20]. We assumed selection coefficients for recessive deleterious mutations are drawn from a gamma distribution of fitness effects (DFE) with a shape parameter of 0.186 and an average selection coefficient $s$ = -0.01315 [93]. We set the ratio of nonsynonymous (recessive deleterious mutations) to synonymous mutations (neutral mutations) to 2.31:1 based on Kim et al. [93] and we used the recombination map for this region as defined by [80]. The per base pair mutation rate was fixed at 1.315e-8 which we obtained by using estimates of the mutation rate scaled over 5kb windows provided in Hubisz and Siepel [86]. We obtained 10,000 simulation replicates for each selection model—i.e., neutral or negative. Each replicate simulation was conditioned on having at least one Denisovan and Neanderthal allele segregating in the MXL population in the present at a bi-allelic site within the simulated 72kb region. Additionally, we conducted further simulations to evaluate the robustness of the *PBS* statistic to detect positive selection under a realistic demography for MXL. We extended our baseline neutral scenario described above to model a mutation in the simulation that has positive effects on fitness. To do so, we first select neutral mutations segregating in MXB right after the EAS-MXB split. Then, we randomly choose one of the neutral mutations based on 1) being of archaic origin (i.e. a mutation that originated in the archaic lineage, either Neanderthal or Denisovan) and 2) being present in our 72kb focal region. For the chosen mutation we modeled positive selection with varying selection coefficient to $s$ = 0.1, $s$ = 0.01, $s$ = 0.0015, respectively, in MXB, while keeping its fitness effects neutral in EAS and EUR.

For simulations with positive selection we obtained 1,000 simulations conditioned on the selected mutation not being lost and at highest frequency in the sampled MXL population in addition to archaic allele condition for the 72kb region previously described for the neutral and heterosis simulations.

To assess the probability of observing a SNP segregating at a frequency of 0.3 or higher in MXL, we first computed the frequency of every SNP across all 10,000 replicates for all SNPs and archaic SNP partitions—where archaic SNPs are the union of Denisovan and Neanderthal SNPs. We calculated *P-values* as the proportion of SNPs segregating at a frequency greater than or equal to 0.3 in MXL compared to the total number of bi-allelic SNPs across all 10,000 replicates per simulated model—i.e., neutral or negative—considering *P-values* less 0.05 as statistically significant (Figure S50; Table S57). Across both simulated models, Archaic SNPs segregating at a frequency of 0.3 or higher are significantly rare (Figure S50; Table S57). However, this is not true when considering all SNPs in the simulated 742kb region and is not significant (Figure S50; Table S57). These results suggest that observing Archaic SNPs at a high frequency (≥ 0.3) is not expected under neutral demographic processes or heterosis.

In our empirical per-SNP *PBS* analysis, after correcting for multiple comparisons using the BHP, we found that 485 SNPs in the observed 742kb region have a *P-value* less than to 0.01 (Table S54). To account for the expected number of false positives among these 485 tests, we consider only the 417 SNPs with *PBS* values greater than the 99.95th percentile as statistically significant, with *PBS* values ranging from ~0.264 to ~0.429. Notably, among these 417 outlier SNPs, there are only 57 unique *PBS* values, including 20 unique *PBS* values for archaic SNPs, four unique *PBS* values for Denisovan-specific SNPs, and 17 unique *PBS* values for Neanderthal-specific SNPs—further indicating that these SNPs are not independent of one another. We computed uncorrected *P-values* for all 417 outlier *PBS* values using our simulated data. We calculated uncorrected *P-values* as the proportion of simulated per-SNP *PBS* values greater than or equal to the observed *PBS* value compared to the total number of SNPs with defined *PBS* values across all 10,000 replicates per simulated model—i.e., neutral or negative. We then corrected for multiple comparisons using both the Bonferroni correction and BHP, as previously described, where the number of multiple comparisons corresponds to the number of *PBS* values for each SNP partition.

Using a significance threshold of 0.05 for Bonferroni-corrected *P-values* and a FDR threshold of 0.01 for BHP-corrected *P-values*, when using the neutral simulated distribution we find that 97 out of 417 SNPs have statistically significant *PBS* values and when using the heterosis simulated distribution we find that 184 out of 417 SNPs have statistically significant *PBS* values when using the Bonferroni method to correct for 417 multiple comparisons (Table S58). Conversely, when we correct for multiple comparisons using the BHP we find that all 417 SNPs have statistically significant *PBS* value when compared to both simulated distributions (Table S58). Notably, for the 208 archaic SNP outliers we find that all 208 have statistically significant *PBS* values when correcting for 208 multiple comparisons using both the Bonferroni method and BHP (Table S58). Furthermore, given that there are only 57 unique outlier per-SNP *PBS* values if we were to correct for 57 multiple comparisons, then all 417 outlier per-SNP *PBS* values would always be significant regardless of the method used to correct for multiple comparisons or the simulation model (Table S59). These results suggest that the 417 SNPs with *PBS* values greater than the 99.95th percentile are not expected under neutral demographic processes or heterosis, providing additional evidence that these SNPs have statistically significant *PBS* values, cocordant with our empirical analyses.

As previously mentioned, it's uncommon to assess hypotheses about positive selection for an entire region using per-SNP *P-values*, as SNPs within a region are correlated due to LD. However, if a region has experienced positive selection, one would expect an excess of SNPs segregating at high frequencies and a per-region *PBS* value greater than expected when compared to neutral demographic processes and a scenario of heterosis. To assess these hypotheses using our simulated data, we first computed *P-values* for the number of SNPs in MXL segregating at high frequency (≥ 0.3). This was calculated as the proportion of the 10,000 simulated replicates where the number of SNPs segregating at high frequency (≥ 0.3) is greater than or equal to the number of observed SNPs segregating at high frequency (≥ 0.3) for both focal regions (i.e., 742kb and 72kb) per simulated model—i.e., neutral or negative. We considered *P-values* less than 0.05 as statistically significant (Figure S51-S52; Table S60). Next, for each simulated replicate, we computed per-region *PBS* values for both the simulated 742kb region and 72kb region. We then computed *P-values* for each observed per-region *PBS* value as the proportion of the 10,000 simulated replicates where the per-region *PBS* value is greater than or equal to the observed per-region *PBS* value per simulated model—i.e., neutral or negative—considering *P-values* less than 0.05 as statistically significant (Figure S53-S54; Table S61).

We find that across all SNP partitions, the observed 742kb region has a significantly large density of SNPs segregating at high frequency (≥ 0.3) compared to all simulated models: all high frequency SNPs = 1311 and archaic high frequency SNPs = 208 (Figure S51; Table S60). This is also true for the observed 72kb region: all high frequency SNPs = 300 and archaic high frequency SNPs = 140 (Figure S52; Table S60). These results suggest that both observed focal regions (i.e., 742kb and 72kb) have a larger density of high frequency SNPs, including archaic SNPs, than expected under neutral demographic processes or heterosis (Figure S51-S52; Table S60). Notably, for both focal regions and both SNP partitions (i.e., all SNPs and archaic SNPs), we find that the observed per-region *PBS* values are always statistically significant regardless of the simulated model (Figure S53-S54; Table S61): all SNPs *PBS* values: 742kb = 0.066 and 72kb = 0.127, archaic SNPs *PBS* values: 742kb = 0.291 and 72kb = 0.280. These results suggest that *PBS* values for the entire 742kb and 72kb regions are larger than expected under neutral demographic processes or heterosis (Figure S53-S54; Table S61).

Lastly, to ensure that per-region *PBS* estimates are robust to detect positive selection under the demographic for MXL we computed the *PBS* values for both the simulated 742kb and 72kb regions for all SNPs and archaic SNP partitions per simulated replicate. We repeated this process for all 1,000 simulated replicates for all three sets of simulated selection coefficients. To compute the power of *PBS* per selection coefficient we computed the proportion of the 1,000 replicates with per-region *PBS* values greater than the upper 5% quantile from its respective neutral distribution of per-region *PBS* values consisting of 10,000 replicates. We find that *PBS* has sufficient power to detect instances of positive selection for scenarios of strong positive selection (i.e., $s = 0.1$) and weak positive selection (i.e., $s = 0.01$), but has much less power when the effects of positive selection are extremely weak (i.e., $s = 0.0015$; Table S62). These results suggest that we have power to correctly identify instances of positive selection in MXL using per-region *PBS* values under the assumptions of strong or weak positive selection (Table S62).

In summary, an evolutionary history of introgression and positive selection will result in a region with an elevated number of SNPs at high frequency and elevated per-region *PBS* compared to expectations under neutral demographic processes and a scenario of heterosis. Using the empirical per-SNP *PBS* distribution, we have identified 417 outlier SNPs in the focal 742kb region with *PBS* values greater than the 99.95th genome-wide percentile (> ~0.259) that are statistically significant using the BHP at a FDR threshold of 0.01 (Table S53-S54). Additionally, using the empirical non-overlapping window distribution, we find that both the focal 742kb and 72kb region *PBS* values are statistically significant using a significance threshold of 0.05 (Figure S7; Table S10-S11). These empirical distributions are designed to control for demographic processes, as demography shapes global patterns of variation while selection shapes local patterns of variation. We further assessed if our observed results could be explained by neutrality or a scenario of heterosis using forward-in-time simulations, the most up-to-date demographic model of the MXL population, and the genetic structure of the 742kb *MUC19* region. Our simulations further bolster our empirical per-SNP and per-region *PBS* results, and highlight that we would not expect the observed density of high frequency archaic SNPs under any of our simulated scenarios (Figures S50-S54; Table S57-S62). Taken together, our simulations corroborate our empirically based claim that the 417 per-SNP *PBS* outliers (including all 135 Denisovan-specific) found in the focal 742kb region are statistically significant. Furthermore, both focal regions (742kb and 72kb) exhibit signals consistent with an evolutionary history of introgression and positive selection, which cannot be explained by neutral demographic processes or heterosis and are further corroborated by our orthogonal lines of evidence for adaptive introgression using $U_{AFR,MXL,Denisovan}(1\%, 30\%, 100\%)$ and positive selection using *iHS*.

**Section S2: Identifying Clusters of Extreme *iHS* Scores**

As previously mentioned in the Methods section, there is no formal way to assess the significance of *iHS* [79]. To determine if the focal 742kb and 72kb regions exhibit signals consistent with positive selection, we assessed whether these regions harbored clusters of extreme *iHS* (i.e., normalized $|iHS| > 2$). To do so, we first binned the genome into non-overlapping windows of size 742kb and 72kb, and then computed the proportion of SNPs with extreme normalized $|iHS|$ for windows containing more than 10 SNPs with defined values. This allowed us to build a genome-wide distribution of the proportion of SNPs with extreme normalized $|iHS|$ and to determine if the focal 742kb and 72kb regions are among the top 1% of windows with the highest fractions of extreme normalized $|iHS|$ per population (Figure S9; Table S12-S13). We performed this analysis for all 1KG populations using all SNPs with a minor allele frequency > 5%. We found that in MXL the 742kb focal region falls within the top ~0.5% of 742kb windows (Percentile Rank: 99.527) with the highest fractions of extreme normalized $|iHS|$ (Figure S9; Table S12). Similarly, the 72kb focal region falls within the top ~0.6% of 72kb windows (Percentile Rank: 99.426) with the highest fractions of extreme normalized $|iHS|$ (see the grey shaded region in Figure S9; Table S13). Notably, no other 1KG population showed signals consistent with positive selection using clusters of extreme normalized $|iHS|$ (Table S12-S13).

We then repeated this analysis for the archaic SNPs—i.e., the union of Denisovan-specific, Neanderthal-specific, and shared archaic SNPs—in every non-African 1KG population. For each population, we built a distribution of the proportion of archaic SNPs with normalized |*iHS*| > 2 for non-overlapping windows with more than 10 archaic SNPs, using window sizes of 742kb and 72kb. We assumed that a population's focal region (i.e., 742kb and 72kb) shows signals consistent with positive selection if the observed proportion of extreme normalized |*iHS*| values falls within the top 1% of windows with the highest fractions of extreme normalized |*iHS*| for each of its respective window size distribution. We set a minimum threshold of at least 10 archaic SNPs observed in each respective focal region (i.e., 742kb and 72kb) for the analysis of clusters with extreme normalized |*iHS*|.

Similar to the *iHS* results using all SNPs, MXL is the only population where we identify clusters of extreme normalized |*iHS*| at the focal 742kb and 72kb regions. At the focal 742kb region: 208 out of 217 archaic SNPs have normalized |*iHS*| > 2 in MXL, which resides in the top 1% of 742kb windows with the highest fractions of extreme normalized |*iHS*| for archaic SNPs (Table S63). At the focal 72kb region: 135 out of 140 archaic SNPs have normalized |*iHS*| > 2 in MXL, which resides in the top 1% of 72kb windows with the highest fractions of extreme *iHS* scores for archaic SNPs (Table S64).

It should be noted that the absence of clusters of extreme normalized |*iHS*| within the top 1% of the non-overlapping window distribution in other AMR populations (i.e., PEL, CLM, and PUR) may be a consequence of recent admixture [97]. Specifically, *iHS* has been shown to be underpowered to identify signals of positive selection in admixed populations, as it does not account for changes in LD due to admixture events [97]. On the other hand, it is notable that we are still able to detect signatures of selection at the focal regions in MXL despite *iHS* being underpowered in admixed populations, likely owing to the high frequency of introgressed haplotypes in this population.

**Section S3: Benchmarking Phasing at the Focal 72kb Region**

Our goal is to phase the Chagyrskaya and Vindija Neanderthals at the focal 72kb region to determine if these late Neanderthals harbor one *Denisovan-like* haplotype. The lack of population-level sampling for archaic humans and the short fragmented nature of ancient DNA reads currently limit reliable genome-wide phasing of high-coverage archaic data. However, we believe that phasing the Chagyrskaya and

Vindija Neanderthals at the focal 72kb region is feasible due to three key reasons: 1) the Altai Neanderthal and Denisovan are homozygous at most sites in this region and thus are effectively already phased, 2) the Chagyrskaya and Vindija Neanderthals exhibit more heterozygous sites than expected, and 3) two divergent haplotypes are segregating among 1KG individuals in this region. Our approach relies on using the 1KG as a reference panel of haplotypes because the introgressed haplotype shows high affinity to the Denisovan, while the non-introgressed haplotypes have an affinity to the Altai Neanderthal in the focal 72kb region.

Since phasing archaic genomes is not a standard practice in the field, we first evaluated the feasibility of phasing a *hypothetical* late Neanderthal. To do so, requires *a priori* knowledge of the true haplotypes to determine the accuracy of phasing using the 1KG as a reference panel. Therefore, we generated a Synthetic Neanderthal with characteristics mirroring those of the late Neanderthals in the focal 72kb region. We created this Synthetic Neanderthal using the 1KG and a combined archaic dataset, focusing on the intersection of four different sets of segregating sites: 1) 184 sites where either late Neanderthal has a heterozygous genotype, 2) 170 sites where the Altai Neanderthal and Denisovan are fixed for different allelic states, 3) 1,587 sites variable in the 1KG, and 4) 82 sites variable among archaics but invariant in the 1KG. This results in 155 segregating sites where the Altai Neanderthal and Denisovan are homozygous for different alleles (i.e., fixed differences) to use as our ground truth haplotypes to assess the accuracy of phasing. We then generated genotypes for phasing by sampling one allele from the Altai Neanderthal and one from the Denisovan, creating a Synthetic Neanderthal with 155 heterozygous genotypes, which is comparable to what we observe in the Chagyrskaya and Vindija Neanderthals at the focal 72kb region (Figure S32; Table S43).

We then used `BEAGLE v5.4` to phase this Synthetic Neanderthal, using 1KG individuals as the reference panel and including the Synthetic Neanderthal, Altai Neanderthal, and Denisovan in the phasing panel [2]. We included the Altai Neanderthal and Denisovan in the phasing panel because `BEAGLE v5.4` utilizes information from all individuals in the phasing panel in its two-stage phasing approach [2]. Our results show that `BEAGLE v5.4` perfectly phased the Synthetic Neanderthal by correctly assigning the Altai Neanderthal and Denisovan alleles to separate haplotypes in the Synthetic Neanderthal at all 155 heterozygous genotypes (Figure S55). This result strongly suggests that BEAGLE v5.4 is able to confidently and accurately phase each of the late Neanderthals at the focal 72kb region by using the 1KG individuals as the reference panel and including the Altai Neanderthal and Denisovan in the phasing panel.

#### Section S4: Phasing the Late Neanderthals at the Focal 72kb Region

Given that we were able to perfectly phase our Synthetic Neanderthal, we proceeded to phase the Chagyrskaya and Vindija Neanderthals using the combined 1KG and archaic dataset at the focal 72kb region. For each late Neanderthal, we only attempted to statistically phase the variable sites in the 1KG that also passed quality control in the late Neanderthal, and the invariant sites in the 1KG where the late Neanderthal is heterozygous and the Denisovan and Altai Neanderthal are fixed for different allelic states. For the aforementioned sites, we used `BEAGLE v5.4` for statistical phasing; using the 1KG individuals as the reference panel and including the late Neanderthal, Altai Neanderthal, and Denisovan in the phasing panel [2]. For the remaining heterozygous sites in each late Neanderthal that were not included in the statistical phasing dataset (i.e., sites invariant in the 1KG and not fixed differences between the Denisovan and Altai Neanderthal), we attempted to resolve these sites by using read-based phasing. Specifically, we manually inspected the reads from the BAM files in `IGV v2.8.10` [98] to determine the haplotypes from reads overlapping adjacent heterozygous sites that were statistically phased. We then validated these read-based haplotypes by ensuring that they were concordant with the phase of the adjacent heterozygous sites that were determined by `BEAGLE v5.4`.

In the combined dataset, the Chagyrskaya Neanderthal has a total of 167 heterozygous sites in the focal 72kb region. We were able to statistically phase 163 of these heterozygous sites, and attempted to manually resolve the remaining four heterozygous sites from the reads (Table S65). At position 40767590, the phase could not be resolved due to a lack of overlapping reads with the adjacent heterozygous sites at positions 40761946 (5644bp away) and 40768905 (1315bp away). However, the phase at position 40821807 was resolved with 11 reads overlapping its adjacent heterozygous sites 40821795 and 40821847. Of these 11 reads, five supported a G-C-C:40821795-40821807-40821847 haplotype, while six supported an A-T-A:40821795-40821807-40821847 haplotype. These haplotypes were further corroborated by the phase of the adjacent sites determined by `BEAGLE v5.4`, which are G-C:40821795-40821847 and A-A:40821795-40821847. Similarly, the phase at position 40826155 was resolved with 17 reads overlapping its preceding heterozygous site, 40826138, six reads overlapping its proceeding heterozygous site, 40826201, and three reads overlapping both. Of the 17 reads overlapping the preceding heterozygous site 40826138, 12 supported a T-A:40826138-40826155 haplotype and five supported a G-G:40826138-40826155 haplotype. Of the six reads overlapping the proceeding heterozygous site, 40826201, five supported an A-G:40826155-40826201 haplotype and one supported an A-A:40826155-40826201 haplotype. All three reads overlapping both adjacent heterozygous sites supported a T-A-G:40826138-40826155-40826201 haplotype. Based on the phase of the adjacent sites reported by `BEAGLE v5.4`, which were T-G:40826138-40826201 and G-A:40826138-40826201, the haplotypes at these three positions were determined to be T-A-G:40826138-40826155-40826201 and G-G-A:40826138-40826155-40826201. Lastly, the phase at position 40828121 could not be resolved, as the two overlapping reads with the preceding heterozygous site, 40761946, supported non-concordant haplotypes (i.e., T-A:40761946-40828121 and C-A :40761946-40828121), and no reads overlapped with the proceeding heterozygous site 40828306 (185bp away).

In the combined dataset, the Vindija Neanderthal has a total of 170 heterozygous sites in the focal 72kb region. We were able to statistically phase 165 of these heterozygous sites, and attempted to manually resolve the remaining five heterozygous sites from the reads (Table S66). At position 40765370, the phase could not be resolved due to the absence of overlapping reads with the adjacent heterozygous sites at positions 40761946 (3424bp away) and 40768905 (3535bp away). Similarly, at position 40783344, the phase could not be resolved due to the lack of overlapping reads with the adjacent heterozygous sites at positions 40775127 (8217bp away) and 40784418 (1074bp away). However, the phase at position 40821807 was resolved with 25 reads overlapping its preceding heterozygous site 40821795, 11 reads overlapping its proceeding heterozygous site 40821847, and seven reads overlapping both. Of the 25 reads overlapping the preceding heterozygous site 40821795, seven supported a G-C:40821795-40821807 haplotype, and 18 supported an A-T:40821795-40821807 haplotype. Of the 11 reads overlapping the proceeding heterozygous site 40821847, eight supported a T-A:40821807-40821847 haplotype, and three supported a C-C:40821807-40821847 haplotype, while all seven reads overlapping both adjacent heterozygous sites supported an A-T-A:40821795-40821807-40821847 haplotype. Based on the phase of the adjacent sites reported by `BEAGLE v5.4`, which were A-A:40821795-40821847 and G-C:40821795-40821847, the haplotypes at these three positions were determined to be A-T-A:40821795-40821807-40821847 and G-C-C:40821795-40821807-40821847, which is concordant with the haplotypes found in the Chagyrskaya Neanderthal at these exact positions (Table S65). Additionally, the phase at position 40824215 was resolved with four reads overlapping its preceding heterozygous site 40824154, and no reads overlapping its proceeding heterozygous site 40824266 (51bp away). Of the four reads overlapping the proceeding heterozygous site 40824154, three supported a G-A:40824154-40824215 haplotype, and one supported an A-C:40824154-40824215 haplotype, which support two valid haplotypes. Lastly, the phase at position 40829306 could not be resolved due to the lack of overlapping reads with the preceding heterozygous site 40828306 (1000bp away), and the absence of a proceeding heterozygous site, as position 40829306 is the last heterozygous site in the Vindija Neanderthal at the focal 72kb region.

In summary, we were able to statistically phase the majority of heterozygous sites in the late Neanderthals (Chagyrskaya: 163 out of 167; Vindija: 165 out of 170). For the remaining sites we could not statistically phased (Chagyrskaya: 4 out of 167; Vindija: 5 out of 170), we successfully resolved roughly half of these heterozygous sites through read-based phasing (Chagyrskaya: 2 out of 4; Vindija: 2 out of 5; Table S65-S66). For subsequent analyses involving the phased late Neanderthals we set the unresolved heterozygous genotypes to missing and adjusted the effective sequence lengths by 2bp and 3bp for the Chagyrskaya and Vindija Neanderthals, respectively. Overall, integrating statistical and read-based phasing approaches allowed us to maximize the number of phased heterozygous sites to achieve near-complete phasing, resolving all but two sites in the Chagyrskaya Neanderthal (165 out of 167), all but three in the Vindija Neanderthal (167 out of 170), and in turn providing a robust set of phased haplotypes for subsequent genetic analyses.

#### Section S5: Patterns of Sequence Divergence at the Focal 72kb Region

The focal 72kb region exhibits unique patterns of sequence divergence—defined as the number of pairwise differences normalized by effective sequence length—between modern human haplotypes and the unphased genotypes of high-coverage archaic individuals. Notably, introgressed haplotypes display a low sequence divergence from the Altai Denisovan, a high sequence divergence from the Altai Neanderthal, and an intermediate sequence divergence from the Chagyrskaya and Vindija Neanderthals (Figure S20-S22; Figure S24; Dataset 1). Given that haplotypes among modern humans exhibit signatures consistent with introgression—i.e., the bimodal distribution of sequence divergence relative to the Altai Denisovan in all non-African 1KG populations, a pattern not observed in African populations (Figure S20)—we sought to determine whether haplotypes in the 1KG were significantly closer than expected to the Altai Denisovan based on the genomic background distribution. To this end, we calculated the sequence divergence between each chromosome in a modern human individual and the unphased archaic genotype per 72kb non-overlapping windows of comparable effective sequence length—i.e., within one standard deviation of the mean 72kb effective sequence length distribution—to construct the genomic background distribution of sequence divergence for each high-coverage archaic individual. To determine whether a modern human individual's haplotype at the focal 72kb region is significantly closer to a high-coverage archaic genome than expected, we computed *P-values* as the proportion of windows from the genomic background with a sequence divergence less than that observed at the focal 72kb region. After correcting for two multiple comparisons—i.e., one per haplotype using the Bonferroni correction—we considered *P-values* less than 0.025 to be statistically significant.

Interestingly, our analysis revealed that no haplotypes in the 1KG are significantly closer to any high-coverage archaic individual at the focal 72kb region (Dataset 1). This finding is particularly striking given that the focal 72kb region harbors an exceptionally high density of Denisovan-specific alleles (Table S4), and at remarkably high frequencies in the MXL (Table S8). At first glance, given that no 1KG haplotypes are significantly closer to the Altai Denisovan than expected, it seems contradictory to a scenario of Denisovan introgression. However, consistent with previous findings from introgressed segments in Papuans [95], we believe that the sequenced Altai Denisovan belongs to a population that is divergent from the donor Denisovan population that interbred with modern humans. To determine whether the sequence divergence between the introgressed haplotype and the Altai Denisovan at the focal 72kb region aligns with expectations relative to other Denisovan introgressed fragments across the genome, we analyzed the well-known Denisovan ancestry tracts in Papuans, who received multiple pulses of introgression from distinct Denisovan populations. Using 1KG individuals from the YRI, MSL, and ESN populations as an outgroup, we used `hmmix` to infer introgressed tracts per haplotype for all Papuan individuals in the phased SGDP dataset across all autosomes [26]. We then filtered these inferred Papuan tracts, retaining only archaic tracts with a posterior probability greater than or equal to 0.8 and those that shared more SNPs with the Altai Denisovan than with any of the three high-coverage Neanderthals, as reported by `hmmix` [26]. These filtered tracts were considered to be Denisovan introgressed tracts. Next, we queried these inferred Denisovan introgressed tracts within a Papuan individual and computed the sequence divergence between the Papuan individual's haplotype and the Altai Denisovan. Repeating this process for all Papuan individuals allowed us to construct a distribution of sequence divergence for all Denisovan introgressed tracts among Papuans (Figure S56).

To illustrate that the sequence divergence between the introgressed haplotype and the Altai Denisovan at the focal 72kb region is consistent with expectations relative to other Denisovan introgressed tracts, we compared the observed sequence divergence between the Altai Denisovan and two focal individuals—MXL (NA19664) and Papuan (B_Papuan-15)—that harbor two introgressed haplotypes at the focal 72kb region based on our sequence divergence thresholds as described in *"Modern Human Haplotype-Archaic Human Sequence Divergence at the Focal 72kb Region"* Methods subsection. We find that the sequence divergence at the focal 72kb region between these individuals and the Altai Denisovan (NA19664: 0.00097; B_Papuan-15: 0.00104) falls within the distribution of sequence divergence for Denisovan introgressed tracts among Papuans (mean: 0.00083; standard deviation: 0.00072; Figure S56). Specifically, the observed sequence divergence at the focal 72kb region falls within the top ~29% (NA19664 Percentile Rank: 70.607) and ~26% (B_Papuan-15 Percentile Rank: 74.338) of the Denisovan introgressed tracts in Papuans (Figure S56). In summary, by analyzing the sequence divergence distribution of Denisovan introgressed tracts in Papuans, we confirm that the sequence divergence observed between the introgressed haplotypes and the Altai Denisovan at the focal 72kb region is consistent with a scenario of Denisovan introgression. However, the source of this introgressed segment likely did not originate directly from the population represented by the Altai Denisovan.

As previously mentioned, the focal 72kb region exhibits unique patterns of sequence divergence (Figure S20-S24). We have already addressed why the introgressed haplotypes are closer to—albeit not significantly—the Altai Denisovan. The other notable pattern is the high sequence divergence between the introgressed haplotypes and the Altai Neanderthal, coupled with an intermediate sequence divergence from the two late Neanderthals—i.e., Chagyrskaya and Vindija (Figure S20-S22). This intermediate sequence divergence with respect to the late Neanderthals can be attributed to the significantly elevated number of heterozygous sites observed at the focal 72kb region in both individuals (Chagyrskaya: 168 heterozygous sites, *P-value*: 0.0002; Vindija: 171 heterozygous sites, *P-value*: 0.0003; Figure S32; Table S43). We hypothesized that the excess heterozygosity observed in the late Neanderthals is consistent with an evolutionary history involving Denisovan introgression, as introgression introduces genetic variation into the recipient population. Furthermore, we believe two orthogonal lines of evidence support this hypothesis: 1) 1KG individuals harboring exactly one introgressed haplotype at the focal 72kb region exhibit significantly more heterozygous sites than expected (average number of heterozygous sites: ~287, *P-value*: 3.157e-5; Figure S33; Table S44), and 2) *D+* tests for local introgression, using the Altai Neanderthal as *P1*, the late Neanderthals as *P2*, and the Altai Denisovan as *P3*, indicate that both late Neanderthals have an excess of allele sharing with the Altai Denisovan at the focal 72kb region than expected under a scenario of no gene flow (Chagyrskaya *D+*: 0.783, *P-value*: 0.029; Vindija *D+*: 0.819, *P-value*: 0.018; Figure S35; Table S46).

To assess this hypothesis, we used a combination of statistical and read-based phasing approaches to achieve near-complete phasing of the late Neanderthals at the focal 72kb region, resolving all but two heterozygous sites in the Chagyrskaya Neanderthal and all but three in the Vindija Neanderthal (see Supplemental Sections S3-S4). Following phasing, we computed the number of pairwise differences between all pairwise combinations of the phased late Neanderthal haplotypes and the unphased genotypes of the Altai Denisovan and Altai Neanderthal. It should be noted that these unphased archaic genotypes are effectively phased due to the low number of heterozygous sites these archaic individuals harbor in the focal 72kb region (Altai Denisovan: 6 heterozygous sites; Altai Neanderthal: 1 heterozygous site; Table S43). Phasing the late Neanderthals revealed that each individual possesses a haplotype with patterns of sequence divergence similar to those observed among *Denisovan-like* haplotypes in the 1KG (Dataset 1). Specifically, the second haplotypes of both the Chagyrskaya and Vindija Neanderthals show a low number of pairwise differences with the Altai Denisovan—43 and 41, respectively (Table S47)—which is similar to the average number of pairwise differences observed between *Denisovan-like* haplotypes and the Altai Denisovan (mean: ~49, standard deviation: ~2; Table S67). Likewise, the late Neanderthals' second haplotypes exhibit a high number of pairwise differences with the Altai Neanderthal (Chagyrskaya: 159.5, Vindija: 161; Table S47), mirroring the pattern seen between *Denisovan-like* haplotypes and the Altai Neanderthal (mean: ~180, standard deviation: ~2.5; Table S67). Conversely, the first haplotypes of both late Neanderthals display a low number of pairwise differences with the Altai Neanderthal (Chagyrskaya: 3.5, Vindija: 4; Table S47) and a high number of differences with the Altai Denisovan (Chagyrskaya: 164, Vindija: 170; Table S47). These findings collectively support our hypothesis that both the Chagyrskaya and Vindija Neanderthals carry a single *Denisovan-like* haplotype at the focal 72kb region, which is consistent with their elevated levels of heterozygosity and our *D*+ tests for Altai Denisovan introgression.

Motivated by our finding that the longest introgressed tract in MXL (i.e., the focal 742 kb region) is closer to—and significantly closer than expected—the two late Neanderthals, and given that our phasing analysis revealed that these Neanderthals harbor a *Denisovan-like* haplotype, we next aimed to determine whether the introgressed haplotype in MXL is significantly closer to the *Denisovan-like* haplotype found in the late Neanderthals. To do so, we computed the number of pairwise differences between the focal MXL individual (NA19664), who carries two introgressed haplotypes at the focal 72kb region, and the *Denisovan-like* haplotype present in the late Neanderthals. Remarkably, we observed that the focal MXL individual exhibits 5 and 4 pairwise differences with the *Denisovan-like* haplotypes found in the Chagyrskaya and Vindija Neanderthals, respectively (Figure S36; Table S48). Since phasing the late Neanderthals genome-wide is not feasible, we assessed if the focal MXL individual's haplotypes are closer than expected to the *Denisovan-like* haplotypes in the late Neanderthals by computing the sequence divergence between each chromosome in a modern human individual and a late Neanderthal's pseudo-haplotype across 72kb non-overlapping windows of comparable effective sequence length—i.e., within one standard deviation of the mean 72kb effective sequence length distribution. Pseudo-haplotypes were generated by randomly sampling one allele at each position from the unphased late Neanderthal genotypes. To determine whether the focal MXL individual's introgressed haplotype at the 72kb region is significantly closer to the *Denisovan-like* haplotype in the late Neanderthals, we computed *P-values* as the proportion of windows from the genomic background with a pseudo-haplotype sequence divergence less than the observed sequence divergence between haplotypes at the focal 72kb region. After applying the Bonferroni correction for four multiple comparisons—i.e., two per haplotype—we considered *P-values* less than 0.0125 to be statistically significant.

Notably, we find that the introgressed haplotype in MXL is significantly closer than expected to the *Denisovan-like* haplotype found in both late Neanderthals (Chagyrskaya: 0.000104, P-value: 0.003; Vindija: 0.000083, P-value: 0.002; Figure S36; Table S48). This result suggests that the *Denisovan-like* segment observed at the focal 72kb region, while of Denisovan origin, was inherited by modern humans through introgression with a population closely related to the Chagyrskaya and Vindija Neanderthals. To further corroborate this conclusion, we recomputed the sequence divergence between each 1KG haplotype and the haplotypes of the phased late Neanderthals at the focal 72kb region (Figure S57-S60; Dataset 3). The increased phase resolution reveals that sequence divergence with respect to the Chagyrskaya and Vindija Neanderthals is no longer intermediate. Instead, introgressed haplotypes in modern humans show a low sequence divergence from the Altai Denisovan, a high sequence divergence from the Altai Neanderthal and the first haplotype (i.e., "Hap. 1"; Figure S57-S58) in the late Neanderthals, and an even lower sequence divergence from the second haplotype of the late Neanderthals (i.e., "Hap. 2"; Figure S59-S60). In summary, our analyses of sequence divergence at the focal 72kb region suggest that the introgressed haplotype present in non-African populations and the two late Neanderthals is likely of Denisovan origin. Although the sequence divergence between the introgressed haplotype in modern humans and the Altai Denisovan is not significantly closer than expected, having phase resolution for the late Neanderthals revealed that the introgressed haplotype in modern humans is significantly closer than expected to the *Denisovan-like* haplotype found in the late Neanderthals. Our results suggest a complex evolutionary history of recurrent introgression at the focal 72kb region, beginning with introgression from a Denisovan population into a late Neanderthal population, and followed by subsequent introgression from a late Neanderthal population into modern humans (Figure S38).

**Section S6: ILS Calculation**

As at least one *MUC19* haplotype is shared by modern humans, Neanderthals, and Denisovans, it is possible that this haplotype has been retained ancestrally in all three branches due to incomplete lineage sorting (ILS), rather than through archaic admixture. Because recombination is expected to break apart haplotypes, and this process is probabilistic given the recombination rate of a genomic region and the length of time experienced between branches, we can calculate the expected length of a haplotype under ILS using a previously published equation [18]:

$$L = 1/(R \; x \; b/G)$$

The expected length of a shared ancestral sequence haplotype ($L$) is calculated as the inverse of regional recombination rate ($R$) multiplied by the branch length ($b$) over the generation time ($G$). We can then calculate the probability of the observed haplotypes: 742kb for the longest introgressed tract, and 72kb for the focal region. The probability ($p$) of seeing a shared haplotype of length ($m$) follows a Gamma distribution with a shape ($S$) of 2 and a rate ($r$) of 1/$L$:

$$p = 1 - GammaCDF(m, S = 2, r = 1/L)$$

Here, we determined the regional recombination rates ($R$) using HapMap [80], used a generation time ($G$) of 29 years, assumed a branch length ($b$) of 550,000 years corresponding to the split time between modern humans and Neanderthals [99], and computed the probability ($p$) of seeing a shared haplotype of length ($m$) using the `scipy.stats.gamma.cdf` function implemented in `scipy v1.7.2` [81].

For the 742kb for the longest introgressed tract region the regional recombination rate is 5.99967e-9, which results in an expected length of the haplotype shared through ILS is 8788bp, and the probability of observing a shared haplotype as long or longer than 742kb is <5e-324, thus a model without gene flow can be rejected for the 742kb region. For the focal 72kb region the regional recombination rate is 1.03395e-8, which results in an expected length of the haplotype shared through ILS is 5100bp, and the probability of observing a shared haplotype as long or longer than 72kb is <5e-324, thus a model without gene flow can be rejected for the 72kb region as well. Taken together, we can confidently conclude that the observed signals we see at both the 742kb and 72kb *MUC19* region are not likely a result of ILS.

**Supplemental Figures**

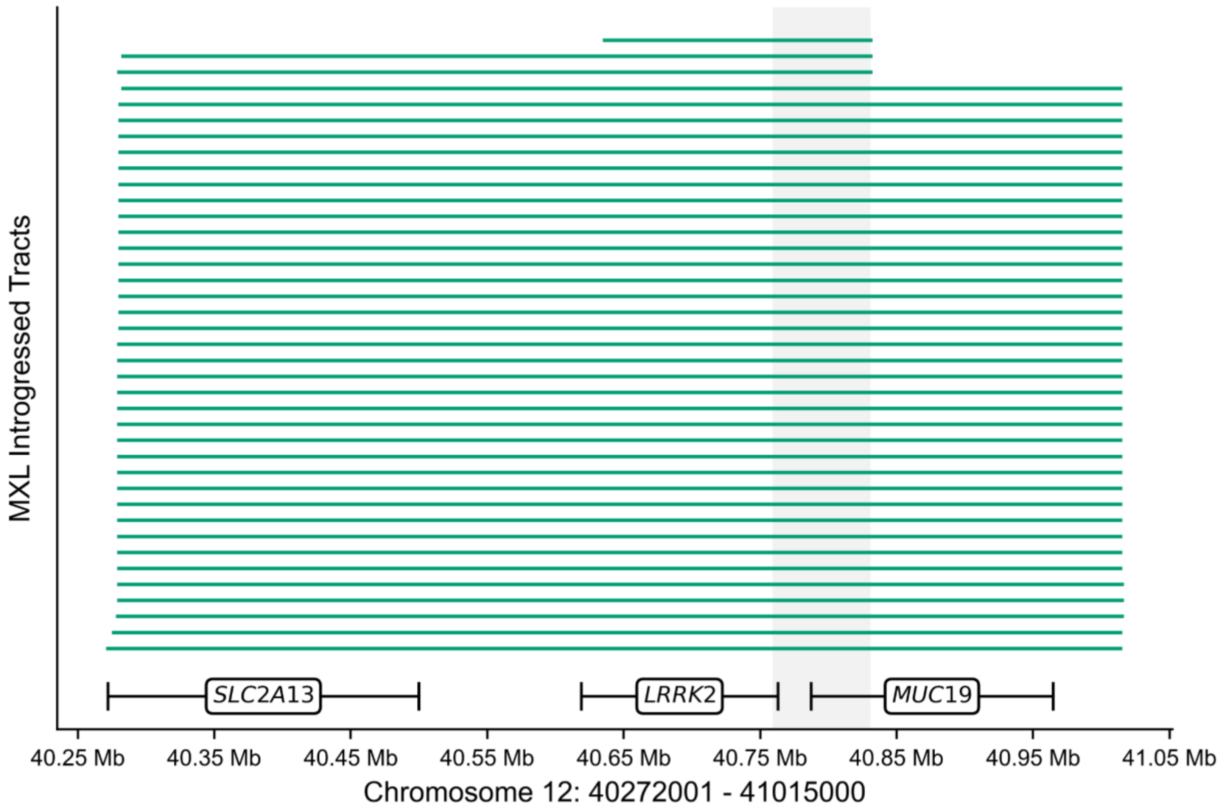


**Fig. S1. Introgressed tracts overlapping *MUC19* in MXL.**
Introgressed tracts—sorted from longest to shortest—overlapping the *MUC19* NCBI RefSeq coordinates (hg19, Chr12:40787196-40964559) among MXL individuals. The gray shaded region corresponds to the focal 72kb region, which is the densest contiguous region of introgressed tracts longer than 40kb. The three box and whiskers plots correspond to the genomic interval for each of the hg19 NCBI RefSeq genes found within the longest introgressed tract found in MXL (i.e., the focal 742kb region) individuals. The *MUC19* and *LRRK2* genes are fully encompassed within the 742kb region, while ~65% of *SLC2A13* overlaps the 742kb region.

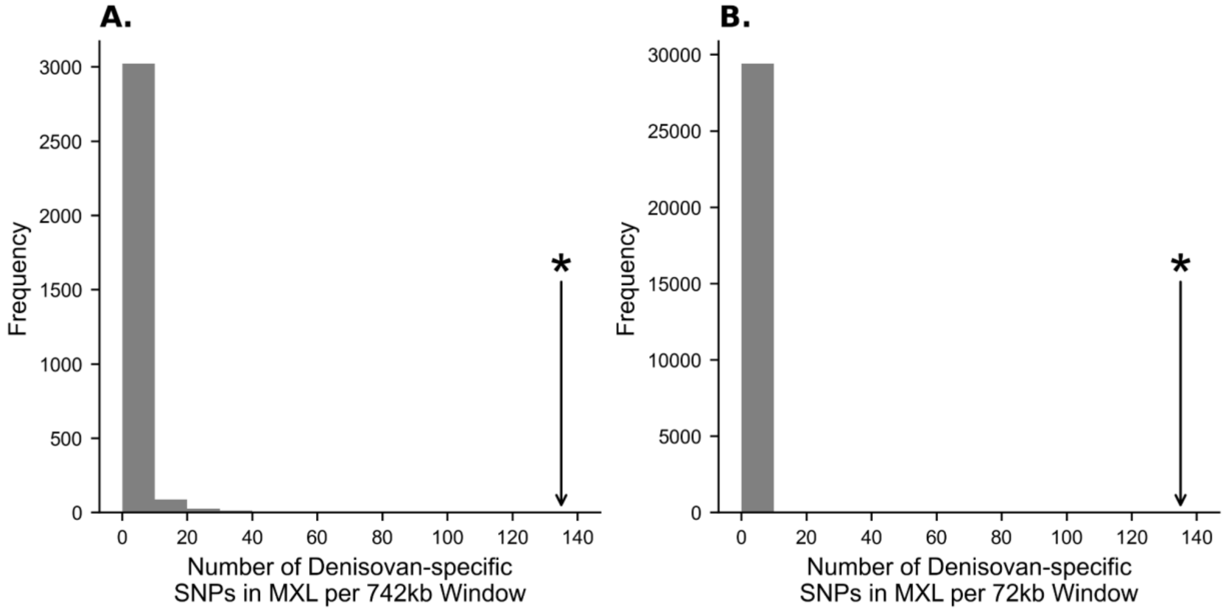


**Fig. S2. High density of Denisovan-specific SNPs in MXL found within the 72kb region of interest.**

Distribution of Denisovan-specific SNPs in MXL for non-overlapping **(A)** 742kb and **(B)** 72kb windows. Denisovan-specific SNPs are defined as those rare or absent in Africa (<1%), present in MXL (>1%), and uniquely shared with the Denisovan. Arrows indicate the observed values for **(A)** the focal 742kb region which corresponds to the longest introgressed tract found in MXL, and **(B)** the focal 72kb region, corresponding to the densest contiguous region (>40kb) of introgressed tracts overlapping *MUC19*. Asterisks above arrows indicate statistical significance when compared to the genome-wide distribution of non-overlapping windows; "ns" denotes non-significance. Note that all 135 Denisovan-specific SNPs found in **(A)** the 742kb region are sequestered within **(B)** the core 72kb region.

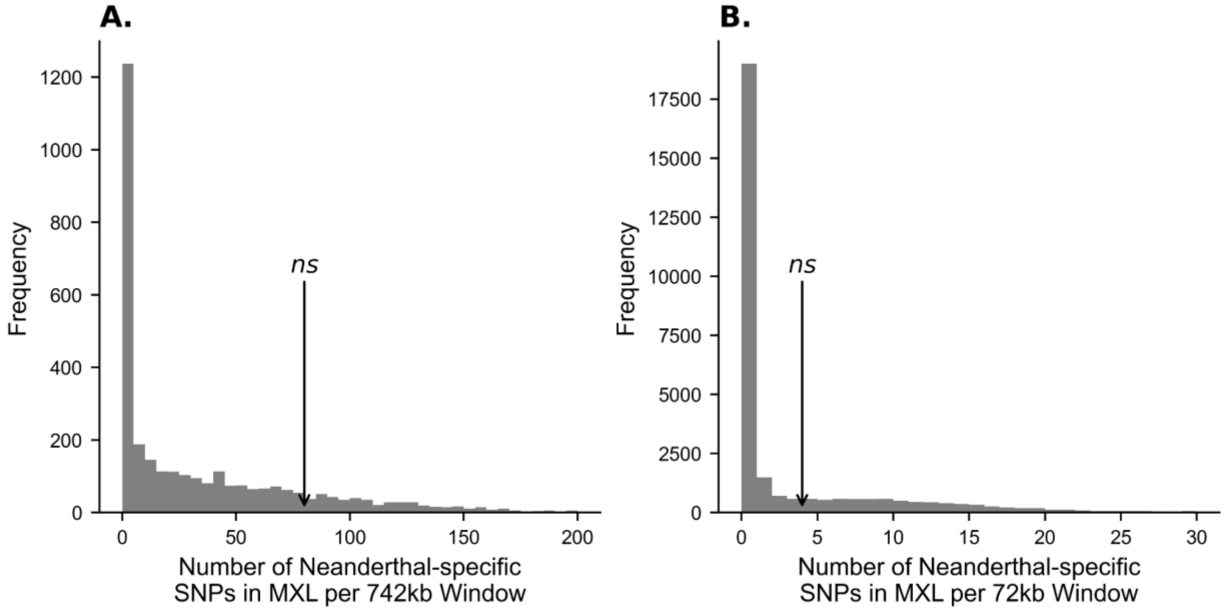


**Fig. S3. The density of Neanderthal-specific SNPs in MXL is within the genome-wide distributions for both focal regions.**
Distribution of Neanderthal-specific SNPs in MXL for non-overlapping **(A)** 742kb and **(B)** 72kb windows. Neanderthal-specific SNPs are defined as those rare or absent in Africa (<1%), present in MXL (>1%), and uniquely shared with at least one of the three high-coverage Neanderthals. Arrows indicate the observed values for **(A)** the focal 742kb region corresponding to the longest introgressed tract found in MXL, and **(B)** the focal 72kb region, which is the densest contiguous region (>40kb) of introgressed tracts overlapping *MUC19*. Asterisks above arrows indicate statistical significance when compared to the genome-wide distribution of non-overlapping windows; "ns" denotes non-significance. Note that of the 80 Neanderthal-specific SNPs found in **(A)** the 742kb region only four are found within **(B)** the core 72kb region.

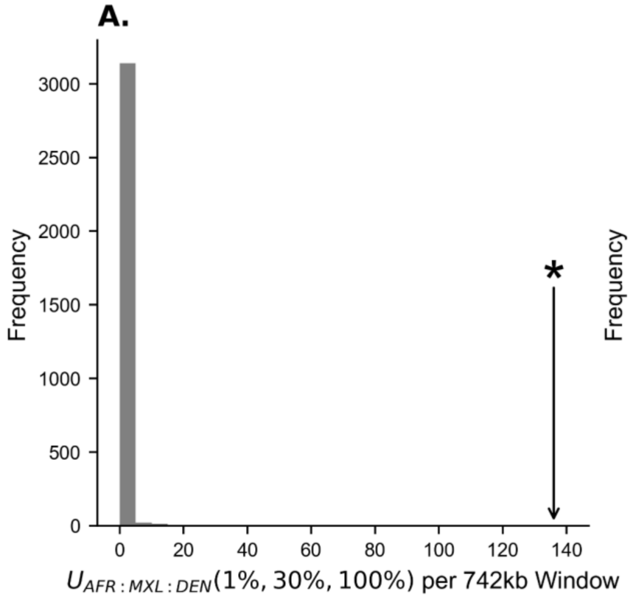


**Fig. S4. Denisovan SNPs within the focal 72kb region are segregating at high frequencies among individuals in MXL.**

Distribution of $U_{AFR,MXL,Denisovan}(1\%, 30\%, 100\%)$ values for non-overlapping **(A)** 742kb and **(B)** 72kb windows. $U_{AFR,MXL,Denisovan}(1\%, 30\%, 100\%)$ quantifies the number of sites where: 1) the Denisovan allele is found in the homozygous state, 2) the Denisovan allele is at low frequency (<1%) in the African super population, and 3) the Denisovan allele is at high frequency (>30%) in MXL. Arrows indicate the observed values for **(A)** the focal 742kb region corresponding to the longest introgressed tract found in MXL, and **(B)** the focal 72kb region, which corresponds to the densest contiguous region (>40kb) of introgressed tracts overlapping *MUC19*. Asterisks above arrows indicate statistical significance when compared to the genome-wide distribution of non-overlapping windows; "ns" denotes non-significance. Note that the $U_{AFR,MXL,Denisovan}(1\%, 30\%, 100\%)$ values for **(A)** the 742kb region and **(B)** the 72kb region are identical.

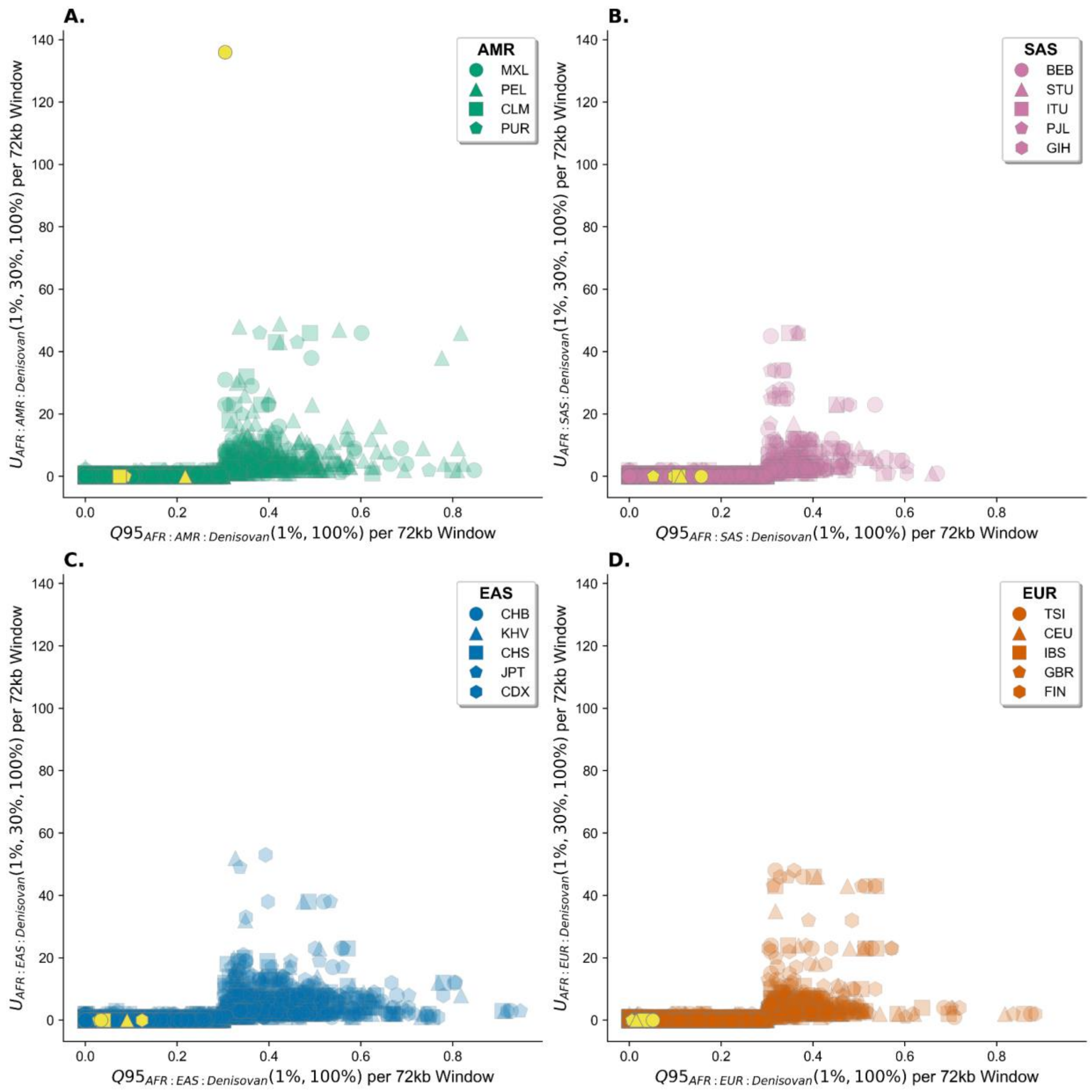


**Fig. S5. Complementary signals of Denisovan adaptive introgression exclusively in MXL at the focal 72kb region.**

Joint distribution of $Q95_{AFR,B,Denisovan}(1\%, 100\%)$ and $U_{AFR,B,Denisovan}(1\%, 30\%, 100\%)$ values for non-overlapping 72kb windows stratified by super population: panel **(A)** - Admixed Americans (AMR), panel **(B)** - South Asians (SAS), panel **(C)** - East Asians (EAS), and panel **(D)** - Europeans (EUR). $Q95_{AFR,B,Denisovan}(1\%, 100\%)$—i.e., the x-axis—quantifies the 95th percentile of Denisovan allele frequencies in the non-African population *B* for sites where: 1) the Denisovan allele is found in the homozygous state, and 2) the Denisovan allele is at low frequency (<1%) in the African super population. $U_{AFR,B,Denisovan}(1\%, 30\%, 100\%)$—i.e., the y-axis—quantifies the number of sites where: 1) the Denisovan allele is found in the homozygous state, 2) the Denisovan allele is at low frequency (<1%) in the African super population, and 3) the Denisovan allele is at high frequency

(>30%) in the non-African population *B*. Note that for windows where $Q95_{AFR,B,Denisovan}(1\%, 100\%)$ is below 30% it is unlikely that there are many, if any, $U_{AFR,B,Denisovan}(1\%, 30\%, 100\%)$ sites. The observed values for each non-African population at the focal 72kb region are denoted by the yellow points, and we only observe signals consistent with Denisovan adaptive introgression in MXL.

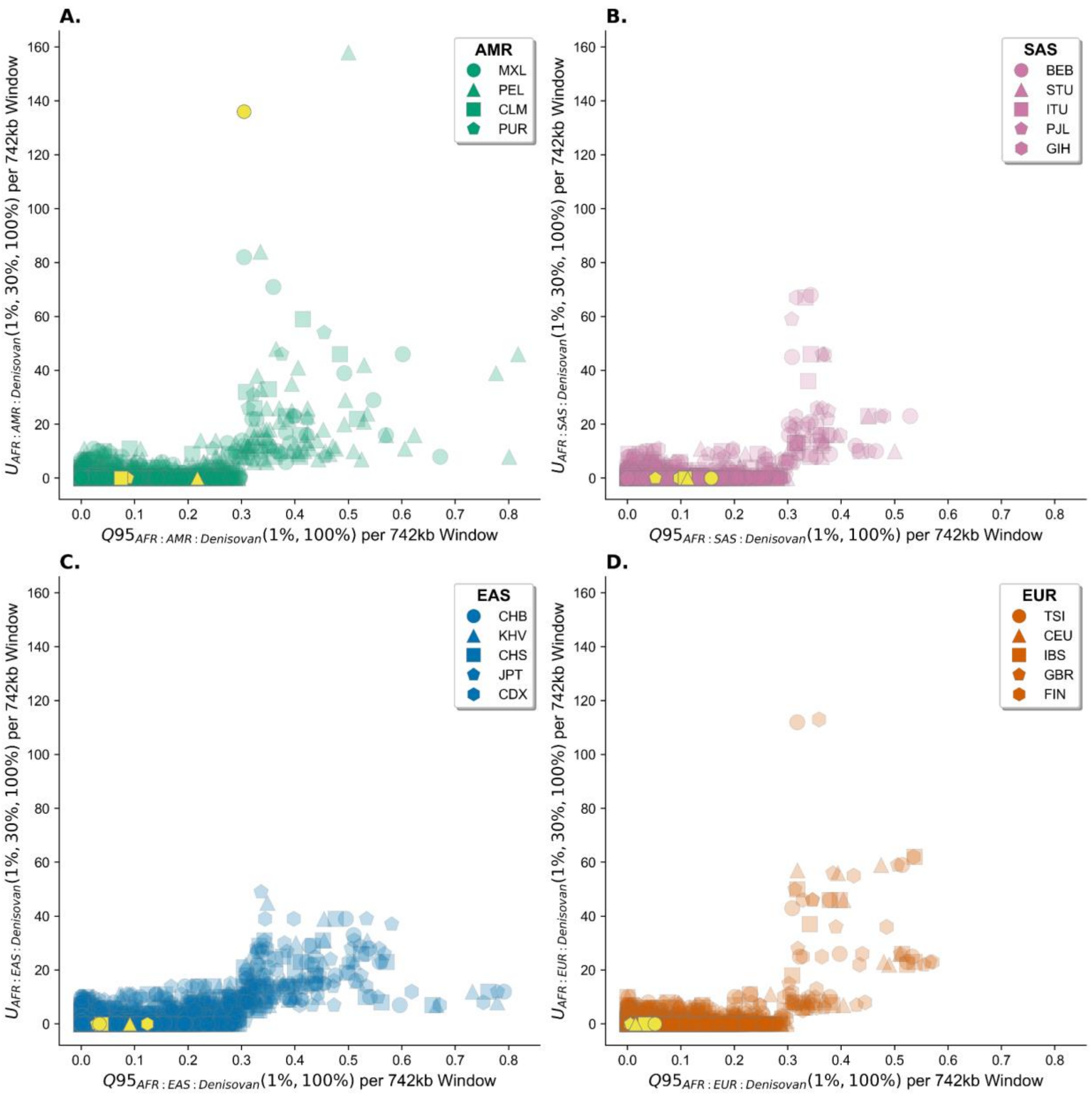


**Fig. S6. Complementary signals of Denisovan adaptive introgression exclusively in MXL at the focal 742kb region.**

Joint distribution of $Q95_{AFR,B,Denisovan}$*(1%, 100%)* and $U_{AFR,B,Denisovan}$*(1%, 30%, 100%)* values for non-overlapping 742kb windows stratified by super population: panel **(A)** - Admixed Americans (AMR), panel **(B)** - South Asians (SAS), panel **(C)** - East Asians (EAS), and panel **(D)** - Europeans (EUR). $Q95_{AFR,B,Denisovan}$*(1%, 100%)*—i.e., the x-axis—quantifies the 95th percentile of Denisovan allele frequencies in the non-African population *B* for sites where: 1) the Denisovan allele is found in the homozygous state, and 2) the Denisovan allele is at low frequency (<1%) in the African super population. $U_{AFR,B,Denisovan}$*(1%, 30%, 100%)*—i.e., the y-axis—quantifies the number of sites where: 1) the Denisovan allele is found in the homozygous state, 2) the Denisovan allele is at low frequency (<1%) in the African super population, and 3) the Denisovan allele is at high frequency

(>30%) in the non-African population *B*. Note that for windows where $Q95_{AFR,B,Denisovan}(1\%, 100\%)$ is below 30% it is unlikely that there are many, if any, $U_{AFR,B,Denisovan}(1\%, 30\%, 100\%)$ sites. The observed values for each non-African population at the focal 742kb region are denoted by the yellow points, and we only observe signals consistent with Denisovan adaptive introgression in MXL.

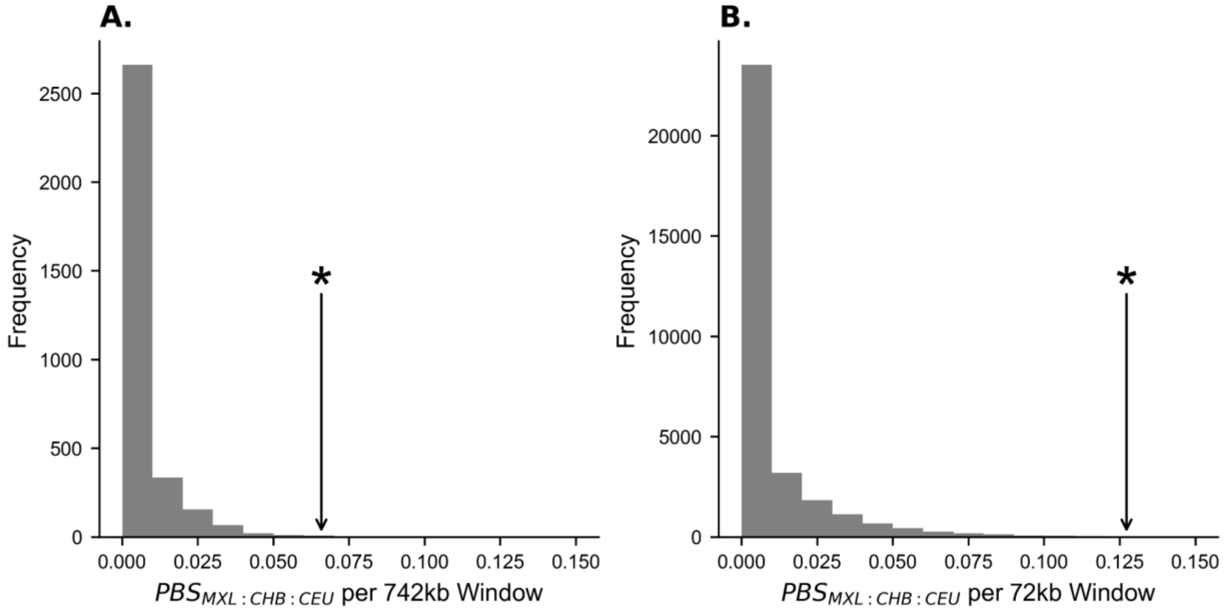


**Fig. S7. Both 742kb and 72kb focal regions exhibit signals of positive selection using the Population Branch Statistic (*PBS*).**

Distribution of $PBS_{MXL:CHB:CEU}$ values for non-overlapping **(A)** 742kb and **(B)** 72kb windows. $PBS_{MXL:CHB:CEU}$ uses the logarithmic transformation of pairwise estimates of $F_{ST}$ to measure the branch length in MXL since its divergence from the two control populations: CHB and CEU. Arrows indicate the observed values for **(A)** the focal 742kb region corresponding to the longest introgressed tract found in MXL, and **(B)** the focal 72kb region, which is the densest contiguous region (>40kb) of introgressed tracts overlapping *MUC19*. Asterisks above arrows indicate statistical significance when compared to the genome-wide distribution of non-overlapping windows; "ns" denotes non-significance. Both panels demonstrate that **(A)** the 742kb region and **(B)** the 72kb region exhibit population genetic signals consistent with positive selection.

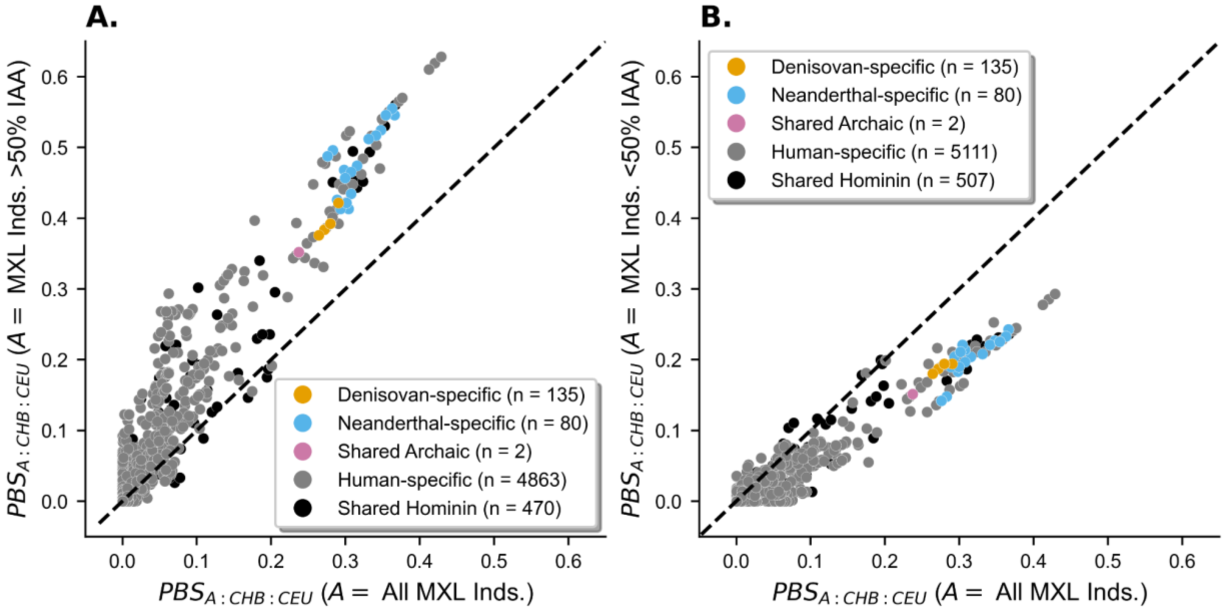


**Fig. S8. Elevated Population Branch Statistic (*PBS*) values among MXL individuals with more than 50% Indigenous American ancestry (IAA).**

Comparison of $PBS_{A:CHB:CEU}$ values for SNPs in the focal 742kb region between: x-axis) all individuals in the MXL population ($n = 64$; $A$ = All MXL Inds.), y-axis) **(A)** MXL individuals with genome-wide IAA > 50% ($n = 27$; $A$ = MXL Inds. >50% IAA) and **(B)** MXL individuals with genome-wide IAA < 50% ($n = 37$; $A$ = MXL Inds. <50% IAA). The orange, sky blue, and reddish purple points represent SNPs that are rare or absent in Africa (<1%), present in MXL (>1%), and are, respectively, either shared uniquely with the Denisovan, uniquely with at least one of the three high-coverage Neanderthals, or shared with both the Denisovan and Neanderthals. The black points represent SNPs present across both modern and archaic human populations, while the gray points represent SNPs private to modern humans. The black dashed line represents the y = x trend line. Points above the dashed line indicate elevated $PBS_{A:CHB:CEU}$ values in the IAA-partitioned deme, while points below indicate higher $PBS_{A:CHB:CEU}$ values when using the entire MXL population. In general, $PBS_{A:CHB:CEU}$ values tend to be higher for **(A)** $A$ = MXL Inds. >50% IAA and lower for **(B)** $A$ = MXL Inds. <50% IAA with respect to $A$ = All MXL Inds.

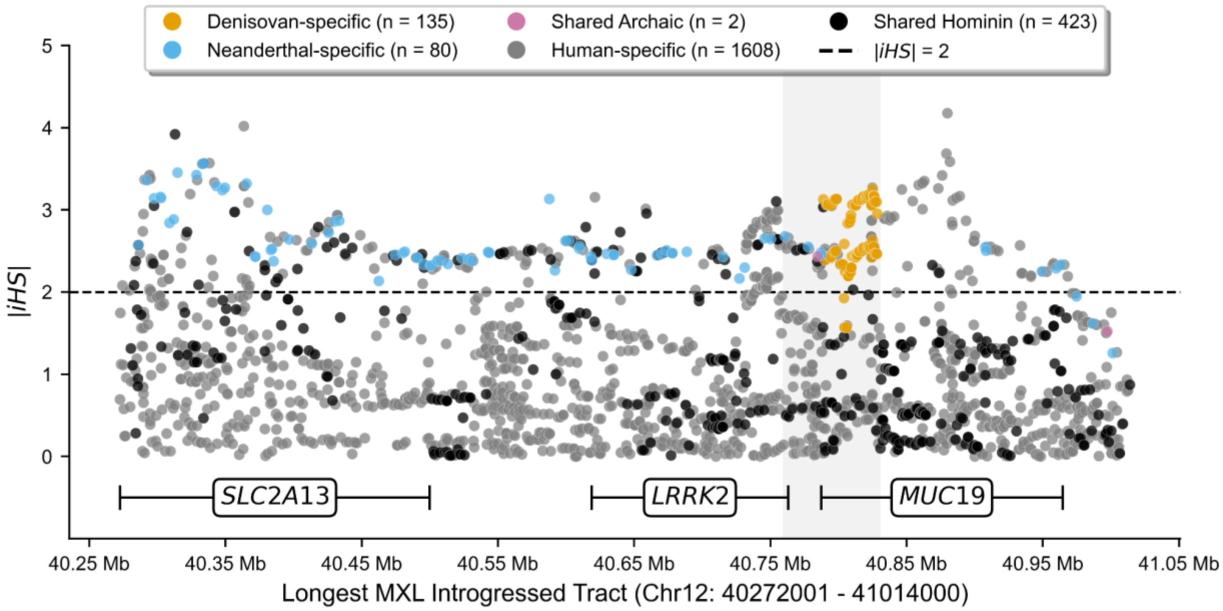


**Fig. S9. Elevated density of extreme Integrated Haplotype Scores (*iHS*) within the focal 742kb region in MXL individuals.**

Normalized |*iHS*| scores computed for every SNP in MXL segregating at a minor allele frequency >5% in the focal 742kb region, which corresponds to the longest introgressed tract found in MXL. The orange, sky blue, and reddish purple points represent SNPs that are rare or absent in Africa (<1%), present in MXL (>1%), and are, respectively, either shared uniquely with the Denisovan, uniquely with at least one of the three high-coverage Neanderthals, or shared with both the Denisovan and Neanderthals. The black points represent SNPs present across both modern human and archaic populations, while the gray points represent SNPs private to modern humans. The black dashed line represents the critical value threshold at |*iHS*| = 2. Points above the |*iHS*| threshold indicate that the haplotype in MXL is unusually long, consistent with signals of positive selection. Note that of the 2248 SNPs with a minor allele frequency >5%, 599 are above the critical value threshold, which include 130 out of the 135 Denisovan-specific SNPs and 77 of the 80 Neanderthal-specific SNPs.

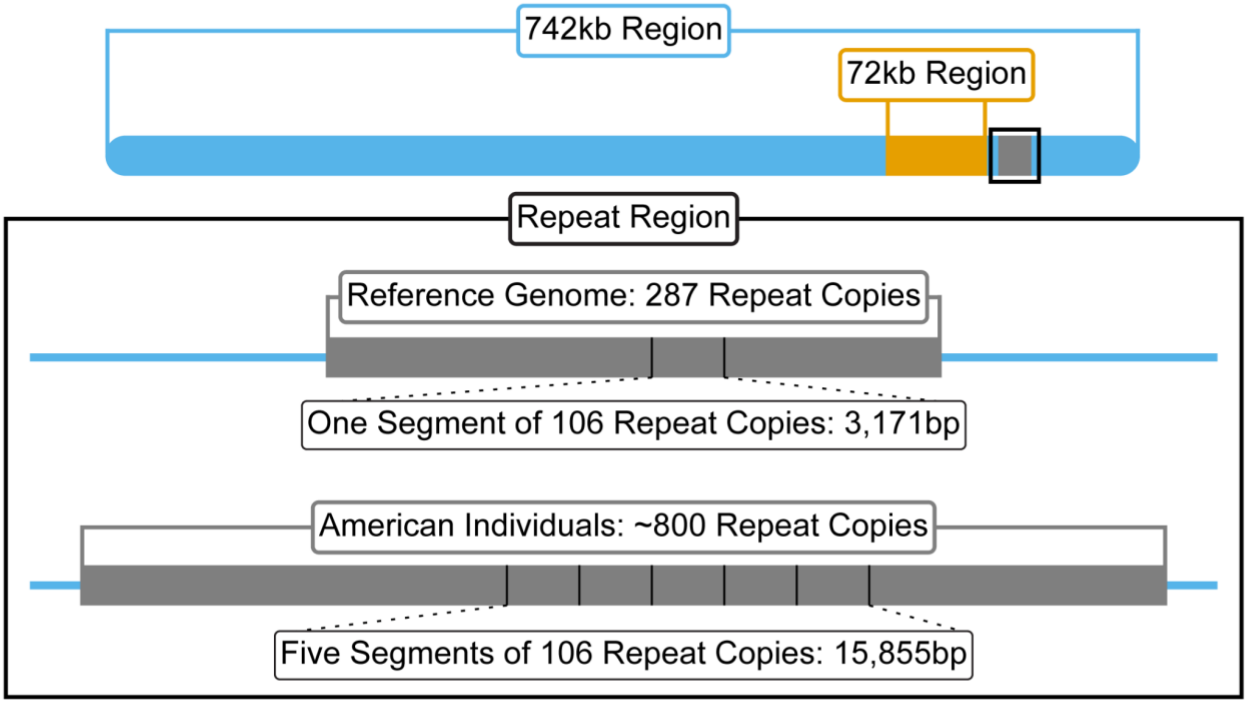


**Fig. S10. Schematic of the *MUC19* tandem repeat expansion in exon 56.**
The top panel denotes the 742kb longest introgressed tract found in MXL in sky blue, the focal 72kb *MUC19* region containing all 135 Denisovan-specific SNPs in orange, and the *MUC19* tandem repeat region in grey. The bottom panel zooms in tandem repeat region where the human reference genome contains 287 copies of a 30bp repeat and within this region contains a single segment consisting of a homologous set of 106 repeats, three of which are 27bp long, resulting in this homologous set having a total repeat length of 3,171bp—i.e., (3 x 27bp repeat) + (103 x 30bp repeat) = 3,171bp. Within this repeat region Admixed American individuals harbor roughly 800 copies of the tandem repeat, notably the segment of 106 repeats is repeated four additional times for a total of five copies of this homologous set (i.e., 1 segment + 4 segments = 5 segments) which has a total repeat length of 15,855bp—i.e., 5 segments x 3,171bp = 15,855bp.

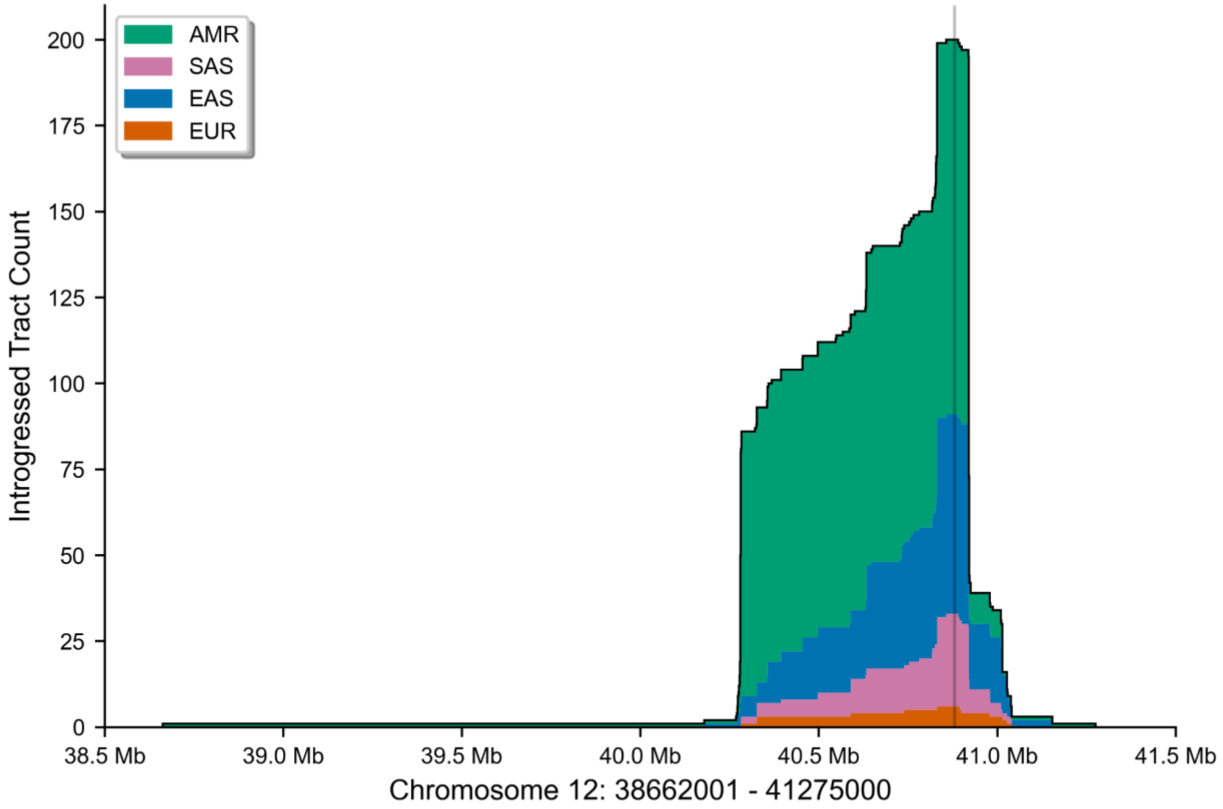


**Fig. S11. Introgressed tracts overlapping the *MUC19* repeat region.**
Density of introgressed tracts that overlap *MUC19* repeat region for the 1KG (black outline) and stratified by super population—Admixed Americans (AMR) in bluish green, South Asians (SAS) in reddish purple, East Asians (EAS) in blue, and Europeans (EUR) in vermillion. The gray shaded region corresponds to the *MUC19* repeat region (hg19, Chr12:40876395-40885001).

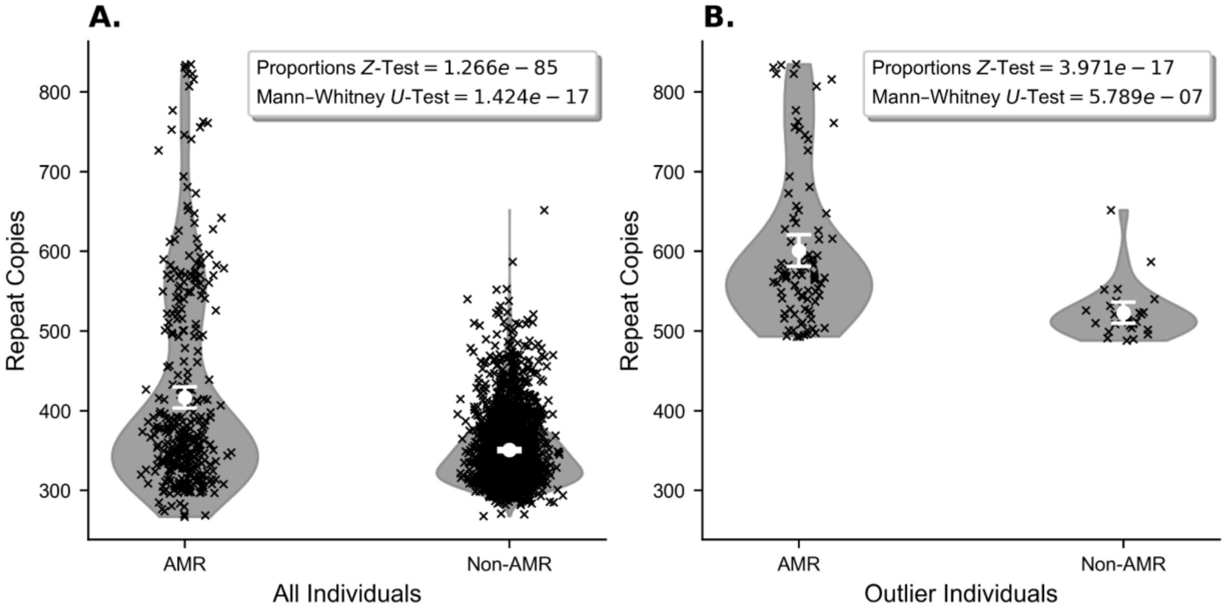


**Fig. S12. Elevated repeat copies among Admixed Americans individuals (AMR) individuals.**

Distributions (represented as gray violin plots) of the estimated number of repeat copies from short-read data for AMR individuals compared to non-AMR individuals—individuals from all other super populations in 1KG. An individual's number of repeat copies is denoted as a black X, the white points represent the mean of the repeat copies distribution, and the white error bars represent the 95% confidence intervals. Panel **(A)** shows the results when considering all individuals in the 1000 Genomes Project (1KG) and demonstrates that between AMR and non-AMR individuals, there is an enrichment of outlier individuals (>487 repeat copies) among the AMR super population (Proportions *Z*-tests) and that AMR individuals have an elevated number of repeat copies (Mann-Whitney *U*-test). Panel **(B)** shows the results when considering only outlier 1KG individuals (>487 repeat copies) and demonstrates that there is an enrichment of outlier individuals within the AMR super population (Proportions *Z*-tests) and that AMR outlier individuals have an elevated number of repeat copies compared to non-AMR outlier individuals (Mann-Whitney *U*-test).

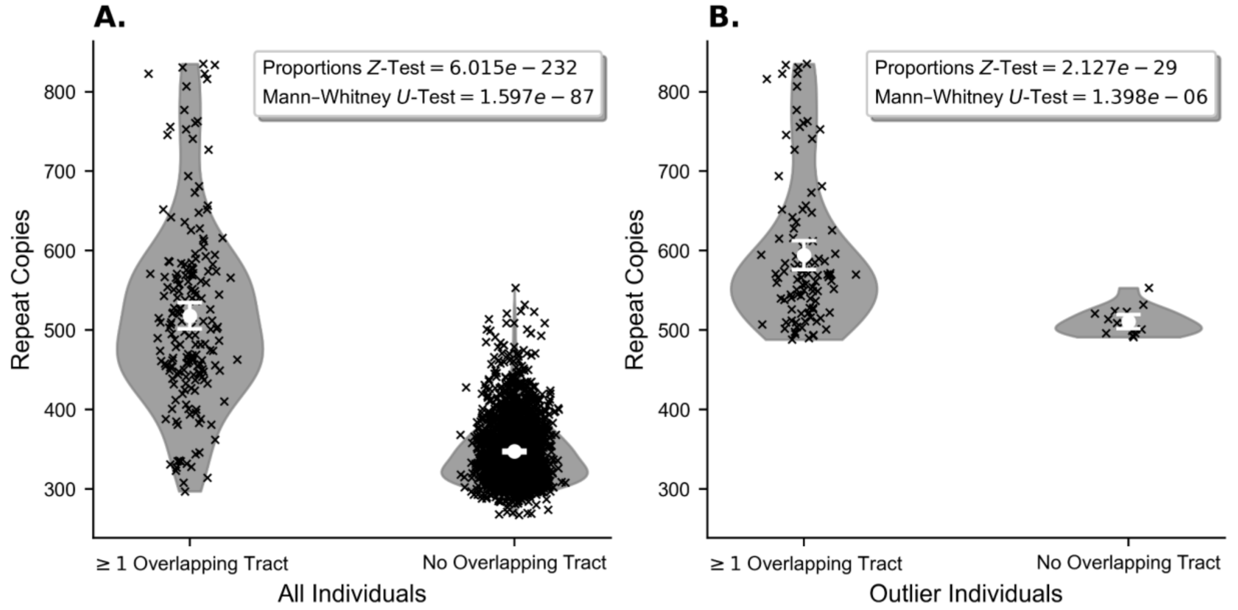


**Fig. S13. Elevated repeat copies among individuals with at least one introgressed tract overlapping the short-read repeat region.**

Distributions (represented as gray violin plots) of the estimated number of repeat copies from short-read data for individuals with one or more introgressed tracts overlapping the repeat region compared to individuals who do not harbor an introgressed tract. An individual's number of repeat copies is denoted as a black X, the white points represent the mean of the repeat copies distribution, and the white error bars represent the 95% confidence intervals. Panel **(A)** shows the results for all individuals in the 1000 Genomes Project (1KG). Note that between individuals with at least one overlapping tract compared to individuals harboring no introgressed tracts, there is an enrichment of outlier individuals (>487 repeat copies) among those with at least one overlapping tract (Proportions *Z*-tests) and that these individuals have an elevated number of repeat copies (Mann-Whitney *U*-test). Panel **(B)** shows the results when considering only outlier 1KG individuals (>487 repeat copies). This panel demonstrates that there is an enrichment of outlier individuals with at least one overlapping tract (Proportions *Z*-tests) and that these outlier individuals have an elevated number of repeat copies compared to outlier individuals who do not harbor an introgressed tract (Mann-Whitney *U*-test).

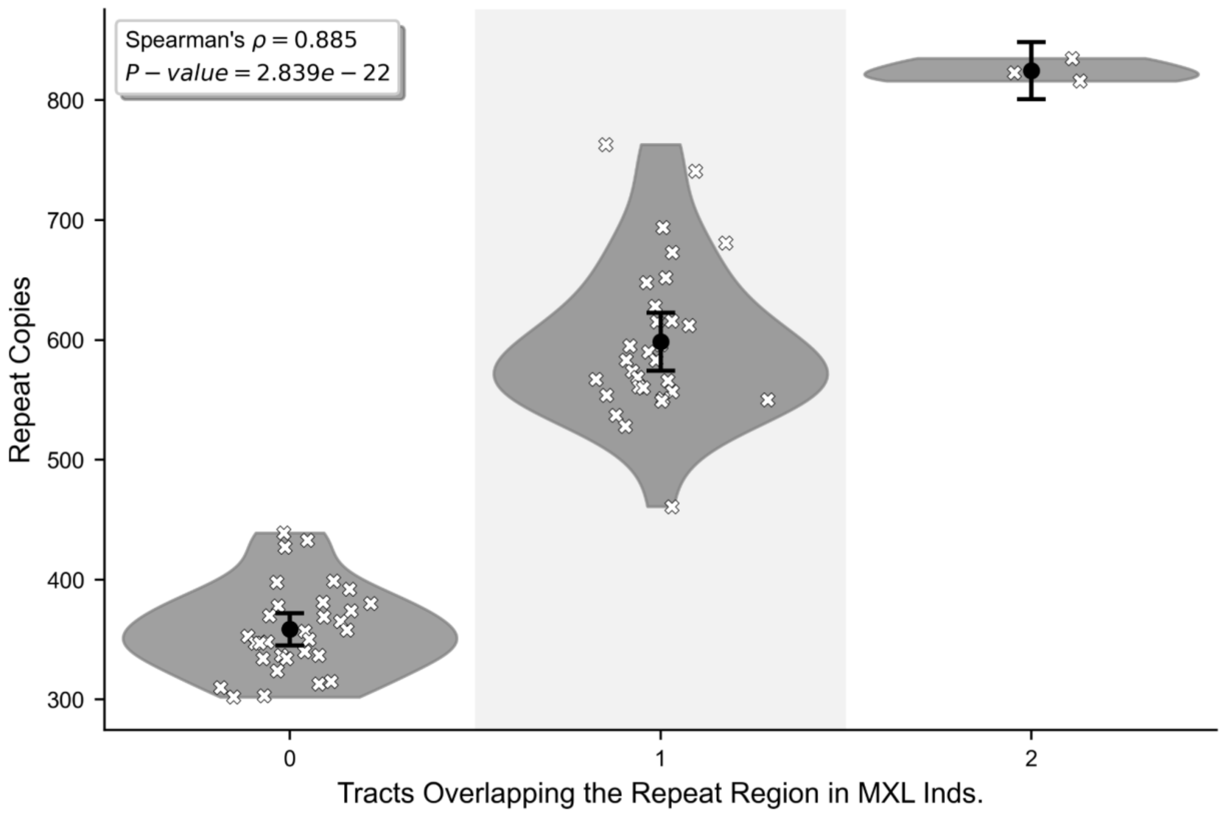


**Fig. S14. Repeat copies are positively correlated with the number of introgressed tracts overlapping the short-read repeat region among MXL individuals.**

Distributions (represented as gray violin plots) of the estimated number of repeat copies from short-read data for MXL individuals partitioned by the number of introgressed tracts overlapping the short-read repeat region. An individual's number of repeat copies is denoted as a white X, the black points represent the mean of the repeat copies distribution, and the black error bars represent the 95% confidence intervals. This figure demonstrates that the number of overlapping tracts is a good predictor of the number of repeat copies for MXL individuals. This suggests that individuals harboring an introgressed tract have an elevated number of repeat copies.

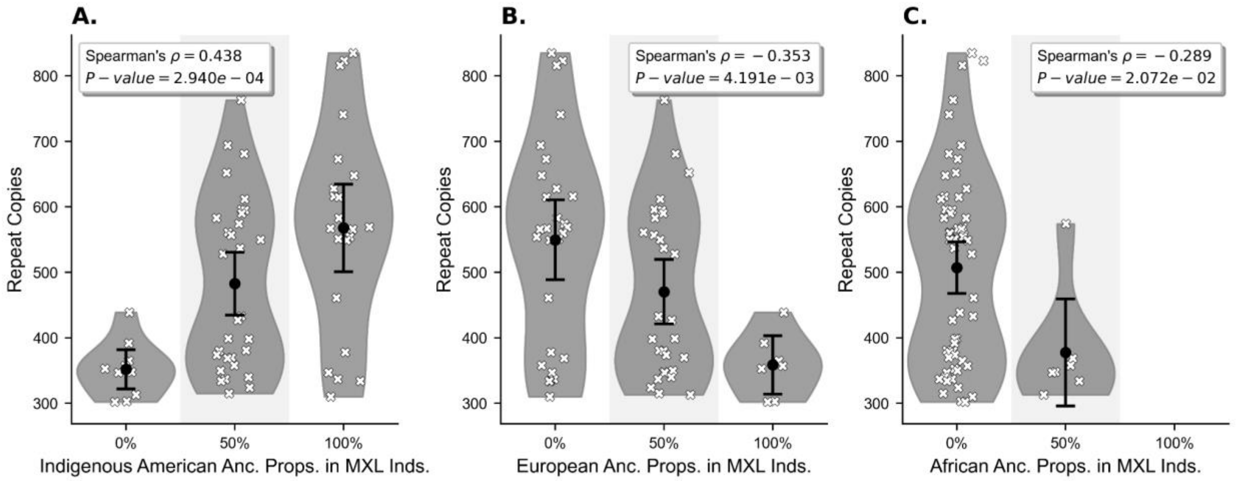


**Fig. S15. The number of repeat copies is positively correlated with the proportion of Indigenous American ancestry in the short-read repeat region among MXL individuals.** Distributions (represented as gray violin plots) of the estimated number of repeat copies from short-read data for MXL individuals partitioned by ancestry component at the short-read repeat region. An individual's number of repeat copies is denoted as a white X, the black points represent the mean of the repeat copies distribution, and the black error bars represent the 95% confidence intervals. The figure demonstrates that the number of repeat copies among MXL individuals are **(A)** positively correlated with an individual's Indigenous American ancestry component, and negatively correlated with **(B)** an individual's European ancestry component, and **(C)** an individual's African ancestry component.

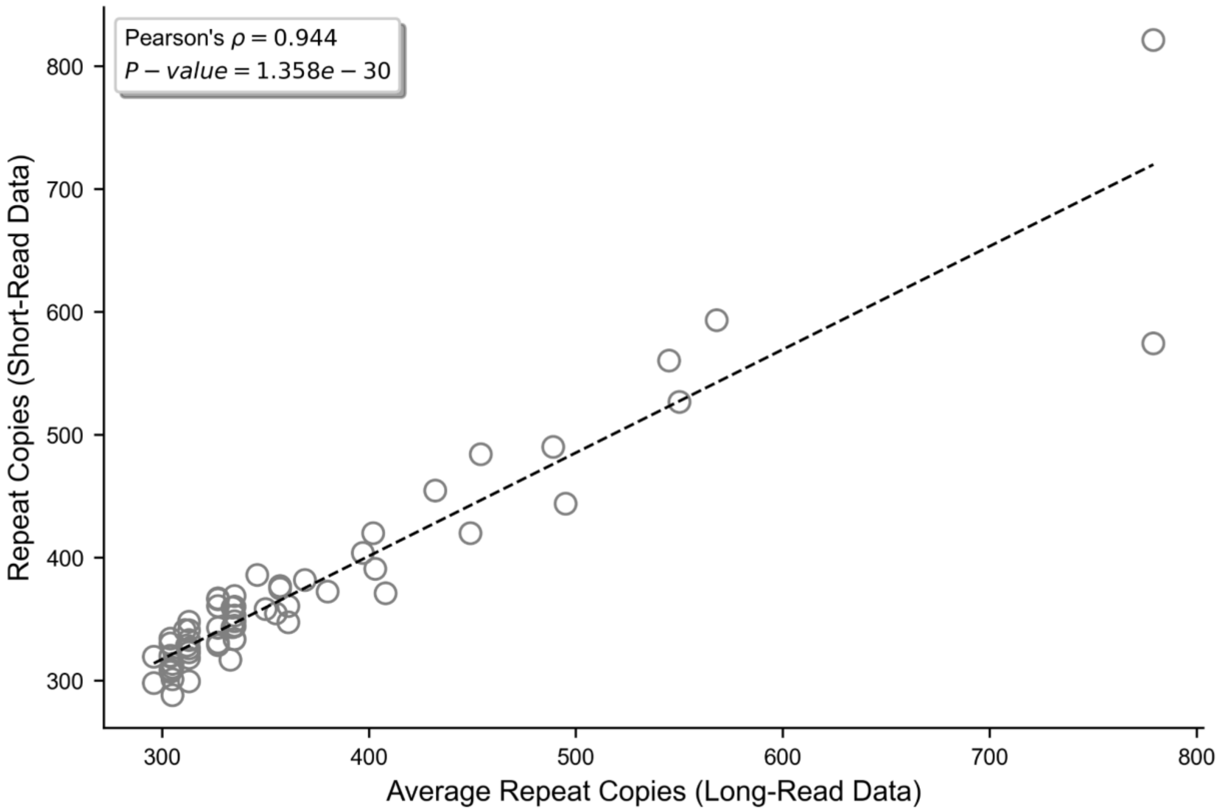


**Fig. S16. The estimated number of repeat copies is correlated between long-read and short-read sequencing technologies.**

The gray circles denote the average number of repeat copies estimated from long-read (x-axis) and short-read (y-axis) sequencing data for the 63 individuals from the Human Pangenome Reference Consortium and the Human Genome Structural Variant Consortium who have publicly available data for both sequencing technologies. The black dashed line denotes the line of best fit from a linear least-squares regression. This figure demonstrates that the estimated number of repeat copies is correlated between different sequencing approaches, indicating that copy number variation analyses are largely robust to sequencing technology.

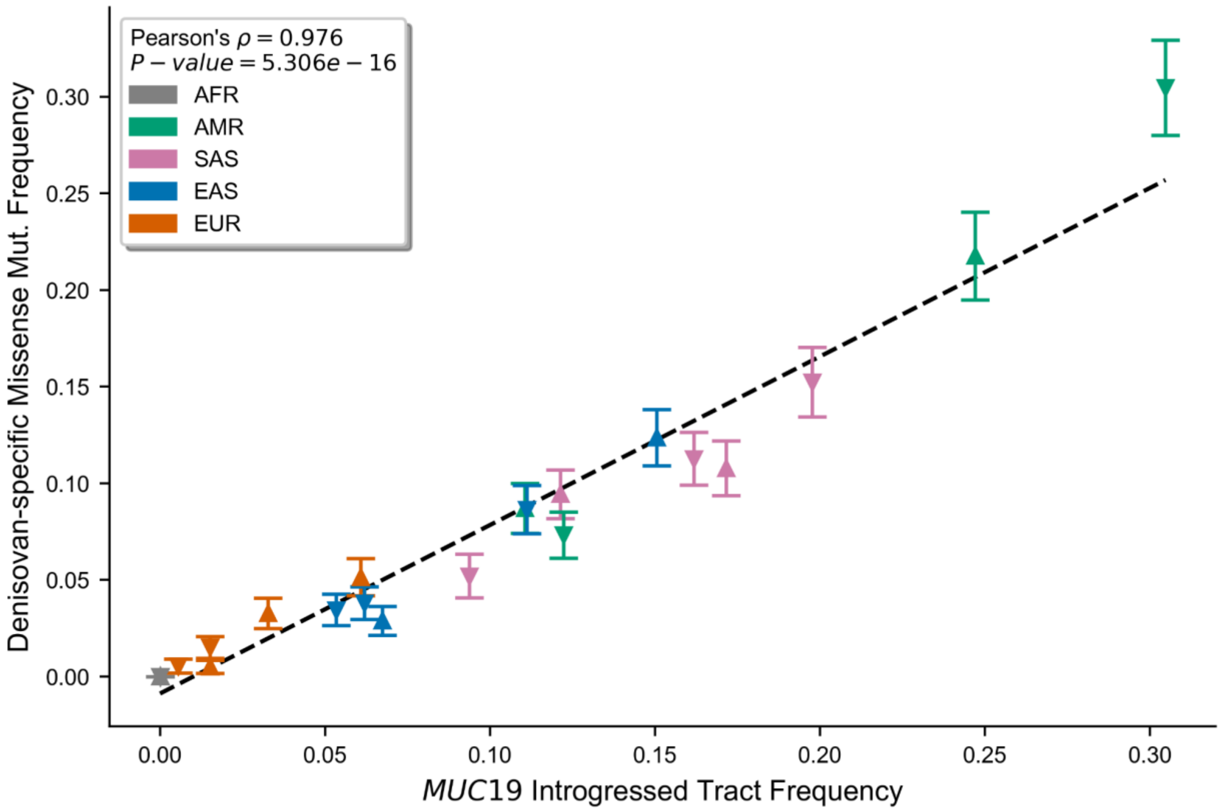


**Fig. S17. The frequency of the Denisovan-specific missense variants is correlated with the frequency of introgressed tracts overlapping *MUC19*.**

Comparison of the frequency of introgressed tracts overlapping the *MUC19* NCBI RefSeq coordinates (x-axis) against the frequency of the nine Denisovan-specific missense variants found within the focal 72kb region (y-axis). Frequencies are stratified by population in the 1KG—African (AFR) populations in gray, Admixed Americans (AMR) in bluish green, South Asians (SAS) in reddish purple, East Asians (EAS) in blue, and Europeans (EUR) in vermillion. Triangles represent the mean Denisovan-specific missense mutation frequency for each population, with error bars indicating 95% confidence intervals. For visual clarity, triangle markers alternate in orientation. The black dashed line denotes the line of best fit from a linear least-squares regression.

| Dataset | Population | Sample Size |
|---|---|---|
| 1000 Genomes Project | Mexico | 64 |
| | Puerto Rico | 104 |
| | Colombia | 94 |
| | Perú | 85 |
| Simons Genome Diversity Project | Pima | 2 |
| | Maya | 2 |
| | Mixe | 3 |
| | Mixtec | 2 |
| | Zapotec | 2 |
| | Piapoco | 2 |
| | Karitiana | 3 |
| | Quechua | 3 |
| | Surui | 2 |
| | Chané | 1 |
| Ancient Americans | Alaska | 1 |
| | Ontario | 1 |
| | Montana | 1 |
| | California | 9 |
| | Querétaro | 3 |
| | Perú | 4 |
| | Patagonia | 4 |

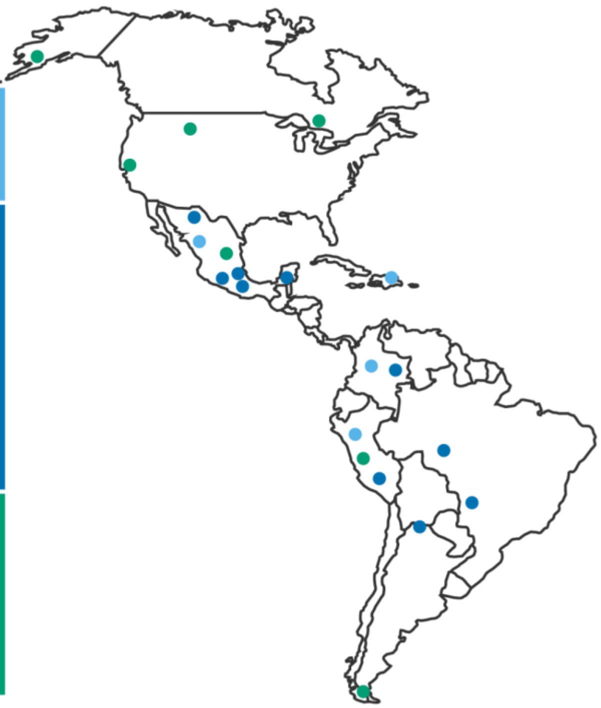

**Fig. S18. Geographic distribution of American individuals used in this study.**
Geographic location and the number of individuals for the modern populations from the 1000 Genomes Project (in sky blue) and Simons Genome Diversity Project (in blue), along with ancient Indigenous American individuals (in bluish green) used in this study.

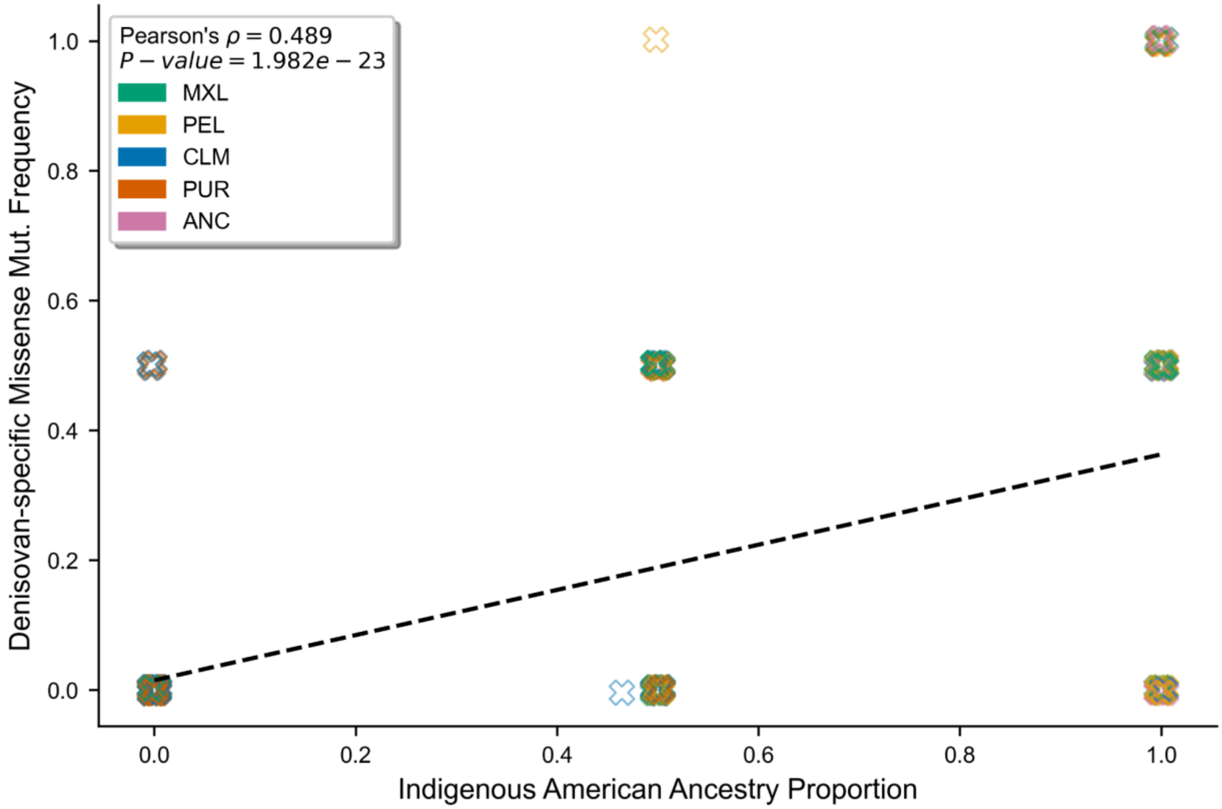


**Fig. S19. The frequency of the Denisovan-specific missense variants is correlated with the Indigenous American Ancestry proportion at the focal 72kb region.**

Comparison of an individual's Indigenous American ancestry proportion at the focal 72kb region (x-axis) against the frequency of the Denisovan-specific missense variant (i.e., 0, 0.5, or 1) at position Chr12:40808726 (y-axis) for all Admixed American individuals in the 1KG and the ancient American individuals. Note that this Denisovan-specific missense variant was chosen given that 20 our of 23 ancient American individuals had genotype information for this position and since all ancient American individuals pre-date colonization we assumed their Indigenous American ancestry proportion is one. The black dashed line denotes the line of best fit from a linear least-squares regression.

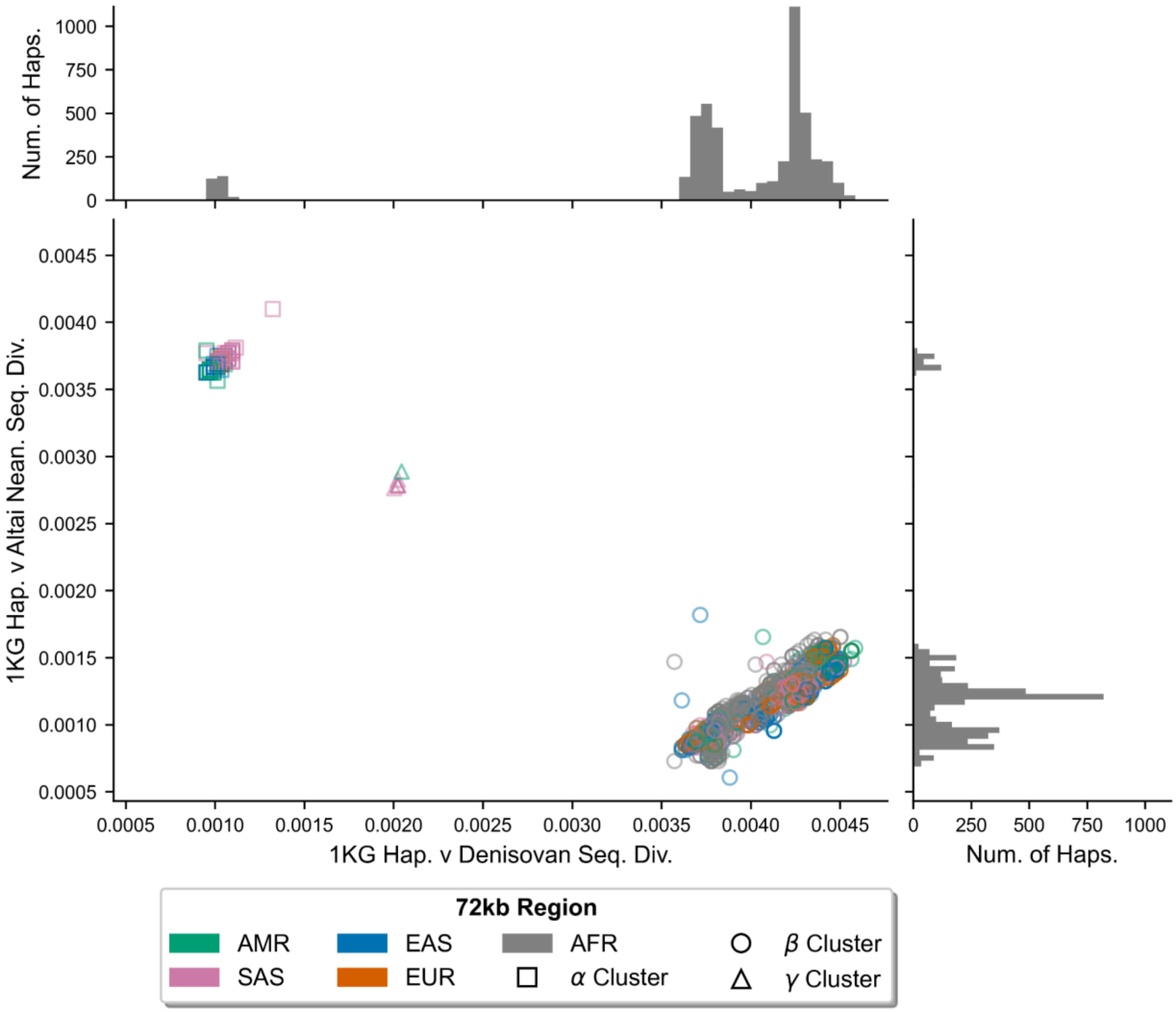


**Fig. S20. Joint distribution of sequence divergence from the Denisovan and Altai Neanderthal at the focal 72kb region.**
The joint distribution of sequence divergence—the number of pairwise differences between a modern human haplotype and an archaic genotype normalized by the effective sequence length—from the Denisovan (x-axis) and the Altai Neanderthal (y-axis) for all haplotypes in the 1000 Genomes Project (1KG) at the focal 72kb region. Points correspond to 1KG haplotypes and are colored according to super population: Africans (AFR) in grey, admixed Americans (AMR) in bluish green, South Asians (SAS) in reddish purple, East Asians (EAS) in blue, and Europeans (EUR) in vermillion. The shape of each point indicates the cluster the haplotype falls within based on the ellipses in Figure 4: squares represent haplotypes within the $\alpha$ cluster, circles represent haplotypes within the $\beta$ cluster, and triangles represent haplotypes within the $\gamma$ cluster. Marginal distributions of sequence divergence from the Denisovan and Altai Neanderthal are depicted by the histograms above and to the right of the joint distribution, respectively. This figure demonstrates that the patterns of haplotype divergence observed in AMR are also observed in every non-African super population. This indicates that all non-African 1KG super populations harbor a *Denisovan-like* haplotype at the focal 72kb region.

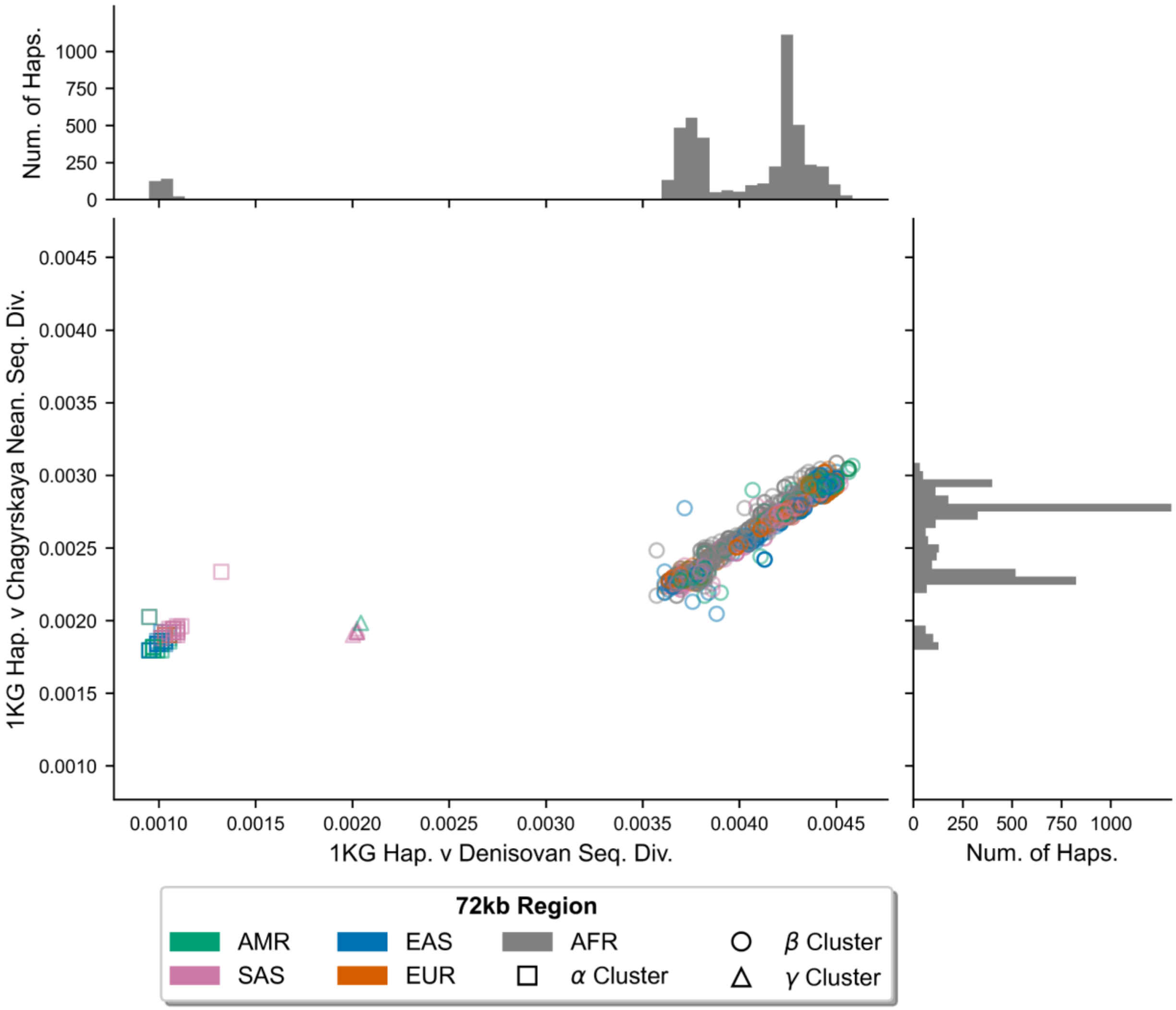


**Fig. S21. Joint distribution of sequence divergence from the Denisovan and Chagyrskaya Neanderthal at the focal 72kb region.**

The joint distribution of sequence divergence—the number of pairwise differences between a modern human haplotype and an archaic genotype normalized by the effective sequence length—from the Denisovan (x-axis) and the Chagyrskaya Neanderthal (y-axis) for all haplotypes in the 1000 Genomes Project (1KG) at the focal 72kb region. Points correspond to 1KG haplotypes and are colored according to super population: Africans (AFR) in grey, admixed Americans (AMR) in bluish green, South Asians (SAS) in reddish purple, East Asians (EAS) in blue, and Europeans (EUR) in vermillion. The shape of each point indicates the cluster the haplotype falls within based on the ellipses in Figure 4: squares represent haplotypes within the $\alpha$ cluster, circles represent haplotypes within the $\beta$ cluster, and triangles represent haplotypes within the $\gamma$ cluster. Marginal distributions of sequence divergence from the Denisovan and Chagyrskaya Neanderthal are depicted by the histograms above and to the right of the joint distribution, respectively. This figure demonstrates that the patterns of haplotype divergence observed in AMR are also observed in every non-African super population. This indicates that all non-African 1KG super populations harbor a *Denisovan-like* haplotype at the focal 72kb region.

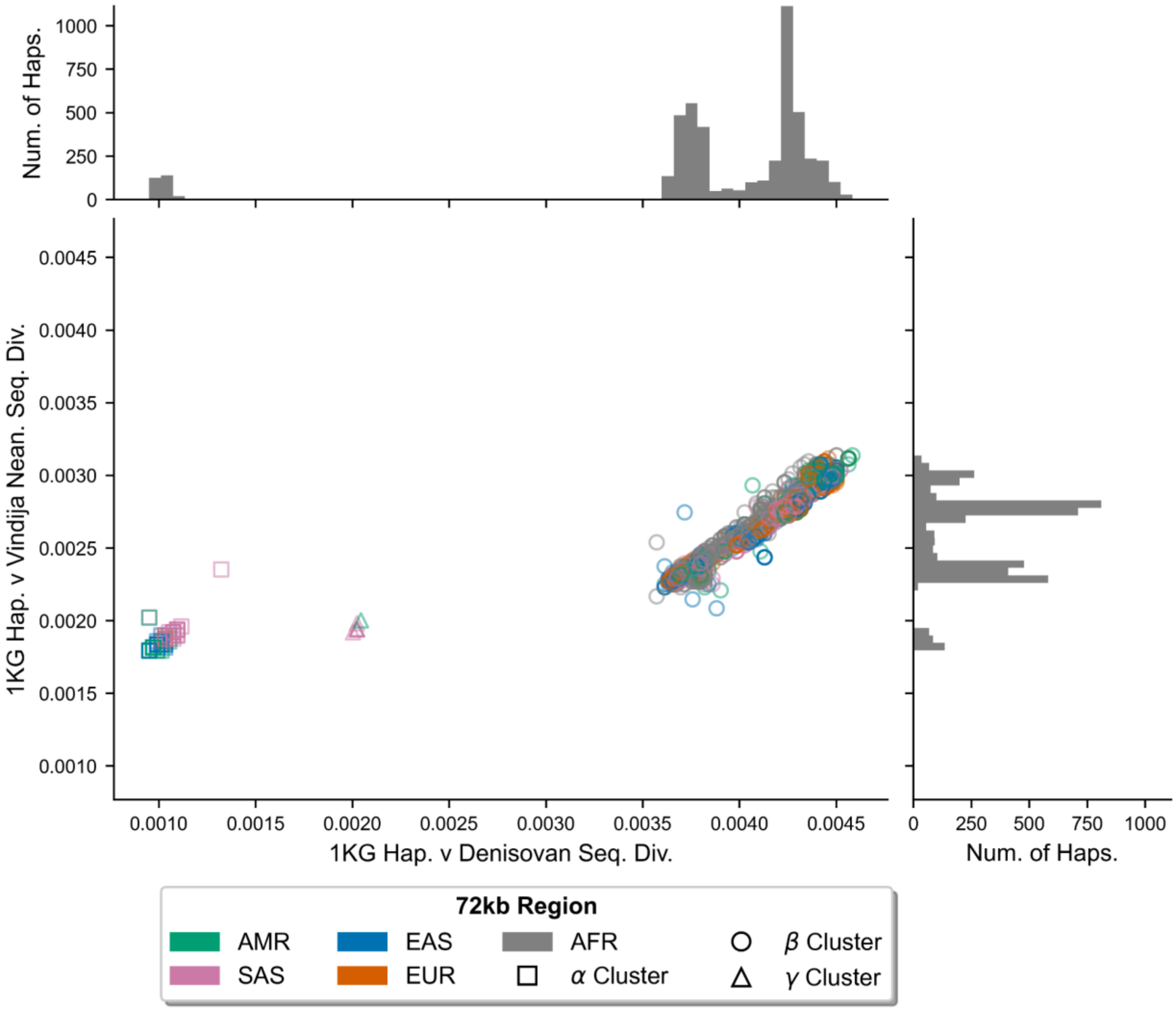


**Fig. S22. Joint distribution of sequence divergence from the Denisovan and Vindija Neanderthal at the focal 72kb region.**

The joint distribution of sequence divergence—the number of pairwise differences between a modern human haplotype and an archaic genotype normalized by the effective sequence length—from the Denisovan (x-axis) and the Vindija Neanderthal (y-axis) for all haplotypes in the 1000 Genomes Project (1KG) at the focal 72kb region. Points correspond to 1KG haplotypes and are colored according to super population: Africans (AFR) in grey, admixed Americans (AMR) in bluish green, South Asians (SAS) in reddish purple, East Asians (EAS) in blue, and Europeans (EUR) in vermillion. The shape of each point indicates the cluster the haplotype falls within based on the ellipses in Figure 4: squares represent haplotypes within the $\alpha$ cluster, circles represent haplotypes within the $\beta$ cluster, and triangles represent haplotypes within the $\gamma$ cluster. Marginal distributions of sequence divergence from the Denisovan and Vindija Neanderthal are depicted by the histograms above and to the right of the joint distribution, respectively. This figure demonstrates that the patterns of haplotype divergence observed in AMR are also observed in every non-African super population. This indicates that all non-African 1KG super populations harbor a *Denisovan-like* haplotype at the focal 72kb region.

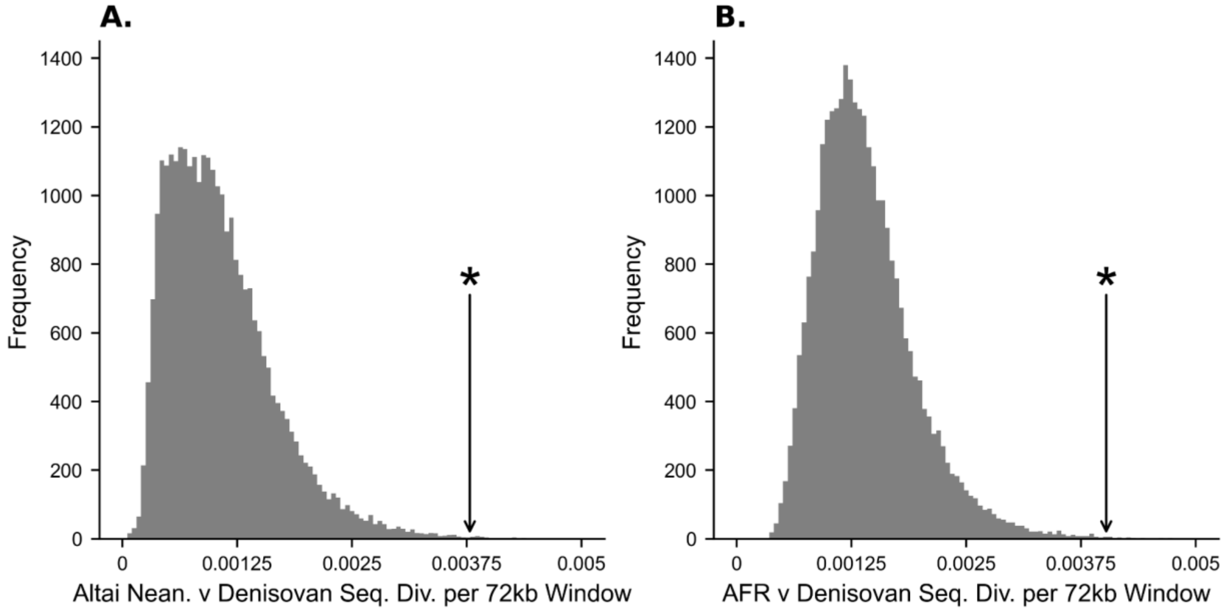


**Fig. S23. The Altai Neanderthal and all Africans in the 1000 Genomes Project (1KG) exhibit elevated sequence divergence from the Denisovan at the focal 72kb region.** Distributions of sequence divergence from the Denisovan for non-overlapping 72kb windows: **(A)** the number of pairwise differences between the Altai Neanderthal's and Denisovan's unphased genotypes normalized by the effective sequence length and **(B)** the average number of pairwise differences between all African haplotypes in the 1KG and the Denisovan's unphased genotypes normalized by the effective sequence length. Arrows indicate the observed sequence divergence between the Denisovan and **(A)** the Altai Neanderthal or **(B)** all Africans in the 1KG at the focal 72kb region. Asterisks above arrows indicate statistical significance when compared to the genome-wide distribution of non-overlapping windows; "ns" denotes non-significance. The figure demonstrates that the Altai Neanderthal and all Africans have a significantly elevated sequence divergence from the Denisovan at the focal 72kb region. This indicates that this shared elevated sequence divergence can best explain the observed affinity Africans have for the Altai Neanderthal, as no African haplotype is significantly closer than expected to any Neanderthal at the unphased or phased 72kb region.

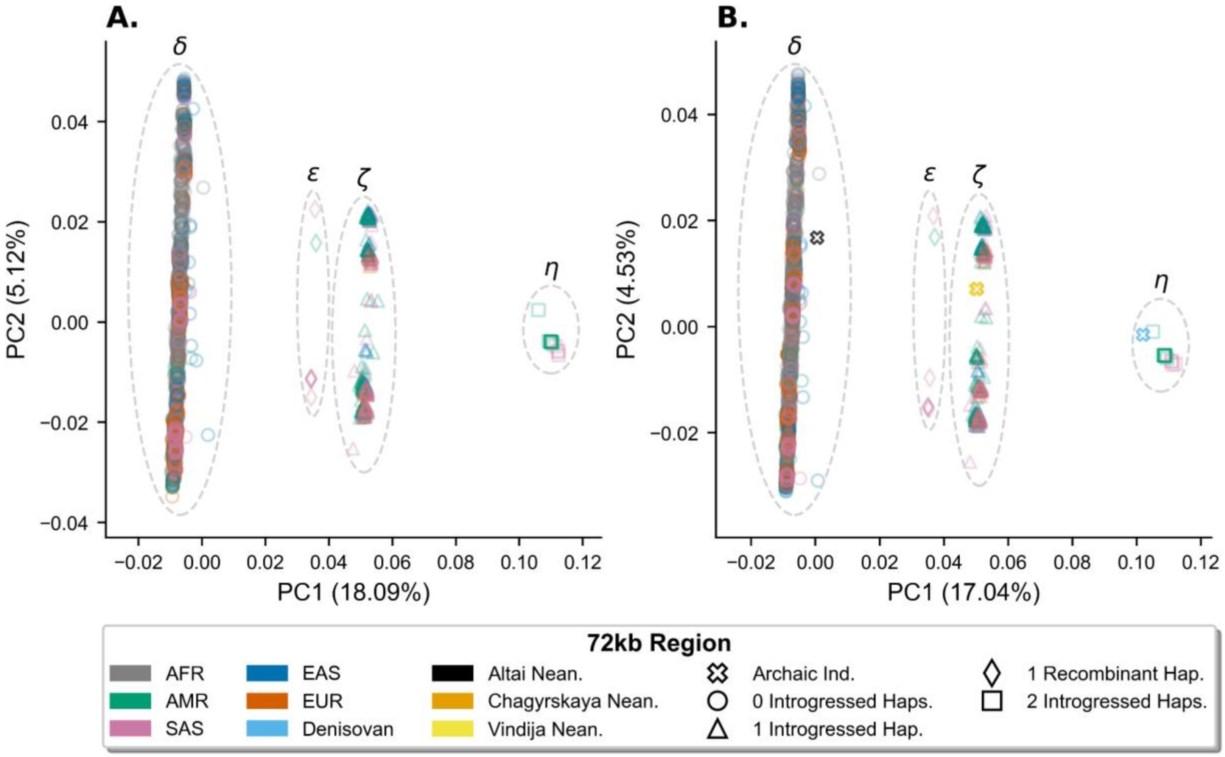


**Fig. S24. PCA reveals haplotype structure at the focal 72kb region.**

Principal Component Analysis (PCA) of the focal 72kb region considering **(A)** only 1KG individuals and **(B)** both 1KG individuals and the four archaic individuals. For each PCA, we plot the first two principal components (PCs) and report the percentage of the total variance explained by each PC. Points correspond to individuals and are colored according to super population: Africans (AFR) in grey, admixed Americans (AMR) in bluish green, South Asians (SAS) in reddish purple, East Asians (EAS) in blue, and Europeans (EUR) in vermillion. The shape of each point indicates the number of Denisovan-like haplotypes a 1KG individual harbors at the 72kb region: circles represent individuals with no introgressed haplotypes, triangles represent individuals with one introgressed haplotype, diamonds represent individuals with one recombinant haplotype, and squares represent individuals with two introgressed haplotypes. Additionally, in panel **(B)**, the Denisovan, Altai Neanderthal, Chagyrskaya Neanderthal, and Vindija Neanderthal are indicated by sky blue X, black X, orange X, and yellow X, respectively. The four grey ellipses highlight the four haplotype groups among the 1KG individuals. The δ ellipse corresponds to individuals lacking the introgressed haplotype at the 72kb region—note these individuals harbor two haplotypes in the $\beta$ ellipse in Figure 4 of the main text. The ε ellipse corresponds to individuals with one recombinant haplotype at the 72kb region—these individuals have a single haplotype in the $\gamma$ ellipse in Figure 4 of the main text. The ζ ellipse corresponds to individuals with one introgressed haplotype at the 72kb region—these individuals have a single haplotype in the $\alpha$ ellipse in Figure 4 of the main text. Lastly, the η ellipse corresponds to individuals with two introgressed haplotypes at the 72kb region—these individuals have two haplotypes in the $\alpha$ ellipse in Figure 4 of the main text. Panel **(A)** demonstrates that PCA recovers the observed haplotype structure at the 72kb region even when the archaic individuals are not included in the analysis. Panel **(B)** further validates these observations as the Altai Neanderthal (black X), which harbors no Denisovan-like haplotypes, falls within the δ cluster; the two late Neanderthals (orange and yellow Xs), each harboring one Denisovan-like haplotype, fall within the ζ cluster; and the Denisovan (sky blue X) falls within the η cluster. Note that due to their high similarity at the 72kb region, the Chagyrskaya and Vindija Neanderthal points overlap in panel **(B)**.

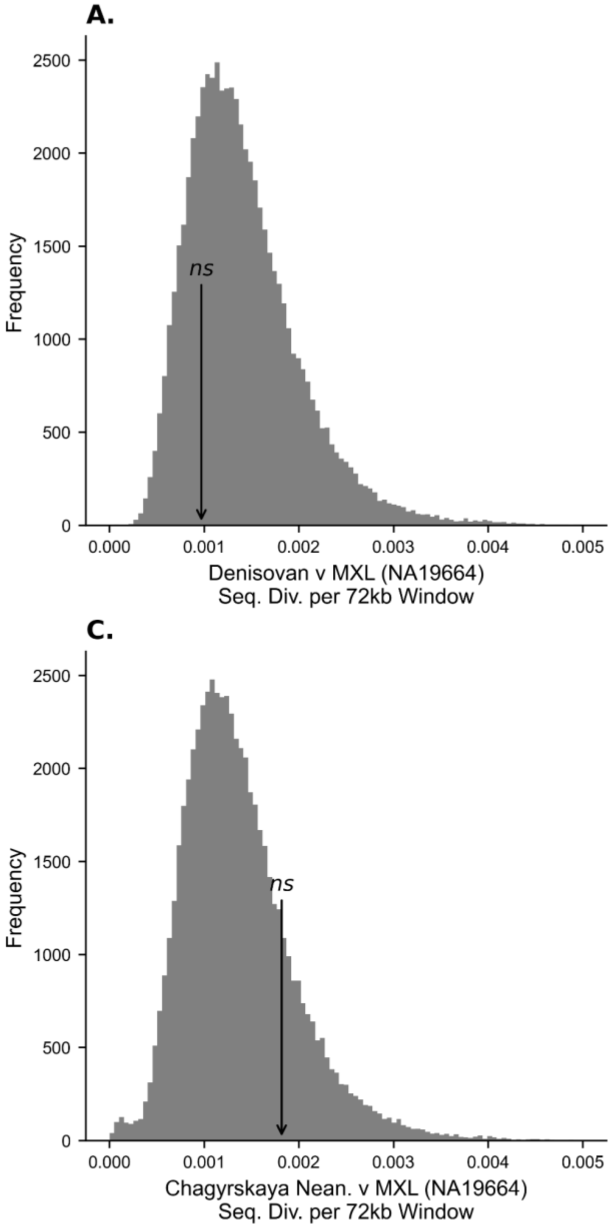


**Fig. S25. The *Denisovan-like* haplotype in MXL is not closer than expected to any of the unphased archaic individuals.**

Distributions of sequence divergence, i.e., the number of pairwise differences between the MXL individual's haplotypes (NA19664), who harbors two *Denisovan-like* haplotypes and has no heterozygous sites at the focal 72kb region, and the unphased archaic genotypes normalized by the effective sequence length, for non-overlapping 72kb windows. Arrows indicate the observed sequence divergence between the *Denisovan-like* haplotype in MXL and the **(A)** Denisovan, **(B)** Altai Neanderthal, **(C)** Chagyrskaya Neanderthal, and **(D)** Vindija Neanderthal at the focal 72kb region. Asterisks above arrows indicate statistical significance when compared to the genome-wide distribution of non-overlapping windows; "ns" denotes non-significance. The figure demonstrates that the *Denisovan-like* haplotype in MXL is not closer than expected to any of the unphased archaic individuals.

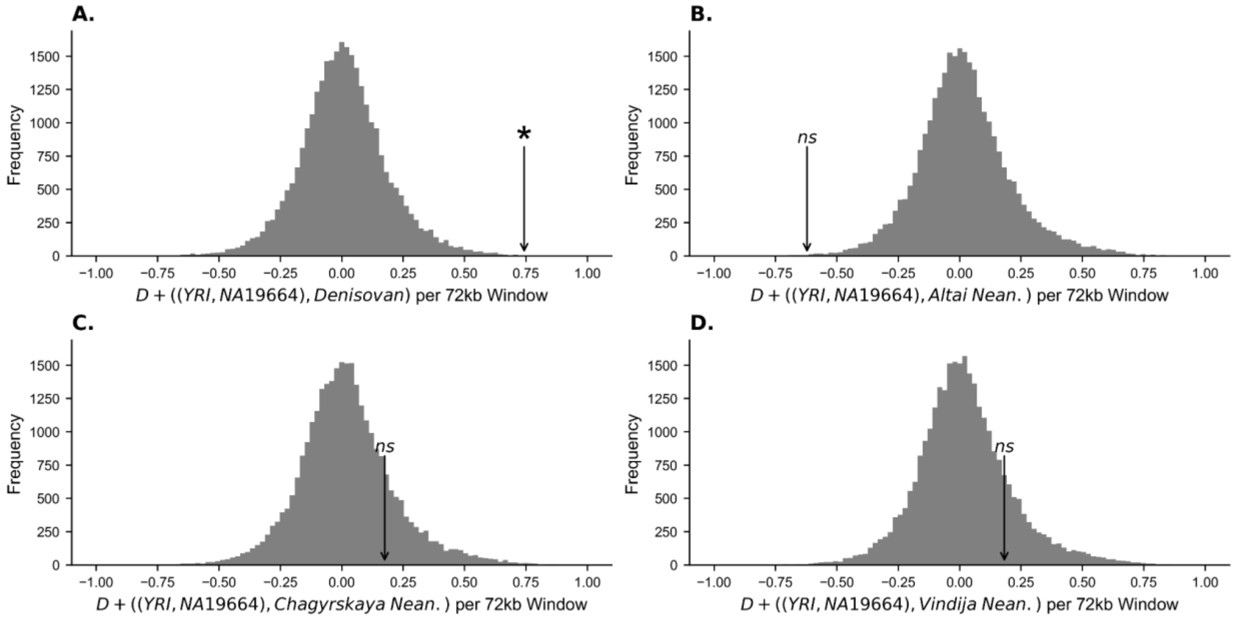


**Fig. S26. Excess of allele sharing between the introgressed haplotype in MXL and the sequenced Denisovan at the focal 72kb region.**

Distributions of *D*+ tests of introgression using the YRI population as *P1*, the focal MXL individual (NA19664) with two copies of the introgressed haplotype with an affinity to the Altai Denisovan as *P2*, the four high-coverage archaic individuals as *P3*, and the *EPO* ancestral sequence as *P4* for non-overlaping 72kb windows. Arrows indicate the observed sequence *D*+ value at the focal 72kb region when: **(A)** *P3* = Denisovan, **(B)** *P3* = Altai Neanderthal*,* **(C)** *P3* = Chagyrskaya Neanderthal*,* and **(D)** *P3* = Vindija Neanderthal. Asterisks above arrows indicate statistical significance when compared to the genome-wide distribution of non-overlapping windows; "ns" denotes non-significance. Panel **(A)** shows that the introgressed haplotype in MXL exhibits the largest positive and significant *D*+ value relative to any other *P2* population when *P3* = Denisovan. Panels **(B)** through **(D)** demonstrate that there is no evidence for an excess of allele sharing between any non-African population when **(B)** *P3* = Altai Neanderthal, **(C)** *P3* = Chagyrskaya Neanderthal, or **(D)** *P3* = Vindija Neanderthal.

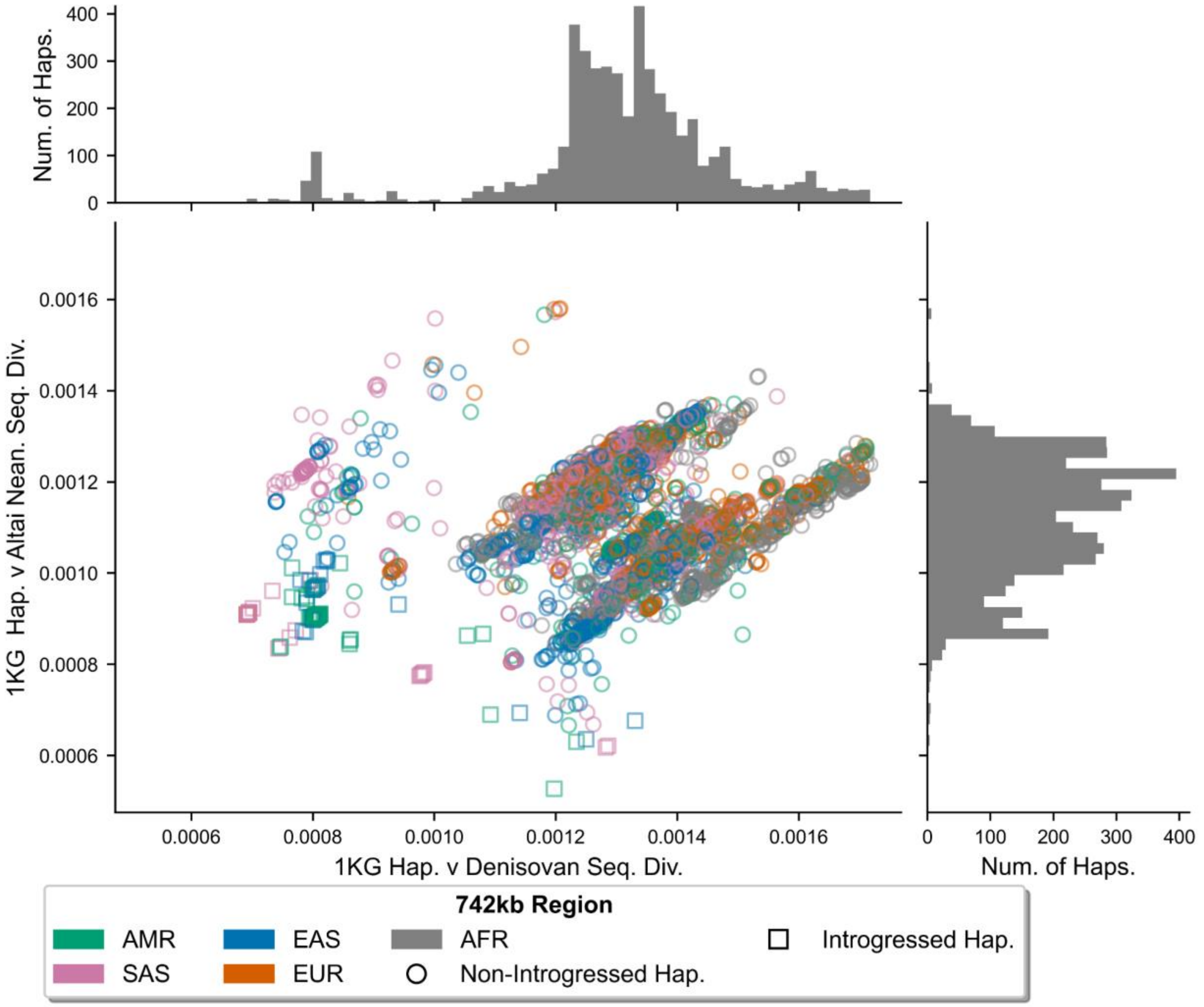


**Fig. S27. Joint distribution of sequence divergence from the Denisovan and Altai Neanderthal at the focal 742kb region.**

The joint distribution of sequence divergence—the number of pairwise differences between a modern human haplotype and an archaic genotype normalized by the effective sequence length—from the Denisovan (x-axis) and the Altai Neanderthal (y-axis) for all haplotypes in the 1000 Genomes Project (1KG) at the focal 742kb region. Points correspond to 1KG haplotypes and are colored according to super population: Africans (AFR) in grey, admixed Americans (AMR) in bluish green, South Asians (SAS) in reddish purple, East Asians (EAS) in blue, and Europeans (EUR) in vermillion. The shape of each point indicates whether the 1KG haplotype is significantly closer to a late Neanderthal after correcting for multiple comparisons: circles represent non-introgressed haplotypes, and circles represent introgressed haplotypes. Marginal distributions of sequence divergence from the Denisovan and Altai Neanderthal are depicted by the histograms above and to the right of the joint distribution, respectively.

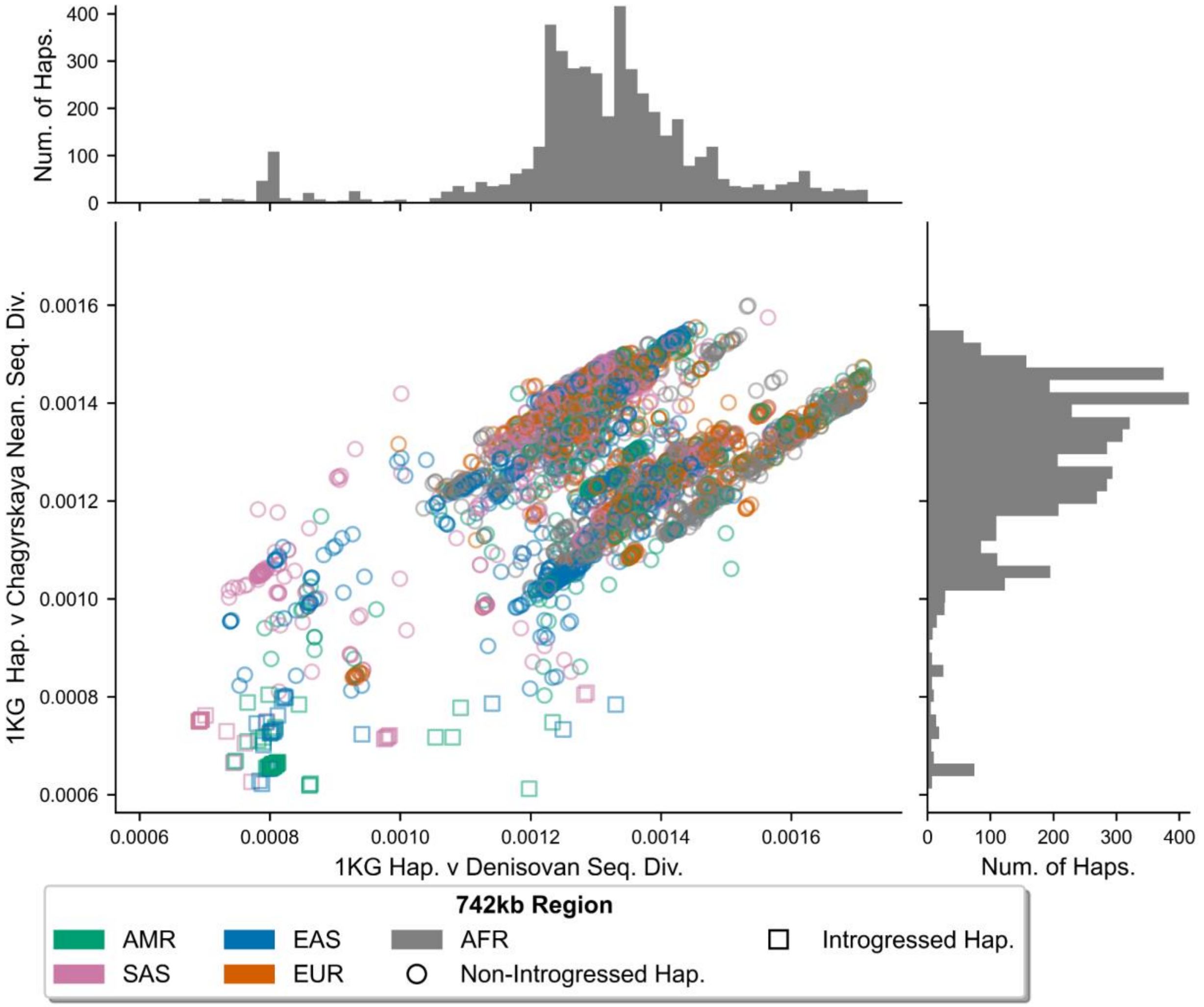


**Fig. S28. Joint distribution of sequence divergence from the Denisovan and Chagyrskaya Neanderthal at the focal 742kb region.**

The joint distribution of sequence divergence—the number of pairwise differences between a modern human haplotype and an archaic genotype normalized by the effective sequence length—from the Denisovan (x-axis) and the Chagyrskaya Neanderthal (y-axis) for all haplotypes in the 1000 Genomes Project (1KG) at the focal 742kb region. Points correspond to 1KG haplotypes and are colored according to super population: Africans (AFR) in grey, admixed Americans (AMR) in bluish green, South Asians (SAS) in reddish purple, East Asians (EAS) in blue, and Europeans (EUR) in vermillion. The shape of each point indicates whether the 1KG haplotype is significantly closer to a late Neanderthal after correcting for multiple comparisons: circles represent non-introgressed haplotypes, and circles represent introgressed haplotypes. Marginal distributions of sequence divergence from the Denisovan and Chagyrskaya Neanderthal are depicted by the histograms above and to the right of the joint distribution, respectively.

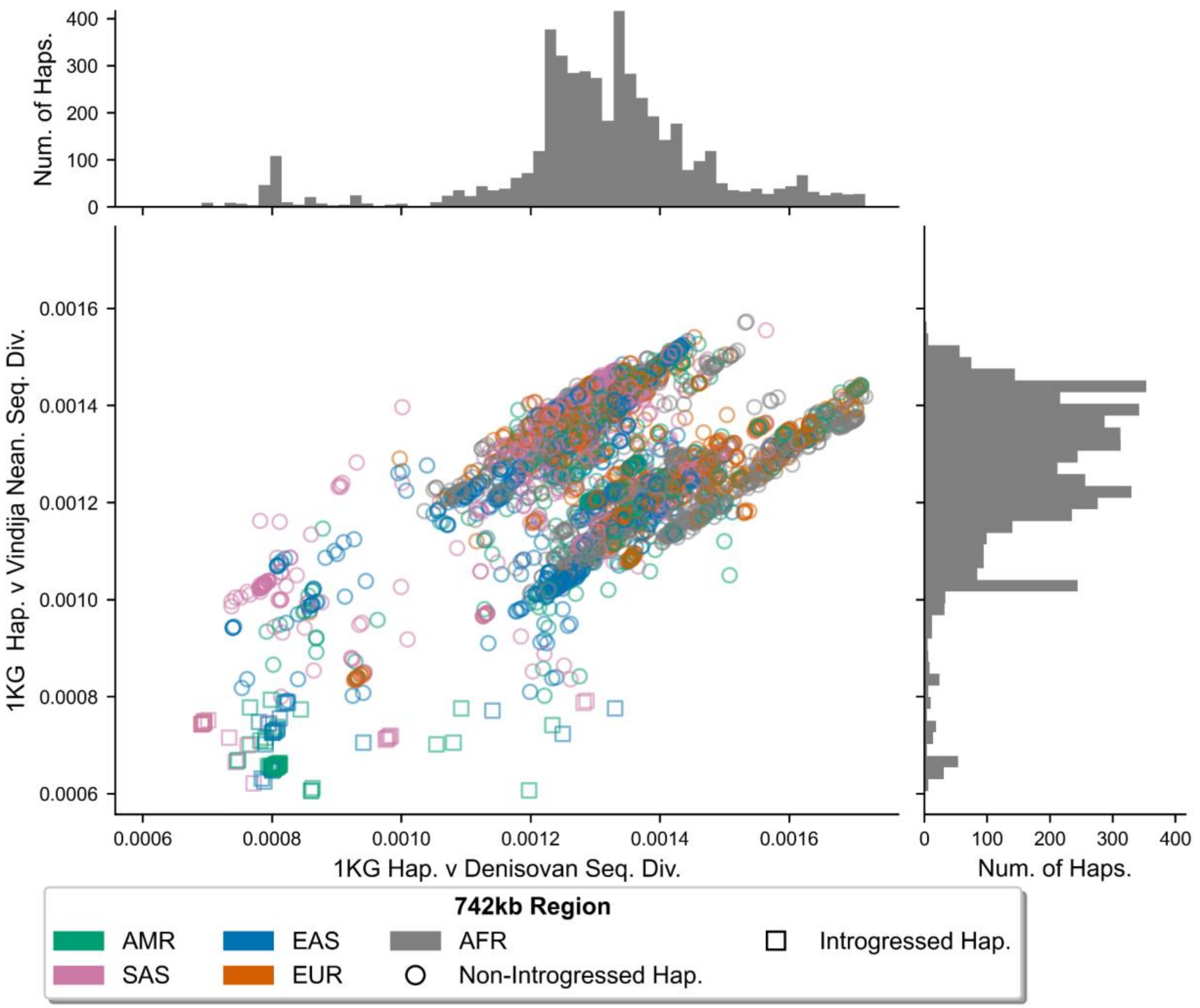


**Fig. S29. Joint distribution of sequence divergence from the Denisovan and Vindija Neanderthal at the focal 742kb region.**

The joint distribution of sequence divergence—the number of pairwise differences between a modern human haplotype and an archaic genotype normalized by the effective sequence length—from the Denisovan (x-axis) and the Vindija Neanderthal (y-axis) for all haplotypes in the 1000 Genomes Project (1KG) at the focal 742kb region. Points correspond to 1KG haplotypes and are colored according to super population: Africans (AFR) in grey, admixed Americans (AMR) in bluish green, South Asians (SAS) in reddish purple, East Asians (EAS) in blue, and Europeans (EUR) in vermillion. The shape of each point indicates whether the 1KG haplotype is significantly closer to a late Neanderthal after correcting for multiple comparisons: circles represent non-introgressed haplotypes, and circles represent introgressed haplotypes. Marginal distributions of sequence divergence from the Denisovan and Vindija Neanderthal are depicted by the histograms above and to the right of the joint distribution, respectively.

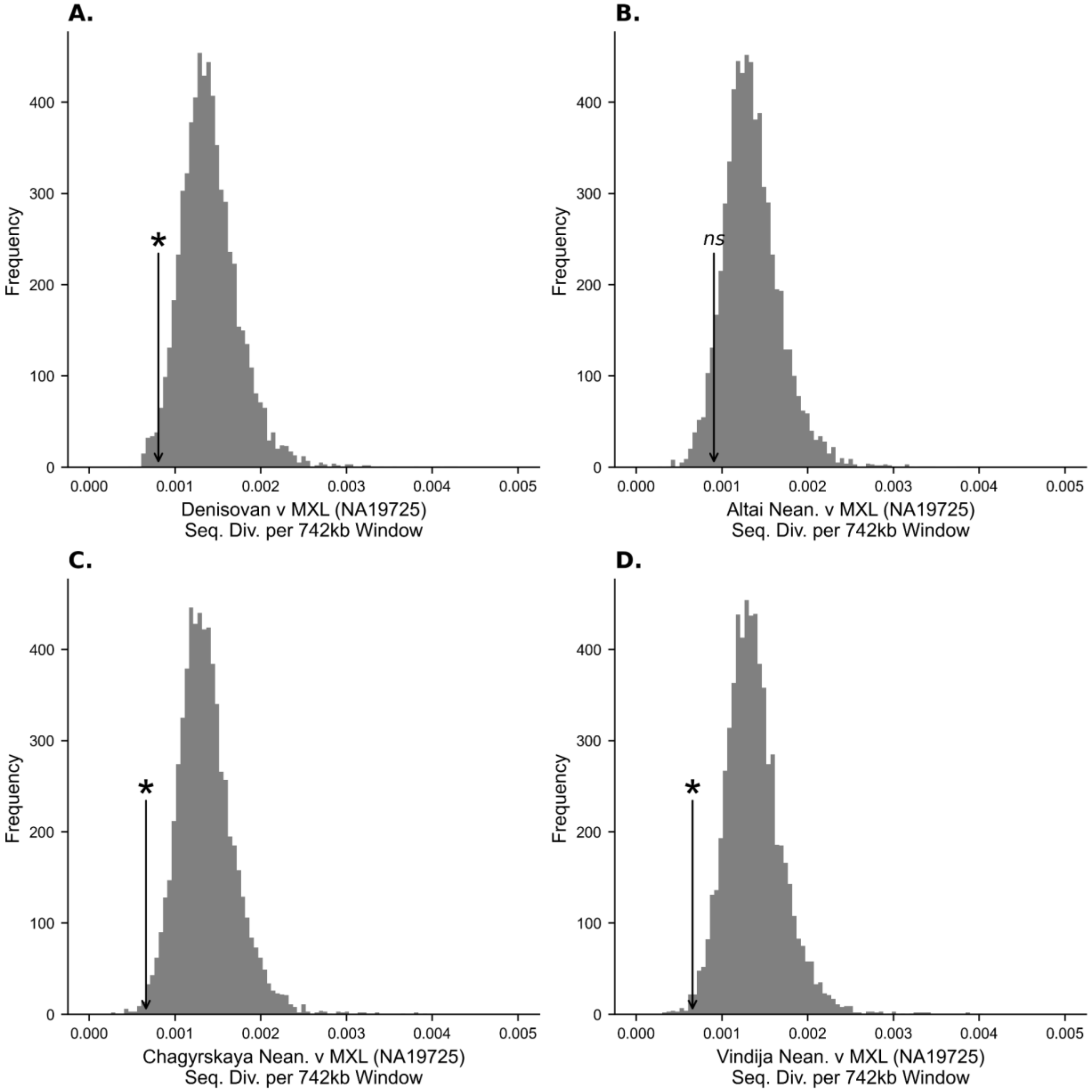


**Fig. S30. The longest introgressed tract in MXL is closer than expected to the Denisovan and the late Neanderthals.**
Distributions of sequence divergence, i.e., the number of pairwise differences between the MXL individual's haplotypes (NA19725), who harbors the 742kb longest introgressed tract overlapping *MUC19*, and the unphased archaic genotypes normalized by the effective sequence length, for non-overlapping 742kb windows. Arrows indicate the observed sequence divergence between the longest introgressed tract overlapping *MUC19* in MXL and the **(A)** Denisovan, **(B)** Altai Neanderthal, **(C)** Chagyrskaya Neanderthal, and **(D)** Vindija Neanderthal at the focal 742kb region. Asterisks above arrows indicate statistical significance when compared to the genome-wide distribution of non-overlapping windows; "ns" denotes non-significance. Panels **(A)**, **(C)**,

and **(D)** demonstrate that the longest introgressed tract in MXL is closer than expected to the Denisovan and both of the late Neanderthals.

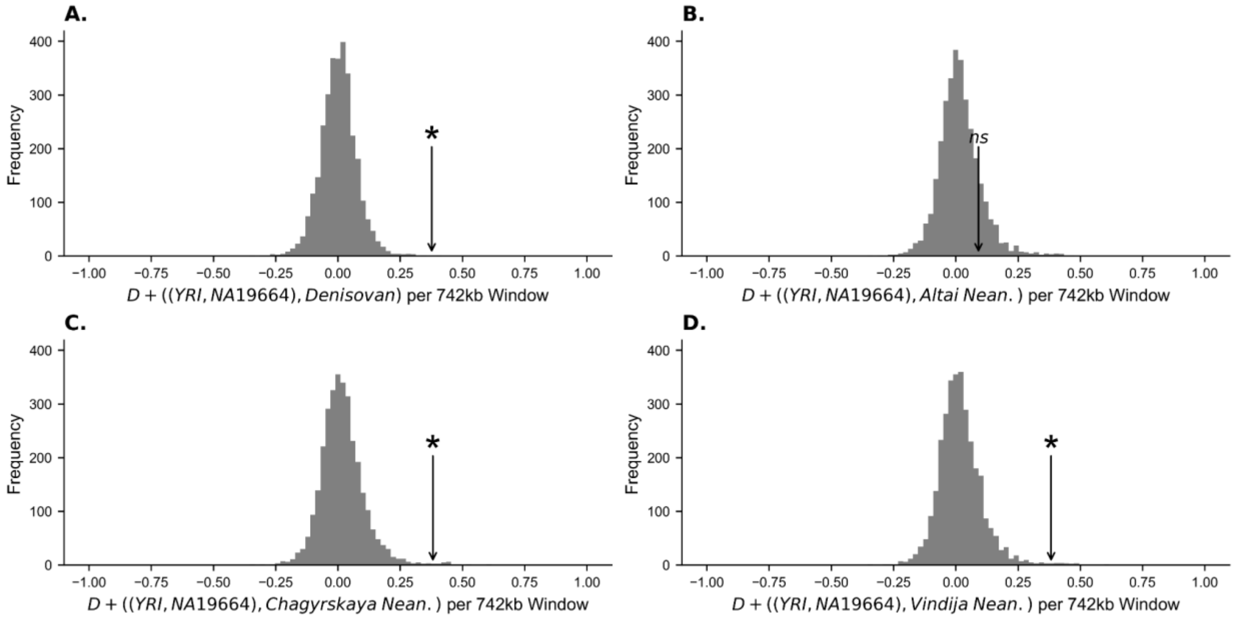


**Fig. S31. Excess of allele sharing between the introgressed haplotype in MXL and the sequenced Denisovan and late Neanderthals at the focal 742kb region.**

Distributions of *D*+ tests of introgression using the YRI population as *P1*, the focal MXL individual (NA19664) with two copies of the introgressed haplotype with an affinity to the Altai Denisovan as *P2*, the four high-coverage archaic individuals as *P3*, and the *EPO* ancestral sequence as *P4* for non-overlaping 742kb windows. Arrows indicate the observed sequence *D*+ value at the focal 742kb region when: **(A)** *P3* = Denisovan, **(B)** *P3* = Altai Neanderthal, **(C)** *P3* = Chagyrskaya Neanderthal, and **(D)** *P3* = Vindija Neanderthal. Asterisks above arrows indicate statistical significance when compared to the genome-wide distribution of non-overlapping windows; "ns" denotes non-significance. Panel **(A)** demonstrates that the introgressed haplotype in MXL exhibits the largest positive and significant *D*+ value relative to any other *P2* population when *P3* = Denisovan. Panel **(B)** demonstrates that there is no evidence for an excess of allele sharing when *P3* = Altai Neanderthal, while panels **(C)** through **(D)** demonstrate that the introgressed haplotype in MXL also exhibits a large positive and significant *D*+ values when **(C)** *P3* = Chagyrskaya Neanderthal or **(D)** *P3* = Vindija Neanderthal.

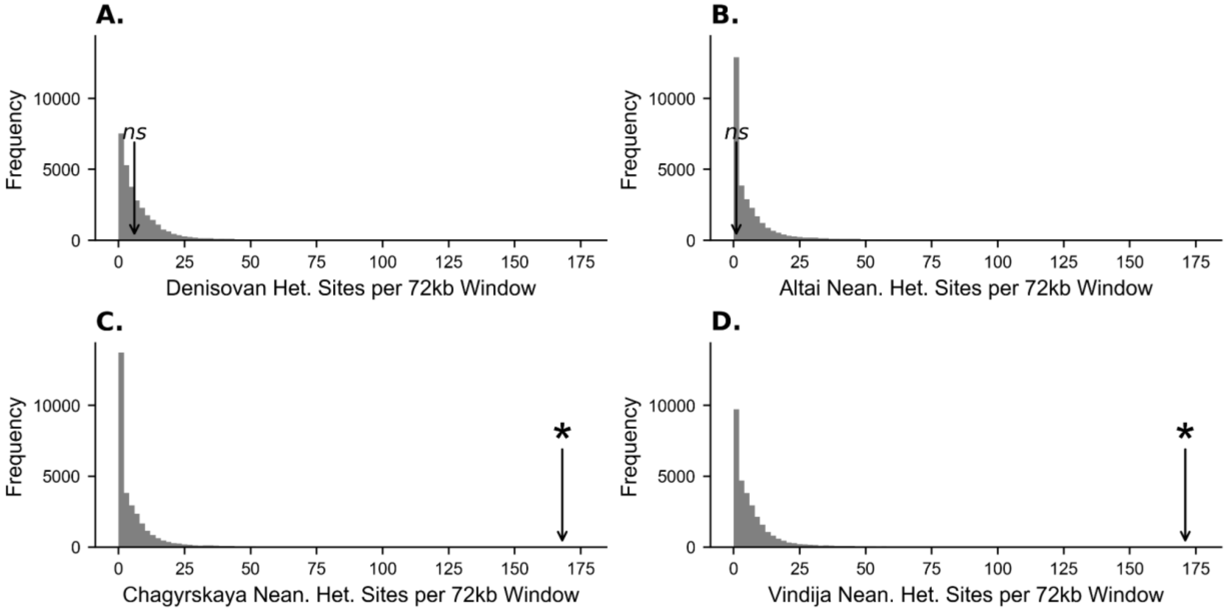


**Fig. S32. The Chagyrskaya and Vindija Neanderthals harbor an elevated number of heterozygous sites at the focal 72kb region.**
Distribution of the number of heterozygous in the **(A)** Denisovan, **(B)** Altai Neanderthal, **(C)** Chagyrskaya Neanderthal, and **(D)** Vindija Neanderthal genomes for non-overlapping 72kb windows. Arrows indicate the observed number of heterozygous sites at the focal 72kb region. Asterisks above arrows indicate statistical significance when compared to the genome-wide distribution of non-overlapping windows; "ns" denotes non-significance. Panels **(A)** and **(B)** demonstrate that number of heterozygous sites observed in the Denisovan and Altai Neanderthal are within the genome-wide expectation for 72kb windows, while panels **(C)** and **(D)** indicate that the focal 72kb region contains an excess of heterozygous sites in both the Chagyrskaya and Vindija Neanderthals.

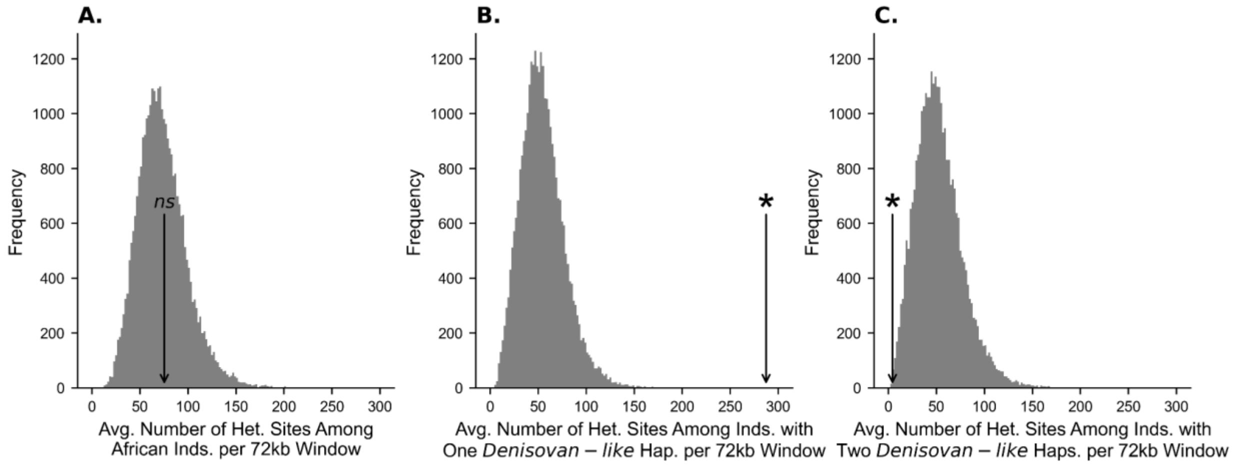


**Fig. S33. Individuals with exactly one *Denisovan-like* haplotype harbor an elevated number of heterozygous sites at the focal 72kb region.**

Distribution of the average number of heterozygous among **(A)** all African individuals, **(B)** individuals with exactly one *Denisovan-like* haplotype, and **(C)** individuals harboring two *Denisovan-like* haplotypes for non-overlapping 72kb windows. Arrows indicate the observed number of heterozygous sites at the focal 72kb region. Asterisks above arrows indicate statistical significance when compared to the genome-wide distribution of non-overlapping windows; "ns" denotes non-significance. This figure demonstrates that at the focal 72kb region **(A)** African individuals harbor an average number of heterozygous sites within the genome-wide expectation for 72kb windows. However, **(B)** individuals with one introgressed haplotype have an elevated average number of heterozygous sites, while **(C)** individuals harboring two introgressed haplotypes have a lower average number of heterozygous sites.

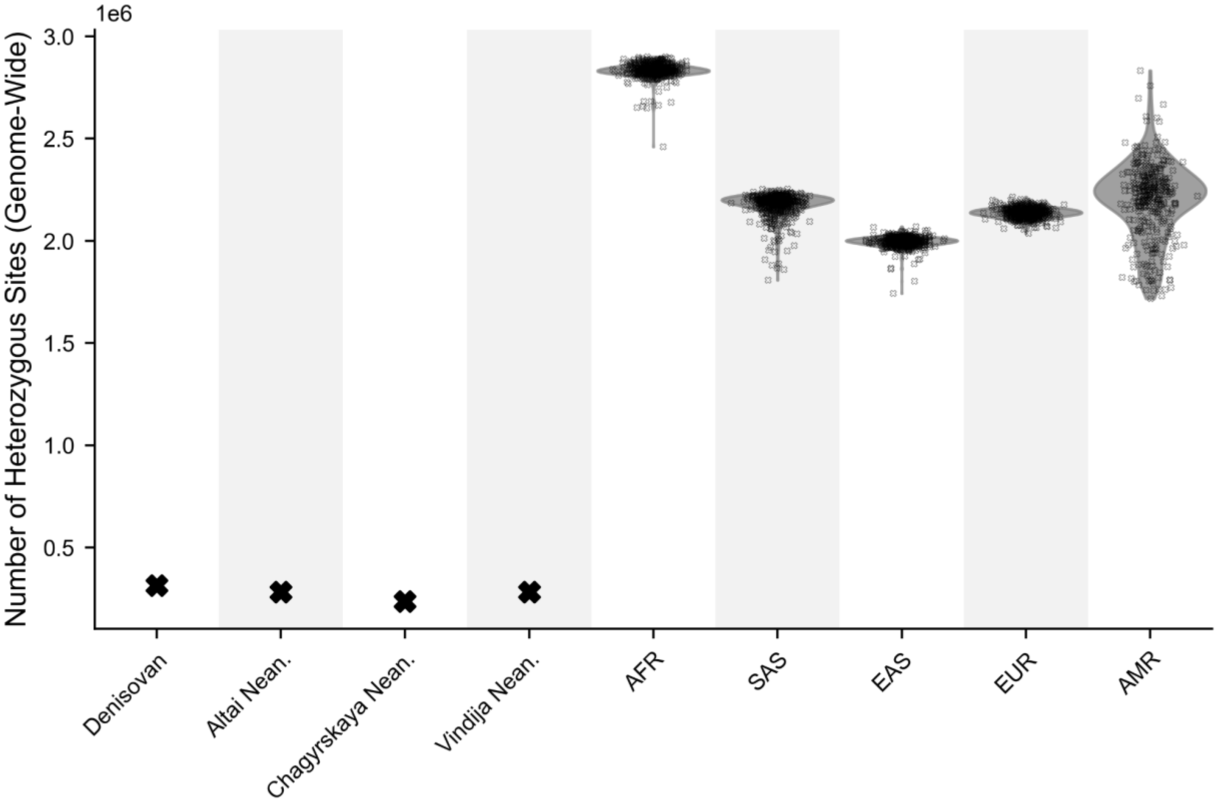


**Fig. S34. Genome-wide distribution of heterozygous sites among the archaics and 1000 Genomes Project (1KG) individuals.**

The average number of heterozygous sites per high-coverage archaic genome (filled black X) and 1KG individual (unfilled black X). The grey violin plots represent the distribution of heterozygous sites per individual stratified by super population: Africans (AFR), South Asians (SAS), East Asians (EAS), Europeans (EUR), and Admixed Americans (AMR).

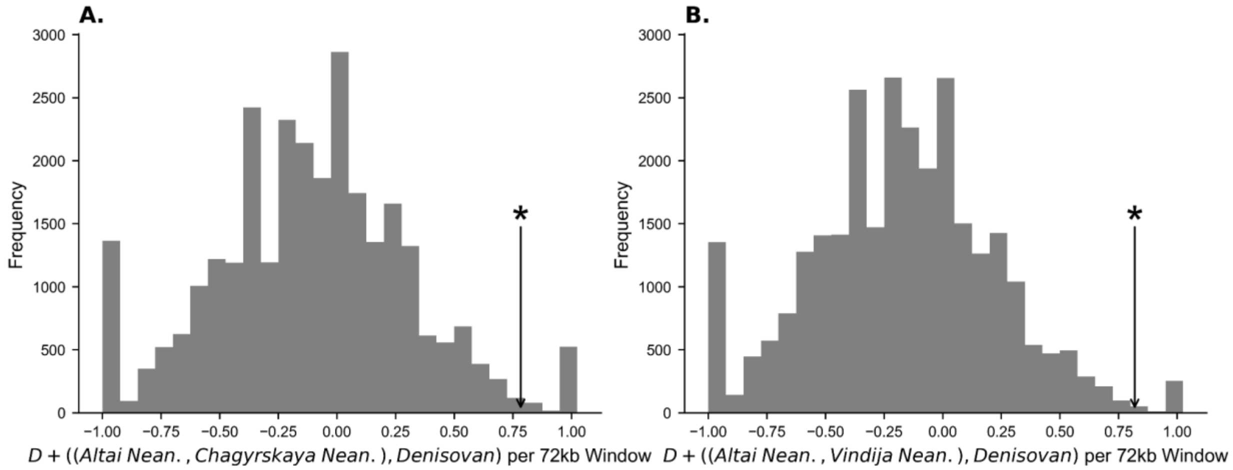


**Fig. S35. Excess of allele sharing between the late Neanderthals (Chagyrskaya and Vindija) and the sequenced Denisovan at the focal 72kb region.**

Distributions of $D+$ tests of introgression using the Altai Neanderthal as *P1*, **(A)** the Chagyrskaya Neanderthal as *P2,* **(B)** the Vindija Neanderthal as *P2*, the Denisovan as *P3*, and the *EPO* ancestral sequence as *P4* for non-overlapping 72kb windows. Arrows indicate the observed $D+$ value at the focal 72kb region. Asterisks above arrows indicate statistical significance when compared to the genome-wide distribution of non-overlapping windows; "ns" denotes non-significance. Panels **(A)** and **(B)** demonstrate that the Chagyrskaya and Vindija Neanderthals share more alleles than expected with the Denisovan.

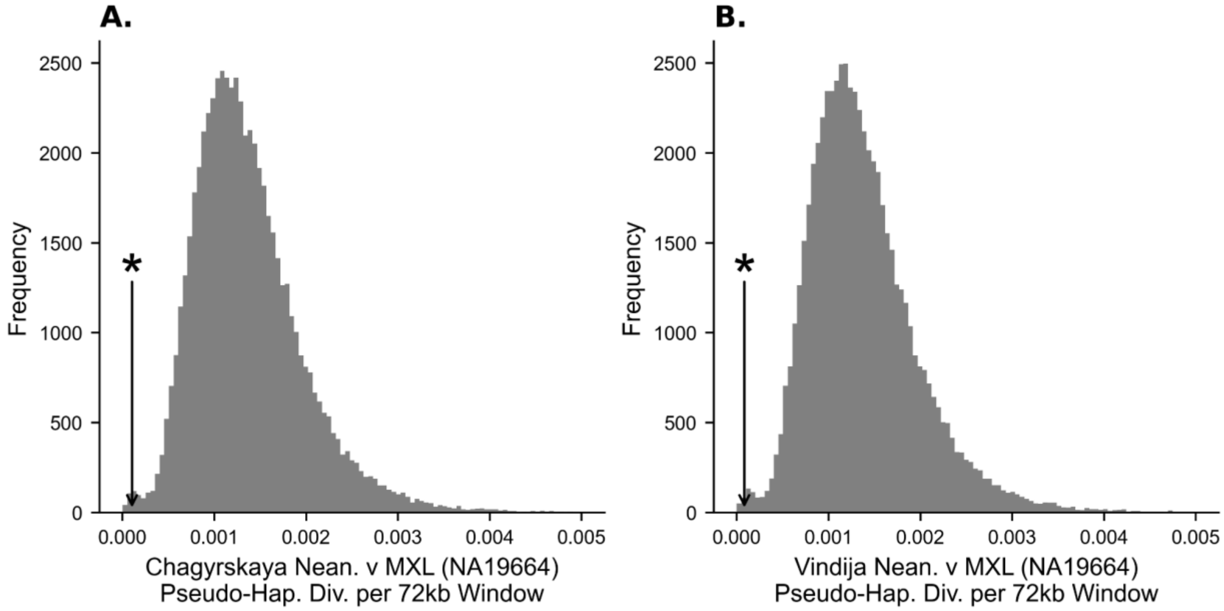


**Fig. S36. The *Denisovan-like* haplotype in MXL is closer than expected to the *Denisovan-like* haplotype in the Chagyrskaya and Vindija Neanderthals.**
Distributions of pseudo-haplotype divergence, i.e., the number of pairwise differences between the MXL individual's haplotypes (NA19664), who harbors two *Denisovan-like* haplotypes and has no heterozygous sites at the focal 72kb region, and pseudo-haplotypes from the two late Neanderthals normalized by the effective sequence length, for non-overlapping 72kb windows. Pseudo-haplotypes for **(A)** the Chagyrskaya Neanderthal and **(B)** the Vindija Neanderthal were generated by randomly sampling an allele for every genotype in a window. Arrows indicate the observed sequence divergence between the *Denisovan-like* haplotype in the MXL individual and the haplotype found in the late Neanderthal that is closest to the Denisovan after phasing the focal 72kb region. Asterisks above arrows indicate statistical significance when compared to the genome-wide distribution of non-overlapping windows; "ns" denotes non-significance. Panels **(A)** and **(B)** demonstrate that the introgressed haplotype in MXL is closer than expected to the haplotype in the Chagyrskaya and Vindija Neanderthals.

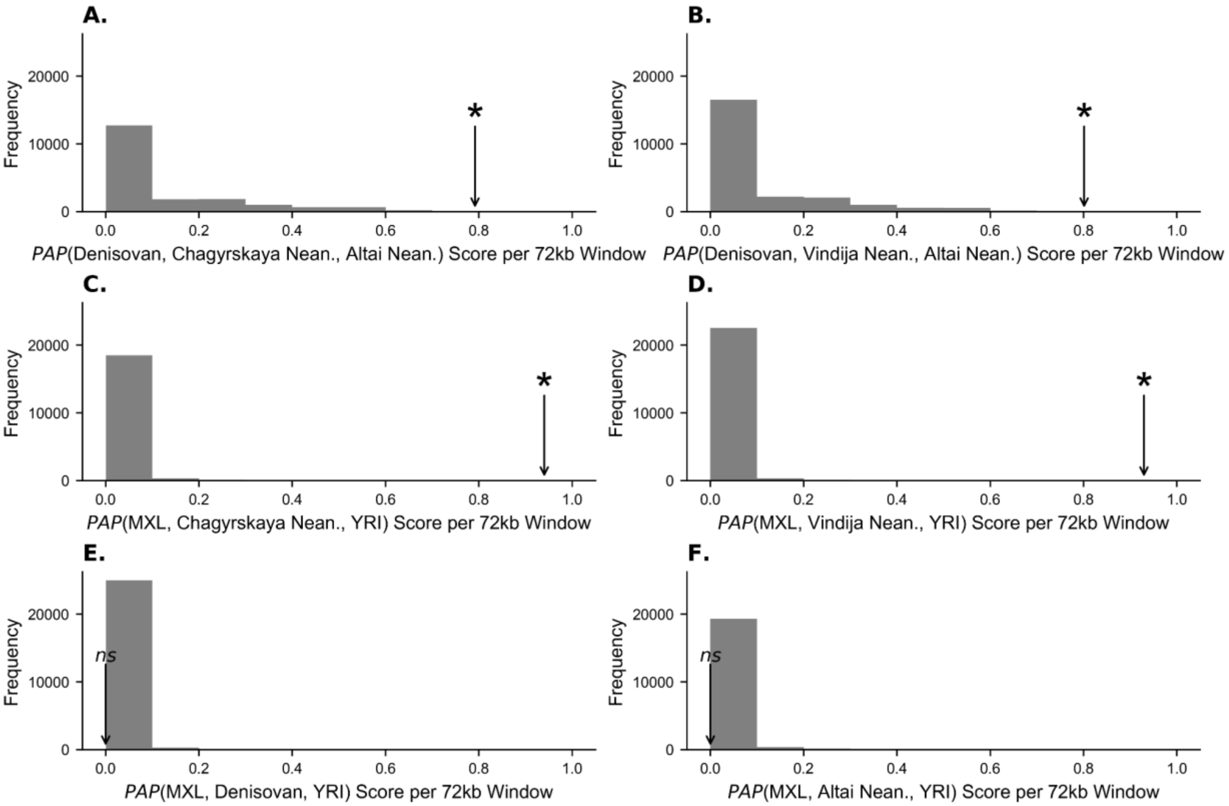


**Fig. S37. Late Neanderthal (Chagyrskaya and Vindija) heterozygous genotypes can be best explained as a mixture of a *Denisovan-like* haplotype and a non-introgressed haplotype at the focal 72kb region.**

Distribution of Pseudo-Ancestry Painting (*PAP*) scores for non-overlapping 72kb windows. *PAP*($Source^1$, *Target*, $Source^2$) scores quantify the number of heterozygous sites in the *Target* individual that can be explained by the two source individuals ($Source^1$ and $Source^2$) being fixed for different allelic states normalized by the total number of heterozygous sites in the *Target* individual. Arrows indicate the observed *PAP* score at the focal 72kb region. Asterisks above arrows indicate statistical significance when compared to the genome-wide distribution of non-overlapping windows; "ns" denotes non-significance. Panels **(A)** and **(B)** demonstrate that heterozygous sites in the Chagyrskaya and Vindija Neanderthals can be explained as a mixture of Denisovan and Altai Neanderthal sources. Panels **(C)** and **(D)** indicate that these heterozygous sites in the late Neanderthal are best explained as a mixture between a *Denisovan-like* haplotype in the MXL individual (NA19664 - who harbors two *Denisovan-like* haplotypes and has no heterozygous sites at the focal 72kb region) and the non-introgressed haplotype in the YRI individual (NA19190 - who does not harbor any introgressed haplotype and has no heterozygous sites at the focal 72kb region). Panels **(E)** and **(F)** demonstrate that *PAP* scores behave properly as neither the Denisovan nor Altai Neanderthal can be a mixture between a Denisovan-like haplotype in the MXL individual and the non-introgressed haplotype in the YRI individual.

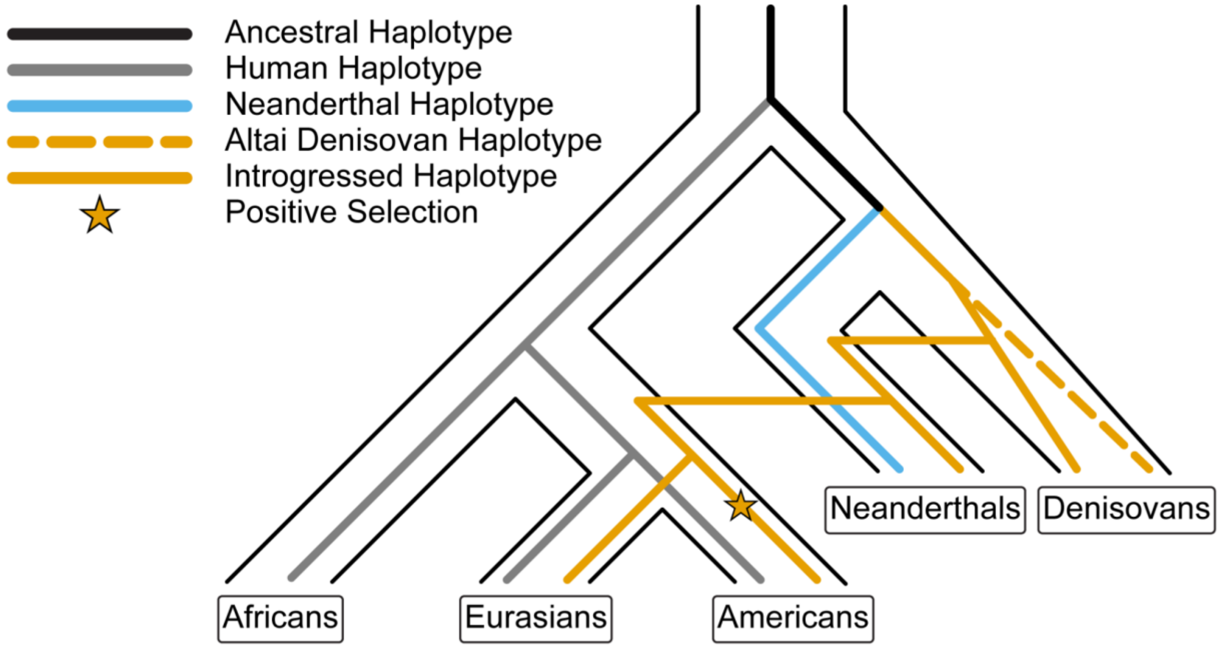


**Fig. S38. Evolutionary history of recurrent introgression and positive selection at *MUC19*.** The proposed evolutionary history of *MUC19* where the *Denisovan-like* haplotype was first introgressed from a Denisovan population that split from the sequenced Altai Denisovan into a Neanderthal population that was closely related to the two late Neanderthals, but not the Altai Neanderthal. This introgression event between archaic populations was then followed by an introgression event between the late Neanderthal population and an ancestral non-African population. Lastly, the introgressed haplotype then experienced positive selection in populations from the Americas.

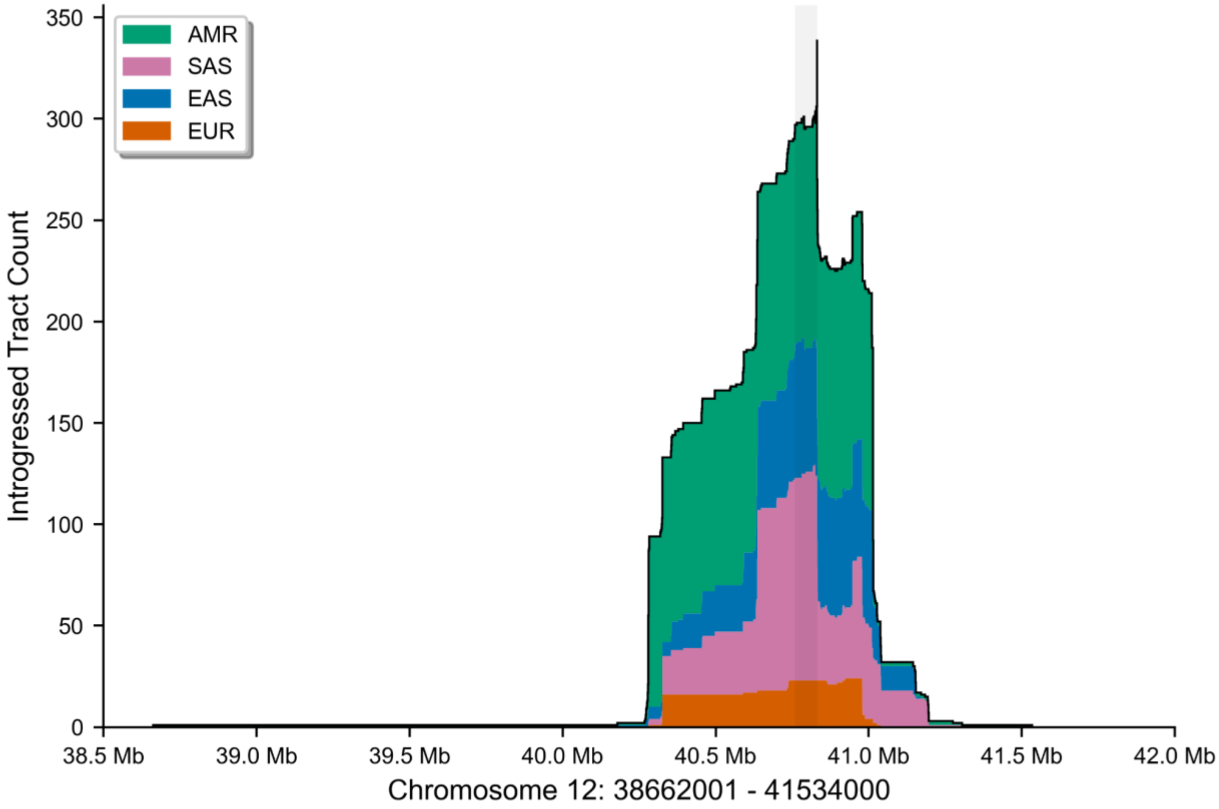


**Fig. S39. Density of introgressed tracts overlapping *MUC19* with the 2.613 Mb tract found in the PUR individual HG01108.**

Density of introgressed tracts that overlap *MUC19* for the 1KG (black outline) and stratified by super population—Admixed Americans (AMR) in bluish green, South Asians (SAS) in reddish purple, East Asians (EAS) in blue, and Europeans (EUR) in vermillion. The gray shaded region corresponds to the focal 72kb region, which is the densest contiguous region of introgressed tracts longer than 40kb. This figure differs from Figure 1A as it includes the 2.613 Mb tract found in the PUR individual HG01108, which was removed from Figure 1A for visual clarity.

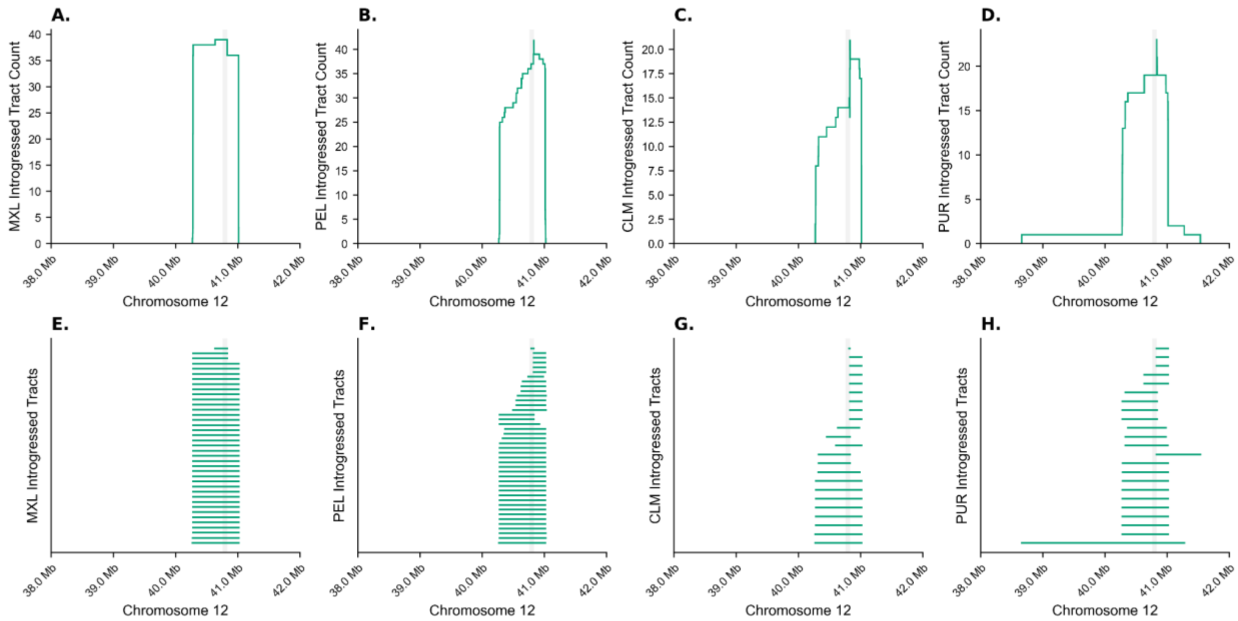


**Fig. S40. Introgressed tracts overlapping *MUC19* among AMR populations.**

Panels **(A)** through **(D)** show the density of introgressed tracts that overlap *MUC19*. Panels **(E)** through **(H)** show the introgressed tracts—sorted from longest to shortest—that overlap *MUC19* stratified by AMR populations. The gray shaded region corresponds to the focal 72kb region, which is the densest contiguous region of introgressed tracts longer than 40kb.

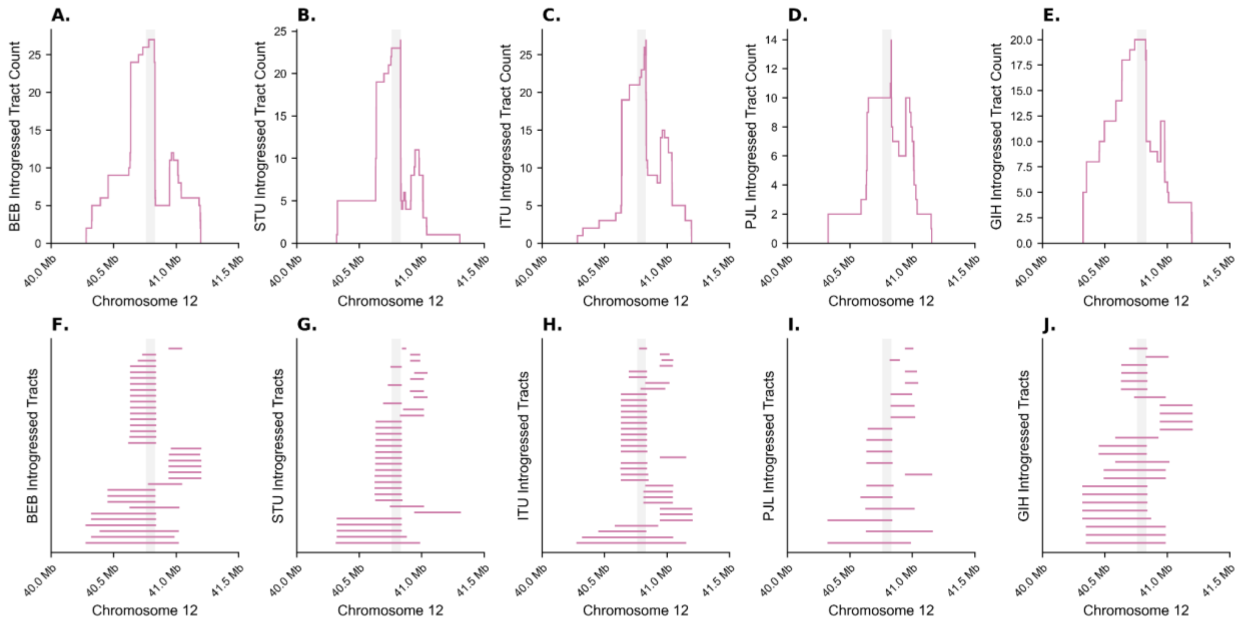


**Fig. S41. Introgressed tracts overlapping *MUC19* among SAS populations.**

Panels **(A)** through **(E)** show the density of introgressed tracts that overlap *MUC19*. Panels **(F)** through **(J)** show the introgressed tracts—sorted from longest to shortest—that overlap *MUC19* stratified by SAS populations. The gray shaded region corresponds to the focal 72kb region, which is the densest contiguous region of introgressed tracts longer than 40kb.

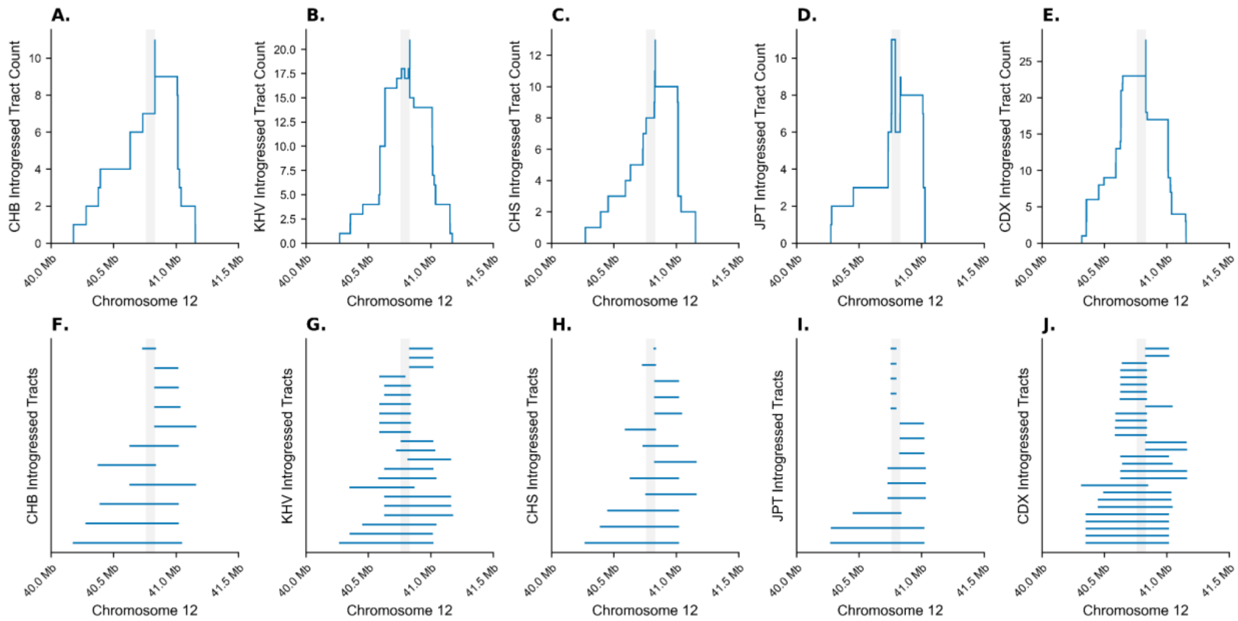


**Fig. S42. Introgressed tracts overlapping *MUC19* among EAS populations.**
Panels **(A)** through **(E)** show the density of introgressed tracts that overlap *MUC19*. Panels **(F)** through **(J)** show the introgressed tracts—sorted from longest to shortest—that overlap *MUC19* stratified by EAS populations. The gray shaded region corresponds to the focal 72kb region, which is the densest contiguous region of introgressed tracts longer than 40kb.

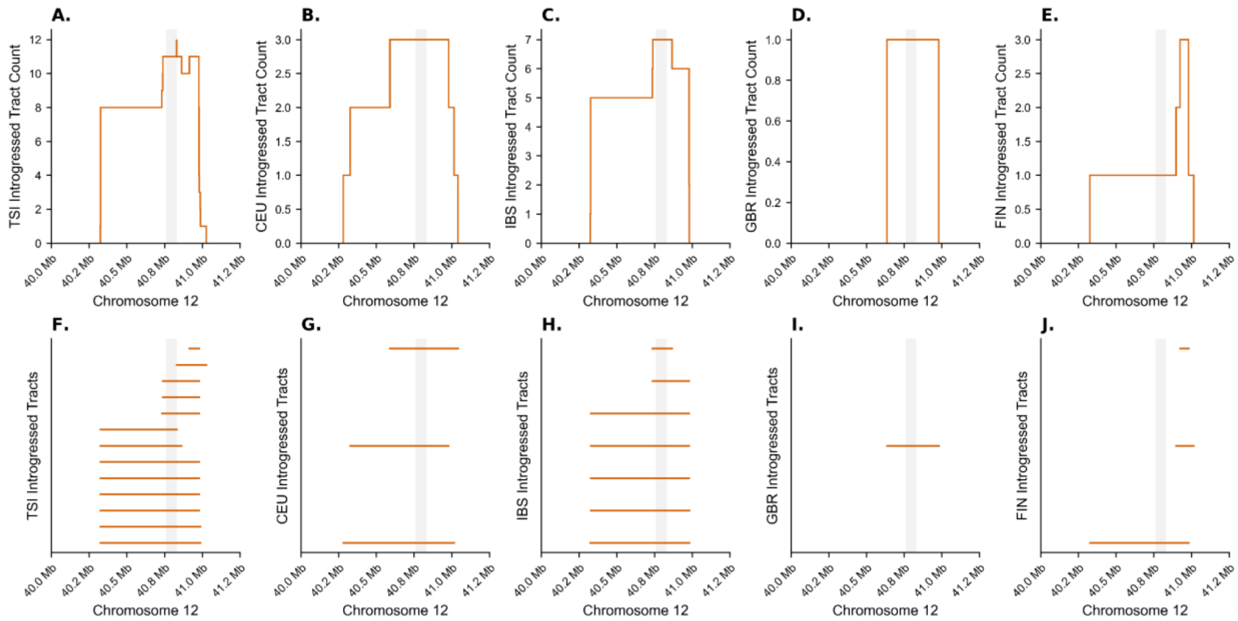


**Fig. S43. Introgressed tracts overlapping *MUC19* among EUR populations.**
Panels **(A)** through **(E)** show the density of introgressed tracts that overlap *MUC19*. Panels **(F)** through **(J)** show the introgressed tracts—sorted from longest to shortest—that overlap *MUC19* stratified by EUR populations. The gray shaded region corresponds to the focal 72kb region, which is the densest contiguous region of introgressed tracts longer than 40kb.

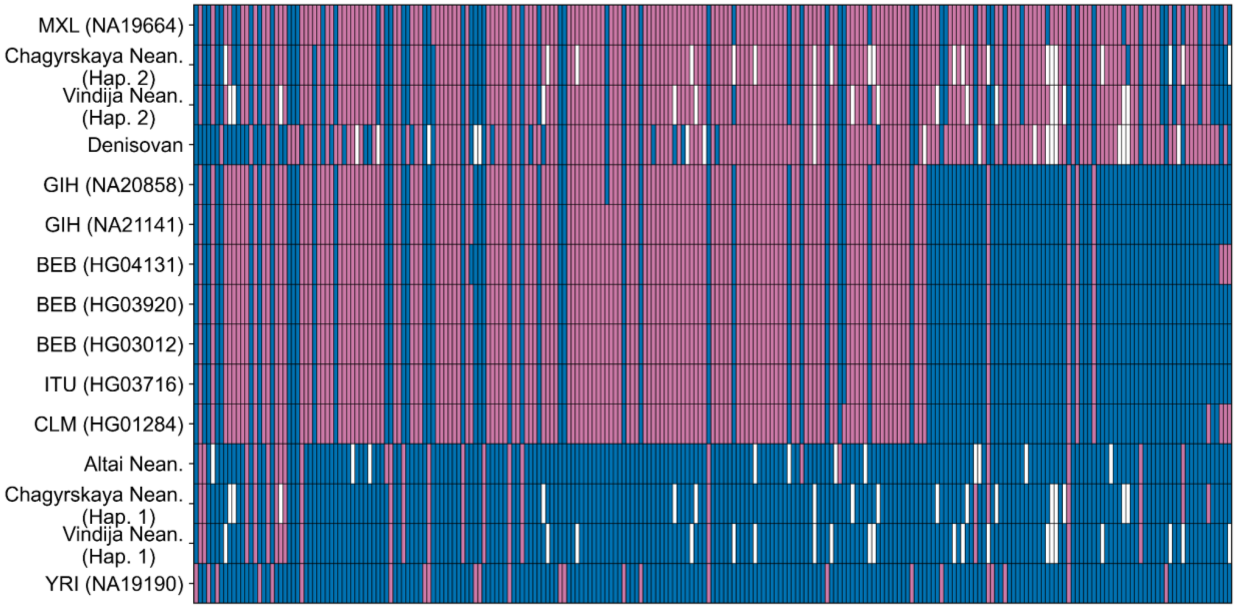


**Fig. S44. Seven recombinant introgressed haplotypes at the focal 72kb region.**
Haplotype matrix of the 245 segregating sites (columns) amongst the seven individuals who harbor a recombinant haplotype at the focal 72kb region; the focal MXL individual (NA19664) with two copies of the introgressed haplotype; the focal YRI individual (NA19190) without the introgressed haplotype; the Altai Denisovan; the Altai Neanderthal; and the two phased haplotypes for the Chagyrskaya and Vindija Neanderthals, respectively. Cells shaded blue denote the hg19 reference allele, cells shaded reddish purple denote the alternative allele, and cells shaded white represent sites that did not pass quality control in the given archaic individual. Note that the focal MXL and YRI individuals are homozygous for every position in the 72kb region in *MUC19* and that the heterozygous sites for the Altai Denisovan and Altai Neanderthal—six and one heterozygous sites, respectively—are omitted.

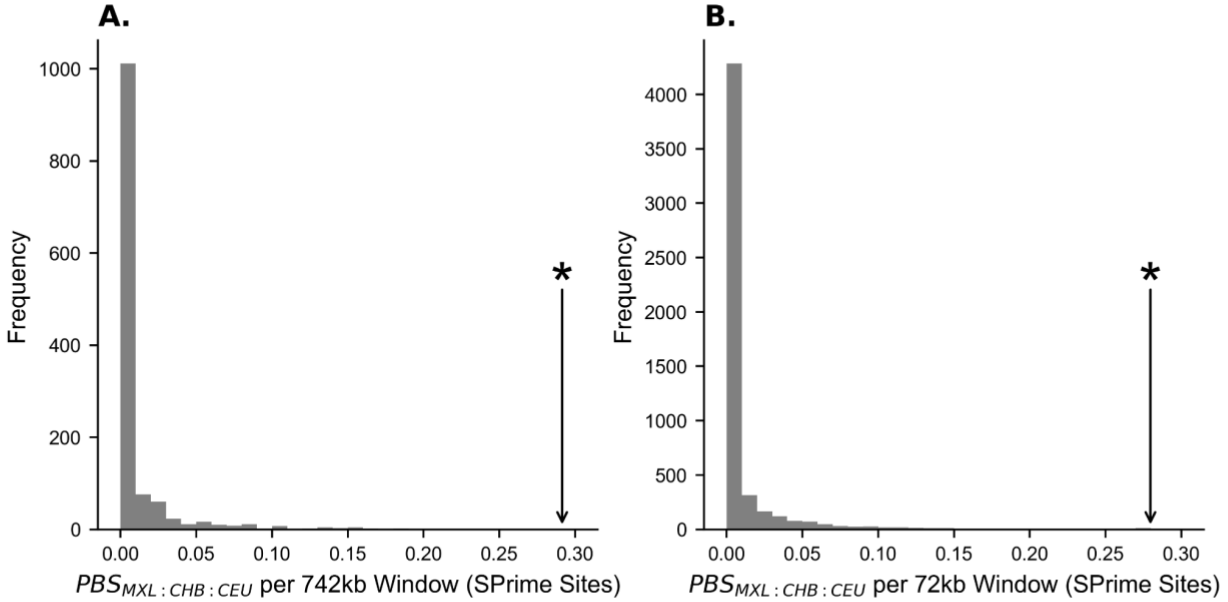


**Fig. S45. Both 742kb and 72kb focal regions exhibit signals of positive selection using the Population Branch Statistic (*PBS*) when considering only putatively introgressed sites.** Distribution of $PBS_{MXL:CHB:CEU}$ values for non-overlapping **(A)** 742kb and **(B)** 72kb windows for only the putatively introgressed sites in MXL denoted as a match with the Altai Denisovan or Altai Neanderthal as defined by the `SPrime` introgression maps. Arrows indicate the observed values for **(A)** the focal 742kb region corresponding to the longest introgressed tract found in MXL, and **(B)** the focal 72kb region, which is the densest contiguous region (>40kb) of introgressed tracts overlapping *MUC19*. Asterisks above arrows indicate statistical significance when compared to the genome-wide distribution of non-overlapping windows; "ns" denotes non-significance. Both panels demonstrate that **(A)** the 742kb region and **(B)** the 72kb region exhibit population genetic signals consistent with positive selection when only considering sites that are putatively introgressed in MXL as defined by SPrime.

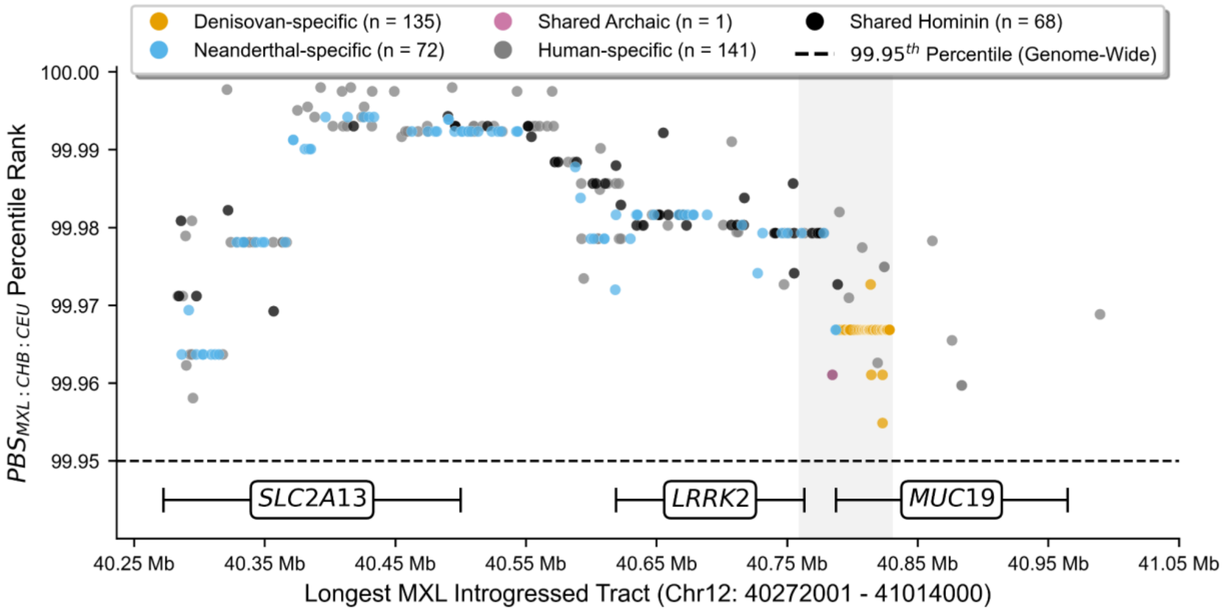


**Fig. S46. Percentile ranks for the 417 per-SNP $PBS_{MXL:CHB:CEU}$ outliers at the focal 742kb region.**

Percentile Rank for $PBS_{MXL:CHB:CEU}$ values greater than the 99.95th percentile of $PBS_{MXL:CHB:CEU}$ values for all SNPs genome-wide (black dashed line) in the focal 742kb region. The orange, sky blue, and reddish purple points represent SNPs that are rare or absent in Africa (<1%), present in MXL (>1%), and are, respectively, either shared uniquely with the Denisovan, uniquely with the Neanderthals, or shared with both the Denisovan and Neanderthals. The black points represent SNPs present across both modern human and archaic populations, while the gray points represent SNPs private to modern humans. The gray shaded region corresponds to the focal 72kb region.

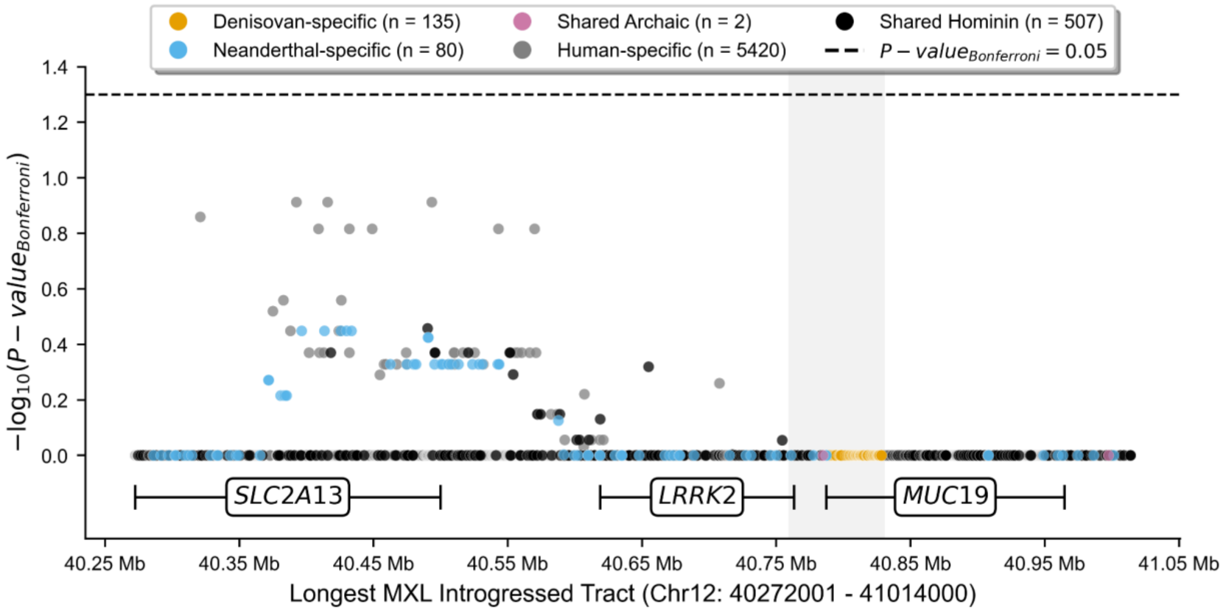


**Fig. S47. Bonferroni corrected *P-values* for the per-SNP $PBS_{MXL:CHB:CEU}$ values at the focal 742kb region.**

Bonferroni corrected *P-values* for all $PBS_{MXL:CHB:CEU}$ values across all SNPs in the 742kb region that corresponds to the longest introgressed tract found in MXL. The orange, sky blue, and reddish purple points represent SNPs that are rare or absent in Africa (<1%), present in MXL (>1%), and are, respectively, either shared uniquely with the Denisovan, uniquely with the Neanderthals, or shared with both the Denisovan and Neanderthals. The black points represent SNPs present across both modern human and archaic populations, while the gray points represent SNPs private to modern humans. The black dashed line represents the significance threshold of 0.05, and the gray shaded region corresponds to the focal 72kb region.

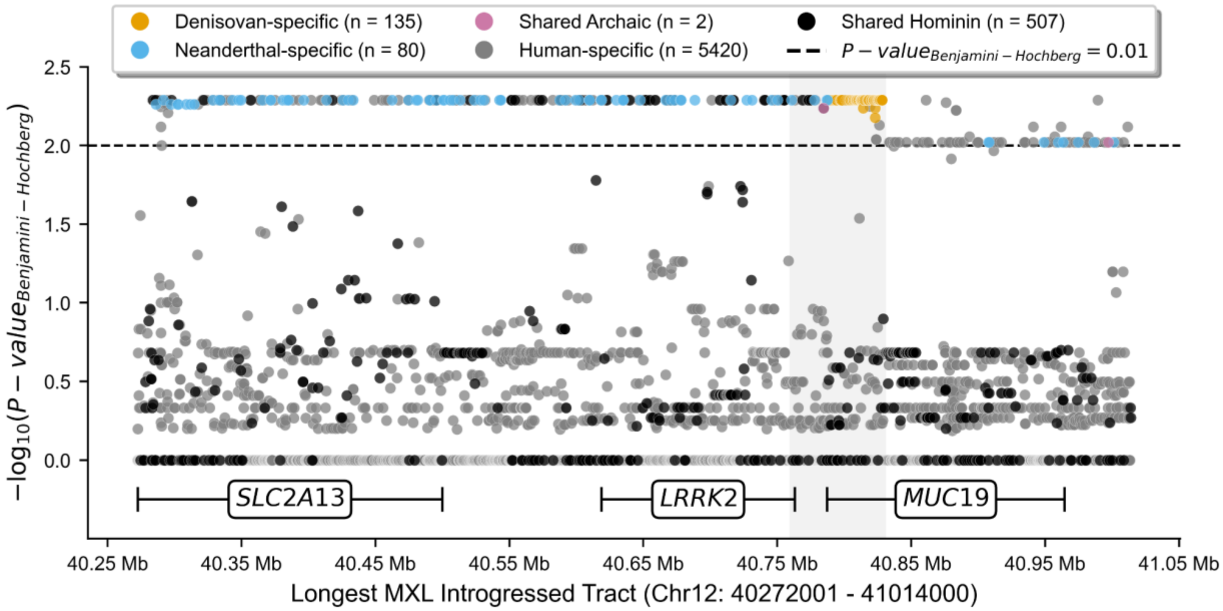


**Fig. S48. Benjamini–Hochberg corrected *P-values* for the per-SNP $PBS_{MXL:CHB:CEU}$ values at the focal 742kb region.**

Benjamini–Hochberg corrected *P-values* for per-SNP $PBS_{MXL:CHB:CEU}$ values across all SNPs in the 742kb region that corresponds to the longest introgressed tract found in MXL. The orange, sky blue, and reddish purple points represent SNPs that are rare or absent in Africa (<1%), present in MXL (>1%), and are, respectively, either shared uniquely with the Denisovan, uniquely with the Neanderthals, or shared with both the Denisovan and Neanderthals. The black points represent SNPs present across both modern human and archaic populations, while the gray points represent SNPs private to modern humans. The black dashed line represents the false discovery rate threshold of 0.01, and the gray shaded region corresponds to the focal 72kb region.

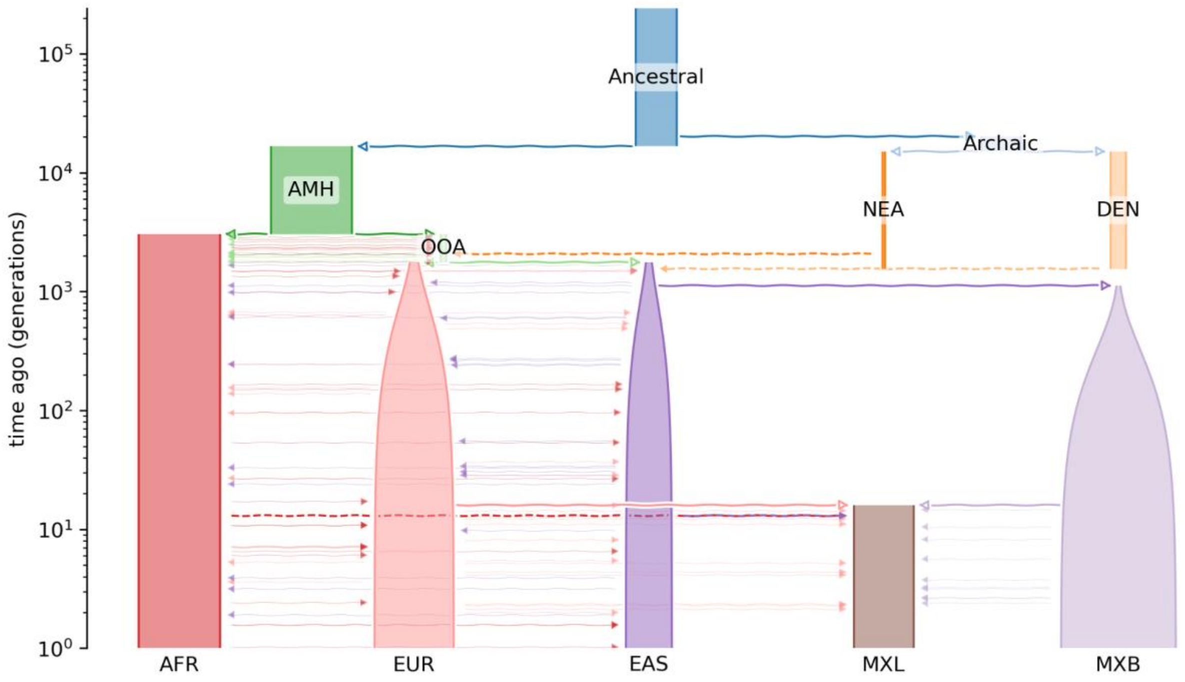


**Fig. S49. Demographic model used in *PBS* simulations.**
A graphical depiction of the extended Medina-Muñoz et al. 2023 demographic model for MXL in 1KG including archaic introgression. We used this model for all the forward-in-time simulations. Demographic parameters are described in Table S56.

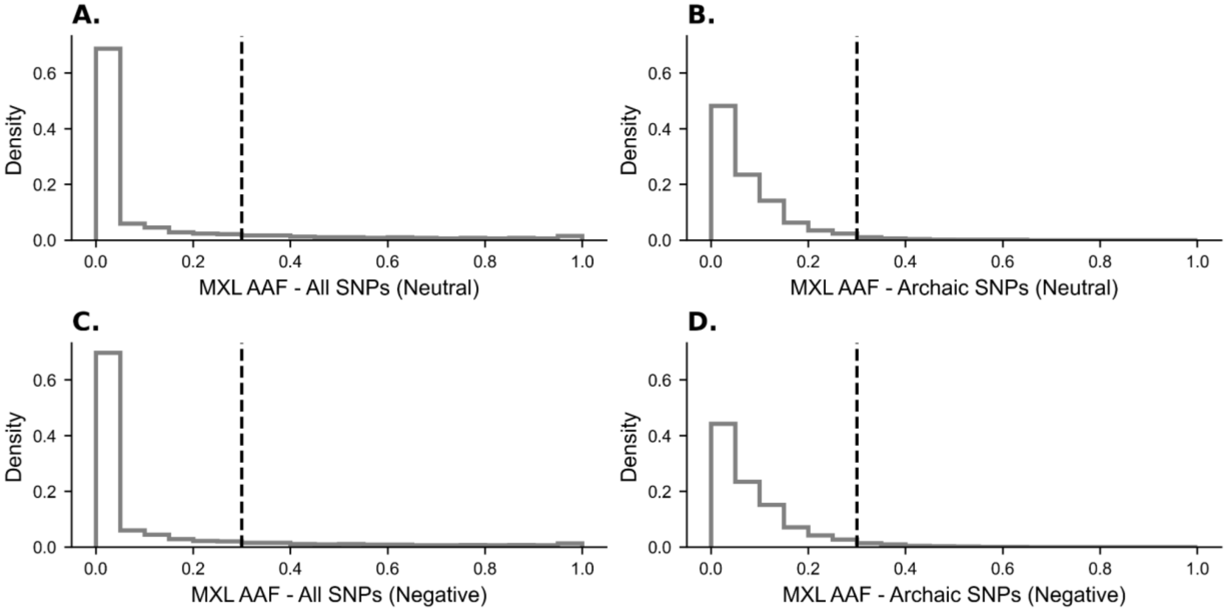


**Fig. S50. The probability of observing high frequency Denisovan-specific SNPs in MXL is low in all simulated scenarios.**

Distribution of simulated alternative allele frequencies (AAF) in a 742kb region for MXL across 10,000 replicates per simulation scenario. Rows represent the different simulation scenarios: **(A-B)** through neutral, **(C-D)** negative selection. Columns show AAF for the different SNP partitions: All SNPs or Archaic SNPs. Gray outlines indicate the probability of observing SNPs at each AAF bin (i.e., the number of SNPs in a given AAF bin divided by the total number of SNPs), with bin heights summing to one. Black dashed lines indicate an AAF of 0.3.

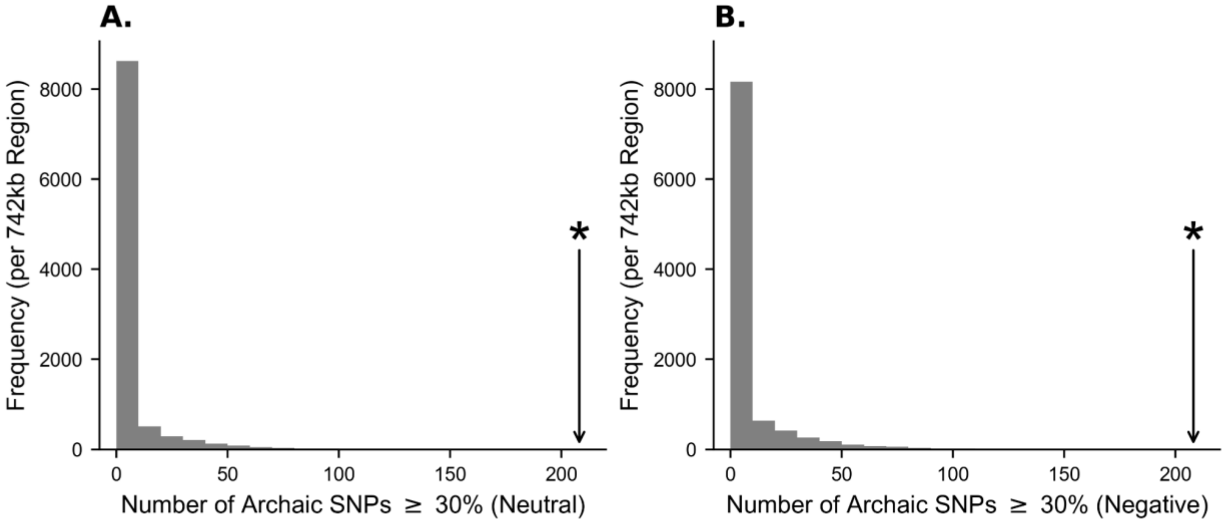


**Fig. S51. The observed number of archaic SNPs segregating at high frequency within the focal 742kb region in MXL is not expected under a neutral demographic process or a scenario of heterosis.**

Distribution of the number of SNPs in a 742kb region with an alternative allele frequency (AAF) greater than or equal to 0.3 in MXL per simulated replicate per simulation scenario—ie., **(A)** neutral and **(B)** negative selection. Arrows indicate the number of SNPs in MXL segregating at a frequency greater than or equal to 0.3 in the observed 742kb region. Asterisks above arrows indicate statistical significance for the observed value when compared to the distribution of 10,000 simulated replicates; "ns" denotes non-significance. Note that across all simulation scenarios, the "All SNPs" SNP partition is also significant (Table S60) but was omitted from the figure for visual clarity.

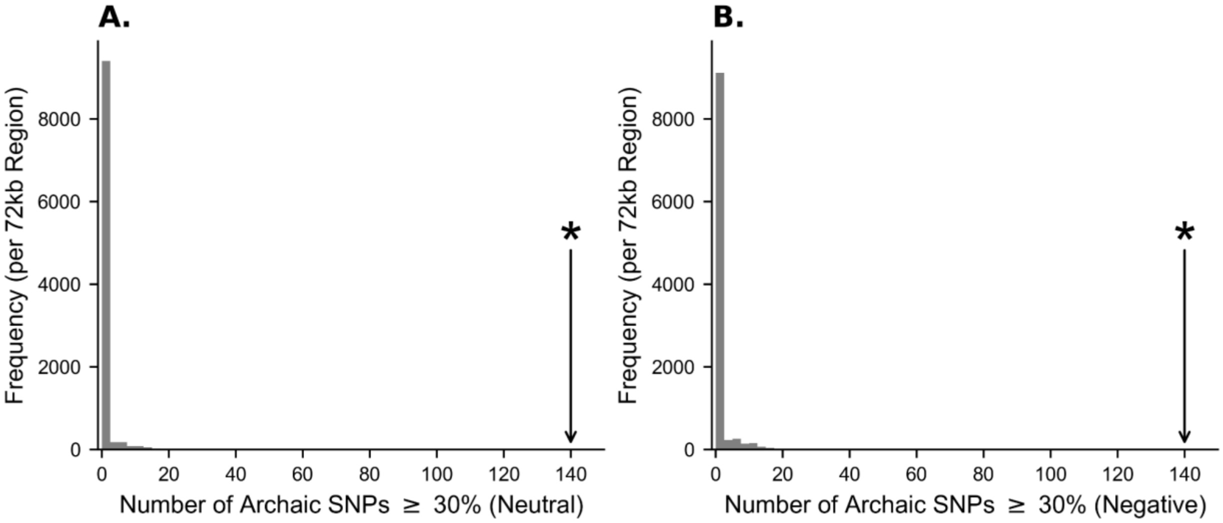


**Fig. S52. The observed number of archaic SNPs segregating at high frequency within the focal 72kb region in MXL is not expected under a neutral demographic process or a scenario of heterosis.**

Distribution of the number of SNPs in a 72kb region with an alternative allele frequency (AAF) greater than or equal to 0.3 in MXL per simulated replicate per simulation scenario—ie., **(A)** neutral and **(B)** negative selection. Arrows indicate the number of SNPs in MXL segregating at a frequency greater than or equal to 0.3 in the observed 72kb region. Asterisks above arrows indicate statistical significance for the observed value when compared to the distribution of 10,000 simulated replicates; "ns" denotes non-significance. Note that across all simulation scenarios, the "All SNPs" SNP partition is also significant (Table S60) but was omitted from the figure for visual clarity.

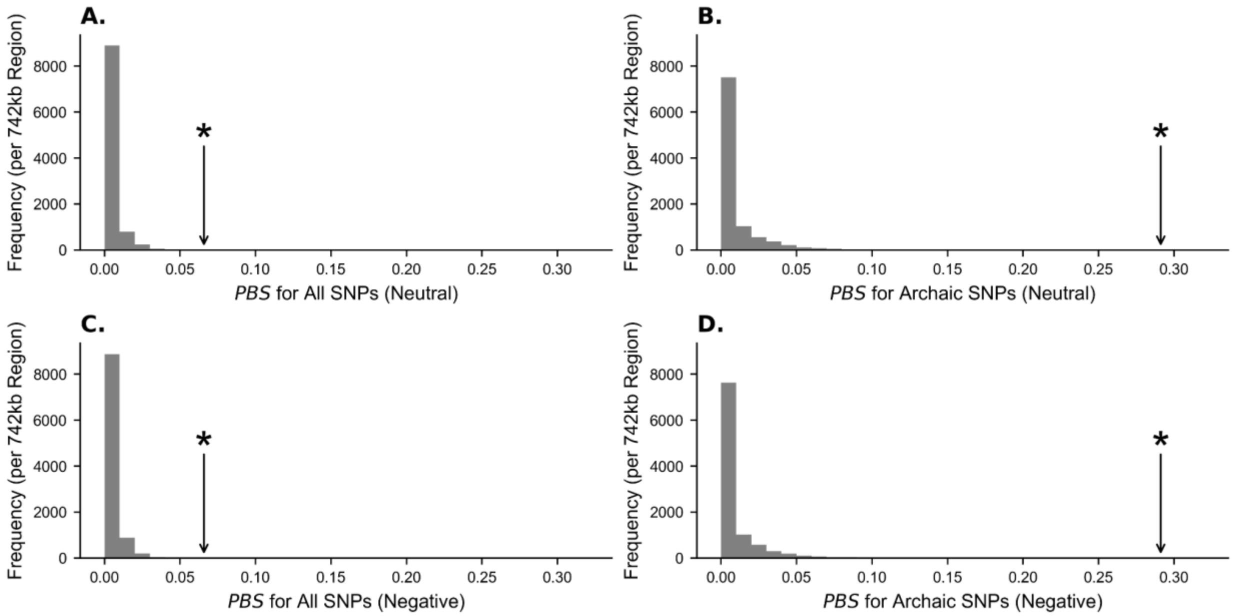


**Fig. S53. The observed number of per-region *PBS* values for the focal 742kb region is not expected under neutral demographic processes or a scenario of heterosis.**

Distribution of the per-region *PBS* values for each simulated replicate in a 742kb region per simulation scenario. Rows represent the different simulation scenarios: **(A-B)** through neutral, **(C-D)** negative selection. Columns show AAF for the different SNP partitions: All SNPs or Archaic SNPs. Arrows indicate the observed *PBS* value for the 742kb region in MXL. Asterisks above arrows indicate statistical significance for the observed value when compared to the distribution of 10,000 simulated replicates; "ns" denotes non-significance.

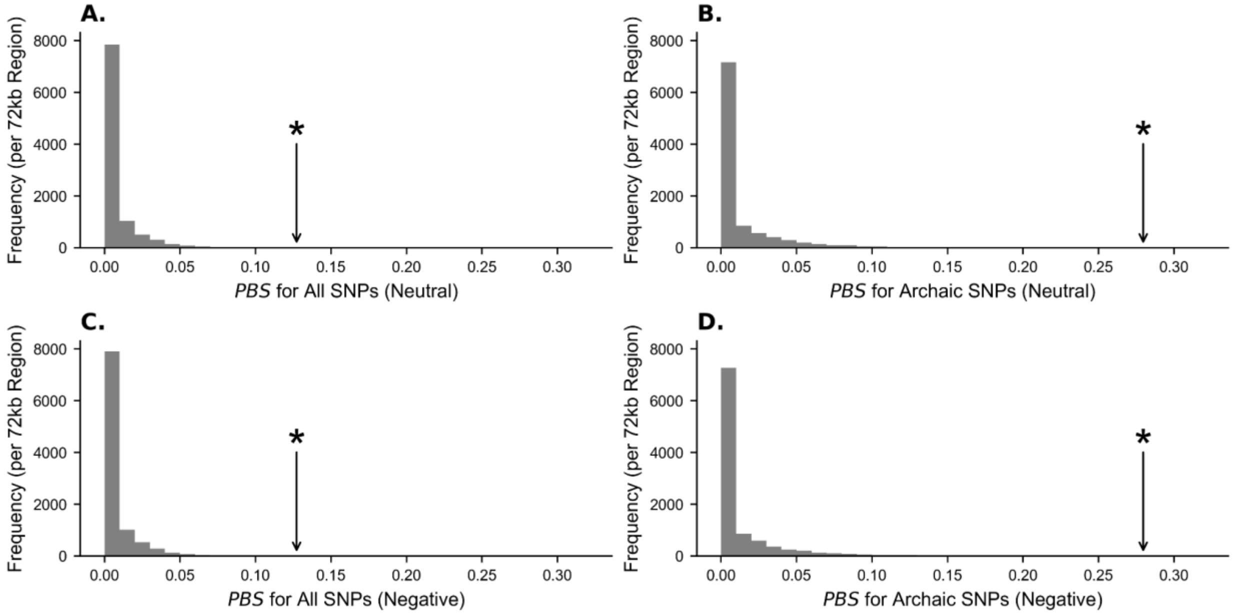


**Fig. S54. The observed number of per-region *PBS* values for the focal 72kb region is not expected under neutral demographic processes or a scenario of heterosis.**
Distribution of the per-region *PBS* values for each simulated replicate in a 72kb region per simulation scenario. Rows represent the different simulation scenarios: **(A-B)** through neutral, **(C-D)** negative selection. Columns show AAF for the different SNP partitions: All SNPs or Archaic SNPs. Arrows indicate the observed *PBS* value for the 72kb region in MXL. Asterisks above arrows indicate statistical significance for the observed value when compared to the distribution of 10,000 simulated replicates; "ns" denotes non-significance.

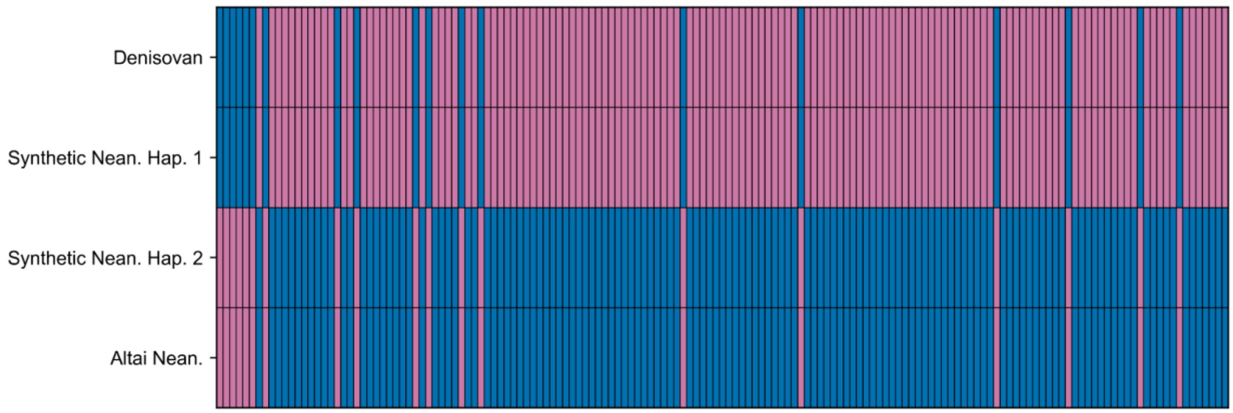


**Fig. S55. Perfect phasing of a Synthetic Neanderthal at the focal 72kb region.**
Haplotype matrix of the 155 heterozygous sites (columns) for the Denisovan, Synthetic Neanderthal, and Altai Neanderthal haplotypes (rows). Blue cells indicate the reference allele, while reddish purple cells represent the alternative allele. The Synthetic Neanderthal's "Hap. 1" aligns with the Denisovan haplotype and "Hap. 2" aligns with the Altai Neanderthal haplotype, which shows perfect phasing. Note that the Denisovan and Altai Neanderthal haplotypes are depicted as a single row since they are homozygous at every position.

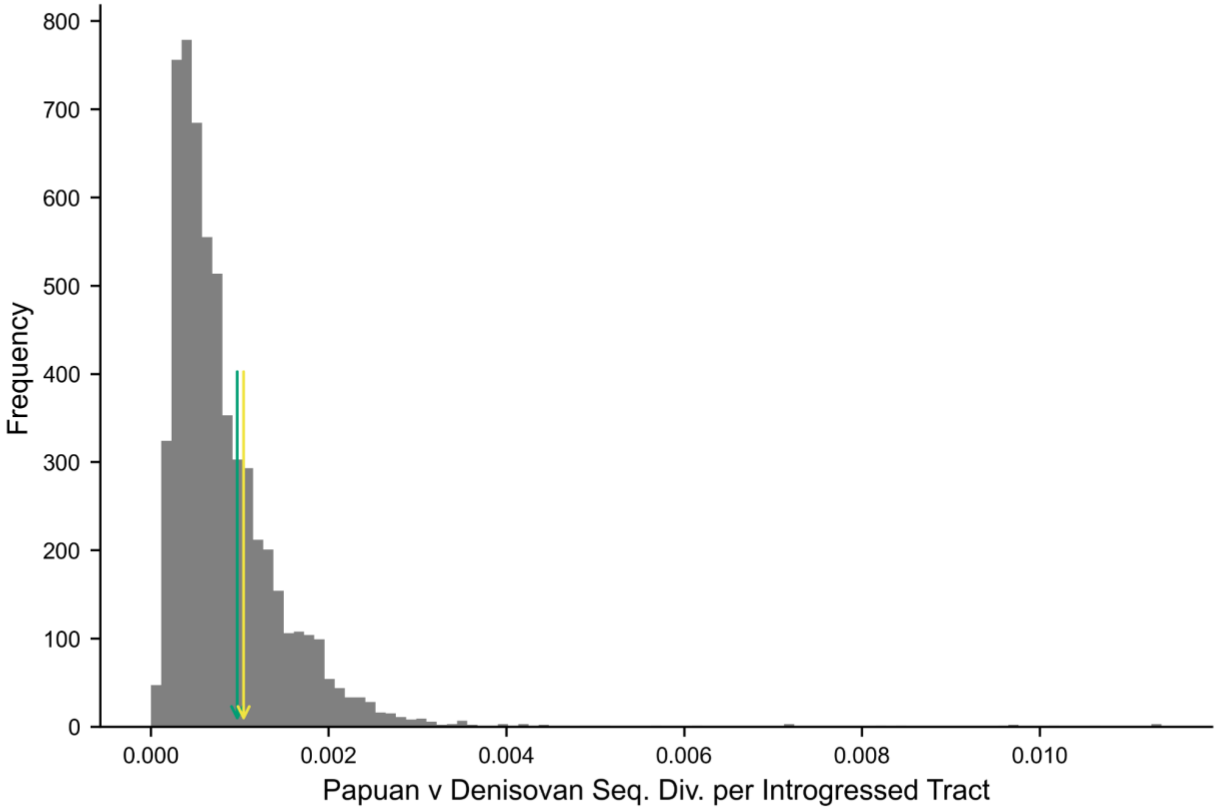


**Fig. S56. Sequence divergence between the introgressed haplotype and the Altai Denisovan fall within the expectation relative to other Denisovan introgressed segments.** Distribution of sequence divergence—number of pairwise differences between a Denisovan introgressed tract found in a Papuan individual and the Altai Denisovan's genotypes normalized by the effective sequence length—across all inferred Denisovan introgressed tracts in Papuans. The blue green and yellow arrows indicate the observed sequence divergence between the Altai Denisovan and the focal MXL (NA19664; Sequence Divergence: 0.00097; Percentile Rank: 70.607) and Papuan (B_Papuan-15; Sequence Divergence: 0.00104; Percentile Rank: 74.338) individuals, respectively.

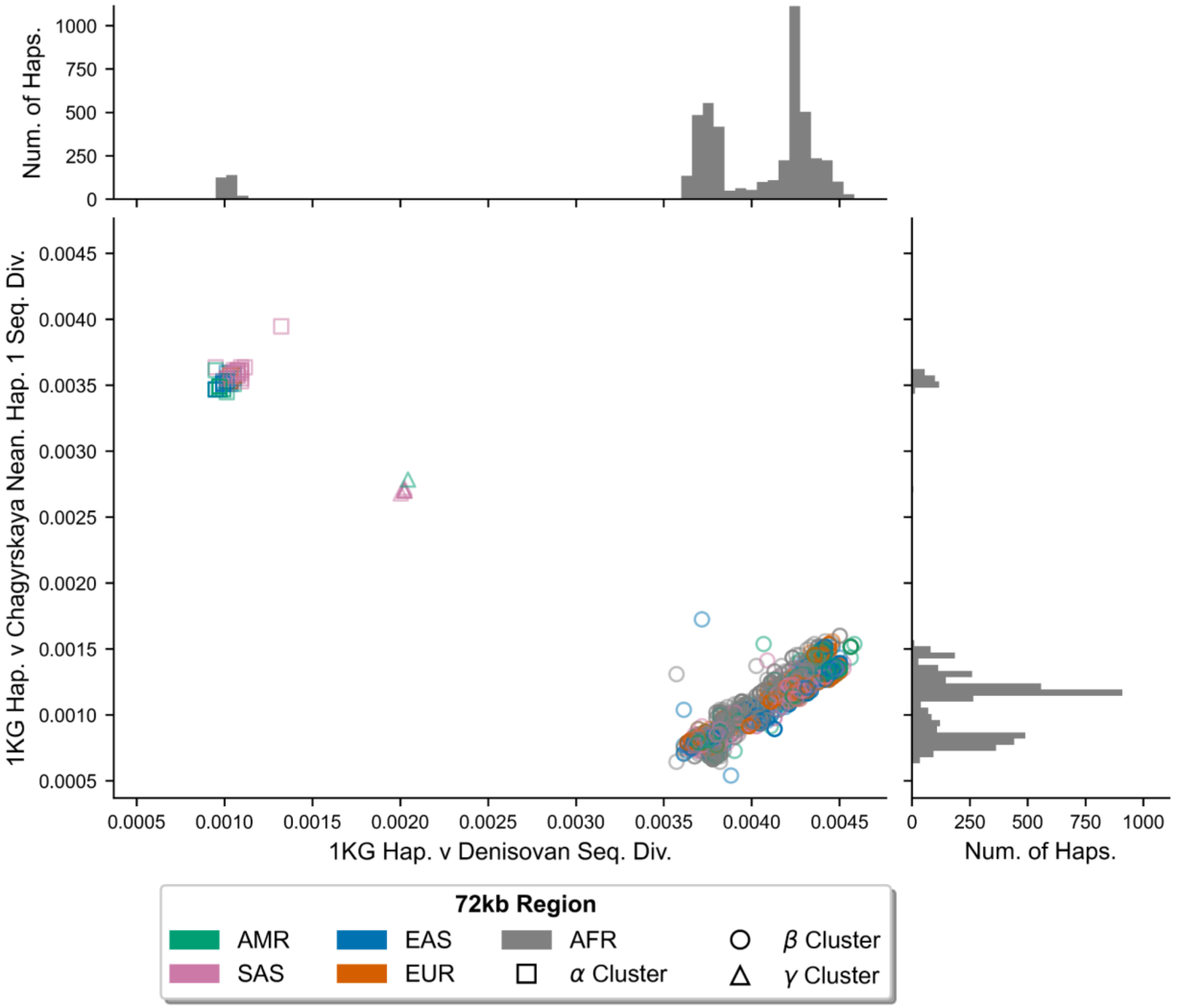


**Fig. S57. Joint distribution of sequence divergence from the Denisovan and the *Neanderthal-like* haplotype found in Chagyrskaya Neanderthal at the phased 72kb region.** The joint distribution of sequence divergence—number of pairwise differences between a modern human haplotype and an archaic genotype or phased late Neanderthal haplotype normalized by the effective sequence length—from the Denisovan (x-axis) and the *Neanderthal-like* haplotype found in Chagyrskaya Neanderthal (y-axis) for all haplotypes in the 1000 Genomes Project (1KG) at the focal 72kb region. Points correspond to 1KG haplotypes and are colored according to super population: Africans (AFR) in grey, admixed Americans (AMR) in bluish green, South Asians (SAS) in reddish purple, East Asians (EAS) in blue, and Europeans (EUR) in vermillion. The shape of each point indicates the cluster the haplotype falls within based on the ellipses in Figure 4: squares represent haplotypes within the alpha cluster, circles represent haplotypes within the beta cluster, and triangles represent haplotypes within the gamma cluster. Marginal distributions of sequence divergence from the Denisovan and the *Neanderthal-like* haplotype found in Chagyrskaya Neanderthal are depicted by the histograms above and to the right of the joint distribution, respectively.

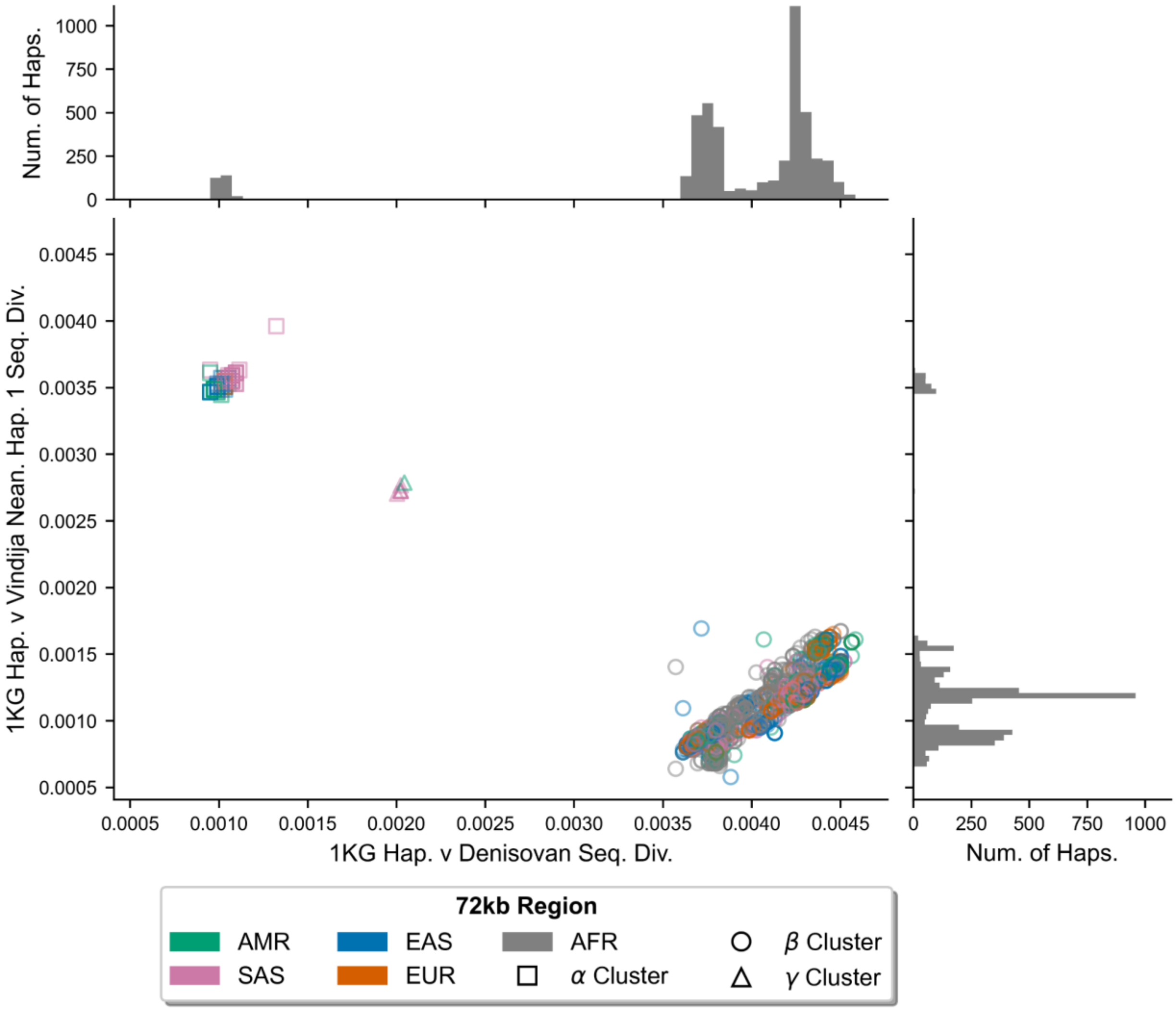


**Fig. S58. Joint distribution of sequence divergence from the Denisovan and the *Neanderthal-like* haplotype found in Vindija Neanderthal at the phased 72kb region.** The joint distribution of sequence divergence—number of pairwise differences between a modern human haplotype and an archaic genotype or phased late Neanderthal haplotype normalized by the effective sequence length—from the Denisovan (x-axis) and the *Neanderthal-like* haplotype found in Vindija Neanderthal (y-axis) for all haplotypes in the 1000 Genomes Project (1KG) at the focal 72kb region. Points correspond to 1KG haplotypes and are colored according to super population: Africans (AFR) in grey, admixed Americans (AMR) in bluish green, South Asians (SAS) in reddish purple, East Asians (EAS) in blue, and Europeans (EUR) in vermillion. The shape of each point indicates the cluster the haplotype falls within based on the ellipses in Figure 4: squares represent haplotypes within the alpha cluster, circles represent haplotypes within the beta cluster, and triangles represent haplotypes within the gamma cluster. Marginal distributions of sequence divergence from the Denisovan and the *Neanderthal-like* haplotype found in Vindija Neanderthal are depicted by the histograms above and to the right of the joint distribution, respectively.

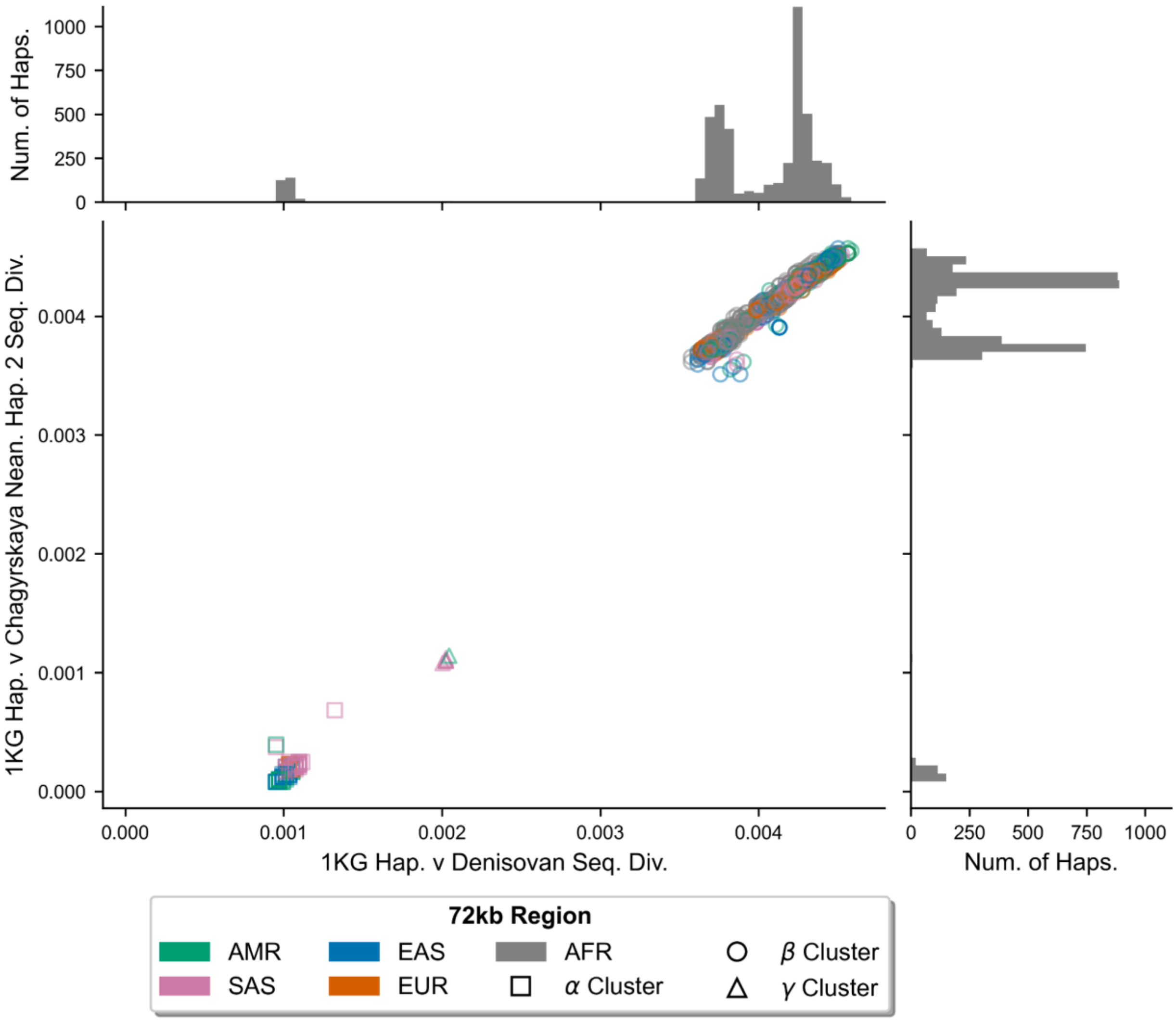


**Fig. S59. Joint distribution of sequence divergence from the Denisovan and the *Denisovan-like* haplotype found in Chagyrskaya Neanderthal at the phased 72kb region.**
The joint distribution of sequence divergence—number of pairwise differences between a modern human haplotype and an archaic genotype or phased late Neanderthal haplotype normalized by the effective sequence length—from the Denisovan (x-axis) and the *Denisovan-like* haplotype found in Chagyrskaya Neanderthal (y-axis) for all haplotypes in the 1000 Genomes Project (1KG) at the focal 72kb region. Points correspond to 1KG haplotypes and are colored according to super population: Africans (AFR) in grey, admixed Americans (AMR) in bluish green, South Asians (SAS) in reddish purple, East Asians (EAS) in blue, and Europeans (EUR) in vermillion. The shape of each point indicates the cluster the haplotype falls within based on the ellipses in Figure 4: squares represent haplotypes within the alpha cluster, circles represent haplotypes within the beta cluster, and triangles represent haplotypes within the gamma cluster. Marginal distributions of sequence divergence from the Denisovan and the *Denisovan-like* haplotype found in Chagyrskaya Neanderthal are depicted by the histograms above and to the right of the joint distribution, respectively.

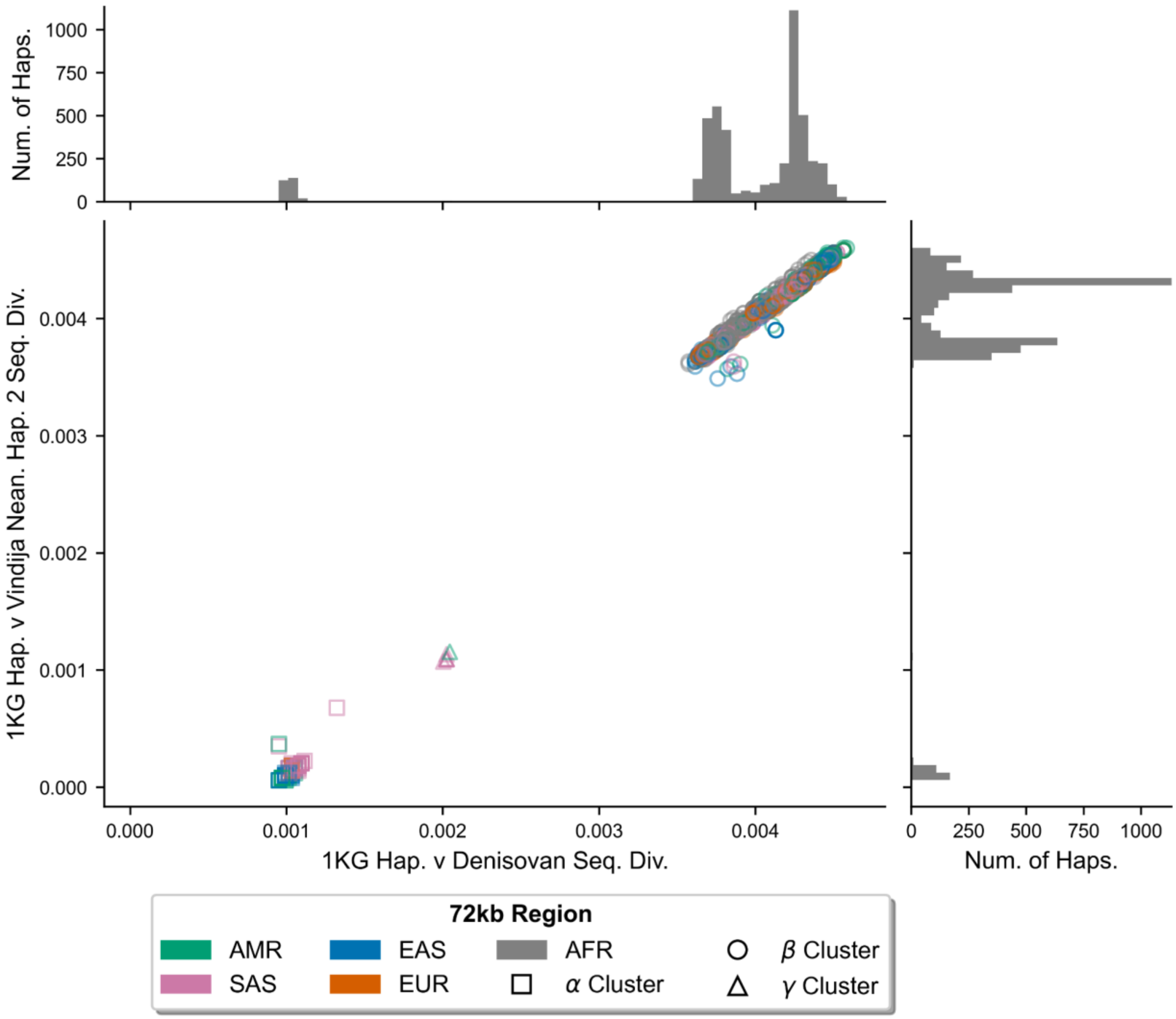


**Fig. S60. Joint distribution of sequence divergence from the Denisovan and the *Denisovan-like* haplotype found in Vindija Neanderthal at the phased 72kb region.**
The joint distribution of sequence divergence—number of pairwise differences between a modern human haplotype and an archaic genotype or phased late Neanderthal haplotype normalized by the effective sequence length—from the Denisovan (x-axis) and the *Denisovan-like* haplotype found in Vindija Neanderthal (y-axis) for all haplotypes in the 1000 Genomes Project (1KG) at the focal 72kb region. Points correspond to 1KG haplotypes and are colored according to super population: Africans (AFR) in grey, admixed Americans (AMR) in bluish green, South Asians (SAS) in reddish purple, East Asians (EAS) in blue, and Europeans (EUR) in vermillion. The shape of each point indicates the cluster the haplotype falls within based on the ellipses in Figure 4: squares represent haplotypes within the alpha cluster, circles represent haplotypes within the beta cluster, and triangles represent haplotypes within the gamma cluster. Marginal distributions of sequence divergence from the Denisovan and the *Denisovan-like* haplotype found in Vindija Neanderthal are depicted by the histograms above and to the right of the joint distribution, respectively.

**Supplemental Material References:**


[54] 1000 Genomes Project Consortium et al. A global reference for human genetic variation. Nature, 526(7571):68, 2015.

[55] Swapan Mallick, Heng Li, Mark Lipson, Iain Mathieson, Melissa Gymrek, Fernando Racimo, Mengyao Zhao, Niru Chennagiri, Susanne Nordenfelt, Arti Tandon, Pontus Skoglund, Iosif Lazaridis, Sriram Sankararaman, Qiaomei Fu, Nadin Rohland, Gabriel Renaud, Yaniv Erlich, Thomas Willems, Carla Gallo, Jeffrey P. Spence, Yun S. Song, Giovanni Poletti, Francois Balloux, George van Driem, Peter de Knijff, Irene Gallego Romero, Aashish R. Jha, Doron M. Behar, Claudio M. Bravi, Cristian Capelli, Tor Hervig, Andres Moreno-Estrada, Olga L. Posukh, Elena Balanovska, Oleg Balanovsky, Sena Karachanak-Yankova, Hovhannes Sahakyan, Draga Toncheva, Levon Yepiskoposyan, Chris Tyler-Smith, Yali Xue, M. Syafiq Abdullah, Andres Ruiz-Linares, Cynthia M. Beall, Anna Di Rienzo, Choongwon Jeong, Elena B. Starikovskaya, Ene Metspalu, Jüri Parik, Richard Villems, Brenna M. Henn, Ugur Hodoglugil, Robert Mahley, Antti Sajantila, George Stamatoyannopoulos, Joseph T. S. Wee, Rita Khusainova, Elza Khusnutdinova, Sergey Litvinov, George Ayodo, David Comas, Michael F. Hammer, Toomas Kivisild, William Klitz, Cheryl A. Winkler, Damian Labuda, Michael Bamshad, Lynn B. Jorde, Sarah A. Tishkoff, W. Scott Watkins, Mait Metspalu, Stanislav Dryomov, Rem Sukernik, Lalji Singh, Kumarasamy Thangaraj, Svante Pääbo, Janet Kelso, Nick Patterson & David Reich The simons genome diversity project: 300 genomes from 142 diverse populations. Nature, 538(7624):201–206, 2016.

[56] Karen H. Y. Wong, Walfred Ma, Chun-Yu Wei, Erh-Chan Yeh, Wan-Jia Lin, Elin H. F. Wang, Jen-Ping Su, Feng-Jen Hsieh, Hsiao-Jung Kao, Hsiao-Huei Chen, Stephen K. Chow, Eleanor Young, Catherine Chu, Annie Poon, Chi-Fan Yang, Dar-Shong Lin, Yu-Feng Hu, Jer-Yuarn Wu, Ni-Chung Lee, Wuh-Liang Hwu, Dario Boffelli, David Martin, Ming Xiao & Pui-Yan Kwok. Towards a reference genome that captures global genetic diversity. Nature communications, 11(1):5482, 2020.

[57] Benedict Paten, Javier Herrero, Kathryn Beal, Stephen Fitzgerald, and Ewan Birney. Enredo and pecan: genome-wide mammalian consistency based multiple alignment with paralogs. Genome research, 18(11):1814– 1828, 2008.

[58] Benedict Paten, Javier Herrero, Stephen Fitzgerald, Kathryn Beal, Paul Flicek, Ian Holmes, and Ewan Birney. Genome-wide nucleotide-level mammalian ancestor reconstruction. Genome research, 18(11):1829–1843, 2008.

[59] Wen-Wei Liao, Mobin Asri, Jana Ebler, Daniel Doerr, Marina Haukness, Glenn Hickey, Shuangjia Lu, Julian K. Lucas, Jean Monlong, Haley J. Abel, Silvia Buonaiuto, Xian H. Chang, Haoyu Cheng, Justin Chu, Vincenza Colonna, Jordan M. Eizenga,

Xiaowen Feng, Christian Fischer, Robert S. Fulton, Shilpa Garg, Cristian Groza, Andrea Guarracino, William T. Harvey, Simon Heumos, Kerstin Howe, Miten Jain, Tsung-Yu Lu, Charles Markello, Fergal J. Martin, Matthew W. Mitchell, Katherine M. Munson, Moses Njagi Mwaniki, Adam M. Novak, Hugh E. Olsen, Trevor Pesout, David Porubsky, Pjotr Prins, Jonas A. Sibbesen, Jouni Sirén, Chad Tomlinson, Flavia Villani, Mitchell R. Vollger, Lucinda L. Antonacci-Fulton, Gunjan Baid, Carl A. Baker, Anastasiya Belyaeva, Konstantinos Billis, Andrew Carroll, Pi-Chuan Chang, Sarah Cody, Daniel E. Cook, Robert M. Cook-Deegan, Omar E. Cornejo, Mark Diekhans, Peter Ebert, Susan Fairley, Olivier Fedrigo, Adam L. Felsenfeld, Giulio Formenti, Adam Frankish, Yan Gao, Nanibaa' A. Garrison, Carlos Garcia Giron, Richard E. Green, Leanne Haggerty, Kendra Hoekzema, Thibaut Hourlier, Hanlee P. Ji, Eimear E. Kenny, Barbara A. Koenig, Alexey Kolesnikov, Jan O. Korbel, Jennifer Kordosky, Sergey Koren, HoJoon Lee, Alexandra P. Lewis, Hugo Magalhães, Santiago Marco-Sola, Pierre Marijon, Ann McCartney, Jennifer McDaniel, Jacquelyn Mountcastle, Maria Nattestad, Sergey Nurk, Nathan D. Olson, Alice B. Popejoy, Daniela Puiu, Mikko Rautiainen, Allison A. Regier, Arang Rhie, Samuel Sacco, Ashley D. Sanders, Valerie A. Schneider, Baergen I. Schultz, Kishwar Shafin, Michael W. Smith, Heidi J. Sofia, Ahmad N. Abou Tayoun, Françoise Thibaud-Nissen, Francesca Floriana Tricomi, Justin Wagner, Brian Walenz, Jonathan M. D. Wood, Aleksey V. Zimin, Guillaume Bourque, Mark J. P. Chaisson, Paul Flicek, Adam M. Phillippy, Justin M. Zook, Evan E. Eichler, David Haussler, Ting Wang, Erich D. Jarvis, Karen H. Miga, Erik Garrison, Tobias Marschall, Ira M. Hall, Heng Li & Benedict Paten. "A draft human pangenome reference." Nature 617, no. 7960: 312-324, 2023.

[60] Jonas A. Gustafson, Sophia B. Gibson, Nikhita Damaraju, Miranda P.G. Zalusky, Kendra Hoekzema, David Twesigomwe, Lei Yang, Anthony A. Snead, Phillip A. Richmond, Wouter De Coster, Nathan D. Olson, Andrea Guarracino, Qiuhui Li, Angela L. Miller, Joy Goffena, Zachary B. Anderson, Sophie H.R. Storz, Sydney A. Ward, Maisha Sinha, Claudia Gonzaga-Jauregui, Wayne E. Clarke, Anna O. Basile, André Corvelo, Catherine Reeves, Adrienne Helland, Rajeeva Lochan Musunuri, Mahler Revsine, Karynne E. Patterson, Cate R. Paschal, Christina Zakarian, Sara Goodwin, Tanner D. Jensen, Esther Robb, The Genomes ONT Sequencing Consortium, University of Washington Center for Rare Disease Research (UW-CRDR), Genomics Research to Elucidate the Genetics of Rare Diseases (GREGoR) Consortium, William Richard McCombie, Fritz J. Sedlazeck, Justin M. Zook, Stephen B. Montgomery, Erik Garrison, Mikhail Kolmogorov, Michael C. Schatz, Richard N. McLaughlin Jr., Harriet Dashnow, Michael C. Zody, Matt Loose, Miten Jain, Evan E. Eichler, and Danny E. Miller. "High-coverage nanopore sequencing of samples from the 1000 Genomes Project to build a comprehensive catalog of human genetic variation." Genome Research 34, no. 11 (2024): 2061-2073.

[61] Javier Herrero , Matthieu Muffato , Kathryn Beal , Stephen Fitzgerald , Leo Gordon , Miguel Pignatelli , Albert J. Vilella , Stephen M. J. Searle , Ridwan Amode , Simon Brent , William Spooner , Eugene Kulesha , Andrew Yates , Paul Flicek. Ensembl comparative genomics resources. Database, 2016:bav096, 2016.

[62] Prüfer K, De Filippo C, Grote S, Mafessoni F, Korlević P, Hajdinjak M, Vernot B, Skov L, Hsieh P, Peyrégne S, Reher D. A high-coverage Neandertal genome from Vindija Cave in Croatia. Science, 358(6363):655-8, 2017.

[63] Mafessoni F, Grote S, De Filippo C, Slon V, Kolobova KA, Viola B, Markin SV, Chintalapati M, Peyrégne S, Skov L, Skoglund P. A high-coverage Neandertal genome from Chagyrskaya Cave. Proceedings of the National Academy of Sciences, 117(26):15132-6, 2020.

[64] Pruitt KD, Tatusova T, Maglott DR. NCBI reference sequences (RefSeq): a curated non-redundant sequence database of genomes, transcripts and proteins. Nucleic acids research, 35(suppl_1):D61-5, 2007.

[65] Pablo Cingolani, Adrian Platts, Le Lily Wang, Melissa Coon, Tung Nguyen, Luan Wang, Susan J Land, Xiangyi Lu, and Douglas M Ruden. A program for annotating and predicting the effects of single nucleotide polymorphisms, snpeff: Snps in the genome of drosophila melanogaster strain w1118; iso-2; iso-3. fly, 6(2):80–92, 2012.

[66] CL Scheib, Hongjie Li, Tariq Desai, Vivian Link, Christopher Kendall, Genevieve Dewar, Peter William Griffith, Alexander Mörseburg, John R Johnson, Amiee Potter, Susan L. Kerr, Phillip Endicott, John Lindo,Marc Haber, Yali Xue, Chris Tyler-Smith, Manjinder S. Sandhu, Joseph G. Lorenz, Tori D. Randall, Zuzana Faltyskova, Luca Pagani, Petr Danecek, Tamsin C. O'Connell, Patricia Martz, Alan S. Boraas, Brian F. Byrd, Alan Leventhal, Rosemary Cambra, Ronald Williamson, Louis Lesage, Brian Holguin, Ernestine Ygnacio-De Soto, JohnTommy Rosas, Mait Metspalu, Jay T. Stock, Andrea Manica, Aylwyn Scally, Daniel Wegmann, Ripan S. Malhi, Toomas Kivisild. Ancient human parallel lineages within North America contributed to a coastal expansion. Science, 360(6392):1024–1027, 2018.

[67] John Lindo, Randall Haas, Courtney Hofman, Mario Apata, Mauricio Moraga, Ricardo A Verdugo, James T Watson, Carlos Viviano Llave, David Witonsky, Cynthia Beall, Christina Warinner, John Novembre, Mark Aldenderfer, and Anna Di Rienzo. The genetic prehistory of the andean highlands 7000 years bp through european contact. Science advances, 4(11): eaau4921, 2018.

[68] Constanza De la Fuente, María C Ávila-Arcos, Jacqueline Galimany, Meredith L Carpenter, Julian R Homburger, Alejandro Blanco, Paloma Contreras, Diana Cruz D´avalos, Omar Reyes, Manuel San Roman, Andrés Moreno-Estrada, Paula F. Campos, Celeste Eng, Scott Huntsman, Esteban G. Burchard, Anna-Sapfo Malaspinas, Carlos D. Bustamante, Eske Willerslev, Elena Llop, Ricardo A. Verdugo, and Mauricio Moraga.

Genomic insights into the origin and diversification of late maritime hunter gatherers from the Chilean patagonia. Proceedings of the National Academy of Sciences, 115(17):E4006–E4012, 2018.

[69] J. Víctor Moreno-Mayar, Ben A. Potter, Lasse Vinner, Matthias Steinrücken, Simon Rasmussen, Jonathan Terhorst, John A. Kamm, Anders Albrechtsen, Anna-Sapfo Malaspinas, Martin Sikora, Joshua D. Reuther, Joel D. Irish, Ripan S. Malhi, Ludovic Orlando, Yun S. Song, Rasmus Nielsen, David J. Meltzer & Eske Willerslev. Terminal pleistocene alaskan genome reveals first founding population of native americans. Nature, 553 (7687):203, 2018.

[70] Morten Rasmussen, Sarah L. Anzick, Michael R. Waters, Pontus Skoglund, Michael DeGiorgio, Thomas W. Stafford Jr, Simon Rasmussen, Ida Moltke, Anders Albrechtsen, Shane M. Doyle, G. David Poznik, Valborg Gudmundsdottir, Rachita Yadav, Anna-Sapfo Malaspinas, Samuel Stockton White V, Morten E. Allentoft, Omar E. Cornejo, Kristiina Tambets, Anders Eriksson, Peter D. Heintzman, Monika Karmin, Thorfinn Sand Korneliussen, David J. Meltzer, Tracey L. Pierre, Jesper Stenderup, Lauri Saag, Vera M. Warmuth, Margarida C. Lopes, Ripan S. Malhi, Søren Brunak, Thomas Sicheritz-Ponten, Ian Barnes, Matthew Collins, Ludovic Orlando, Francois Balloux, Andrea Manica, Ramneek Gupta, Mait Metspalu, Carlos D. Bustamante, Mattias Jakobsson, Rasmus Nielsen & Eske Willerslev. The genome of a late pleistocene human from a clovis burial site in western montana. Nature, 506(7487): 225, 2014.

[71] Viridiana Villa-Islas, Alan Izarraras-Gomez, Maximilian Larena, Elizabeth Mej´ıa Perez Campos, Marcela Sandoval-Velasco, Juan Esteban Rodríguez Rodríguez, Miriam Bravo-Lopez, Barbara Moguel, Rosa Fregel, Ernesto Garfias-Morales, Jazeps Medina Tretmanis, David Alberto Velázquez-Ramírez,Alberto Herrera-Muñóz, Karla Sandoval, Maria A. Nieves-Colón, Gabriela Zepeda García Moreno,Fernando A. Villanea, Eugenia Fernández Villanueva Medina, Ramiro Aguayo-Haro,Cristina Valdiosera, Alexander G. Ioannidis, Andrés Moreno-Estrada, Flora Jay,Emilia Huerta-Sanchez, J. Víctor Moreno-Mayar, Federico Sánchez-Quinto, María C. Ávila-Arco. Demographic history and genetic structure in pre hispanic central mexico. Science, 380(6645):eadd6142, 2023.

[77] Xin Yi, Yu Liang, Emilia Huerta-Sanchez, Xin Jin, Zha Xi Ping Cuo, John E. Pool, Xun Xu, Hui Jiang, Nicolas Vinckenbosch, Thorfinn Sand Korneliussen, Hancheng Zheng, Tao Liu, Weiming He, Kui Li, Ruibang Luo, Xifang Nie, Honglong Wu, Meiru Zhao, Hongzhi Cao, Jing Zou, Ying Shan, Shuzheng Li, Qi Yang, Asan, Peixiang Ni, Geng Tian, Junming Xu, Xiao Liu, Tao Jiang, Renhua Wu, Guangyu Zhou, Meifang Tang, Junjie Qin, Tong Wang, Shuijian Feng, Guohong Li, Huasang, Jiangbai Luosang, Wei Wang, Fang Chen, Yading Wang, Xiaoguang Zheng, Zhuo Li, Zhuoma Bianba, Ge Yang, Xinping Wang, Shuhui Tang, Guoyi Gao, Yong Chen, Zhen Luo, Lamu Gusang, Zheng Cao, Qinghui Zhang, Weihan Ouyang, Xiaoli Ren, Huiqing Liang, Huisong Zheng, Yebo Huang, Jingxiang Li, Lars Bolund, Karsten Kristiansen, Yingrui Li, Yong Zhang, Xiuqing Zhang, Ruiqiang Li, Songgang Li, Huanming Yang, Rasmus Nielsen, Jun Wang, and Jian Wang. Sequencing of 50 human exomes reveals adaptation to high altitude. science, 329(5987):75–78, 2010.

[78] Gaurav Bhatia, Nick Patterson, Sriram Sankararaman, and Alkes L Price. Estimating and interpreting fst: the impact of rare variants. Genome research, 23(9):1514–1521, 2013.

[79] Voight BF, Kudaravalli S, Wen X, Pritchard JK. A map of recent positive selection in the human genome. PLoS biology, 4(3):e72, 2006.

[80] International HapMap Consortium. A second generation human haplotype map of over 3.1 million SNPs. Nature, 449(7164):851, 2007.

[81] Pauli Virtanen, Ralf Gommers, Travis E Oliphant, Matt Haberland, Tyler Reddy, David Cournapeau, Evgeni Burovski, Pearu Peterson, Warren Weckesser, Jonathan Bright, Stéfan J. van der Walt, Matthew Brett, Joshua Wilson, K. Jarrod Millman, Nikolay Mayorov, Andrew R. J. Nelson, Eric Jones, Robert Kern, Eric Larson, C J Carey, İlhan Polat, Yu Feng, Eric W. Moore, Jake VanderPlas, Denis Laxalde, Josef Perktold, Robert Cimrman, Ian Henriksen, E. A. Quintero, Charles R. Harris, Anne M. Archibald, Antônio

H. Ribeiro, Fabian Pedregosa, Paul van Mulbregt & SciPy 1.0 Contributors. Scipy 1.0: fundamental algorithms for scientific computing in python. Nature methods, 17(3):261–272, 2020.

[88] Jennifer Harrow, Adam Frankish, Jose M Gonzalez, Electra Tapanari, Mark Diekhans, Felix Kokocinski, Bronwen L Aken, Daniel Barrell, Amonida Zadissa, Stephen Searle, If Barnes, Alexandra Bignell, Veronika Boychenko, Toby Hunt, Mike Kay, Gaurab Mukherjee, Jeena Rajan, Gloria Despacio-Reyes, Gary Saunders, Charles Steward, Rachel Harte, Michael Lin, Cédric Howald, Andrea Tanzer, Thomas Derrien, Jacqueline Chrast, Nathalie Walters, Suganthi Balasubramanian, Baikang Pei, Michael Tress, Jose Manuel Rodriguez, Iakes Ezkurdia, Jeltje van Baren, Michael Brent, David Haussler, Manolis Kellis, Alfonso Valencia, Alexandre Reymond, Mark Gerstein, Roderic Guigó and Tim J. Hubbard. Gencode: the reference human genome annotation for the encode project. Genome research, 22(9):1760–1774, 2012.

[89] Ryan N Gutenkunst, Ryan D Hernandez, Scott H Williamson, and Carlos D Bustamante. Inferring the joint demographic history of multiple populations from multidimensional snp frequency data. PLoS genetics, 5 (10):e1000695, 2009.

[90] Richard E. Green, Johannes Krause, Adrian W. Briggs, Tomislav Maricic, Udo Stenzel, Martin Kircher, Nick Patterson, Heng Li, Weiwei Zhai, Markus Hsi-Yang Fritz, Nancy F. Hansen, Eric Y. Durand, Anna-Sapfo Malaspinas, Jeffrey D. Jensen, Tomas Marques-Bonet, Can Alkan, Kay Prüfer, Matthias Meyer, Hernán A. Burbano, Jeffrey M. Good, Rigo Schultz, Ayinuer Aximu-Petri, Anne Butthof, Barbara Höber, Barbara Höffner, Madlen Siegemund, Antje Weihmann, Chad Nusbaum, Eric S. Lander, Carsten Russ, Nathaniel Novod, Jason Affourtit, Michael Egholm, Christine Verna, Pavao Rudan, Dejana Brajkovic, Željko Kucan, Ivan Gušic, Vladimir B. Doronichev, Liubov V. Golovanova, Carles Lalueza-Fox, Marco de la Rasilla, Javier Fortea, Antonio Rosas, Ralf W. Schmitz, Philip L. F. Johnson, Evan E. Eichler, Daniel Falush, Ewan Birney, James C. Mullikin, Montgomery Slatkin, Rasmus Nielsen, Janet Kelso, Michael Lachmann, David Reich, and Svante Pääbo. A draft sequence of the Neandertal genome. Science 328, (5979): 710-722, 2010.

[91] Simon Gravel, Brenna M Henn, Ryan N Gutenkunst, Amit R Indap, Ga bor T Marth, Andrew G Clark, Fuli Yu, Richard A Gibbs, 1000 Genomes Project, Carlos D Bustamante, et al. Demographic history and rare allele sharing among human populations. Proceedings of the National Academy of Sciences, 108(29):11983–11988, 2011.

[92] David Reich, Richard E Green, Martin Kircher, Johannes Krause, Nick Patterson, Eric Y Durand, Bence Viola, Adrian W Briggs, Udo Stenzel, Philip LF Johnson, Tomislav Maricic, Jeffrey M. Good, Tomas Marques-Bonet, Can Alkan, Qiaomei Fu, Swapan Mallick, Heng Li, Matthias Meyer, Evan E. Eichler, Mark Stoneking, Michael Richards, Sahra Talamo, Michael V. Shunkov, Anatoli P. Derevianko, Jean-Jacques Hublin, Janet Kelso, Montgomery Slatkin & Svante Pääbo. Genetic history of an archaic hominin group from denisova cave in siberia. Nature, 468(7327):1053–1060, 2010.

[93] Bernard Y Kim, Christian D Huber, and Kirk E Lohmueller. Inference of the distribution of selection coefficients for new nonsynonymous mutations using large samples. Genetics, 206(1):345–361, 2017.

[94] Alistair Miles and NJ Harding. scikit-allel: A python package for exploring and analysing genetic variation data, 2016.

[95] Guy S Jacobs, Georgi Hudjashov, Lauri Saag, Pradiptajati Kusuma, Chelzie C Darusallam, Daniel J Lawson, Mayukh Mondal, Luca Pagani, Fran¸cois-Xavier Ricaut,

Mark Stoneking, Mait Metspalu, Herawati Sudoyo, J. Stephen Lansing, Murray P. Cox. Multiple deeply divergent denisovan ancestries in papuans. Cell, 2019.

[96] Pontus Skoglund & Matthias Jakobsson. Archaic human ancestry in East Asia. Proceedings of the National Academy of Sciences, 108(45), 18301-18306, 2011.

[97] Hamid, I., Korunes, K. L., Beleza, S., & Goldberg, A. Rapid adaptation to malaria facilitated by admixture in the human population of Cabo Verde. Elife, 10, e63177, 2021.

[98] Robinson, James T., Helga Thorvaldsdóttir, Wendy Winckler, Mitchell Guttman, Eric S. Lander, Gad Getz, and Jill P. Mesirov. "Integrative genomics viewer." Nature biotechnology 29, no. 1: 24-26, 2011.

[99] Prüfer K, Racimo F, Patterson N, Jay F, Sankararaman S, Sawyer S, Heinze A, Renaud G, Sudmant PH, De Filippo C, Li H. The complete genome sequence of a Neanderthal from the Altai Mountains. Nature;505(7481):43-9, 2014.
