## Supplemental Tables for "The *MUC19* Gene: An Evolutionary History of Recurrent Introgression and Natural Selection"

### Supplementary Tables & Datasets

**Table S1. Frequency of introgressed tracts overlapping *MUC19* among 1000 Genomes Project super populations.** The frequency of introgressed tracts—i.e., the number of introgressed tracts normalized by the total number of chromosomes—and the mean tract length stratified by super population: Admixed Americans (AMR), South Asians (SAS), East Asians (EAS), Europeans (EUR), and Africans (AFR). Introgressed tracts at *MUC19* are significantly enriched in AMR individuals (Fisher's Exact Test, Odds Ratio: 2.354, *P-value*: $2.144e^{-12}$), with AMR populations exhibiting a higher proportion of introgressed tracts compared to non-AMR populations, excluding AFR (Proportions $Z$-Test, $Z$-statistic: 7.441, *P-value*: $5.011e^{-14}$). Note that a "—" denotes that there are no introgressed tracts overlapping *MUC19* for that group.

Table S1:

| Super Population | Total Number of Chromosomes | Number of Introgressed Tracts | Introgressed Tract Frequency | Mean Tract Length |
|---|---|---|---|---|
| AMR | 694 | 127 | 0.183 | 609692.913 |
| SAS | 978 | 145 | 0.148 | 266268.966 |
| EAS | 1008 | 88 | 0.087 | 349102.273 |
| EUR | 1006 | 27 | 0.027 | 467037.037 |
| AFR | 1008 | 0 | 0 | — |

**Table S2. Frequency of introgressed tracts overlapping *MUC19* among 1000 Genomes Project populations.** The frequency of introgressed tracts—i.e., the number of introgressed tracts normalized by the total number of chromosomes—and the mean tract length for each population stratified by super population: Admixed Americans (AMR), South Asians (SAS), East Asians (EAS), Europeans (EUR), and Africans (AFR). Note that a "—" denotes that there are no introgressed tracts overlapping *MUC19* for that group.

Table S2:

| Super Population | Population | Total Number of Chromosomes | Number of Introgressed Tracts | Introgressed Tract Frequency | Mean Tract Length |
|---|---|---|---|---|---|
| AMR | MXL | 128 | 39 | 0.305 | 710666.667 |
| AMR | PEL | 170 | 42 | 0.247 | 574714.286 |
| AMR | CLM | 188 | 23 | 0.122 | 443304.348 |
| AMR | PUR | 208 | 23 | 0.111 | 668739.13 |
| SAS | BEB | 172 | 34 | 0.198 | 289941.176 |
| SAS | STU | 204 | 33 | 0.162 | 220878.788 |
| SAS | ITU | 204 | 35 | 0.172 | 227800 |
| SAS | PJL | 192 | 18 | 0.094 | 238277.778 |
| SAS | GIH | 206 | 25 | 0.121 | 368000 |
| EAS | CHB | 206 | 11 | 0.053 | 413454.545 |
| EAS | KHV | 198 | 22 | 0.111 | 357818.182 |
| EAS | CHS | 210 | 13 | 0.062 | 324692.308 |
| EAS | JPT | 208 | 14 | 0.067 | 245500 |
| EAS | CDX | 186 | 28 | 0.151 | 380107.143 |
| EUR | TSI | 214 | 13 | 0.061 | 458769.231 |
| EUR | CEU | 198 | 3 | 0.015 | 612333.333 |
| EUR | IBS | 214 | 7 | 0.033 | 520000 |

Continued on next page

Table S2: (Continued)

| Super Population | Population | Total Number of Chromosomes | Number of Introgressed Tracts | Introgressed Tract Frequency | Mean Tract Length |
|---|---|---|---|---|---|
| EUR | GBR | 182 | 1 | 0.005 | 343000 |
| EUR | FIN | 198 | 3 | 0.015 | 275333.333 |
| AFR | LWK | 198 | 0 | 0 | — |
| AFR | GWD | 226 | 0 | 0 | — |
| AFR | MSL | 170 | 0 | 0 | — |
| AFR | ESN | 198 | 0 | 0 | — |
| AFR | YRI | 216 | 0 | 0 | — |

**Table S3. Number of Denisovan-specific SNPs within the focal 742kb *MUC19* region among non-African populations in the 1000 Genomes Project.**
The number of Denisovan-specific SNPs—i.e., SNPs rare or absent in African populations ($< 1\%$), present in the non-African population ($> 1\%$), and uniquely shared with the Denisovan—observed in the focal 742kb region for each non-African population, stratified by super population: Admixed Americans (AMR), South Asians (SAS), East Asians (EAS), and Europeans (EUR). For each non-African population, the mean ($\mu$), standard deviation ($\sigma$), standard error of the mean ($SEM$), and 95% confidence intervals ($\pm CI_{95\%}$) of the Denisovan-specific SNP genomic background distribution used to compute the *P-value* are also reported. The *P-value* represents the proportion of non-overlapping 742kb windows of comparable effective sequence length where the number of Denisovan-specific SNPs is greater than or equal to what is observed at the focal 742kb region. A *P-value* less than 0.05 is considered statistically significant.

Table S3:

| Super Population | Population | Focal 742kb Region (Denisovan-specific SNPs) | 742kb Non-overlapping Windows ($\mu$) | 742kb Non-overlapping Windows ($\sigma$) | 742kb Non-overlapping Windows ($SEM$) | 742kb Non-overlapping Windows ($\pm CI_{95\%}$) | $P-value$ |
|---|---|---|---|---|---|---|---|
| AMR | MXL | 135 | 1.75 | 5.623 | 0.1 | 0.196 | $< 3.164e^{-4}$ |
| AMR | PEL | 135 | 1.663 | 5.824 | 0.104 | 0.203 | $< 3.164e^{-4}$ |
| AMR | CLM | 135 | 2.001 | 6.004 | 0.107 | 0.209 | $< 3.164e^{-4}$ |
| AMR | PUR | 135 | 1.608 | 4.909 | 0.087 | 0.171 | $< 3.164e^{-4}$ |
| SAS | BEB | 135 | 4.687 | 10.991 | 0.196 | 0.383 | $3.164e^{-4}$ |
| SAS | STU | 135 | 3.724 | 9.675 | 0.172 | 0.337 | $3.164e^{-4}$ |
| SAS | ITU | 135 | 3.767 | 9.997 | 0.178 | 0.349 | $3.164e^{-4}$ |
| SAS | PJL | 135 | 4.422 | 10.531 | 0.187 | 0.367 | $3.164e^{-4}$ |
| SAS | GIH | 135 | 3.39 | 8.954 | 0.159 | 0.312 | $3.164e^{-4}$ |
| EAS | CHB | 135 | 2.757 | 9.943 | 0.177 | 0.347 | $6.327e^{-4}$ |
| EAS | KHV | 135 | 3.297 | 10.724 | 0.191 | 0.374 | $9.491e^{-4}$ |
| EAS | CHS | 135 | 2.693 | 10.166 | 0.181 | 0.355 | $9.491e^{-4}$ |

Continued on next page

Table S3: (Continued)

| Super Population | Population | Focal 742kb Region (Denisovan-specific SNPs) | 742kb Non-overlapping Windows ($\mu$) | 742kb Non-overlapping Windows ($\sigma$) | 742kb Non-overlapping Windows ($SEM$) | 742kb Non-overlapping Windows ($\pm CI_{95\%}$) | $P-value$ |
|---|---|---|---|---|---|---|---|
| EAS | JPT | 135 | 2.481 | 8.651 | 0.154 | 0.302 | $3.164e^{-4}$ |
| EAS | CDX | 135 | 3.079 | 10.265 | 0.183 | 0.358 | $9.491e^{-4}$ |
| EUR | TSI | 135 | 1.321 | 4.473 | 0.08 | 0.156 | $< 3.164e^{-4}$ |
| EUR | CEU | 135 | 1.633 | 5.242 | 0.093 | 0.183 | $< 3.164e^{-4}$ |
| EUR | IBS | 135 | 1.368 | 4.821 | 0.086 | 0.168 | $< 3.164e^{-4}$ |
| EUR | GBR | 0 | 1.548 | 5.149 | 0.092 | 0.18 | $> 0.99968$ |
| EUR | FIN | 2 | 1.83 | 5.443 | 0.097 | 0.19 | 0.238 |

**Table S4. Number of Denisovan-specific SNPs within the focal 72kb *MUC19* region among non-African populations in the 1000 Genomes Project.**
The number of Denisovan-specific SNPs—i.e., SNPs rare or absent in African populations ($< 1\%$), present in the non-African population ($> 1\%$), and uniquely shared with the Denisovan—observed in the focal 72kb region for each non-African population, stratified by super population: Admixed Americans (AMR), South Asians (SAS), East Asians (EAS), and Europeans (EUR). For each non-African population, the mean ($\mu$), standard deviation ($\sigma$), standard error of the mean ($SEM$), and 95% confidence intervals ($\pm CI_{95\%}$) of the Denisovan-specific SNP genomic background distribution used to compute the *P-value* are also reported. The *P-value* represents the proportion of non-overlapping 72kb windows of comparable effective sequence length where the number of Denisovan-specific SNPs is greater than or equal to what is observed at the focal 72kb region. A *P-value* less than 0.05 is considered statistically significant.

Table S4:

| Super Population | Population | Focal 72kb Region (Denisovan-specific SNPs) | 72kb Non-overlapping Windows ($\mu$) | 72kb Non-overlapping Windows ($\sigma$) | 72kb Non-overlapping Windows ($SEM$) | 72kb Non-overlapping Windows ($\pm CI_{95\%}$) | $P-value$ |
|---|---|---|---|---|---|---|---|
| AMR | MXL | 135 | 0.178 | 1.536 | 0.009 | 0.018 | $< 3.389e^{-5}$ |
| AMR | PEL | 135 | 0.168 | 1.503 | 0.009 | 0.017 | $< 3.389e^{-5}$ |
| AMR | CLM | 135 | 0.201 | 1.631 | 0.009 | 0.019 | $< 3.389e^{-5}$ |
| AMR | PUR | 135 | 0.162 | 1.441 | 0.008 | 0.016 | $< 3.389e^{-5}$ |
| SAS | BEB | 135 | 0.467 | 2.449 | 0.014 | 0.028 | $< 3.389e^{-5}$ |
| SAS | STU | 135 | 0.371 | 2.12 | 0.012 | 0.024 | $< 3.389e^{-5}$ |
| SAS | ITU | 135 | 0.368 | 2.159 | 0.013 | 0.025 | $< 3.389e^{-5}$ |
| SAS | PJL | 135 | 0.433 | 2.292 | 0.013 | 0.026 | $< 3.389e^{-5}$ |
| SAS | GIH | 135 | 0.334 | 2.064 | 0.012 | 0.024 | $< 3.389e^{-5}$ |
| EAS | CHB | 135 | 0.28 | 2.01 | 0.012 | 0.023 | $< 3.389e^{-5}$ |
| EAS | KHV | 135 | 0.336 | 2.153 | 0.013 | 0.025 | $< 3.389e^{-5}$ |
| EAS | CHS | 135 | 0.275 | 2.009 | 0.012 | 0.023 | $< 3.389e^{-5}$ |

Continued on next page

Table S4: (Continued)

| Super Population | Population | Focal 72kb Region (Denisovan-specific SNPs) | 72kb Non-overlapping Windows ($\mu$) | 72kb Non-overlapping Windows ($\sigma$) | 72kb Non-overlapping Windows ($SEM$) | 72kb Non-overlapping Windows ($\pm CI_{95\%}$) | $P - value$ |
|---|---|---|---|---|---|---|---|
| EAS | JPT | 135 | 0.256 | 1.907 | 0.011 | 0.022 | $< 3.389e^{-5}$ |
| EAS | CDX | 135 | 0.312 | 2.068 | 0.012 | 0.024 | $< 3.389e^{-5}$ |
| EUR | TSI | 135 | 0.129 | 1.291 | 0.008 | 0.015 | $< 3.389e^{-5}$ |
| EUR | CEU | 135 | 0.157 | 1.388 | 0.008 | 0.016 | $< 3.389e^{-5}$ |
| EUR | IBS | 135 | 0.134 | 1.358 | 0.008 | 0.015 | $< 3.389e^{-5}$ |
| EUR | GBR | 0 | 0.151 | 1.283 | 0.007 | 0.015 | $> 0.9999966$ |
| EUR | FIN | 2 | 0.183 | 1.371 | 0.008 | 0.016 | 0.024 |

**Table S5. Number of Neanderthal-specific SNPs within the focal 742kb *MUC19* region among non-African populations in the 1000 Genomes Project.**

The number of Neanderthal-specific SNPs—i.e., SNPs rare or absent in African populations ($< 1\%$), present in the non-African population ($> 1\%$), and uniquely shared with at least one of the three high-coverage Neanderthals—observed in the focal 742kb region for each non-African population, stratified by super population: Admixed Americans (AMR), South Asians (SAS), East Asians (EAS), and Europeans (EUR). For each non-African population, the mean ($\mu$), standard deviation ($\sigma$), standard error of the mean ($SEM$), and 95% confidence intervals ($\pm CI_{95\%}$) of the Neanderthal-specific SNP genomic background distribution used to compute the *P-value* are also reported. The *P-value* represents the proportion of non-overlapping 742kb windows of comparable effective sequence length where the number of Neanderthal-specific SNPs is greater than or equal to what is observed at the focal 742kb region. A *P-value* less than 0.05 is considered statistically significant.

Table S5:

| Super Population | Population | Focal 742kb Region (Neanderthal-specific SNPs) | 742kb Non-overlapping Windows ($\mu$) | 742kb Non-overlapping Windows ($\sigma$) | 742kb Non-overlapping Windows ($SEM$) | 742kb Non-overlapping Windows ($\pm CI_{95\%}$) | $P-value$ |
|---|---|---|---|---|---|---|---|
| AMR | MXL | 80 | 35.183 | 45.21 | 0.804 | 1.577 | 0.159 |
| AMR | PEL | 95 | 31.07 | 42.422 | 0.755 | 1.48 | 0.097 |
| AMR | CLM | 95 | 40.198 | 48.246 | 0.858 | 1.683 | 0.146 |
| AMR | PUR | 95 | 35.2 | 45.629 | 0.812 | 1.592 | 0.117 |
| SAS | BEB | 95 | 52.222 | 56.294 | 1.001 | 1.964 | 0.231 |
| SAS | STU | 95 | 44.079 | 52.299 | 0.93 | 1.824 | 0.177 |
| SAS | ITU | 93 | 43.83 | 52.051 | 0.926 | 1.816 | 0.185 |
| SAS | PJL | 95 | 50.68 | 55.329 | 0.984 | 1.93 | 0.217 |
| SAS | GIH | 94 | 42.474 | 51.58 | 0.918 | 1.799 | 0.17 |
| EAS | CHB | 82 | 32.609 | 47.829 | 0.851 | 1.668 | 0.168 |
| EAS | KHV | 84 | 38.057 | 51.288 | 0.912 | 1.789 | 0.2 |
| EAS | CHS | 69 | 32.633 | 47.634 | 0.847 | 1.661 | 0.2 |

Continued on next page

Table S5: (Continued)

| Super Population | Population | Focal 742kb Region (Neanderthal-specific SNPs) | 742kb Non-overlapping Windows ($\mu$) | 742kb Non-overlapping Windows ($\sigma$) | 742kb Non-overlapping Windows ($SEM$) | 742kb Non-overlapping Windows ($\pm CI_{95\%}$) | $P-value$ |
|---|---|---|---|---|---|---|---|
| EAS | JPT | 83 | 32.509 | 47.424 | 0.844 | 1.654 | 0.164 |
| EAS | CDX | 70 | 35.868 | 49.539 | 0.881 | 1.728 | 0.22 |
| EUR | TSI | 87 | 34.63 | 45.715 | 0.813 | 1.595 | 0.142 |
| EUR | CEU | 86 | 38.689 | 48.181 | 0.857 | 1.681 | 0.168 |
| EUR | IBS | 87 | 34.864 | 46.257 | 0.823 | 1.613 | 0.138 |
| EUR | GBR | 33 | 37.39 | 47.164 | 0.839 | 1.645 | 0.388 |
| EUR | FIN | 23 | 39.594 | 47.67 | 0.848 | 1.663 | 0.482 |

**Table S6. Number of Neanderthal-specific SNPs within the focal 72kb *MUC19* region among non-African populations in the 1000 Genomes Project.**
The number of Neanderthal-specific SNPs—i.e., SNPs rare or absent in African populations ($< 1\%$), present in the non-African population ($> 1\%$), and uniquely shared with at least one of the three high-coverage Neanderthals—observed in the focal 72kb region for each non-African population, stratified by super population: Admixed Americans (AMR), South Asians (SAS), East Asians (EAS), and Europeans (EUR). For each non-African population, the mean ($\mu$), standard deviation ($\sigma$), standard error of the mean ($SEM$), and 95% confidence intervals ($\pm CI_{95\%}$) of the Neanderthal-specific SNP genomic background distribution used to compute the *P-value* are also reported. The *P-value* represents the proportion of non-overlapping 72kb windows of comparable effective sequence length where the number of Neanderthal-specific SNPs is greater than or equal to what is observed at the focal 72kb region. A *P-value* less than 0.05 is considered statistically significant.

Table S6:

| Super Population | Population | Focal 72kb Region (Neanderthal-specific SNPs) | 72kb Non-overlapping Windows ($\mu$) | 72kb Non-overlapping Windows ($\sigma$) | 72kb Non-overlapping Windows ($SEM$) | 72kb Non-overlapping Windows ($\pm CI_{95\%}$) | $P-value$ |
|---|---|---|---|---|---|---|---|
| AMR | MXL | 4 | 3.427 | 7.031 | 0.041 | 0.08 | 0.263 |
| AMR | PEL | 4 | 3.025 | 6.599 | 0.038 | 0.075 | 0.235 |
| AMR | CLM | 4 | 3.914 | 7.441 | 0.043 | 0.085 | 0.296 |
| AMR | PUR | 4 | 3.427 | 7.039 | 0.041 | 0.08 | 0.262 |
| SAS | BEB | 4 | 5.11 | 8.377 | 0.049 | 0.096 | 0.375 |
| SAS | STU | 4 | 4.306 | 7.852 | 0.046 | 0.09 | 0.32 |
| SAS | ITU | 4 | 4.292 | 7.801 | 0.045 | 0.089 | 0.32 |
| SAS | PJL | 4 | 4.965 | 8.337 | 0.049 | 0.095 | 0.364 |
| SAS | GIH | 4 | 4.159 | 7.776 | 0.045 | 0.089 | 0.311 |
| EAS | CHB | 4 | 3.178 | 6.906 | 0.04 | 0.079 | 0.238 |
| EAS | KHV | 4 | 3.709 | 7.365 | 0.043 | 0.084 | 0.275 |
| EAS | CHS | 4 | 3.177 | 6.878 | 0.04 | 0.078 | 0.238 |

Continued on next page

Table S6: (Continued)

| Super Population | Population | Focal 72kb Region (Neanderthal-specific SNPs) | 72kb Non-overlapping Windows ($\mu$) | 72kb Non-overlapping Windows ($\sigma$) | 72kb Non-overlapping Windows ($SEM$) | 72kb Non-overlapping Windows ($\pm CI_{95\%}$) | $P - value$ |
|---|---|---|---|---|---|---|---|
| EAS | JPT | 4 | 3.161 | 6.907 | 0.04 | 0.079 | 0.236 |
| EAS | CDX | 4 | 3.509 | 7.218 | 0.042 | 0.082 | 0.26 |
| EUR | TSI | 4 | 3.39 | 7.037 | 0.041 | 0.08 | 0.257 |
| EUR | CEU | 4 | 3.779 | 7.359 | 0.043 | 0.084 | 0.284 |
| EUR | IBS | 4 | 3.401 | 7.123 | 0.041 | 0.081 | 0.256 |
| EUR | GBR | 0 | 3.654 | 7.272 | 0.042 | 0.083 | $> 0.9999966$ |
| EUR | FIN | 0 | 3.854 | 7.408 | 0.043 | 0.085 | $> 0.9999966$ |

**Table S7. $U_{AFR:B:DEN}(1\%, 30\%, 100\%)$ values for the focal 742kb *MUC19* region among non-African populations in the 1000 Genomes Project.**

The $U_{AFR:B:DEN}(1\%, 30\%, 100\%)$ values observed in the focal 742kb region for each non-African population, stratified by super population: Admixed Americans (AMR), South Asians (SAS), East Asians (EAS), and Europeans (EUR). The $U_{AFR:B:DEN}(1\%, 30\%, 100\%)$ statistic quantifies the number of sites where: 1) the Denisovan allele is found in the homozygous state, 2) the Denisovan allele is at low frequency ($< 1\%$) in the African super population, and 3) the Denisovan allele is at high frequency ($> 30\%$) in the non-African population ($B$). For each non-African population, the mean ($\mu$), standard deviation ($\sigma$), standard error of the mean ($SEM$), and 95% confidence intervals ($\pm CI_{95\%}$) of the $U_{AFR:B:DEN}(1\%, 30\%, 100\%)$ genomic background distribution used to compute the *P-value* are also reported. The *P-value* represents the proportion of non-overlapping 742kb windows of comparable effective sequence length where the $U_{AFR:B:DEN}(1\%, 30\%, 100\%)$ value is greater than or equal to that observed at the focal 742kb region. A *P-value* less than 0.05 is considered statistically significant.

Table S7:

| Super Population | Population ($B$) | Focal 742kb Region ($U_{AFR:B:DEN}$ $(1\%, 30\%, 100\%)$) | 742kb Non-overlapping Windows ($\mu$) | 742kb Non-overlapping Windows ($\sigma$) | 742kb Non-overlapping Windows ($SEM$) | 742kb Non-overlapping Windows ($\pm CI_{95\%}$) | $P-value$ |
|---|---|---|---|---|---|---|---|
| AMR | MXL | 136 | 0.312 | 2.779 | 0.049 | 0.097 | $< 3.139e^{-4}$ |
| AMR | PEL | 0 | 0.667 | 4.358 | 0.077 | 0.151 | $> 0.999686$ |
| AMR | CLM | 0 | 0.156 | 1.829 | 0.032 | 0.064 | $> 0.999686$ |
| AMR | PUR | 0 | 0.125 | 1.64 | 0.029 | 0.057 | $> 0.999686$ |
| SAS | BEB | 0 | 0.175 | 1.829 | 0.032 | 0.064 | $> 0.999686$ |
| SAS | STU | 0 | 0.13 | 1.299 | 0.023 | 0.045 | $> 0.999686$ |
| SAS | ITU | 0 | 0.181 | 1.896 | 0.034 | 0.066 | $> 0.999686$ |
| SAS | PJL | 0 | 0.162 | 1.772 | 0.031 | 0.062 | $> 0.999686$ |
| SAS | GIH | 0 | 0.175 | 1.844 | 0.033 | 0.064 | $> 0.999686$ |
| EAS | CHB | 0 | 0.438 | 2.281 | 0.04 | 0.079 | $> 0.999686$ |
| EAS | KHV | 0 | 0.415 | 2.308 | 0.041 | 0.08 | $> 0.999686$ |

Continued on next page

Table S7: (Continued)

| Super Population | Population ($B$) | Focal 742kb Region ($U_{AFR:B:DEN}$ $(1\%, 30\%, 100\%)$) | 742kb Non-overlapping Windows ($\mu$) | 742kb Non-overlapping Windows ($\sigma$) | 742kb Non-overlapping Windows ($SEM$) | 742kb Non-overlapping Windows ($\pm CI_{95\%}$) | $P-value$ |
|---|---|---|---|---|---|---|---|
| EAS | CHS | 0 | 0.416 | 2.257 | 0.04 | 0.078 | $> 0.999686$ |
| EAS | JPT | 0 | 0.444 | 2.482 | 0.044 | 0.086 | $> 0.999686$ |
| EAS | CDX | 0 | 0.452 | 2.405 | 0.043 | 0.084 | $> 0.999686$ |
| EUR | TSI | 0 | 0.215 | 2.73 | 0.048 | 0.095 | $> 0.999686$ |
| EUR | CEU | 0 | 0.213 | 2.19 | 0.039 | 0.076 | $> 0.999686$ |
| EUR | IBS | 0 | 0.211 | 2.191 | 0.039 | 0.076 | $> 0.999686$ |
| EUR | GBR | 0 | 0.241 | 2.417 | 0.043 | 0.084 | $> 0.999686$ |
| EUR | FIN | 0 | 0.283 | 2.965 | 0.053 | 0.103 | $> 0.999686$ |

**Table S8. $U_{AFR:B:DEN}(1\%, 30\%, 100\%)$ values for the focal 72kb *MUC19* region among non-African populations in the 1000 Genomes Project.**

The $U_{AFR:B:DEN}(1\%, 30\%, 100\%)$ values observed in the focal 72kb region for each non-African population, stratified by super population: Admixed Americans (AMR), South Asians (SAS), East Asians (EAS), and Europeans (EUR). The $U_{AFR:B:DEN}(1\%, 30\%, 100\%)$ statistic quantifies the number of sites where: 1) the Denisovan allele is found in the homozygous state, 2) the Denisovan allele is at low frequency ($< 1\%$) in the African super population, and 3) the Denisovan allele is at high frequency ($> 30\%$) in the non-African population ($B$). For each non-African population, the mean ($\mu$), standard deviation ($\sigma$), standard error of the mean ($SEM$), and 95% confidence intervals ($\pm CI_{95\%}$) of the $U_{AFR:B:DEN}(1\%, 30\%, 100\%)$ genomic background distribution used to compute the *P-value* are also reported. The *P-value* represents the proportion of non-overlapping 72kb windows of comparable effective sequence length where the $U_{AFR:B:DEN}(1\%, 30\%, 100\%)$ value is greater than or equal to that observed at the focal 72kb region. A *P-value* less than 0.05 is considered statistically significant.

Table S8:

| Super Population | Population ($B$) | Focal 72kb Region ($U_{AFR:B:DEN}$ $(1\%, 30\%, 100\%)$) | 72kb Non-overlapping Windows ($\mu$) | 72kb Non-overlapping Windows ($\sigma$) | 72kb Non-overlapping Windows ($SEM$) | 72kb Non-overlapping Windows ($\pm CI_{95\%}$) | $P-value$ |
|---|---|---|---|---|---|---|---|
| AMR | MXL | 136 | 0.03 | 0.86 | 0.005 | 0.01 | $< 3.284e^{-5}$ |
| AMR | PEL | 0 | 0.066 | 0.932 | 0.005 | 0.01 | $> 0.999967$ |
| AMR | CLM | 0 | 0.016 | 0.504 | 0.003 | 0.006 | $> 0.999967$ |
| AMR | PUR | 0 | 0.012 | 0.457 | 0.003 | 0.005 | $> 0.999967$ |
| SAS | BEB | 0 | 0.017 | 0.482 | 0.003 | 0.005 | $> 0.999967$ |
| SAS | STU | 0 | 0.013 | 0.38 | 0.002 | 0.004 | $> 0.999967$ |
| SAS | ITU | 0 | 0.016 | 0.473 | 0.003 | 0.005 | $> 0.999967$ |
| SAS | PJL | 0 | 0.016 | 0.476 | 0.003 | 0.005 | $> 0.999967$ |
| SAS | GIH | 0 | 0.018 | 0.487 | 0.003 | 0.005 | $> 0.999967$ |
| EAS | CHB | 0 | 0.041 | 0.546 | 0.003 | 0.006 | $> 0.999967$ |
| EAS | KHV | 0 | 0.041 | 0.646 | 0.004 | 0.007 | $> 0.999967$ |

Continued on next page

Table S8: (Continued)

| Super Population | Population ($B$) | Focal 72kb Region ($U_{AFR:B:DEN}$ $(1\%, 30\%, 100\%)$) | 72kb Non-overlapping Windows ($\mu$) | 72kb Non-overlapping Windows ($\sigma$) | 72kb Non-overlapping Windows ($SEM$) | 72kb Non-overlapping Windows ($\pm CI_{95\%}$) | $P - value$ |
|---|---|---|---|---|---|---|---|
| EAS | CHS | 0 | 0.04 | 0.556 | 0.003 | 0.006 | $> 0.999967$ |
| EAS | JPT | 0 | 0.042 | 0.624 | 0.004 | 0.007 | $> 0.999967$ |
| EAS | CDX | 0 | 0.044 | 0.66 | 0.004 | 0.007 | $> 0.999967$ |
| EUR | TSI | 0 | 0.022 | 0.596 | 0.003 | 0.007 | $> 0.999967$ |
| EUR | CEU | 0 | 0.021 | 0.544 | 0.003 | 0.006 | $> 0.999967$ |
| EUR | IBS | 0 | 0.02 | 0.551 | 0.003 | 0.006 | $> 0.999967$ |
| EUR | GBR | 0 | 0.023 | 0.601 | 0.003 | 0.007 | $> 0.999967$ |
| EUR | FIN | 0 | 0.029 | 0.649 | 0.004 | 0.007 | $> 0.999967$ |

**Table S9. $Q95_{AFR:B:DEN}(1\%, 100\%)$ values for the focal 72kb and 742kb *MUC19* regions among non-African populations in the 1000 Genomes Project.**
The $Q95_{AFR:B:DEN}(1\%, 100\%)$ values observed in the focal 72kb and 742kb regions for each non-African population, stratified by super population: Admixed Americans (AMR), South Asians (SAS), East Asians (EAS), and Europeans (EUR). The $Q95_{AFR:B:DEN}(1\%, 100\%)$ statistic quantifies $95^{th}$ percentile of the Denisovan alleles in the non-African population ($B$) conditioned on: 1) the Denisovan allele is found in the homozygous state, and 2) the Denisovan allele is at low frequency ($< 1\%$) in the African super population.

Table S9:

| Super Population | Population ($B$) | Focal 72kb Region ($Q95_{AFR:B:DEN}$ $(1\%, 100\%)$) | Focal 742kb Region ($Q95_{AFR:B:DEN}$ $(1\%, 100\%)$) |
|---|---|---|---|
| AMR | MXL | 0.305 | 0.305 |
| AMR | PEL | 0.218 | 0.218 |
| AMR | CLM | 0.074 | 0.074 |
| AMR | PUR | 0.087 | 0.087 |
| SAS | BEB | 0.157 | 0.157 |
| SAS | STU | 0.113 | 0.113 |
| SAS | ITU | 0.108 | 0.108 |
| SAS | PJL | 0.052 | 0.052 |
| SAS | GIH | 0.097 | 0.097 |
| EAS | CHB | 0.034 | 0.034 |
| EAS | KHV | 0.091 | 0.091 |
| EAS | CHS | 0.04 | 0.038 |
| EAS | JPT | 0.029 | 0.029 |
| EAS | CDX | 0.124 | 0.124 |
| EUR | TSI | 0.051 | 0.051 |
| EUR | CEU | 0.015 | 0.015 |

Continued on next page

Table S9: (Continued)

| Super Population | Population ($B$) | Focal 72kb Region ($Q95_{AFR:B:DEN}$ (1%, 100%)) | Focal 742kb Region ($Q95_{AFR:B:DEN}$ (1%, 100%)) |
|---|---|---|---|
| EUR | IBS | 0.033 | 0.033 |
| EUR | GBR | 0.005 | 0.005 |
| EUR | FIN | 0.005 | 0.005 |

**Table S10. $PBS_{A:CHB:CEU}$ values for the focal 742kb *MUC19* region among different Indigenous American ancestry (IAA) demes in MXL.**

The $PBS_{A:CHB:CEU}$ values observed in the focal 742kb region for different IAA-partitioned demes in MXL: all individuals in the MXL population ($n = 64$; $A$ = All MXL Individuals), MXL individuals with genome-wide IAA $> 50\%$ ($n = 27$; $A$ = MXL Individuals $> 50\%$ IAA), and MXL individuals with genome-wide IAA $< 50\%$ ($n = 37$; $A$ = MXL Individuals $< 50\%$ IAA). The $PBS_{A:CHB:CEU}$ statistic uses the logarithmic transformation of pairwise $F_{ST}$ estimates to measure the branch length in deme $A$ since its divergence from the two control populations: CHB and CEU. For each IAA-partitioned deme, the mean ($\mu$), standard deviation ($\sigma$), standard error of the mean ($SEM$), and 95% confidence intervals ($\pm CI_{95\%}$) of the $PBS_{A:CHB:CEU}$ genomic background distribution used to compute the *P-value* are also reported. The *P-value* represents the proportion of non-overlapping 742kb windows of comparable effective sequence length where the $PBS_{A:CHB:CEU}$ value is greater than or equal to that observed at the focal 742kb region. A *P-value* less than 0.05 is considered statistically significant.

Table S10:

| $PBS_{A:CHB:CEU}$ | Focal 742kb Region ($PBS$) | 742kb Non-overlapping Windows ($\mu$) | 742kb Non-overlapping Windows ($\sigma$) | 742kb Non-overlapping Windows ($SEM$) | 742kb Non-overlapping Windows ($\pm CI_{95\%}$) | $P - value$ |
|---|---|---|---|---|---|---|
| $A =$ All MXL Inds. | 0.066 | 0.005 | 0.011 | $1.949e^{-4}$ | $3.822e^{-4}$ | 0.004 |
| $A =$ MXL Inds. $> 50\%$ IAA | 0.133 | 0.018 | 0.024 | $4.128e^{-4}$ | $8.093e^{-4}$ | 0.003 |
| $A =$ MXL Inds. $< 50\%$ IAA | 0.029 | 0.002 | 0.007 | $1.231e^{-4}$ | $2.414e^{-4}$ | 0.01 |

**Table S11. $PBS_{A:CHB:CEU}$ values for the focal 72kb *MUC19* region among different Indigenous American ancestry (IAA) demes in MXL.**

The $PBS_{A:CHB:CEU}$ values observed in the focal 72kb region for different IAA-partitioned demes in MXL: all individuals in the MXL population ($n = 64$; $A$ = All MXL Individuals), MXL individuals with genome-wide IAA > 50% ($n = 27$; $A$ = MXL Individuals > 50% IAA), and MXL individuals with genome-wide IAA < 50% ($n = 37$; $A$ = MXL Individuals < 50% IAA). The $PBS_{A:CHB:CEU}$ statistic uses the logarithmic transformation of pairwise $F_{ST}$ estimates to measure the branch length in deme $A$ since its divergence from the two control populations: CHB and CEU. For each IAA-partitioned deme, the mean ($\mu$), standard deviation ($\sigma$), standard error of the mean ($SEM$), and 95% confidence intervals ($\pm CI_{95\%}$) of the $PBS_{A:CHB:CEU}$ genomic background distribution used to compute the *P-value* are also reported. The *P-value* represents the proportion of non-overlapping 72kb windows of comparable effective sequence length where the $PBS_{A:CHB:CEU}$ value is greater than or equal to that observed at the focal 72kb region. A *P-value* less than 0.05 is considered statistically significant.

Table S11:

| $PBS_{A:CHB:CEU}$ | Focal 72kb Region ($PBS$) | 72kb Non-overlapping Windows ($\mu$) | 72kb Non-overlapping Windows ($\sigma$) | 72kb Non-overlapping Windows ($SEM$) | 72kb Non-overlapping Windows ($\pm CI_{95\%}$) | $P - value$ |
|---|---|---|---|---|---|---|
| $A$ = All MXL Inds. | 0.127 | 0.009 | 0.019 | $1.040e^{-4}$ | $2.038e^{-4}$ | 0.002 |
| $A$ = MXL Inds. > 50% IAA | 0.217 | 0.023 | 0.036 | $2.033e^{-4}$ | $3.984e^{-4}$ | 0.003 |
| $A$ = MXL Inds. < 50% IAA | 0.072 | 0.005 | 0.013 | $7.435e^{-5}$ | $1.457e^{-4}$ | 0.006 |

**Table S12. Integrated Haplotype Scores (*iHS*) for the focal 742kb *MUC19* region among 1000 Genomes Project populations.**

The normalized $| iHS |$ scores observed at the focal 742kb region for each population stratified by super population: Admixed Americans (AMR), South Asians (SAS), East Asians (EAS), Europeans (EUR), and Africans (AFR). For each population, the normalized *iHS* scores were calculated for all SNPs with a minor allele frequency $> 5\%$. For each population, the table reports the total number of *iHS* scores observed, the number of extreme *iHS* scores (i.e., $| iHS |> 2$), the observed proportion of extreme *iHS* scores (calculated as the number of extreme *iHS* scores normalized by the total number of *iHS* scores), the $99^{th}$ percentile of the proportion of extreme *iHS* scores determined from the genomic background distribution (i.e., non-overlapping 742kb windows with more than 10 SNPs), the total number of 742kb windows in the genomic background distribution, the number of 742kb windows where the observed proportion of extreme *iHS* scores is greater than the genomic background distribution, and the percentile rank of the observed proportion of extreme *iHS* scores (i.e., the percentage of the genomic background distribution that is less than the observed proportion of extreme *iHS* scores). An observed proportion of extreme *iHS* scores greater than the $99^{th}$ percentile of the genomic background distribution is considered evidence of positive selection.

Table S12:

| Super Population | Population | Focal 742kb Region (Total SNPs) | Focal 742kb Region (SNPs with $\| iHS \|> 2$) | Focal 742kb Region (Prop. of SNPs with $\| iHS \|> 2$) | 742kb Non-overlapping Windows ($99^{th}$ Percentile) | 742kb Non-overlapping Windows (Total SNPs $> 10$) | Focal 742kb Region $>$ 742kb Non-overlapping Windows | Focal 742kb Region (Percentile Rank) |
|---|---|---|---|---|---|---|---|---|
| AMR | MXL | 2248 | 599 | 0.266 | 0.21 | 3595 | 3578 | 99.527 |
| AMR | PEL | 2158 | 162 | 0.075 | 0.28 | 3591 | 2983 | 83.069 |
| AMR | CLM | 2397 | 53 | 0.022 | 0.237 | 3596 | 1165 | 32.397 |
| AMR | PUR | 2407 | 68 | 0.028 | 0.214 | 3597 | 1503 | 41.785 |
| SAS | BEB | 2177 | 78 | 0.036 | 0.274 | 3595 | 2106 | 58.581 |
| SAS | STU | 2197 | 65 | 0.03 | 0.28 | 3596 | 1781 | 49.527 |
| SAS | ITU | 1976 | 23 | 0.012 | 0.26 | 3596 | 668 | 18.576 |
| SAS | PJL | 2064 | 32 | 0.016 | 0.272 | 3596 | 945 | 26.279 |
| SAS | GIH | 2195 | 24 | 0.011 | 0.262 | 3596 | 596 | 16.574 |

Continued on next page

Table S12: (Continued)

| Super Population | Population | Focal 742kb Region (Total SNPs) | Focal 742kb Region (SNPs with $\mid iHS \mid > 2$) | Focal 742kb Region (Prop. of SNPs with $\mid iHS \mid > 2$) | 742kb Non-overlapping Windows ($99^{th}$ Percentile) | 742kb Non-overlapping Windows (Total SNPs > 10) | Focal 742kb Region > 742kb Non-overlapping Windows | Focal 742kb Region (Percentile Rank) |
|---|---|---|---|---|---|---|---|---|
| EAS | CHB | 1591 | 48 | 0.03 | 0.233 | 3594 | 1795 | 49.944 |
| EAS | KHV | 2010 | 59 | 0.029 | 0.225 | 3594 | 1719 | 47.83 |
| EAS | CHS | 1586 | 40 | 0.025 | 0.249 | 3595 | 1529 | 42.531 |
| EAS | JPT | 1618 | 78 | 0.048 | 0.239 | 3594 | 2481 | 69.032 |
| EAS | CDX | 1955 | 76 | 0.039 | 0.223 | 3594 | 2158 | 60.045 |
| EUR | TSI | 2323 | 83 | 0.036 | 0.226 | 3594 | 1995 | 55.509 |
| EUR | CEU | 2090 | 66 | 0.032 | 0.24 | 3595 | 1824 | 50.737 |
| EUR | IBS | 1944 | 40 | 0.021 | 0.229 | 3595 | 1178 | 32.768 |
| EUR | GBR | 1986 | 58 | 0.029 | 0.214 | 3596 | 1766 | 49.11 |
| EUR | FIN | 2007 | 10 | 0.005 | 0.23 | 3594 | 217 | 6.038 |
| AFR | LWK | 2218 | 4 | 0.002 | 0.193 | 3598 | 12 | 0.334 |
| AFR | GWD | 2324 | 19 | 0.008 | 0.205 | 3597 | 94 | 2.613 |
| AFR | MSL | 2152 | 45 | 0.021 | 0.208 | 3598 | 634 | 17.621 |
| AFR | ESN | 2321 | 28 | 0.012 | 0.194 | 3597 | 280 | 7.784 |
| AFR | YRI | 2277 | 34 | 0.015 | 0.221 | 3598 | 362 | 10.061 |

**Table S13. Integrated Haplotype Scores (*iHS*) for the focal 72kb *MUC19* region among 1000 Genomes Project populations.**

The normalized $| iHS |$ scores observed at the focal 742kb region for each population stratified by super population: Admixed Americans (AMR), South Asians (SAS), East Asians (EAS), Europeans (EUR), and Africans (AFR). For each population, the normalized *iHS* scores were calculated for all SNPs with a minor allele frequency $> 5\%$. For each population, the table reports the total number of *iHS* scores observed, the number of extreme *iHS* scores (i.e., $| iHS | > 2$), the observed proportion of extreme *iHS* scores (calculated as the number of extreme *iHS* scores normalized by the total number of *iHS* scores), the $99^{th}$ percentile of the proportion of extreme *iHS* scores determined from the genomic background distribution (i.e., non-overlapping 72kb windows with more than 10 SNPs), the total number of 72kb windows in the genomic background distribution, the number of 72kb windows where the observed proportion of extreme *iHS* scores is greater than the genomic background distribution, and the percentile rank of the observed proportion of extreme *iHS* scores (i.e., the percentage of the genomic background distribution that is less than the observed proportion of extreme *iHS* scores). An observed proportion of extreme *iHS* scores greater than the $99^{th}$ percentile of the genomic background distribution is considered evidence of positive selection.

Table S13:

| Super Population | Population | Focal 72kb Region (Total SNPs) | Focal 72kb Region (SNPs with $\| iHS \| > 2$) | Focal 72kb Region (Prop. of SNPs with $\| iHS \| > 2$) | 72kb Non-overlapping Windows ($99^{th}$ Percentile) | 72kb Non-overlapping Windows (Total SNPs $> 10$) | Focal 72kb Region > 72kb Non-overlapping Windows | Focal 72kb Region (Percentile Rank) |
|---|---|---|---|---|---|---|---|---|
| AMR | MXL | 425 | 229 | 0.539 | 0.462 | 36411 | 36202 | 99.426 |
| AMR | PEL | 403 | 86 | 0.213 | 0.492 | 36293 | 34299 | 94.506 |
| AMR | CLM | 431 | 7 | 0.016 | 0.473 | 36459 | 19972 | 54.779 |
| AMR | PUR | 441 | 1 | 0.002 | 0.447 | 36474 | 11772 | 32.275 |
| SAS | BEB | 420 | 6 | 0.014 | 0.526 | 36388 | 21536 | 59.184 |
| SAS | STU | 413 | 0 | 0 | 0.524 | 36376 | 0 | 0 |
| SAS | ITU | 403 | 0 | 0 | 0.52 | 36383 | 0 | 0 |
| SAS | PJL | 413 | 0 | 0 | 0.516 | 36409 | 0 | 0 |
| SAS | GIH | 415 | 0 | 0 | 0.5 | 36354 | 0 | 0 |

Continued on next page

Table S13: (Continued)

| Super Population | Population | Focal 72kb Region (Total SNPs) | Focal 72kb Region (SNPs with $\mid iHS \mid > 2$) | Focal 72kb Region (Prop. of SNPs with $\mid iHS \mid > 2$) | 72kb Non-overlapping Windows ($99^{th}$ Percentile) | 72kb Non-overlapping Windows (Total SNPs $> 10$) | Focal 72kb Region > 72kb Non-overlapping Windows | Focal 72kb Region (Percentile Rank) |
|---|---|---|---|---|---|---|---|---|
| EAS | CHB | 168 | 1 | 0.006 | 0.49 | 36193 | 17256 | 47.678 |
| EAS | KHV | 394 | 1 | 0.003 | 0.495 | 36283 | 15128 | 41.694 |
| EAS | CHS | 169 | 3 | 0.018 | 0.496 | 36208 | 22884 | 63.202 |
| EAS | JPT | 180 | 0 | 0 | 0.49 | 36192 | 0 | 0 |
| EAS | CDX | 393 | 0 | 0 | 0.479 | 36219 | 0 | 0 |
| EUR | TSI | 452 | 13 | 0.029 | 0.478 | 36375 | 24780 | 68.124 |
| EUR | CEU | 232 | 6 | 0.026 | 0.483 | 36393 | 24602 | 67.601 |
| EUR | IBS | 217 | 4 | 0.018 | 0.487 | 36372 | 22140 | 60.871 |
| EUR | GBR | 226 | 2 | 0.009 | 0.487 | 36348 | 18741 | 51.56 |
| EUR | FIN | 234 | 1 | 0.004 | 0.48 | 36374 | 14941 | 41.076 |
| AFR | LWK | 217 | 0 | 0 | 0.401 | 36570 | 0 | 0 |
| AFR | GWD | 219 | 1 | 0.005 | 0.43 | 36562 | 9341 | 25.548 |
| AFR | MSL | 208 | 1 | 0.005 | 0.424 | 36576 | 9130 | 24.962 |
| AFR | ESN | 223 | 1 | 0.004 | 0.432 | 36565 | 9545 | 26.104 |
| AFR | YRI | 211 | 0 | 0 | 0.449 | 36554 | 0 | 0 |

**Table S14. Frequency of introgressed tracts overlapping *MUC19* short-read repeat region among 1000 Genomes Project super populations.**

The frequency of introgressed tracts—i.e., the number of introgressed tracts normalized by the total number of chromosomes—and the mean tract length stratified by super population: Admixed Americans (AMR), South Asians (SAS), East Asians (EAS), Europeans (EUR), and Africans (AFR). Introgressed tracts at the *MUC19* short-read repeat region (hg19, Chr12:40876395-40885001) are significantly enriched in AMR individuals (Fisher's Exact Test, Odds Ratio: 5.940, *P-value*: $1.953e^{-31}$), with AMR populations exhibiting a higher proportion of introgressed tracts compared to non-AMR populations, excluding AFR (Proportions $Z$-Test, $Z$-statistic: 13.269, *P-value*: $1.742e^{-40}$). Note that a "—" denotes that there are no introgressed tracts overlapping short-read repeat region for that group.

Table S14:

| Super Population | Total Number of Chromosomes | Number of Introgressed Tracts | Introgressed Tract Frequency | Mean Tract Length |
|---|---|---|---|---|
| AMR | 694 | 109 | 0.157 | 547954.128 |
| SAS | 978 | 27 | 0.028 | 283666.667 |
| EAS | 1008 | 58 | 0.058 | 335327.586 |
| EUR | 1006 | 6 | 0.006 | 460500 |
| AFR | 1008 | 0 | 0 | — |

**Table S15. Frequency of introgressed tracts overlapping *MUC19* short-read repeat region among 1000 Genomes Project populations.**
The frequency of introgressed tracts—i.e., the number of introgressed tracts normalized by the total number of chromosomes—and the mean tract length for each population stratified by super population: Admixed Americans (AMR), South Asians (SAS), East Asians (EAS), Europeans (EUR), and Africans (AFR). Note that a ”—” denotes that there are no introgressed tracts overlapping short-read repeat region for that group.

Table S15:

| Super Population | Population | Total Number of Chromosomes | Number of Introgressed Tracts | Introgressed Tract Frequency | Mean Tract Length |
|---|---|---|---|---|---|
| AMR | MXL | 128 | 36 | 0.281 | 653250 |
| AMR | PEL | 170 | 38 | 0.224 | 511368.421 |
| AMR | CLM | 188 | 18 | 0.096 | 381166.667 |
| AMR | PUR | 208 | 17 | 0.082 | 583352.941 |
| SAS | BEB | 172 | 4 | 0.023 | 427750 |
| SAS | STU | 204 | 3 | 0.015 | 136333.333 |
| SAS | ITU | 204 | 8 | 0.039 | 271250 |
| SAS | PJL | 192 | 7 | 0.036 | 223142.857 |
| SAS | GIH | 206 | 5 | 0.024 | 361400 |
| EAS | CHB | 206 | 9 | 0.044 | 363666.667 |
| EAS | KHV | 198 | 14 | 0.071 | 306214.286 |
| EAS | CHS | 210 | 10 | 0.048 | 283000 |
| EAS | JPT | 208 | 8 | 0.038 | 326750 |
| EAS | CDX | 186 | 17 | 0.091 | 379117.647 |
| EUR | TSI | 214 | 3 | 0.014 | 334666.667 |
| EUR | CEU | 198 | 2 | 0.01 | 592500 |

Continued on next page

Table S15: (Continued)

| Super Population | Population | Total Number of Chromosomes | Number of Introgressed Tracts | Introgressed Tract Frequency | Mean Tract Length |
|---|---|---|---|---|---|
| EUR | IBS | 214 | 1 | 0.005 | 574000 |
| EUR | GBR | 182 | 0 | 0 | — |
| EUR | FIN | 198 | 0 | 0 | — |
| AFR | LWK | 198 | 0 | 0 | — |
| AFR | GWD | 226 | 0 | 0 | — |
| AFR | MSL | 170 | 0 | 0 | — |
| AFR | ESN | 198 | 0 | 0 | — |
| AFR | YRI | 216 | 0 | 0 | — |

**Table S16. Estimated repeat copies of the *MUC19* 30bp variable number tandem repeat from short-read sequencing data among 1000 Genomes Project populations.**
Summary of the distribution of repeat copy numbers for *MUC19* per population stratified by super populations: Admixed Americans (AMR), South Asians (SAS), East Asians (EAS), Europeans (EUR), and Africans (AFR). For each population, the mean ($\mu$), standard deviation ($\sigma$), standard error of the mean ($SEM$), and 95% confidence intervals ($\pm CI_{95\%}$) of the repeat copy distribution are reported. We also report the total number of individuals, the number of individuals with elevated repeat copies (i.e., those with $>$ 487 repeat copies), and the proportion of individuals with elevated repeat copies per population. Populations are sorted by their mean repeat copy number.

Table S16:

| Super Population | Population | Number of Inds. | Repeat Copies ($\mu$) | Repeat Copies ($\sigma$) | Repeat Copies ($SEM$) | Repeat Copies ($\pm CI_{95\%}$) | Number of Inds. with Elevated Repeats | Proportion of Inds. with Elevated Repeats |
|---|---|---|---|---|---|---|---|---|
| AMR | MXL | 64 | 492.844 | 147.128 | 18.536 | 37.042 | 32 | 0.5 |
| AMR | PEL | 85 | 409.941 | 140.779 | 15.36 | 30.546 | 24 | 0.282 |
| AMR | CLM | 94 | 398.883 | 101.087 | 10.482 | 20.816 | 20 | 0.213 |
| AMR | PUR | 104 | 391.346 | 95.726 | 9.432 | 18.706 | 15 | 0.144 |
| EAS | CDX | 93 | 365.032 | 61.999 | 6.464 | 12.838 | 6 | 0.065 |
| EAS | KHV | 99 | 361.263 | 50.017 | 5.052 | 10.026 | 2 | 0.02 |
| EUR | TSI | 107 | 358.804 | 44.556 | 4.328 | 8.58 | 1 | 0.009 |
| EUR | CEU | 99 | 357.051 | 48.05 | 4.854 | 9.632 | 1 | 0.01 |
| SAS | GIH | 103 | 356.903 | 51.543 | 5.104 | 10.123 | 3 | 0.029 |
| EAS | JPT | 104 | 354.788 | 38.662 | 3.809 | 7.555 | 1 | 0.01 |
| SAS | PJL | 96 | 353.615 | 47.445 | 4.868 | 9.664 | 1 | 0.01 |
| EAS | CHS | 105 | 352.076 | 47.589 | 4.667 | 9.254 | 0 | 0 |

Continued on next page

Table S16: (Continued)

| Super Population | Population | Number of Inds. | Repeat Copies ($\mu$) | Repeat Copies ($\sigma$) | Repeat Copies ($SEM$) | Repeat Copies ($\pm CI_{95\%}$) | Number of Inds. with Elevated Repeats | Proportion of Inds. with Elevated Repeats |
|---|---|---|---|---|---|---|---|---|
| EUR | IBS | 107 | 351.925 | 39.528 | 3.839 | 7.612 | 0 | 0 |
| AFR | YRI | 108 | 350.361 | 38.85 | 3.756 | 7.445 | 0 | 0 |
| SAS | ITU | 102 | 350.235 | 47.46 | 4.722 | 9.368 | 3 | 0.029 |
| SAS | STU | 102 | 349.824 | 44.253 | 4.403 | 8.735 | 2 | 0.02 |
| AFR | ESN | 99 | 348.273 | 38.94 | 3.934 | 7.806 | 1 | 0.01 |
| EUR | GBR | 91 | 348.187 | 45.544 | 4.801 | 9.537 | 1 | 0.011 |
| SAS | BEB | 86 | 347.372 | 39.937 | 4.332 | 8.613 | 1 | 0.012 |
| EAS | CHB | 103 | 343.961 | 45.212 | 4.477 | 8.879 | 2 | 0.019 |
| AFR | MSL | 85 | 343.529 | 34.437 | 3.757 | 7.472 | 0 | 0 |
| EUR | FIN | 99 | 341.899 | 35.976 | 3.634 | 7.212 | 0 | 0 |
| AFR | GWD | 113 | 341.726 | 40.331 | 3.811 | 7.551 | 2 | 0.018 |
| AFR | LWK | 99 | 340.707 | 34.856 | 3.521 | 6.987 | 0 | 0 |

**Table S17. Estimated repeat copies of the *MUC19* 30bp variable number tandem repeat from short-read sequencing data among 1000 Genomes Project super populations.**
Summary of the distribution of repeat copy numbers for *MUC19* per super population: Admixed Americans (AMR), South Asians (SAS), East Asians (EAS), Europeans (EUR), and Africans (AFR). For each super population, the mean ($\mu$), standard deviation ($\sigma$), standard error of the mean ($SEM$), and 95% confidence intervals ($\pm CI_{95\%}$) of the repeat copy distribution are reported. We also report the total number of individuals, the number of individuals with elevated repeat copies (i.e., those with $> 487$ repeat copies), and the proportion of individuals with elevated repeat copies per super population. Super populations are sorted by their mean repeat copy number.

Table S17:

| Super Population | Number of Inds. | Repeat Copies ($\mu$) | Repeat Copies ($\sigma$) | Repeat Copies ($SEM$) | Repeat Copies ($\pm CI_{95\%}$) | Number of Inds. with Elevated Repeats | Proportion of Inds. with Elevated Repeats |
|---|---|---|---|---|---|---|---|
| AMR | 347 | 416.663 | 125.383 | 6.741 | 13.258 | 91 | 0.262 |
| EAS | 504 | 355.173 | 49.514 | 2.208 | 4.338 | 11 | 0.022 |
| EUR | 503 | 351.748 | 43.32 | 1.933 | 3.799 | 3 | 0.006 |
| SAS | 489 | 351.714 | 46.597 | 2.109 | 4.145 | 10 | 0.02 |
| AFR | 504 | 344.966 | 37.938 | 1.692 | 3.323 | 3 | 0.006 |

**Table S18. Distribution of repeat copies for different focal groups among all individuals in the 1000 Genome Project.** Summary of the distribution of repeat copy numbers for individuals with at least one introgressed tract overlapping the repeat region vs individuals with no introgressed tracts overlapping the repeat region and admixed American vs non-admixed American individuals. For each focal group, the mean ($\mu$), standard deviation ($\sigma$), standard error of the mean ($SEM$), and 95% confidence intervals ($\pm CI_{95\%}$) of the repeat copy distribution are reported.

Table S18:

| Group 1 | Group 2 | Group 1 Repeat Copies ($\mu$) | Group 2 Repeat Copies ($\mu$) | Group 1 Repeat Copies ($\sigma$) | Group 2 Repeat Copies ($\sigma$) | Group 1 Repeat Copies ($SEM$) | Group 2 Repeat Copies ($SEM$) | Group 1 Repeat Copies ($\pm CI_{95\%}$) | Group 2 Repeat Copies ($\pm CI_{95\%}$) |
|---|---|---|---|---|---|---|---|---|---|
| All Inds.: $\geq 1$ Overlapping Tract | All Inds.: No Overlapping Tract | 517.886 | 347.16 | 114.083 | 39.384 | 8.41 | 0.847 | 16.593 | 1.661 |
| All AMR Inds. | All non-AMR Inds. | 416.663 | 350.894 | 125.383 | 44.69 | 6.741 | 1 | 13.258 | 1.96 |

**Table S19. Comparisons of the distributions of repeat copies between different focal groups among all individuals in the 1000 Genome Project.**
Comparing the distributions of repeat copy numbers between focal groups: individuals with at least one introgressed tract overlapping the repeat region vs individuals with no introgressed tracts overlapping the repeat region and admixed American vs non-admixed American individuals. For each comparison, the total number of individuals, the number of individuals with elevated repeat copies (i.e., those with $> 487$ repeat copies), the proportion of individuals with elevated repeat copies per population, and the results of the Proportions $Z$-Test, and Mann-Whitney $U$-Test is reported. A *P-value* less than 0.05 is considered statistically significant.

Table S19:

| Group 1 | Group 2 | Number of Inds. (Group 1) | Number of Inds. (Group 2) | Number of Inds. with Elevated Repeats (Group 1) | Number of Inds. with Elevated Repeats (Group 2) | Prop. of Inds. with Elevated Repeats (Group 1) | Prop. of Inds. with Elevated Repeats (Group 2) | $Z-Statistic$ | $P-value$ (Prop. $Z$-Test) | $U-Statistic$ | $P-value$ (Mann-Whitney $U$-Test) |
|---|---|---|---|---|---|---|---|---|---|---|---|
| All Inds.: $\geq 1$ Overlapping Tract | All Inds.: No Overlapping Tract | 185 | 2162 | 102 | 16 | 0.551 | 0.007 | 32.496 | $6.015e^{-232}$ | 375108.5 | $1.597e^{-87}$ |
| All AMR Inds. | All non-AMR Inds. | 347 | 2000 | 91 | 27 | 0.262 | 0.013 | 19.575 | $1.266e^{-85}$ | 445495 | $1.424e^{-17}$ |

**Table S20. Distribution of repeat copies for different focal groups among individuals with an elevated number of repeat copies in the 1000 Genome Project.**
Summary of the distribution of repeat copy numbers for different focal groups of outlier individuals: individuals with at least one introgressed tract overlapping the repeat region vs individuals with no introgressed tracts overlapping the repeat region and admixed American vs non-admixed American individuals. For each focal group, the mean ($\mu$), standard deviation ($\sigma$), standard error of the mean ($SEM$), and 95% confidence intervals ($\pm CI_{95\%}$) of the repeat copy distribution are reported.

Table S20:

| Group 1 | Group 2 | Group 1 Repeat Copies ($\mu$) | Group 2 Repeat Copies ($\mu$) | Group 1 Repeat Copies ($\sigma$) | Group 2 Repeat Copies ($\sigma$) | Group 1 Repeat Copies ($SEM$) | Group 2 Repeat Copies ($SEM$) | Group 1 Repeat Copies ($\pm CI_{95\%}$) | Group 2 Repeat Copies ($\pm CI_{95\%}$) |
|---|---|---|---|---|---|---|---|---|---|
| Outlier Inds.: $\geq 1$ Overlapping Tract | Outlier Inds.: No Overlapping Tract | 594.48 | 510.688 | 92.117 | 16.116 | 9.166 | 4.161 | 18.183 | 8.869 |
| Outlier AMR Inds. | Outlier non-AMR Inds. | 600.956 | 523 | 94.368 | 33.26 | 9.947 | 6.523 | 19.762 | 13.408 |

**Table S21. Comparisons of the distributions of repeat copies between different focal groups among individuals with an elevated number of repeat copies in the 1000 Genome Project.**
Comparing the distributions of repeat copy numbers between focal groups of outlier individuals: individuals with at least one introgressed tract overlapping the repeat region vs individuals with no introgressed tracts overlapping the repeat region and admixed American vs non-admixed American individuals. For each comparison, the total number of individuals, the number of individuals with elevated repeat copies (i.e., those with $> 487$ repeat copies), the proportion of individuals with elevated repeat copies per population, and the results of the Proportions $Z$-Test, and Mann-Whitney $U$-Test is reported. A $P$*-value* less than 0.05 is considered statistically significant.

Table S21:

| Group 1 | Group 2 | Total Number of Inds. with Elevated Repeats | Number of Inds. (Group 1) | Number of Inds. (Group 2) | Prop. of Inds. (Group 1) | Prop. of Inds. (Group 2) | $Z-Statistic$ | $P-value$ (Prop. $Z$-Test) | $U-Statistic$ | $P-value$ (Mann-Whitney $U$-Test) |
|---|---|---|---|---|---|---|---|---|---|---|
| Outlier Inds.: $\geq 1$ Overlapping Tract | Outlier Inds.: No Overlapping Tract | 118 | 102 | 16 | 0.864 | 0.136 | 11.196 | $2.127e^{-29}$ | 1412.5 | $1.398e^{-6}$ |
| Outlier AMR Inds. | Outlier non-AMR Inds. | 118 | 91 | 27 | 0.771 | 0.229 | 8.332 | $3.971e^{-17}$ | 1988 | $5.789e^{-7}$ |

**Table S22. Correlation between the introgressed ancestry and repeat copies in the 1000 Genome Project.** Spearman's correlation coefficients and associated *P-values* between the number of introgressed tracts overlapping the repeat region and the number of repeat copies for all individuals in the 1000 Genome Project, individuals stratified by super population, and individuals stratified by population. A *P-value* less than 0.05 is considered statistically significant, and a "—" denotes that there are no introgressed tracts overlapping the repeat region for that group.

Table S22:

| Group | Spearman's $\rho$ | $P - value$ |
|---|---|---|
| TGP | 0.409 | $2.022e^{-95}$ |
| AFR | — | — |
| AMR | 0.736 | $1.741e^{-60}$ |
| SAS | 0.216 | $1.471e^{-6}$ |
| EAS | 0.532 | $3.914e^{-38}$ |
| EUR | 0.104 | 0.019 |
| LWK | — | — |
| GWD | — | — |
| MSL | — | — |
| ESN | — | — |
| YRI | — | — |
| MXL | 0.885 | $2.839e^{-22}$ |
| PEL | 0.812 | $3.815e^{-21}$ |
| CLM | 0.632 | $8.640e^{-12}$ |
| PUR | 0.528 | $8.119e^{-9}$ |
| BEB | 0.36 | $6.532e^{-4}$ |
| STU | 0.255 | 0.01 |

Continued on next page

Table S22: (Continued)

| Group | Spearman's $\rho$ | $P - value$ |
|---|---|---|
| ITU | 0.206 | 0.038 |
| PJL | 0.124 | 0.23 |
| GIH | 0.206 | 0.037 |
| CHB | 0.461 | $9.297e^{-7}$ |
| KHV | 0.586 | $1.887e^{-10}$ |
| CHS | 0.499 | $5.839e^{-8}$ |
| JPT | 0.419 | $9.370e^{-6}$ |
| CDX | 0.653 | $1.270e^{-12}$ |
| TSI | 0.043 | 0.66 |
| CEU | 0.244 | 0.015 |
| IBS | 0.116 | 0.233 |
| GBR | — | — |
| FIN | — | — |

**Table S23. Ancestry component distribution for the *MUC19* short-read repeat region among admixed American populations in 1000 Genomes Project.**
Summary of the per-individual ancestry proportions at the *MUC19* short-read repeat region (hg19, Chr12:40876395-40885001) for each of the admixed American populations in the 1000 Genomes project. For each ancestry component per population, the mean ($\mu$), standard deviation ($\sigma$), standard error of the mean ($SEM$), and 95% confidence intervals ($\pm CI_{95\%}$) of the per-individual ancestry proportion distribution are reported.

Table S23:

| Population | Ancestry Componenet | Ancestry Proportion ($\mu$) | Ancestry Proportion ($\sigma$) | Ancestry Proportion ($SEM$) | Ancestry Proportion ($\pm CI_{95\%}$) |
|---|---|---|---|---|---|
| MXL | Indigenous American | 0.602 | 0.344 | 0.043 | 0.087 |
| MXL | European | 0.336 | 0.331 | 0.042 | 0.083 |
| MXL | African | 0.055 | 0.156 | 0.02 | 0.039 |
| PEL | Indigenous American | 0.694 | 0.344 | 0.038 | 0.075 |
| PEL | European | 0.271 | 0.339 | 0.037 | 0.074 |
| PEL | African | 0.029 | 0.118 | 0.013 | 0.026 |
| CLM | Indigenous American | 0.282 | 0.306 | 0.032 | 0.063 |
| CLM | European | 0.649 | 0.341 | 0.035 | 0.07 |
| CLM | African | 0.064 | 0.182 | 0.019 | 0.037 |
| PUR | Indigenous American | 0.154 | 0.251 | 0.025 | 0.049 |
| PUR | European | 0.683 | 0.333 | 0.033 | 0.065 |
| PUR | African | 0.154 | 0.26 | 0.026 | 0.051 |

**Table S24. Correlation between ancestry components and repeat copies among Admixed American populations in the 1000 Genome Project.**
Spearman's correlation coefficients and associated *P-values* between the ancestry proportion (per ancestry component) and the number of repeat copies for all admixed American individuals in the 1000 Genome Project stratified by population. After correcting for three multiple comparisons per admixed American population—i.e., one per ancestry component—a *P-value* less than 0.0167 is considered statistically significant.

Table S24:

| Population | Ancestry Componenet | Spearman's $\rho$ | $P - value$ |
|---|---|---|---|
| MXL | Indigenous American | 0.438 | $2.940e^{-4}$ |
| MXL | European | -0.353 | 0.004 |
| MXL | African | -0.289 | 0.021 |
| PEL | Indigenous American | 0.391 | $2.178e^{-4}$ |
| PEL | European | -0.313 | 0.004 |
| PEL | African | -0.2 | 0.067 |
| CLM | Indigenous American | 0.553 | $7.378e^{-9}$ |
| CLM | European | -0.495 | $3.957e^{-7}$ |
| CLM | African | -0.047 | 0.651 |
| PUR | Indigenous American | 0.414 | $1.281e^{-5}$ |
| PUR | European | -0.216 | 0.028 |
| PUR | African | -0.127 | 0.198 |

**Table S25. Denisovan-specific coding mutations in the focal 72kb *MUC19* region.**
Allelic and functional information for the 11 Denisovan-specific coding mutations found within the focal 72kb region. Each variant's effect on protein structure is categorized using Grantham scores as follows: Grantham score $< 50$ = conservative, Grantham score 51–100 = moderately conservative, Grantham score 101–150 = moderately radical, and Grantham score $> 150$ = radical. The PhyloP score measures the conservation of the reference base-pair, with higher positive scores indicating stronger conservation.

Table S25:

| Chr12 Position (Hg19) | rsID | Mut. Type | Ref. Allele | Ref. Amino Acid | Denisovan Allele | Denisovan Amino Acid | Protein Position | Grantham Score | PhyloP Score |
|---|---|---|---|---|---|---|---|---|---|
| 40808672 | rs4768252 | Missense | C | Ser | T | Leu | 275 | 145 | -1.221 |
| 40808726 | rs4768253 | Missense | C | Thr | T | Met | 293 | 81 | -0.879 |
| 40809983 | rs142268259 | Missense | C | Asp | A | Glu | 410 | 45 | -0.061 |
| 40814107 | rs2114566 | Missense | A | Ile | G | Val | 511 | 29 | -0.189 |
| 40814197 | rs17467164 | Missense | G | Val | A | Ile | 541 | 29 | -0.336 |
| 40815060 | rs149221842 | Missense | C | His | T | Tyr | 572 | 83 | -0.569 |
| 40820403 | rs11564249 | Synonymous | C | Asp | T | Asp | 711 | 0 | 0.818 |
| 40821795 | rs61736852 | Missense | G | Glu | A | Lys | 766 | 56 | 0.581 |
| 40821871 | rs17467284 | Missense | G | Arg | T | Leu | 791 | 102 | 5.15 |
| 40826201 | rs11564125 | Missense | G | Val | A | Ile | 883 | 29 | -1.722 |
| 40826535 | rs11564104 | Synonymous | T | Ser | C | Ser | 939 | 0 | -0.035 |

**Table S26. Metadata for the ancient Indigenous American genomes.**
The location, sample ID, age, genomic coverage, and reference for the 23 ancient Indigenous American individuals that predate European colonization and the African slave trade used in this study.

Table S26:

| Location | ID | Age (years) | Coverage (x) | Reference |
|---|---|---|---|---|
| Alaska | USR1 | 11,500 | 17.0 | Moreno-Mayar et al. 2018 Nature |
| California | CR-01 | 1,111 | 8.36 | Scheib et al. 2018 Science |
| California | CT-01 | 387 | 6.68 | Scheib et al. 2018 Science |
| California | NC_C | 1,450 | 1.12 | Scheib et al. 2018 Science |
| California | PS-06 | 1,570 | 4.12 | Scheib et al. 2018 Science |
| California | SC-05 | 1,101 | 13.67 | Scheib et al. 2018 Science |
| California | SM-02 | 826 | 7.38 | Scheib et al. 2018 Science |
| California | SN-13 | 881 | 1.74 | Scheib et al. 2018 Science |
| California | SN-17 | 4,517 | 7.46 | Scheib et al. 2018 Science |
| California | SN-44 | 4,646 | 9.36 | Scheib et al. 2018 Science |
| Montana | Clovis | 12,500 | 14.4 | Rasmussen et al. 2014 Nature |

Continued on next page

Table S26: (Continued)

| Location | ID | Age (years) | Coverage (x) | Reference |
|---|---|---|---|---|
| Ontario | CK-13 | 4,267 | 3.02 | Scheib et al. 2018 Science |
| Patagonia | ERR2270782 | 1,000 | 7.8 | de la Fuente et al. 2018 PNAS |
| Patagonia | ERR2270783 | 1,000 | 3.5 | de la Fuente et al. 2018 PNAS |
| Patagonia | ERR2270784 | 1,000 | 1.7 | de la Fuente et al. 2018 PNAS |
| Patagonia | ERR2270785 | 1,000 | 9.1 | de la Fuente et al. 2018 PNAS |
| Peru | SRR8144644 | 1,800 | 3.34 | Lindo et al. 2018 Science Advances |
| Peru | SRR8144646 | 1,800 | 5.89 | Lindo et al. 2018 Science Advances |
| Peru | SRR8144647 | 1,800 | 6.51 | Lindo et al. 2018 Science Advances |
| Peru | SRR8144650 | 1,800 | 5.62 | Lindo et al. 2018 Science Advances |
| Querétaro | 2417J | 1,161 | 1.39 | Villa-Islas et al. 2023 Science |
| Querétaro | 2417Q | 1,257 | 4.74 | Villa-Islas et al. 2023 Science |
| Querétaro | 333B | 1,351 | 2.28 | Villa-Islas et al. 2023 Science |

**Table S27. Comparison of the per-individual frequency distributions for each Denisovan-specific missense mutation in the focal 72kb *MUC19* region between admixed American and non-admixed American individuals in 1000 Genomes Project**

Comparisons of the per-individual frequency distribution for each of the nine Denisovan-specific missense mutations in the focal 72kb *MUC19* region between the 347 admixed American (AMR) individuals ($n = 694$ chromosomes) and 1,496 non-admixed American (Non-AMR) individuals ($n = 2,992$ chromosomes), excluding African populations, in the 1000 Genomes Project. For each comparison, the number of chromosomes harboring the respective Denisovan-specific missense mutation, the frequency of the Denisovan-specific missense mutation, and the results of the Proportions $Z$-Test and Fisher's Exact Test are reported. A *P-value* less than 0.05 is considered statistically significant.

Table S27:

| Position | Denisovan-specific Mis. Mut. Count (AMR) | Denisovan-specific Mis. Freq. (AMR) | Denisovan-specific Mis. Mut. Count (Non-AMR) | Denisovan-specific Mis. Freq. (Non-AMR) | $Z-Statistic$ | $P-value$ (Prop. $Z$-Test) | Odds Ratio | $P-value$ (Fisher's Exact Test) |
|---|---|---|---|---|---|---|---|---|
| 40808672 | 109 | 0.157 | 187 | 0.062 | 8.259 | $7.374e^{-17}$ | 2.795 | $1.939e^{-14}$ |
| 40808726 | 108 | 0.156 | 190 | 0.064 | 8.02 | $5.269e^{-16}$ | 2.718 | $9.213e^{-14}$ |
| 40809983 | 108 | 0.156 | 186 | 0.062 | 8.187 | $1.337e^{-16}$ | 2.78 | $3.175e^{-14}$ |
| 40814107 | 108 | 0.156 | 186 | 0.062 | 8.187 | $1.337e^{-16}$ | 2.78 | $3.175e^{-14}$ |
| 40814197 | 108 | 0.156 | 188 | 0.063 | 8.103 | $2.670e^{-16}$ | 2.749 | $5.430e^{-14}$ |
| 40815060 | 108 | 0.156 | 186 | 0.062 | 8.187 | $1.337e^{-16}$ | 2.78 | $3.175e^{-14}$ |
| 40821795 | 107 | 0.154 | 180 | 0.06 | 8.328 | $4.116e^{-17}$ | 2.848 | $1.320e^{-14}$ |
| 40821871 | 107 | 0.154 | 180 | 0.06 | 8.328 | $4.116e^{-17}$ | 2.848 | $1.320e^{-14}$ |
| 40826201 | 108 | 0.156 | 181 | 0.06 | 8.399 | $2.248e^{-17}$ | 2.862 | $8.011e^{-15}$ |

**Table S28. Frequency of the Denisovan-specific missense mutations in the focal 72kb *MUC19* region among super populations in the 1000 Genomes Project.**
Frequencies of the nine Denisovan-specific missense mutations found within the focal 72kb *MUC19* region per super population in the 1000 Genomes Project.

Table S28:

| Super Population | 40808672 | 40808726 | 40809983 | 40814107 | 40814197 | 40815060 | 40821795 | 40821871 | 40826201 |
|---|---|---|---|---|---|---|---|---|---|
| AFR | 0.0 | 0.0 | 0.0 | 0.0 | 0.0 | 0.0 | 0.0 | 0.0 | 0.0 |
| AMR | 0.157 | 0.156 | 0.156 | 0.156 | 0.156 | 0.156 | 0.154 | 0.154 | 0.156 |
| SAS | 0.105 | 0.108 | 0.104 | 0.104 | 0.105 | 0.104 | 0.098 | 0.098 | 0.098 |
| EAS | 0.061 | 0.061 | 0.061 | 0.061 | 0.062 | 0.061 | 0.061 | 0.061 | 0.062 |
| EUR | 0.023 | 0.023 | 0.023 | 0.023 | 0.023 | 0.023 | 0.023 | 0.023 | 0.023 |

**Table S29. Frequency of the Denisovan-specific missense mutations in the focal 72kb *MUC19* region among populations in the 1000 Genomes Project.**
Frequencies of the nine Denisovan-specific missense mutations found within the focal 72kb *MUC19* region per population in the 1000 Genomes Project.

Table S29:

| Population | 40808672 | 40808726 | 40809983 | 40814107 | 40814197 | 40815060 | 40821795 | 40821871 | 40826201 |
|---|---|---|---|---|---|---|---|---|---|
| GWD | 0.0 | 0.0 | 0.0 | 0.0 | 0.0 | 0.0 | 0.0 | 0.0 | 0.0 |
| ESN | 0.0 | 0.0 | 0.0 | 0.0 | 0.0 | 0.0 | 0.0 | 0.0 | 0.0 |
| MSL | 0.0 | 0.0 | 0.0 | 0.0 | 0.0 | 0.0 | 0.0 | 0.0 | 0.0 |
| YRI | 0.0 | 0.0 | 0.0 | 0.0 | 0.0 | 0.0 | 0.0 | 0.0 | 0.0 |
| LWK | 0.0 | 0.0 | 0.0 | 0.0 | 0.0 | 0.0 | 0.0 | 0.0 | 0.0 |
| PUR | 0.087 | 0.087 | 0.087 | 0.087 | 0.087 | 0.087 | 0.087 | 0.087 | 0.091 |
| CLM | 0.08 | 0.074 | 0.074 | 0.074 | 0.074 | 0.074 | 0.069 | 0.069 | 0.069 |
| PEL | 0.218 | 0.218 | 0.218 | 0.218 | 0.218 | 0.218 | 0.218 | 0.218 | 0.218 |
| MXL | 0.305 | 0.305 | 0.305 | 0.305 | 0.305 | 0.305 | 0.305 | 0.305 | 0.305 |
| PJL | 0.052 | 0.052 | 0.052 | 0.052 | 0.052 | 0.052 | 0.052 | 0.052 | 0.052 |
| BEB | 0.157 | 0.169 | 0.157 | 0.157 | 0.157 | 0.157 | 0.14 | 0.14 | 0.14 |
| STU | 0.113 | 0.113 | 0.113 | 0.113 | 0.113 | 0.113 | 0.113 | 0.113 | 0.113 |
| ITU | 0.113 | 0.113 | 0.108 | 0.108 | 0.113 | 0.108 | 0.103 | 0.103 | 0.103 |
| GIH | 0.097 | 0.102 | 0.097 | 0.097 | 0.097 | 0.097 | 0.087 | 0.087 | 0.087 |
| CHS | 0.038 | 0.038 | 0.038 | 0.038 | 0.038 | 0.038 | 0.038 | 0.038 | 0.038 |
| CDX | 0.124 | 0.124 | 0.124 | 0.124 | 0.124 | 0.124 | 0.124 | 0.124 | 0.124 |
| KHV | 0.086 | 0.086 | 0.086 | 0.086 | 0.086 | 0.086 | 0.086 | 0.086 | 0.091 |
| CHB | 0.034 | 0.034 | 0.034 | 0.034 | 0.039 | 0.034 | 0.034 | 0.034 | 0.034 |

Continued on next page

Table S29: (Continued)

| Population | 40808672 | 40808726 | 40809983 | 40814107 | 40814197 | 40815060 | 40821795 | 40821871 | 40826201 |
|---|---|---|---|---|---|---|---|---|---|
| JPT | 0.029 | 0.029 | 0.029 | 0.029 | 0.029 | 0.029 | 0.029 | 0.029 | 0.029 |
| GBR | 0.005 | 0.005 | 0.005 | 0.005 | 0.005 | 0.005 | 0.005 | 0.005 | 0.005 |
| FIN | 0.005 | 0.005 | 0.005 | 0.005 | 0.005 | 0.005 | 0.005 | 0.005 | 0.005 |
| IBS | 0.033 | 0.033 | 0.033 | 0.033 | 0.033 | 0.033 | 0.033 | 0.033 | 0.033 |
| CEU | 0.015 | 0.015 | 0.015 | 0.015 | 0.015 | 0.015 | 0.015 | 0.015 | 0.015 |
| TSI | 0.051 | 0.051 | 0.051 | 0.051 | 0.051 | 0.051 | 0.051 | 0.051 | 0.051 |

**Table S30. Denisovan-specific missense mutation allele counts in the focal 72kb *MUC19* region among 23 ancient Indigenous American individuals.**
Allele counts (i.e., 0, 1, or 2) of the nine Denisovan-specific missense mutations found within the focal 72kb *MUC19* region for each of the 23 ancient Indigenous American individuals that predate European colonization and the African slave trade. A "—" denotes that the site did not pass quality control.

Table S30:

| ID | 40808672 | 40808726 | 40809983 | 40814107 | 40814197 | 40815060 | 40821795 | 40821871 | 40826201 |
|---|---|---|---|---|---|---|---|---|---|
| USR1 | 1 | 0 | 0 | 0 | 0 | 0 | 0 | 0 | 0 |
| CK-13 | — | 2 | 0 | — | 0 | — | — | — | — |
| Anzick | 1 | 0 | 0 | 0 | 0 | 0 | 0 | 0 | 0 |
| CR-01 | 2 | 2 | 2 | 2 | — | 2 | 2 | 2 | 2 |
| CT-01 | 0 | 0 | — | 0 | 0 | 0 | — | 0 | 0 |
| NC_C | — | 0 | — | — | — | — | 0 | 0 | — |
| PS-06 | 0 | 0 | 0 | — | — | 0 | 0 | 0 | — |
| SC-05 | 1 | 1 | 1 | 1 | 1 | 2 | 1 | 1 | 1 |
| SM-02 | 2 | 2 | 2 | 2 | 2 | — | — | 2 | 2 |
| SN-13 | — | 2 | — | 2 | — | 2 | — | — | — |
| SN-17 | 0 | 1 | 1 | 0 | 2 | 2 | 2 | 0 | 1 |
| SN-44 | 1 | 1 | 1 | 1 | 1 | 1 | — | — | 1 |
| 2417J | — | — | — | — | — | — | — | 2 | — |
| 2417Q | 0 | — | — | 0 | — | — | 0 | — | — |
| 333B | — | — | — | 0 | 2 | — | — | 1 | — |
| SRR8144644 | 0 | 0 | 2 | 0 | 2 | 0 | — | 1 | 0 |
| SRR8144646 | 0 | 0 | 0 | 0 | 0 | 0 | 0 | 0 | 0 |

Continued on next page

Table S30: (Continued)

| ID | 40808672 | 40808726 | 40809983 | 40814107 | 40814197 | 40815060 | 40821795 | 40821871 | 40826201 |
|---|---|---|---|---|---|---|---|---|---|
| SRR8144647 | 0 | 0 | 0 | 0 | 0 | 0 | 0 | 0 | 0 |
| SRR8144650 | 0 | 0 | 0 | 0 | 0 | 0 | 0 | 0 | 0 |
| ERR2270782 | 2 | 2 | 2 | 2 | 2 | 2 | 2 | 2 | 2 |
| ERR2270783 | 2 | 1 | — | 0 | 0 | 1 | 1 | 0 | 1 |
| ERR2270784 | 0 | 0 | 0 | — | — | 0 | — | 0 | — |
| ERR2270785 | 1 | 1 | 1 | 1 | 1 | 1 | 1 | 1 | 1 |

**Table S31. Per-individual frequency distribution for each Denisovan-specific missense mutation in the focal 72kb *MUC19* region among Mexicans in 1000 Genomes Project, 23 ancient Indigenous American individuals, and Indigenous American individuals in the Simons Genome Diversity Project.**

Summary of the per-individual frequency (i.e., 0, 0.5, or 1) distribution for each of the nine Denisovan-specific missense mutations in the focal 72kb *MUC19* region per group: 64 Mexican individuals in the 1000 Genomes project ($n = 128$ chromosomes), 23 ancient Indigenous American individuals ($n = 46$ chromosomes), and 22 Indigenous American individuals in the Simons Genome Diversity Project ($n = 44$ chromosomes). For each group, the total number of chromosomes that passed quality control, the mean (i.e., frequency), standard deviation ($\sigma$), standard error of the mean ($SEM$), and 95% confidence intervals ($\pm CI_{95\%}$) of the per-individual Denisovan-specific missense mutation frequency distribution are reported.

Table S31:

| Position | Group | Number of Chroms. | Denisovan-specific Mis. Mut. Freq. | Denisovan-specific Mis. Mut. ($\sigma$) | Denisovan-specific Mis. Mut. ($SEM$) | Denisovan-specific Mis. Mut. ($\pm CI_{95\%}$) |
|---|---|---|---|---|---|---|
| 40808672 | MXL | 128 | 0.305 | 0.301 | 0.038 | 0.076 |
| 40808672 | SDGP AMR | 44 | 0.364 | 0.308 | 0.067 | 0.14 |
| 40808672 | Ancient Americans | 36 | 0.361 | 0.402 | 0.097 | 0.205 |
| 40808726 | MXL | 128 | 0.305 | 0.301 | 0.038 | 0.076 |
| 40808726 | SDGP AMR | 44 | 0.364 | 0.308 | 0.067 | 0.14 |
| 40808726 | Ancient Americans | 40 | 0.375 | 0.415 | 0.095 | 0.199 |
| 40809983 | MXL | 128 | 0.305 | 0.301 | 0.038 | 0.076 |
| 40809983 | SDGP AMR | 44 | 0.364 | 0.308 | 0.067 | 0.14 |
| 40809983 | Ancient Americans | 32 | 0.375 | 0.415 | 0.107 | 0.228 |
| 40814107 | MXL | 128 | 0.305 | 0.301 | 0.038 | 0.076 |
| 40814107 | SDGP AMR | 44 | 0.364 | 0.308 | 0.067 | 0.14 |

Continued on next page

Table S31: (Continued)

| Position | Group | Number of Chroms. | Denisovan-specific Mis. Mut. Freq. | Denisovan-specific Mis. Mut. ($\sigma$) | Denisovan-specific Mis. Mut. ($SEM$) | Denisovan-specific Mis. Mut. ($\pm CI_{95\%}$) |
|---|---|---|---|---|---|---|
| 40814107 | Ancient Americans | 36 | 0.306 | 0.413 | 0.1 | 0.211 |
| 40814197 | MXL | 128 | 0.305 | 0.301 | 0.038 | 0.076 |
| 40814197 | SDGP AMR | 44 | 0.364 | 0.308 | 0.067 | 0.14 |
| 40814197 | Ancient Americans | 32 | 0.406 | 0.441 | 0.114 | 0.243 |
| 40815060 | MXL | 128 | 0.305 | 0.301 | 0.038 | 0.076 |
| 40815060 | SDGP AMR | 44 | 0.364 | 0.308 | 0.067 | 0.14 |
| 40815060 | Ancient Americans | 34 | 0.382 | 0.438 | 0.11 | 0.232 |
| 40821795 | MXL | 128 | 0.305 | 0.301 | 0.038 | 0.076 |
| 40821795 | SDGP AMR | 44 | 0.364 | 0.308 | 0.067 | 0.14 |
| 40821795 | Ancient Americans | 28 | 0.321 | 0.406 | 0.113 | 0.243 |
| 40821871 | MXL | 128 | 0.305 | 0.301 | 0.038 | 0.076 |
| 40821871 | SDGP AMR | 44 | 0.364 | 0.308 | 0.067 | 0.14 |
| 40821871 | Ancient Americans | 38 | 0.316 | 0.404 | 0.095 | 0.2 |
| 40826201 | MXL | 128 | 0.305 | 0.301 | 0.038 | 0.076 |
| 40826201 | SDGP AMR | 44 | 0.364 | 0.308 | 0.067 | 0.14 |
| 40826201 | Ancient Americans | 30 | 0.367 | 0.386 | 0.103 | 0.221 |

**Table S32. Comparison of the per-individual frequency distributions for each Denisovan-specific missense mutation in the focal 72kb *MUC19* region between Mexicans in 1000 Genomes Project, 23 ancient Indigenous American individuals, and Indigenous American individuals in the Simons Genome Diversity Project.**
Comparisons of the per-individual frequency distribution for each of the nine Denisovan-specific missense mutations in the focal 72kb *MUC19* region between groups: 64 Mexican individuals in the 1000 Genomes project ($n = 128$ chromosomes), 23 ancient Indigenous American individuals ($n = 46$ chromosomes), and 22 Indigenous American individuals in the Simons Genome Diversity Project ($n = 44$ chromosomes). For each comparison, the total number of chromosomes that passed quality control, the number of chromosomes harboring the respective Denisovan-specific missense mutation, and the results of the Proportions $Z$-Test and Fisher's Exact Test are reported. A $P$-*value* less than 0.05 is considered statistically significant.

Table S32:

| Position | Group 1 | Group 2 | Number of Chroms. (Group 1) | Denisovan-specific Mis. Mut. Count (Group 1) | Number of Chroms. (Group 2) | Denisovan-specific Mis. Mut. Count (Group 2) | $Z-Statistic$ | $P-value$ (Prop. $Z$-Test) | Odds Ratio | $P-value$ (Fisher's Exact Test) |
|---|---|---|---|---|---|---|---|---|---|---|
| 40808672 | MXL | Ancient Americans | 128 | 39 | 36 | 13 | -0.643 | 0.52 | 0.775 | 0.547 |
| 40808726 | MXL | Ancient Americans | 128 | 39 | 40 | 15 | -0.831 | 0.406 | 0.73 | 0.441 |
| 40809983 | MXL | Ancient Americans | 128 | 39 | 32 | 12 | -0.763 | 0.445 | 0.73 | 0.525 |
| 40814107 | MXL | Ancient Americans | 128 | 39 | 36 | 11 | -0.01 | 0.992 | 0.996 | 1 |

Continued on next page

Table S32: (Continued)

| Position | Group 1 | Group 2 | Number of Chroms. (Group 1) | Denisovan-specific Mis. Mut. Count (Group 1) | Number of Chroms. (Group 2) | Denisovan-specific Mis. Mut. Count (Group 2) | $Z-Statistic$ | $P-value$ (Prop. $Z$-Test) | Odds Ratio | $P-value$ (Fisher's Exact Test) |
|---|---|---|---|---|---|---|---|---|---|---|
| 40814197 | MXL | Ancient Americans | 128 | 39 | 32 | 13 | -1.097 | 0.273 | 0.64 | 0.296 |
| 40815060 | MXL | Ancient Americans | 128 | 39 | 34 | 13 | -0.862 | 0.389 | 0.708 | 0.413 |
| 40821795 | MXL | Ancient Americans | 128 | 39 | 28 | 9 | -0.174 | 0.862 | 0.925 | 0.826 |
| 40821871 | MXL | Ancient Americans | 128 | 39 | 38 | 12 | -0.13 | 0.896 | 0.949 | 1 |
| 40826201 | MXL | Ancient Americans | 128 | 39 | 30 | 11 | -0.657 | 0.511 | 0.757 | 0.519 |
| 40808672 | MXL | SDGP AMR | 128 | 39 | 44 | 16 | -0.723 | 0.47 | 0.767 | 0.461 |
| 40808726 | MXL | SDGP AMR | 128 | 39 | 44 | 16 | -0.723 | 0.47 | 0.767 | 0.461 |
| 40809983 | MXL | SDGP AMR | 128 | 39 | 44 | 16 | -0.723 | 0.47 | 0.767 | 0.461 |

Continued on next page

Table S32: (Continued)

| Position | Group 1 | Group 2 | Number of Chroms. (Group 1) | Denisovan-specific Mis. Mut. Count (Group 1) | Number of Chroms. (Group 2) | Denisovan-specific Mis. Mut. Count (Group 2) | $Z-Statistic$ | $P-value$ (Prop. $Z$-Test) | Odds Ratio | $P-value$ (Fisher's Exact Test) |
|---|---|---|---|---|---|---|---|---|---|---|
| 40814107 | MXL | SDGP AMR | 128 | 39 | 44 | 16 | -0.723 | 0.47 | 0.767 | 0.461 |
| 40814197 | MXL | SDGP AMR | 128 | 39 | 44 | 16 | -0.723 | 0.47 | 0.767 | 0.461 |
| 40815060 | MXL | SDGP AMR | 128 | 39 | 44 | 16 | -0.723 | 0.47 | 0.767 | 0.461 |
| 40821795 | MXL | SDGP AMR | 128 | 39 | 44 | 16 | -0.723 | 0.47 | 0.767 | 0.461 |
| 40821871 | MXL | SDGP AMR | 128 | 39 | 44 | 16 | -0.723 | 0.47 | 0.767 | 0.461 |
| 40826201 | MXL | SDGP AMR | 128 | 39 | 44 | 16 | -0.723 | 0.47 | 0.767 | 0.461 |
| 40808672 | SDGP AMR | Ancient Americans | 44 | 16 | 36 | 13 | 0.023 | 0.981 | 1 | 1 |
| 40808726 | SDGP AMR | Ancient Americans | 44 | 16 | 40 | 15 | -0.108 | 0.914 | 0.952 | 1 |
| 40809983 | SDGP AMR | Ancient Americans | 44 | 16 | 32 | 12 | -0.101 | 0.919 | 0.952 | 1 |

Continued on next page

Table S32: (Continued)

| Position | Group 1 | Group 2 | Number of Chroms. (Group 1) | Denisovan-specific Mis. Mut. Count (Group 1) | Number of Chroms. (Group 2) | Denisovan-specific Mis. Mut. Count (Group 2) | $Z-Statistic$ | $P-value$ (Prop. $Z$-Test) | Odds Ratio | $P-value$ (Fisher's Exact Test) |
|---|---|---|---|---|---|---|---|---|---|---|
| 40814107 | SDGP AMR | Ancient Ameri-cans | 44 | 16 | 36 | 11 | 0.547 | 0.585 | 1 | 0.64 |
| 40814197 | SDGP AMR | Ancient Ameri-cans | 44 | 16 | 32 | 13 | -0.378 | 0.706 | 0.835 | 0.812 |
| 40815060 | SDGP AMR | Ancient Ameri-cans | 44 | 16 | 34 | 13 | -0.17 | 0.865 | 0.923 | 1 |
| 40821795 | SDGP AMR | Ancient Ameri-cans | 44 | 16 | 28 | 9 | 0.367 | 0.714 | 1 | 0.802 |
| 40821871 | SDGP AMR | Ancient Ameri-cans | 44 | 16 | 38 | 12 | 0.456 | 0.649 | 1 | 0.816 |
| 40826201 | SDGP AMR | Ancient Ameri-cans | 44 | 16 | 30 | 11 | -0.027 | 0.979 | 0.987 | 1 |

**Table S33. Frequency of the Denisovan-specific missense mutations in the focal 72kb *MUC19* region among the African super population and African populations in the Simons Genome Diversity Project.**
Frequencies of the nine Denisovan-specific missense mutations found within the focal 72kb *MUC19* region for the entire African super population and stratified by African population in the Simons Genome Diversity Project.

Table S33:

| Group | 40808672 | 40808726 | 40809983 | 40814107 | 40814197 | 40815060 | 40821795 | 40821871 | 40826201 |
|---|---|---|---|---|---|---|---|---|---|
| AFR | 0.011 | 0.011 | 0.011 | 0.011 | 0.011 | 0.011 | 0.011 | 0.011 | 0.011 |
| Khomani San | 0.25 | 0.25 | 0.25 | 0.25 | 0.25 | 0.25 | 0.25 | 0.25 | 0.25 |
| Ju\|'hoan North | 0.0 | 0.0 | 0.0 | 0.0 | 0.0 | 0.0 | 0.0 | 0.0 | 0.0 |
| Dinka | 0.0 | 0.0 | 0.0 | 0.0 | 0.0 | 0.0 | 0.0 | 0.0 | 0.0 |
| Mandenka | 0.0 | 0.0 | 0.0 | 0.0 | 0.0 | 0.0 | 0.0 | 0.0 | 0.0 |
| Mbuti | 0.0 | 0.0 | 0.0 | 0.0 | 0.0 | 0.0 | 0.0 | 0.0 | 0.0 |
| Yoruba | 0.0 | 0.0 | 0.0 | 0.0 | 0.0 | 0.0 | 0.0 | 0.0 | 0.0 |
| Bantu Herero | 0.0 | 0.0 | 0.0 | 0.0 | 0.0 | 0.0 | 0.0 | 0.0 | 0.0 |
| Bantu Kenya | 0.0 | 0.0 | 0.0 | 0.0 | 0.0 | 0.0 | 0.0 | 0.0 | 0.0 |
| Bantu Tswana | 0.0 | 0.0 | 0.0 | 0.0 | 0.0 | 0.0 | 0.0 | 0.0 | 0.0 |
| Biaka | 0.0 | 0.0 | 0.0 | 0.0 | 0.0 | 0.0 | 0.0 | 0.0 | 0.0 |
| Esan | 0.0 | 0.0 | 0.0 | 0.0 | 0.0 | 0.0 | 0.0 | 0.0 | 0.0 |
| Gambian | 0.0 | 0.0 | 0.0 | 0.0 | 0.0 | 0.0 | 0.0 | 0.0 | 0.0 |
| Luhya | 0.0 | 0.0 | 0.0 | 0.0 | 0.0 | 0.0 | 0.0 | 0.0 | 0.0 |
| Luo | 0.0 | 0.0 | 0.0 | 0.0 | 0.0 | 0.0 | 0.0 | 0.0 | 0.0 |

Continued on next page

Table S33: (Continued)

| Group | 40808672 | 40808726 | 40809983 | 40814107 | 40814197 | 40815060 | 40821795 | 40821871 | 40826201 |
|---|---|---|---|---|---|---|---|---|---|
| Masai | 0.0 | 0.0 | 0.0 | 0.0 | 0.0 | 0.0 | 0.0 | 0.0 | 0.0 |
| Mende | 0.0 | 0.0 | 0.0 | 0.0 | 0.0 | 0.0 | 0.0 | 0.0 | 0.0 |
| Mozabite | 0.0 | 0.0 | 0.0 | 0.0 | 0.0 | 0.0 | 0.0 | 0.0 | 0.0 |
| Saharawi | 0.0 | 0.0 | 0.0 | 0.0 | 0.0 | 0.0 | 0.0 | 0.0 | 0.0 |
| Somali | 0.0 | 0.0 | 0.0 | 0.0 | 0.0 | 0.0 | 0.0 | 0.0 | 0.0 |

**Table S34. Frequency of introgressed haplotypes at the focal 72kb *MUC19* region among 1000 Genomes Project super populations.**

The frequency of introgressed haplotypes—i.e., the number of introgressed haplotypes normalized by the total number of chromosomes—stratified by super population: Admixed Americans (AMR), South Asians (SAS), East Asians (EAS), Europeans (EUR), and Africans (AFR). Introgressed haplotypes at the focal 72kb *MUC19* region are significantly enriched in AMR individuals (Fisher's Exact Test, Odds Ratio: 2.848, *P-value*: $1.320e^{-14}$), with AMR populations exhibiting a higher proportion of introgressed tracts compared to non-AMR populations, excluding AFR (Proportions $Z$-Test, $Z$-statistic: 8.328, *P-value*: $4.116e^{-17}$).

Table S34:

| Super Population | Total Number of Chromosomes | Number of Introgressed Haps. | Introgressed Hap. Frequency |
|---|---|---|---|
| AMR | 694 | 107 | 0.154 |
| SAS | 978 | 96 | 0.098 |
| EAS | 1008 | 61 | 0.061 |
| EUR | 1006 | 23 | 0.023 |
| AFR | 1008 | 0 | 0 |

**Table S35. Frequency of introgressed haplotypes at the focal 72kb *MUC19* region among 1000 Genomes Project populations.**
The frequency of introgressed haplotypes—i.e., the number of introgressed haplotypes normalized by the total number of chromosomes—for each population stratified by super population: Admixed Americans (AMR), South Asians (SAS), East Asians (EAS), Europeans (EUR), and Africans (AFR).

Table S35:

| Super Population | Population | Total Number of Chromosomes | Number of Introgressed Haps. | Introgressed Hap. Frequency |
|---|---|---|---|---|
| AMR | MXL | 128 | 39 | 0.305 |
| AMR | PEL | 170 | 37 | 0.218 |
| AMR | CLM | 188 | 13 | 0.069 |
| AMR | PUR | 208 | 18 | 0.087 |
| SAS | BEB | 172 | 24 | 0.14 |
| SAS | STU | 204 | 23 | 0.113 |
| SAS | ITU | 204 | 21 | 0.103 |
| SAS | PJL | 192 | 10 | 0.052 |
| SAS | GIH | 206 | 18 | 0.087 |
| EAS | CHB | 206 | 7 | 0.034 |
| EAS | KHV | 198 | 17 | 0.086 |
| EAS | CHS | 210 | 8 | 0.038 |
| EAS | JPT | 208 | 6 | 0.029 |
| EAS | CDX | 186 | 23 | 0.124 |
| EUR | TSI | 214 | 11 | 0.051 |
| EUR | CEU | 198 | 3 | 0.015 |
| EUR | IBS | 214 | 7 | 0.033 |

Continued on next page

Table S35: (Continued)

| Super Population | Population | Total Number of Chromosomes | Number of Introgressed Haps. | Introgressed Hap. Frequency |
|---|---|---|---|---|
| EUR | GBR | 182 | 1 | 0.005 |
| EUR | FIN | 198 | 1 | 0.005 |
| AFR | LWK | 198 | 0 | 0 |
| AFR | GWD | 226 | 0 | 0 |
| AFR | MSL | 170 | 0 | 0 |
| AFR | ESN | 198 | 0 | 0 |
| AFR | YRI | 216 | 0 | 0 |

**Table S36. Sequence divergence from the Denisovan for the Altai Neanderthal and all African Individuals in the 1000 Genomes Project (1KG) at the focal 72kb *MUC19* region.**

The observed sequence divergence from the Densisovan for the Altai Neanderthal (i.e., the number of pairwise differences between the Altai Neanderthal's and Denisovan's unphased genotypes normalized by the effective sequence length) and all Africans in the 1KG (i.e., the average number of pairwise differences between all African haplotypes in the 1KG and the Denisovan's unphased genotypes normalized by the effective sequence length). For each comparison to the Denisovan, the mean ($\mu$), standard deviation ($\sigma$), standard error of the mean ($SEM$), and 95% confidence intervals ($\pm CI_{95\%}$) of the sequence divergence genomic background distribution used to compute the *P-value* are also reported. The *P-value* represents the proportion of non-overlapping 72kb windows of comparable effective sequence length where the sequence divergence is greater than or equal to that observed at the focal 72kb region. A *P-value* less than 0.05 is considered statistically significant. Note that at the focal 72kb region, the effective sequence length with respect to the Denisovan is 46136bp and 48435bp for the Altai Neanderthal and African comparison, respectively.

Table S36:

| Comparison | Focal 72kb Region (Pairwise Diffs.) | Focal 72kb Region (Seq. Div.) | 72kb Nonoverlapping Windows ($\mu$) | 72kb Nonoverlapping Windows ($\sigma$) | 72kb Nonoverlapping Windows ($SEM$) | 72kb Nonoverlapping Windows ($\pm CI_{95\%}$) | $P-value$ |
|---|---|---|---|---|---|---|---|
| Altai Nean. | 174.5 | 0.003782 | 0.001 | $6.013e^{-4}$ | $3.501e^{-6}$ | $6.862e^{-6}$ | 0.002 |
| AFR | 194.885 | 0.004024 | 0.001 | $5.529e^{-4}$ | $3.168e^{-6}$ | $6.210e^{-6}$ | 0.002 |

**Table S37. Sequence divergence between the *Denisovan-like* haplotype in MXL and unphased archaic individuals at the focal 72kb *MUC19* region.**
The observed sequence divergence—i.e., the number of pairwise differences between a modern human haplotype and an archaic genotype normalized by the effective sequence length—between the *Denisovan-like* haplotype in the focal MXL individual (i.e., NA19664), who harbors two *Denisovan-like* haplotypes and has no heterozygous sites at the focal 72kb region, and the four unphased high-coverage archaic individuals. For each archaic individual, the mean ($\mu$), standard deviation ($\sigma$), standard error of the mean ($SEM$), and 95% confidence intervals ($\pm CI_{95\%}$) of the sequence divergence genomic background distribution used to compute the *P-value* are also reported. The *P-value* represents the proportion of non-overlapping 72kb windows of comparable effective sequence length where the sequence divergence is less than or equal to that observed at the focal 72kb region. After correcting for two multiple comparisons—i.e., one per haplotype—a *P-value* less than 0.025 is considered statistically significant. Note that at the focal 72kb region, the effective sequence length with respect to the NA19664 individual is 48435bp, 48654bp, 48121bp, and 48447bp for the Denisovan, Altai Neanderthal, Chagyrskaya Neanderthal, and Vindija Neanderthal, respectively.

Table S37:

| Archaic | Focal 72kb Region (Pairwise Diffs. Hap. 1) | Focal 72kb Region (Pairwise Diffs. Hap. 2) | Focal 72kb Region (Seq. Div. Hap. 1) | Focal 72kb Region (Seq. Div. Hap. 2) | 72kb Non-overlapping Windows ($\mu$) | 72kb Non-overlapping Windows ($\sigma$) | 72kb Non-overlapping Windows ($SEM$) | 72kb Non-overlapping Windows ($\pm CI_{95\%}$) | $P-value$ (Hap. 1) | $P-value$ (Hap. 2) |
|---|---|---|---|---|---|---|---|---|---|---|
| Denisovan | 47 | 47 | 0.00097 | 0.00097 | 0.001 | $6.132e^{-4}$ | $2.484e^{-6}$ | $4.870e^{-6}$ | 0.237 | 0.237 |
| Altai Nean. | 177.5 | 177.5 | 0.003648 | 0.003648 | 0.001 | $6.028e^{-4}$ | $2.442e^{-6}$ | $4.787e^{-6}$ | 0.995 | 0.995 |
| Chagyrskaya Nean. | 87.5 | 87.5 | 0.001818 | 0.001818 | 0.001 | $6.074e^{-4}$ | $2.465e^{-6}$ | $4.832e^{-6}$ | 0.811 | 0.811 |
| Vindija Nean. | 88 | 88 | 0.001816 | 0.001816 | 0.001 | $6.041e^{-4}$ | $2.447e^{-6}$ | $4.796e^{-6}$ | 0.806 | 0.806 |

**Table S38. *D+* tests for introgression at the focal 72kb *MUC19*.**
*D+* results for the focal 72kb *MUC19* region using the Yoruba in Ibadan, Nigeria population (YRI) as *P1*, the introgressed haplotype in the focal MXL individual (i.e., NA19664) as *P2*, the four high-coverage archaics as *P3*, and the EPO ancestral sequence to polarize ancestral states. For each archaic(*P3*), the mean ($\mu$), standard deviation ($\sigma$), standard error of the mean ($SEM$), and 95% confidence intervals ($\pm CI_{95\%}$) of the *D+* genomic background distribution used to compute the *P-value* are also reported. *P-values* were computed by building a *Z*-distribution of *D+* values from the non-overlapping 72kb windows of comparable effective sequence length. A *P-value* less than 0.05 is considered statistically significant and suggests that introgressed haplotype in MXL (*P2*) shares more derived and ancestral alleles with the given archaic individual (*P3*) than expected under a model of no gene flow.

Table S38:

| $P1$ | $P2$ | $P3$ | Focal 72kb Region ($D+$) | 72kb Non-overlapping Windows ($\mu$) | 72kb Non-overlapping Windows ($\sigma$) | 72kb Non-overlapping Windows ($SEM$) | 72kb Non-overlapping Windows ($\pm CI_{95\%}$) | $P-value$ |
|---|---|---|---|---|---|---|---|---|
| YRI | NA19664 | Denisovan | 0.743 | 0.007 | 0.177 | 0.001 | 0.291 | $1.386e^{-5}$ |
| YRI | NA19664 | Altai Nean. | -0.622 | 0.02 | 0.191 | 0.001 | 0.314 | 0.999 |
| YRI | NA19664 | Chagyrskaya Nean. | 0.175 | 0.02 | 0.193 | 0.001 | 0.317 | 0.183 |
| YRI | NA19664 | Vindija Nean. | 0.182 | 0.021 | 0.194 | 0.001 | 0.318 | 0.174 |

**Table S39. Sequence divergence between the longest introgressed tract in MXL and unphased archaic individuals at the focal 742kb *MUC19* region.**
The observed sequence divergence—i.e., the number of pairwise differences between a modern human haplotype and an archaic genotype normalized by the effective sequence length—between the MXL individual (i.e., NA19725) harboring the longest introgressed tract in MXL (i.e., 742kb) and the four unphased high-coverage archaic individuals. For each archaic individual, the mean ($\mu$), standard deviation ($\sigma$), standard error of the mean ($SEM$), and 95% confidence intervals ($\pm CI_{95\%}$) of the sequence divergence genomic background distribution used to compute the *P-value* are also reported. The *P-value* represents the proportion of non-overlapping 742kb windows of comparable effective sequence length where the sequence divergence is less than or equal to that observed at the focal 742kb region. After correcting for two multiple comparisons—i.e., one per haplotype—a *P-value* less than 0.025 is considered statistically significant. Note that the 742kb longest introgressed tract in MXL corresponds to NA19725's second haplotype (i.e., Hap. 2). Note that at the focal 742kb region, the effective sequence length with respect to the NA19725 individual is 495788bp, 497642bp, 494606bp, and 499368bp for the Denisovan, Altai Neanderthal, Chagyrskaya Neanderthal, and Vindija Neanderthal, respectively.

Table S39:

| Archaic | Focal 742kb Region (Pairwise Diffs. Hap. 1) | Focal 742kb Region (Pairwise Diffs. Hap. 2) | Focal 742kb Region (Seq. Div. Hap. 1) | Focal 742kb Region (Seq. Div. Hap. 2) | 742kb Non-overlapping Windows ($\mu$) | 742kb Non-overlapping Windows ($\sigma$) | 742kb Non-overlapping Windows ($SEM$) | 742kb Non-overlapping Windows ($\pm CI_{95\%}$) | $P-value$ (Hap. 1) | $P-value$ (Hap. 2) |
|---|---|---|---|---|---|---|---|---|---|---|
| Denisovan | 792.5 | 399.5 | 0.001598 | 0.000806 | 0.001 | $3.460e^{-4}$ | $4.335e^{-6}$ | $8.498e^{-6}$ | 0.755 | 0.019 |
| Altai Nean. | 580.5 | 450.5 | 0.001167 | 0.000905 | 0.001 | $3.383e^{-4}$ | $4.233e^{-6}$ | $8.297e^{-6}$ | 0.294 | 0.065 |
| Chagyrskaya Nean. | 662 | 327 | 0.001338 | 0.000661 | 0.001 | $3.403e^{-4}$ | $4.264e^{-6}$ | $8.359e^{-6}$ | 0.512 | 0.006 |
| Vindija Nean. | 658.5 | 327.5 | 0.001319 | 0.000656 | 0.001 | $3.397e^{-4}$ | $4.248e^{-6}$ | $8.328e^{-6}$ | 0.47 | 0.007 |

**Table S40. Frequency of introgressed haplotypes at the focal 742kb *MUC19* region among 1000 Genomes Project super populations.**

The frequency of introgressed haplotypes—i.e., the number of introgressed haplotypes normalized by the total number of chromosomes—stratified by super population: Admixed Americans (AMR), South Asians (SAS), East Asians (EAS), Europeans (EUR), and Africans (AFR). Introgressed haplotypes at the focal 742kb *MUC19* region are significantly enriched in AMR individuals (Fisher's Exact Test, Odds Ratio: 10.033, *P-value*: $1.601e^{-37}$), with AMR populations exhibiting a higher proportion of introgressed tracts compared to non-AMR populations, excluding AFR (Proportions $Z$-Test, $Z$-statistic: 14.909, *P-value*: $1.441e^{-50}$).

Table S40:

| Super Population | Total Number of Chromosomes | Number of Introgressed Haps. | Introgressed Hap. Frequency |
|---|---|---|---|
| AMR | 694 | 94 | 0.135 |
| SAS | 978 | 21 | 0.021 |
| EAS | 1008 | 23 | 0.023 |
| EUR | 1006 | 2 | 0.002 |
| AFR | 1008 | 0 | 0 |

**Table S41. Frequency of introgressed haplotypes at the focal 742kb *MUC19* region among 1000 Genomes Project populations.**
The frequency of introgressed haplotypes—i.e., the number of introgressed haplotypes normalized by the total number of chromosomes—for each population stratified by super population: Admixed Americans (AMR), South Asians (SAS), East Asians (EAS), Europeans (EUR), and Africans (AFR).

Table S41:

| Super Population | Population | Total Number of Chromosomes | Number of Introgressed Haps. | Introgressed Hap. Frequency |
|---|---|---|---|---|
| AMR | MXL | 128 | 40 | 0.312 |
| AMR | PEL | 170 | 30 | 0.176 |
| AMR | CLM | 188 | 10 | 0.053 |
| AMR | PUR | 208 | 14 | 0.067 |
| SAS | BEB | 172 | 6 | 0.035 |
| SAS | STU | 204 | 5 | 0.025 |
| SAS | ITU | 204 | 3 | 0.015 |
| SAS | PJL | 192 | 1 | 0.005 |
| SAS | GIH | 206 | 6 | 0.029 |
| EAS | CHB | 206 | 3 | 0.015 |
| EAS | KHV | 198 | 6 | 0.03 |
| EAS | CHS | 210 | 2 | 0.01 |
| EAS | JPT | 208 | 4 | 0.019 |
| EAS | CDX | 186 | 8 | 0.043 |
| EUR | TSI | 214 | 1 | 0.005 |
| EUR | CEU | 198 | 1 | 0.005 |
| EUR | IBS | 214 | 0 | 0 |

Continued on next page

Table S41: (Continued)

| Super Population | Population | Total Number of Chromosomes | Number of Introgressed Haps. | Introgressed Hap. Frequency |
|---|---|---|---|---|
| EUR | GBR | 182 | 0 | 0 |
| EUR | FIN | 198 | 0 | 0 |
| AFR | LWK | 198 | 0 | 0 |
| AFR | GWD | 226 | 0 | 0 |
| AFR | MSL | 170 | 0 | 0 |
| AFR | ESN | 198 | 0 | 0 |
| AFR | YRI | 216 | 0 | 0 |

**Table S42. *D+* tests for introgression at the focal 742kb *MUC19*.**
*D+* results for the focal 742kb *MUC19* region using the Yoruba in Ibadan, Nigeria population (YRI) as *P1*, the introgressed haplotype in the focal MXL individual (i.e., NA19664) as *P2*, the four high-coverage archaics as *P3*, and the EPO ancestral sequence to polarize ancestral states. For each archaic(*P3*), the mean ($\mu$), standard deviation ($\sigma$), standard error of the mean ($SEM$), and 95% confidence intervals ($\pm CI_{95\%}$) of the *D+* genomic background distribution used to compute the *P-value* are also reported. *P-values* were computed by building a *Z*-distribution of *D+* values from the non-overlapping 742kb windows of comparable effective sequence length. A *P-value* less than 0.05 is considered statistically significant and suggests that introgressed haplotype in MXL (*P2*) shares more derived and ancestral alleles with the given archaic individual (*P3*) than expected under a model of no gene flow.

Table S42:

| $P1$ | $P2$ | $P3$ | Focal 742kb Region ($D+$) | 742kb Non-overlapping Windows ($\mu$) | 742kb Non-overlapping Windows ($\sigma$) | 742kb Non-overlapping Windows ($SEM$) | 742kb Non-overlapping Windows ($\pm CI_{95\%}$) | $P-value$ |
|---|---|---|---|---|---|---|---|---|
| YRI | NA19664 | Denisovan | 0.377 | 0.002 | 0.072 | 0.001 | 0.119 | $9.889e^{-8}$ |
| YRI | NA19664 | Altai Nean. | 0.091 | 0.017 | 0.086 | 0.002 | 0.141 | 0.144 |
| YRI | NA19664 | Chagyrskaya Nean. | 0.381 | 0.018 | 0.088 | 0.002 | 0.145 | $7.375e^{-6}$ |
| YRI | NA19664 | Vindija Nean. | 0.383 | 0.019 | 0.088 | 0.002 | 0.145 | $7.505e^{-6}$ |

**Table S43. The number of heterozygous sites in the focal 72kb *MUC19* region among archaic individuals.** The number of heterozygous observed in the focal 72kb *MUC19* region for each of the high-coverage archaic individuals. For each archaic individual, the mean ($\mu$), standard deviation ($\sigma$), standard error of the mean (*SEM*), and 95% confidence intervals ($\pm CI_{95\%}$) of the heterozygous sites genomic background distribution used to compute the *P-value* are also reported. The *P-value* represents the proportion of non-overlapping 72kb windows of comparable effective sequence length where the number of heterozygous sites is greater than or equal to that observed at the focal 72kb region. A *P-value* less than 0.05 is considered statistically significant.

Table S43:

| Archaic | Focal 72kb Region (Het. Sites) | Non-overlapping Windows ($\mu$) | Non-overlapping Windows ($\sigma$) | Non-overlapping Windows ($SEM$) | Non-overlapping Windows ($\pm CI_{95\%}$) | $P-value$ |
|---|---|---|---|---|---|---|
| Denisovan | 6 | 8.747 | 14.169 | 0.081 | 0.159 | 0.455 |
| Altai Nean. | 1 | 7.862 | 15.261 | 0.087 | 0.171 | 0.679 |
| Chagyrskaya Nean. | 168 | 6.534 | 13.242 | 0.076 | 0.149 | $2.307e^{-4}$ |
| Vindija Nean. | 171 | 7.837 | 13.598 | 0.078 | 0.153 | $3.282e^{-4}$ |

**Table S44. The average number of heterozygous sites in the focal 72kb *MUC19* region among African individuals and individuals carrying a *Denisovan-like* haplotype in the 1000 Genomes Project.**
The average number of heterozygous observed in the focal 72kb *MUC19* region for each focal group in the 1000 Genomes Project: all African individuals ($n = 504$), individuals harboring one *Denisovan-like* haplotype ($n = 255$), and individuals harboring two *Denisovan-like* haplotypes ($n = 16$). For each group, the mean ($\mu$), standard deviation ($\sigma$), standard error of the mean ($SEM$), and 95% confidence intervals ($\pm CI_{95\%}$) of the average number of heterozygous sites genomic background distribution used to compute the *P-value* are also reported. For the African individuals and individuals harboring one *Denisovan-like* haplotype groups, the *P-value* represents the proportion of non-overlapping 72kb windows of comparable effective sequence length where the average number of heterozygous sites is greater than or equal to that observed at the focal 72kb region. For the individuals harboring two *Denisovan-like* haplotypes group, the *P-value* represents the proportion of non-overlapping 72kb windows of comparable effective sequence length where the average number of heterozygous sites is less than or equal to that observed at the focal 72kb region. A *P-value* less than 0.05 is considered statistically significant.

Table S44:

| Group | Focal 72kb Region (Average Number of Het. Sites) | Non-overlapping Windows ($\mu$) | Non-overlapping Windows ($\sigma$) | Non-overlapping Windows ($SEM$) | Non-overlapping Windows ($\pm CI_{95\%}$) | $P-value$ |
|---|---|---|---|---|---|---|
| African Inds. | 75.315 | 73.319 | 25.708 | 0.144 | 0.283 | 0.424 |
| Inds. with One $Denisovan-like$ Hap. | 287.341 | 54.639 | 23.128 | 0.13 | 0.255 | $3.157e^{-5}$ |
| Inds. with Two $Denisovan-like$ Haps. | 4.188 | 52.579 | 24.891 | 0.14 | 0.274 | $6.945e^{-4}$ |

**Table S45. Genome-wide heterozygosity among archaic individuals and African individuals in the 1000 Genomes Project.**
The genome-wide heterozygosity (i.e., the number of heterozygous sites normalized by the effective sequence length) for each of the high coverage archaic individuals and the mean genome-wide heterozygosity among all African individuals in the 1000 Genomes Project.

Table S45:

| Group | Genome-Wide Heterozygosity |
|---|---|
| AFR | 0.000985 |
| Denisovan | 0.000191 |
| Altai Nean. | 0.000171 |
| Chagyrskaya Nean. | 0.000144 |
| Vindija Nean. | 0.00017 |

**Table S46. *D+* tests for Denisovan introgression at the focal 72kb *MUC19* region among the late Neanderthal individuals.**
*D+* results for the focal 72kb *MUC19* region using Altai Neanderthal as *P1*, each of the late Neanderthals as *P2*, the Denisovan as *P3*, and the EPO ancestral sequence to polarize ancestral states. For each late Neanderthal (*P2*), the mean ($\mu$), standard deviation ($\sigma$), standard error of the mean ($SEM$), and 95% confidence intervals ($\pm CI_{95\%}$) of the *D+* genomic background distribution used to compute the *P-value* are also reported. *P-values* were computed by building a *Z*-distribution of *D+* values from the non-overlapping 72kb windows of comparable effective sequence length. A *P-value* less than 0.05 is considered statistically significant and suggests that the late Neanderthal (*P2*) shares more derived and ancestral alleles with the Denisovan (*P3*) than expected under a model of no gene flow.

Table S46:

| $P1$ | $P2$ | $P3$ | Focal 72kb Region ($D+$) | 72kb Non-overlapping Windows ($\mu$) | 72kb Non-overlapping Windows ($\sigma$) | 72kb Non-overlapping Windows ($SEM$) | 72kb Non-overlapping Windows ($\pm CI_{95\%}$) | $P - value$ |
|---|---|---|---|---|---|---|---|---|
| Altai Nean. | Chagyrskaya Nean. | Denisovan | 0.783 | -0.113 | 0.414 | 0.002 | 0.682 | 0.029 |
| Altai Nean. | Vindija Nean. | Denisovan | 0.819 | -0.169 | 0.391 | 0.002 | 0.644 | 0.018 |

**Table S47. The number of pairwise differences at the focal 72kb *MUC19* region among the Denisovan, Altai Neanderthal, and the phased late Neanderthal individuals.**
The number of pairwise differences observed at the focal 72kb *MUC19* region between all pairwise possibilities of the unphased Denisovan genotypes, unphased Altai Neanderthal genotypes, phased Chagyrskaya Neanderthal haplotypes, and phased Vindija Neanderthal haplotypes.

Table S47:

| Archaic 1 | Archaic 2 | Focal 72kb Region (Pairwise Diffs.) |
|---|---|---|
| Denisovan | Altai Nean. | 173.5 |
| Denisovan | Chagyrskaya Nean. Hap. 1 | 164 |
| Denisovan | Chagyrskaya Nean. Hap. 2 | 43 |
| Denisovan | Vindija Nean. Hap. 1 | 170 |
| Denisovan | Vindija Nean. Hap. 2 | 41 |
| Altai Nean. | Chagyrskaya Nean. Hap. 1 | 3.5 |
| Altai Nean. | Chagyrskaya Nean. Hap. 2 | 159.5 |
| Altai Nean. | Vindija Nean. Hap. 1 | 4 |
| Altai Nean. | Vindija Nean. Hap. 2 | 161 |
| Chagyrskaya Nean. Hap. 1 | Chagyrskaya Nean. Hap. 2 | 165 |
| Chagyrskaya Nean. Hap. 1 | Vindija Nean. Hap. 1 | 1 |
| Chagyrskaya Nean. Hap. 1 | Vindija Nean. Hap. 2 | 154 |
| Chagyrskaya Nean. Hap. 2 | Vindija Nean. Hap. 1 | 153 |
| Chagyrskaya Nean. Hap. 2 | Vindija Nean. Hap. 2 | 0 |
| Vindija Nean. Hap. 1 | Vindija Nean. Hap. 2 | 167 |

**Table S48. Sequence divergence between the *Denisovan-like* haplotype in MXL and phased late Neanderthal haplotypes at the focal 72kb *MUC19* region.**

The observed sequence divergence—i.e., the number of pairwise differences between a modern human haplotype and a late Neanderthal haplotype normalized by the effective sequence length—between the *Denisovan-like* haplotype in the focal MXL individual (i.e., NA19664), who harbors two *Denisovan-like* haplotypes and has no heterozygous sites at the focal 72kb region, and the phased haplotypes for both of the late Neanderthals. To assess the significance, we built a distribution of sequence divergence between each of the modern human's two chromosomes and the late Neanderthal's pseudo-haplotype, which was generated by randomly sampling one allele at each position from the unphased late Neanderthal's genotypes. For each late Neanderthal, the mean ($\mu$), standard deviation ($\sigma$), standard error of the mean ($SEM$), and 95% confidence intervals ($\pm CI_{95\%}$) of the pseudo-haplotype divergence genomic background distribution used to compute the *P-value* are also reported. The *P-value* represents the proportion of non-overlapping 72kb windows of comparable effective sequence length where the sequence divergence is less than or equal to that observed at the focal 72kb region. After correcting for four multiple comparisons—i.e., two per each modern human haplotype—a *P-value* less than 0.0125 is considered statistically significant. Note that the *Denisovan-like* in each of the late Neanderthal's corresponds to their second haplotype (i.e., Chagyrskaya Nean. Hap. 2 and Vindija Nean. Hap. 2). Also, note that at the phased 72kb region, the effective sequence length with respect to the NA19664 individual is 48119bp and 48444bp for the Chagyrskaya and Vindija Neanderthal, respectively.

Table S48:

| Archaic Hap. | Focal 72kb Region (Pairwise Diffs. Hap. 1) | Focal 72kb Region (Pairwise Diffs. Hap. 2) | Focal 72kb Region (Seq. Div. Hap. 1) | Focal 72kb Region (Seq. Div. Hap. 2) | 72kb Non-overlapping Windows ($\mu$) | 72kb Non-overlapping Windows ($\sigma$) | 72kb Non-overlapping Windows ($SEM$) | 72kb Nonoverlapping Windows ($\pm CI_{95\%}$) | $P-value$ (Hap. 1) | $P-value$ (Hap. 2) |
|---|---|---|---|---|---|---|---|---|---|---|
| Chagyrskaya Nean. Hap. 1 | 168 | 168 | 0.003491 | 0.003491 | 0.001 | $6.078e^{-4}$ | $2.467e^{-6}$ | $4.836e^{-6}$ | 0.993 | 0.993 |
| Chagyrskaya Nean. Hap. 2 | 5 | 5 | 0.000104 | 0.000104 | 0.001 | $6.078e^{-4}$ | $2.467e^{-6}$ | $4.836e^{-6}$ | 0.003 | 0.003 |

Continued on next page

Table S48: (Continued)

| Archaic Hap. | Focal 72kb Region (Pairwise Diffs. Hap. 1) | Focal 72kb Region (Pairwise Diffs. Hap. 2) | Focal 72kb Region (Seq. Div. Hap. 1) | Focal 72kb Region (Seq. Div. Hap. 2) | 72kb Non-overlapping Windows ($\mu$) | 72kb Non-overlapping Windows ($\sigma$) | 72kb Non-overlapping Windows ($SEM$) | 72kb Nonoverlapping Windows ($\pm CI_{95\%}$) | $P-value$ (Hap. 1) | $P-value$ (Hap. 2) |
|---|---|---|---|---|---|---|---|---|---|---|
| Vindija Nean. Hap. 1 | 169 | 169 | 0.003489 | 0.003489 | 0.001 | $6.046e^{-4}$ | $2.449e^{-6}$ | $4.801e^{-6}$ | 0.993 | 0.993 |
| Vindija Nean. Hap. 2 | 4 | 4 | 0.000083 | 0.000083 | 0.001 | $6.046e^{-4}$ | $2.449e^{-6}$ | $4.801e^{-6}$ | 0.002 | 0.002 |

**Table S49. Pseudo-Ancestry Painting scores for the focal 72kb *MUC19* region.**
The Pseudo-Ancestry Painting (*PAP*) scores observed in the focal 72kb *MUC19* region. The *PAP* score quantifies the number of heterozygous sites in the target individual (i.e., *Target*) that can be explained by the two source individuals (i.e., $Source^1$ and $Source^2$) being fixed different allelic states and then normalized by the total number of heterozygous sites in the target individual. *PAP* scores were first computed for the late Neanderthals as target individuals using the Denisovan and Altai Neanderthal as sources as well as a pairing of an MXL individual (i.e., NA19664) who is homozygous for the *Denisovan-like* haplotype at the 72kb region and a YRI individual (i.e., NA19190) who is homozygous for the *Human-like* haplotype. Additionally, as a negative control, we computed additional *PAP* configurations where the Denisovan and Altai Neanderthal are the target individuals and the focal MXL (i.e., NA19664) and YRI (i.e., NA19190) individuals are sources. For each *PAP* configuration, the mean ($\mu$), standard deviation ($\sigma$), standard error of the mean (*SEM*), and 95% confidence intervals ($\pm CI_{95\%}$) of the *PAP* score genomic background distribution used to compute the *P-value* are also reported. The *P-value* represents the proportion of non-overlapping 72kb windows of comparable effective sequence length where the *PAP* score is greater than or equal to that observed at the focal 72kb region. A *P-value* less than 0.05 is considered statistically significant.

Table S49:

| $Source^1$ | *Target* | $Source^2$ | Focal 72kb Region (*PAP* Score) | 72kb Non-overlapping Windows ($\mu$) | 72kb Non-overlapping Windows ($\sigma$) | 72kb Non-overlapping Windows (*SEM*) | 72kb Non-overlapping Windows ($\pm CI_{95\%}$) | $P-value$ |
|---|---|---|---|---|---|---|---|---|
| Denisovan | Chagyrskaya Nean. | Altai Nean. | 0.792 | 0.105 | 0.173 | 0.001 | 0.002 | 0.005 |
| Denisovan | Vindija Nean. | Altai Nean. | 0.801 | 0.085 | 0.154 | 0.001 | 0.002 | 0.003 |
| MXL (NA19664) | Chagyrskaya Nean. | YRI (NA19190) | 0.94 | 0.008 | 0.046 | $3.348e^{-4}$ | $6.562e^{-4}$ | $3.683e^{-4}$ |
| MXL (NA19664) | Vindija Nean. | YRI (NA19190) | 0.929 | 0.006 | 0.038 | $2.511e^{-4}$ | $4.922e^{-4}$ | $8.679e^{-5}$ |
| MXL (NA19664) | Denisovan | YRI (NA19190) | 0 | 0.005 | 0.032 | $1.998e^{-4}$ | $3.916e^{-4}$ | $> 0.999967$ |

Continued on next page

Table S49: (Continued)

| $Source^1$ | $Target$ | $Source^2$ | Focal 72kb Region ($PAP$ Score) | 72kb Non-overlapping Windows ($\mu$) | 72kb Non-overlapping Windows ($\sigma$) | 72kb Non-overlapping Windows ($SEM$) | 72kb Non-overlapping Windows ($\pm CI_{95\%}$) | $P-value$ |
|---|---|---|---|---|---|---|---|---|
| MXL (NA19664) | Altai Nean. | YRI (NA19190) | 0 | 0.008 | 0.044 | $3.111e^{-4}$ | $6.097e^{-4}$ | $> 0.999967$ |

**Table S50. $PBS_{MXL:CHB:CEU}$ values for the focal 742kb and 72kb *MUC19* using SPrime sites.**
The $PBS_{MXL:CHB:CEU}$ values observed in the focal 742kb and 72kb region for putatively introgressed sites in MXL that is a match with either the Altai Denisovan or the Altai Neanderthal as defined by the SPrime introgression map. For each focal region, the mean ($\mu$), standard deviation ($\sigma$), standard error of the mean ($SEM$), and 95% confidence intervals ($\pm CI_{95\%}$) of the $PBS_{MXL:CHB:CEU}$ genomic background distribution used to compute the *P-value* are also reported. The *P-value* represents the proportion of non-overlapping windows of comparable effective sequence length where the $PBS_{MXL:CHB:CEU}$ value, when only considering putatively introgressed SPrime sites, is greater than or equal to that observed at the focal respective focal region. A *P-value* less than 0.05 is considered statistically significant.

Table S50:

| Focal Region | $PBS_{MXL:CHB:CE}$ | Non-overlapping Windows ($\mu$) | Non-overlapping Windows ($\sigma$) | Non-overlapping Windows ($SEM$) | Non-overlapping Windows ($\pm CI_{95\%}$) | $P-value$ |
|---|---|---|---|---|---|---|
| 742kb | 0.291 | 0.01 | 0.027 | $7.689e^{-4}$ | 0.002 | $7.962e^{-4}$ |
| 72kb | 0.28 | 0.01 | 0.03 | $4.158e^{-4}$ | $8.151e^{-4}$ | 0.002 |

**Table S51.** $\boldsymbol{PBS_{AMR:EAS/SAS:EUR}}$ **values for the focal 742kb** $\boldsymbol{MUC19}$ **region.**
The $PBS_{A:B:C}$ values observed in the focal 742kb region for all unique combinations of: AMR populations as the target population ($A$), EAS and SAS populations as one of the control populations ($B$), and EUR populations as the second control population ($C$). For each unique $PBS_{A:B:C}$ configuration, the mean ($\mu$), standard deviation ($\sigma$), standard error of the mean ($SEM$), and 95% confidence intervals ($\pm CI_{95\%}$) of the $PBS_{A:B:C}$ genomic background distribution used to compute the *P-value* are also reported. The *P-value* represents the proportion of non-overlapping 742kb windows of comparable effective sequence length where the $PBS_{A:B:C}$ value is greater than or equal to that observed at the focal 742kb region. A *P-value* less than 0.05 is considered statistically significant. Note that when the target population is MXL, $PBS_{MXL:B:C}$ is always significant, and is never significant any other AMR population is the target population.

Table S51:

| $A$ Pop. | $B$ Pop. | $C$ Pop. | Focal 742kb Region ($PBS_{A:B:C}$) | 742kb Non-overlapping Windows ($\mu$) | 742kb Non-overlapping Windows ($\sigma$) | 742kb Non-overlapping Windows ($SEM$) | 742kb Non-overlapping Windows ($\pm CI_{95\%}$) | $P-value$ |
|---|---|---|---|---|---|---|---|---|
| CLM | BEB | CEU | 0.006 | 0.004 | 0.006 | $1.089e^{-4}$ | $2.136e^{-4}$ | 0.258 |
| CLM | BEB | FIN | 0 | 0.006 | 0.007 | $1.234e^{-4}$ | $2.419e^{-4}$ | $> 0.99969$ |
| CLM | BEB | GBR | 0.008 | 0.004 | 0.006 | $1.089e^{-4}$ | $2.135e^{-4}$ | 0.201 |
| CLM | BEB | IBS | 0.002 | 0.004 | 0.006 | $1.018e^{-4}$ | $1.996e^{-4}$ | 0.425 |
| CLM | BEB | TSI | 0.006 | 0.005 | 0.006 | $1.092e^{-4}$ | $2.142e^{-4}$ | 0.289 |
| CLM | CDX | CEU | 0 | 0.001 | 0.005 | $8.084e^{-5}$ | $1.585e^{-4}$ | $> 0.99969$ |
| CLM | CDX | FIN | 0 | 0.003 | 0.007 | $1.234e^{-4}$ | $2.420e^{-4}$ | $> 0.99969$ |
| CLM | CDX | GBR | 0 | 0.002 | 0.005 | $8.658e^{-5}$ | $1.698e^{-4}$ | $> 0.99969$ |
| CLM | CDX | IBS | 0 | 0.001 | 0.004 | $7.604e^{-5}$ | $1.491e^{-4}$ | $> 0.99969$ |
| CLM | CDX | TSI | 0 | 0.002 | 0.005 | $8.337e^{-5}$ | $1.635e^{-4}$ | $> 0.99969$ |
| CLM | CHB | CEU | 0 | 0.001 | 0.005 | $8.000e^{-5}$ | $1.568e^{-4}$ | $> 0.99969$ |
| CLM | CHB | FIN | 0 | 0.003 | 0.007 | $1.250e^{-4}$ | $2.451e^{-4}$ | $> 0.99969$ |
| CLM | CHB | GBR | $2.876e^{-4}$ | 0.001 | 0.005 | $8.710e^{-5}$ | $1.708e^{-4}$ | 0.185 |

Continued on next page

Table S51: (Continued)

| $A$ Pop. | $B$ Pop. | $C$ Pop. | Focal 742kb Region ($PBS_{A:B:C}$) | 742kb Non-overlapping Windows ($\mu$) | 742kb Non-overlapping Windows ($\sigma$) | 742kb Non-overlapping Windows ($SEM$) | 742kb Non-overlapping Windows ($\pm CI_{95\%}$) | $P - value$ |
|---|---|---|---|---|---|---|---|---|
| CLM | CHB | IBS | 0.004 | 0.001 | 0.004 | $7.508e^{-5}$ | $1.472e^{-4}$ | 0.095 |
| CLM | CHB | TSI | 0.003 | 0.001 | 0.005 | $8.209e^{-5}$ | $1.609e^{-4}$ | 0.137 |
| CLM | CHS | CEU | 0 | 0.001 | 0.005 | $8.113e^{-5}$ | $1.591e^{-4}$ | $> 0.99969$ |
| CLM | CHS | FIN | 0 | 0.003 | 0.007 | $1.255e^{-4}$ | $2.461e^{-4}$ | $> 0.99969$ |
| CLM | CHS | GBR | 0 | 0.002 | 0.005 | $8.792e^{-5}$ | $1.724e^{-4}$ | $> 0.99969$ |
| CLM | CHS | IBS | 0.003 | 0.001 | 0.004 | $7.564e^{-5}$ | $1.483e^{-4}$ | 0.111 |
| CLM | CHS | TSI | 0.002 | 0.001 | 0.005 | $8.282e^{-5}$ | $1.624e^{-4}$ | 0.16 |
| CLM | GIH | CEU | 0.01 | 0.006 | 0.007 | $1.283e^{-4}$ | $2.517e^{-4}$ | 0.193 |
| CLM | GIH | FIN | $9.950e^{-4}$ | 0.007 | 0.008 | $1.360e^{-4}$ | $2.666e^{-4}$ | 0.751 |
| CLM | GIH | GBR | 0.012 | 0.006 | 0.007 | $1.281e^{-4}$ | $2.512e^{-4}$ | 0.159 |
| CLM | GIH | IBS | 0.004 | 0.005 | 0.007 | $1.202e^{-4}$ | $2.357e^{-4}$ | 0.458 |
| CLM | GIH | TSI | 0.008 | 0.006 | 0.007 | $1.286e^{-4}$ | $2.521e^{-4}$ | 0.295 |
| CLM | ITU | CEU | 0.013 | 0.005 | 0.007 | $1.187e^{-4}$ | $2.327e^{-4}$ | 0.107 |
| CLM | ITU | FIN | 0 | 0.006 | 0.007 | $1.307e^{-4}$ | $2.562e^{-4}$ | $> 0.99969$ |
| CLM | ITU | GBR | 0.013 | 0.005 | 0.007 | $1.201e^{-4}$ | $2.355e^{-4}$ | 0.116 |
| CLM | ITU | IBS | 0.004 | 0.005 | 0.006 | $1.132e^{-4}$ | $2.220e^{-4}$ | 0.385 |
| CLM | ITU | TSI | 0.009 | 0.006 | 0.007 | $1.225e^{-4}$ | $2.403e^{-4}$ | 0.227 |
| CLM | JPT | CEU | 0 | 0.001 | 0.005 | $7.922e^{-5}$ | $1.553e^{-4}$ | $> 0.99969$ |
| CLM | JPT | FIN | 0 | 0.003 | 0.007 | $1.268e^{-4}$ | $2.487e^{-4}$ | $> 0.99969$ |
| CLM | JPT | GBR | 0 | 0.001 | 0.005 | $8.629e^{-5}$ | $1.692e^{-4}$ | $> 0.99969$ |

Continued on next page

Table S51: (Continued)

| $A$ Pop. | $B$ Pop. | $C$ Pop. | Focal 742kb Region ($PBS_{A:B:C}$) | 742kb Non-overlapping Windows ($\mu$) | 742kb Non-overlapping Windows ($\sigma$) | 742kb Non-overlapping Windows ($SEM$) | 742kb Non-overlapping Windows ($\pm CI_{95\%}$) | $P-value$ |
|---|---|---|---|---|---|---|---|---|
| CLM | JPT | IBS | 0.002 | 0.001 | 0.004 | $7.611e^{-5}$ | $1.492e^{-4}$ | 0.117 |
| CLM | JPT | TSI | 0.001 | 0.001 | 0.005 | $8.167e^{-5}$ | $1.601e^{-4}$ | 0.158 |
| CLM | KHV | CEU | 0 | 0.001 | 0.005 | $8.113e^{-5}$ | $1.591e^{-4}$ | $> 0.99969$ |
| CLM | KHV | FIN | 0 | 0.003 | 0.007 | $1.224e^{-4}$ | $2.400e^{-4}$ | $> 0.99969$ |
| CLM | KHV | GBR | 0 | 0.002 | 0.005 | $8.671e^{-5}$ | $1.700e^{-4}$ | $> 0.99969$ |
| CLM | KHV | IBS | 0.003 | 0.001 | 0.004 | $7.651e^{-5}$ | $1.500e^{-4}$ | 0.123 |
| CLM | KHV | TSI | 0.002 | 0.002 | 0.005 | $8.245e^{-5}$ | $1.616e^{-4}$ | 0.17 |
| CLM | PJL | CEU | 0.015 | 0.006 | 0.007 | $1.251e^{-4}$ | $2.453e^{-4}$ | 0.095 |
| CLM | PJL | FIN | 0.003 | 0.007 | 0.008 | $1.322e^{-4}$ | $2.592e^{-4}$ | 0.672 |
| CLM | PJL | GBR | 0.015 | 0.006 | 0.007 | $1.267e^{-4}$ | $2.484e^{-4}$ | 0.097 |
| CLM | PJL | IBS | 0.007 | 0.006 | 0.007 | $1.189e^{-4}$ | $2.331e^{-4}$ | 0.309 |
| CLM | PJL | TSI | 0.009 | 0.007 | 0.007 | $1.264e^{-4}$ | $2.479e^{-4}$ | 0.278 |
| CLM | STU | CEU | 0.009 | 0.005 | 0.007 | $1.189e^{-4}$ | $2.332e^{-4}$ | 0.2 |
| CLM | STU | FIN | 0 | 0.006 | 0.007 | $1.305e^{-4}$ | $2.559e^{-4}$ | $> 0.99969$ |
| CLM | STU | GBR | 0.01 | 0.005 | 0.007 | $1.195e^{-4}$ | $2.343e^{-4}$ | 0.167 |
| CLM | STU | IBS | 0.004 | 0.004 | 0.006 | $1.133e^{-4}$ | $2.221e^{-4}$ | 0.403 |
| CLM | STU | TSI | 0.007 | 0.005 | 0.007 | $1.224e^{-4}$ | $2.400e^{-4}$ | 0.3 |
| MXL | BEB | CEU | 0.074 | 0.017 | 0.015 | $2.548e^{-4}$ | $4.996e^{-4}$ | 0.007 |
| MXL | BEB | FIN | 0.071 | 0.017 | 0.015 | $2.544e^{-4}$ | $4.988e^{-4}$ | 0.007 |
| MXL | BEB | GBR | 0.078 | 0.017 | 0.015 | $2.551e^{-4}$ | $5.002e^{-4}$ | 0.005 |

Continued on next page

Table S51: (Continued)

| $A$ Pop. | $B$ Pop. | $C$ Pop. | Focal 742kb Region ($PBS_{A:B:C}$) | 742kb Non-overlapping Windows ($\mu$) | 742kb Non-overlapping Windows ($\sigma$) | 742kb Non-overlapping Windows ($SEM$) | 742kb Non-overlapping Windows ($\pm CI_{95\%}$) | $P-value$ |
|---|---|---|---|---|---|---|---|---|
| MXL | BEB | IBS | 0.067 | 0.017 | 0.014 | $2.532e^{-4}$ | $4.964e^{-4}$ | 0.01 |
| MXL | BEB | TSI | 0.069 | 0.018 | 0.015 | $2.554e^{-4}$ | $5.007e^{-4}$ | 0.011 |
| MXL | CDX | CEU | 0.055 | 0.006 | 0.012 | $2.074e^{-4}$ | $4.067e^{-4}$ | 0.01 |
| MXL | CDX | FIN | 0.052 | 0.008 | 0.013 | $2.306e^{-4}$ | $4.521e^{-4}$ | 0.015 |
| MXL | CDX | GBR | 0.06 | 0.006 | 0.012 | $2.109e^{-4}$ | $4.135e^{-4}$ | 0.007 |
| MXL | CDX | IBS | 0.053 | 0.006 | 0.012 | $2.123e^{-4}$ | $4.163e^{-4}$ | 0.01 |
| MXL | CDX | TSI | 0.056 | 0.006 | 0.012 | $2.146e^{-4}$ | $4.207e^{-4}$ | 0.01 |
| MXL | CHB | CEU | 0.066 | 0.005 | 0.011 | $1.949e^{-4}$ | $3.822e^{-4}$ | 0.004 |
| MXL | CHB | FIN | 0.061 | 0.007 | 0.013 | $2.222e^{-4}$ | $4.356e^{-4}$ | 0.006 |
| MXL | CHB | GBR | 0.072 | 0.005 | 0.011 | $1.995e^{-4}$ | $3.911e^{-4}$ | 0.004 |
| MXL | CHB | IBS | 0.069 | 0.005 | 0.011 | $1.999e^{-4}$ | $3.920e^{-4}$ | 0.004 |
| MXL | CHB | TSI | 0.066 | 0.006 | 0.012 | $2.031e^{-4}$ | $3.982e^{-4}$ | 0.005 |
| MXL | CHS | CEU | 0.064 | 0.005 | 0.011 | $2.001e^{-4}$ | $3.923e^{-4}$ | 0.005 |
| MXL | CHS | FIN | 0.06 | 0.007 | 0.013 | $2.268e^{-4}$ | $4.447e^{-4}$ | 0.009 |
| MXL | CHS | GBR | 0.07 | 0.006 | 0.012 | $2.041e^{-4}$ | $4.002e^{-4}$ | 0.005 |
| MXL | CHS | IBS | 0.069 | 0.006 | 0.012 | $2.046e^{-4}$ | $4.011e^{-4}$ | 0.004 |
| MXL | CHS | TSI | 0.066 | 0.006 | 0.012 | $2.069e^{-4}$ | $4.056e^{-4}$ | 0.005 |
| MXL | GIH | CEU | 0.087 | 0.021 | 0.016 | $2.773e^{-4}$ | $5.436e^{-4}$ | 0.004 |
| MXL | GIH | FIN | 0.085 | 0.02 | 0.016 | $2.741e^{-4}$ | $5.375e^{-4}$ | 0.006 |
| MXL | GIH | GBR | 0.091 | 0.021 | 0.016 | $2.783e^{-4}$ | $5.457e^{-4}$ | 0.004 |

Continued on next page

Table S51: (Continued)

| $A$ Pop. | $B$ Pop. | $C$ Pop. | Focal 742kb Region ($PBS_{A:B:C}$) | 742kb Non-overlapping Windows ($\mu$) | 742kb Non-overlapping Windows ($\sigma$) | 742kb Non-overlapping Windows ($SEM$) | 742kb Non-overlapping Windows ($\pm CI_{95\%}$) | $P-value$ |
|---|---|---|---|---|---|---|---|---|
| MXL | GIH | IBS | 0.078 | 0.021 | 0.016 | $2.743e^{-4}$ | $5.378e^{-4}$ | 0.008 |
| MXL | GIH | TSI | 0.08 | 0.022 | 0.016 | $2.766e^{-4}$ | $5.424e^{-4}$ | 0.007 |
| MXL | ITU | CEU | 0.085 | 0.019 | 0.015 | $2.709e^{-4}$ | $5.312e^{-4}$ | 0.006 |
| MXL | ITU | FIN | 0.077 | 0.018 | 0.015 | $2.685e^{-4}$ | $5.265e^{-4}$ | 0.008 |
| MXL | ITU | GBR | 0.087 | 0.019 | 0.016 | $2.720e^{-4}$ | $5.334e^{-4}$ | 0.005 |
| MXL | ITU | IBS | 0.073 | 0.019 | 0.015 | $2.693e^{-4}$ | $5.280e^{-4}$ | 0.01 |
| MXL | ITU | TSI | 0.076 | 0.02 | 0.016 | $2.725e^{-4}$ | $5.343e^{-4}$ | 0.008 |
| MXL | JPT | CEU | 0.063 | 0.005 | 0.011 | $1.948e^{-4}$ | $3.820e^{-4}$ | 0.005 |
| MXL | JPT | FIN | 0.059 | 0.007 | 0.013 | $2.250e^{-4}$ | $4.411e^{-4}$ | 0.009 |
| MXL | JPT | GBR | 0.069 | 0.005 | 0.012 | $2.014e^{-4}$ | $3.949e^{-4}$ | 0.004 |
| MXL | JPT | IBS | 0.068 | 0.005 | 0.012 | $2.011e^{-4}$ | $3.943e^{-4}$ | 0.003 |
| MXL | JPT | TSI | 0.066 | 0.005 | 0.012 | $2.026e^{-4}$ | $3.973e^{-4}$ | 0.004 |
| MXL | KHV | CEU | 0.063 | 0.006 | 0.012 | $2.042e^{-4}$ | $4.004e^{-4}$ | 0.006 |
| MXL | KHV | FIN | 0.059 | 0.008 | 0.013 | $2.280e^{-4}$ | $4.470e^{-4}$ | 0.01 |
| MXL | KHV | GBR | 0.067 | 0.006 | 0.012 | $2.073e^{-4}$ | $4.065e^{-4}$ | 0.005 |
| MXL | KHV | IBS | 0.065 | 0.006 | 0.012 | $2.091e^{-4}$ | $4.099e^{-4}$ | 0.006 |
| MXL | KHV | TSI | 0.062 | 0.006 | 0.012 | $2.114e^{-4}$ | $4.145e^{-4}$ | 0.007 |
| MXL | PJL | CEU | 0.096 | 0.022 | 0.016 | $2.728e^{-4}$ | $5.349e^{-4}$ | 0.003 |
| MXL | PJL | FIN | 0.09 | 0.021 | 0.015 | $2.688e^{-4}$ | $5.270e^{-4}$ | 0.004 |
| MXL | PJL | GBR | 0.099 | 0.022 | 0.016 | $2.743e^{-4}$ | $5.378e^{-4}$ | 0.002 |

Continued on next page

Table S51: (Continued)

| $A$ Pop. | $B$ Pop. | $C$ Pop. | Focal 742kb Region ($PBS_{A:B:C}$) | 742kb Non-overlapping Windows ($\mu$) | 742kb Non-overlapping Windows ($\sigma$) | 742kb Non-overlapping Windows ($SEM$) | 742kb Non-overlapping Windows ($\pm CI_{95\%}$) | $P-value$ |
|---|---|---|---|---|---|---|---|---|
| MXL | PJL | IBS | 0.084 | 0.021 | 0.015 | $2.703e^{-4}$ | $5.300e^{-4}$ | 0.005 |
| MXL | PJL | TSI | 0.085 | 0.023 | 0.016 | $2.721e^{-4}$ | $5.335e^{-4}$ | 0.006 |
| MXL | STU | CEU | 0.08 | 0.019 | 0.015 | $2.707e^{-4}$ | $5.308e^{-4}$ | 0.007 |
| MXL | STU | FIN | 0.076 | 0.018 | 0.015 | $2.687e^{-4}$ | $5.269e^{-4}$ | 0.008 |
| MXL | STU | GBR | 0.084 | 0.019 | 0.015 | $2.709e^{-4}$ | $5.312e^{-4}$ | 0.006 |
| MXL | STU | IBS | 0.071 | 0.019 | 0.015 | $2.691e^{-4}$ | $5.277e^{-4}$ | 0.01 |
| MXL | STU | TSI | 0.073 | 0.02 | 0.016 | $2.726e^{-4}$ | $5.344e^{-4}$ | 0.011 |
| PEL | BEB | CEU | 0.039 | 0.06 | 0.035 | $6.051e^{-4}$ | 0.001 | 0.723 |
| PEL | BEB | FIN | 0.039 | 0.058 | 0.034 | $5.977e^{-4}$ | 0.001 | 0.696 |
| PEL | BEB | GBR | 0.042 | 0.06 | 0.035 | $6.056e^{-4}$ | 0.001 | 0.676 |
| PEL | BEB | IBS | 0.032 | 0.061 | 0.035 | $6.038e^{-4}$ | 0.001 | 0.826 |
| PEL | BEB | TSI | 0.036 | 0.062 | 0.035 | $6.063e^{-4}$ | 0.001 | 0.788 |
| PEL | CDX | CEU | 0.02 | 0.034 | 0.031 | $5.445e^{-4}$ | 0.001 | 0.627 |
| PEL | CDX | FIN | 0.02 | 0.036 | 0.032 | $5.508e^{-4}$ | 0.001 | 0.659 |
| PEL | CDX | GBR | 0.023 | 0.034 | 0.031 | $5.480e^{-4}$ | 0.001 | 0.585 |
| PEL | CDX | IBS | 0.018 | 0.035 | 0.031 | $5.482e^{-4}$ | 0.001 | 0.687 |
| PEL | CDX | TSI | 0.022 | 0.036 | 0.031 | $5.507e^{-4}$ | 0.001 | 0.624 |
| PEL | CHB | CEU | 0.023 | 0.033 | 0.031 | $5.384e^{-4}$ | 0.001 | 0.555 |
| PEL | CHB | FIN | 0.021 | 0.034 | 0.031 | $5.470e^{-4}$ | 0.001 | 0.61 |
| PEL | CHB | GBR | 0.027 | 0.033 | 0.031 | $5.418e^{-4}$ | 0.001 | 0.482 |

Continued on next page

Table S51: (Continued)

| $A$ Pop. | $B$ Pop. | $C$ Pop. | Focal 742kb Region ($PBS_{A:B:C}$) | 742kb Non-overlapping Windows ($\mu$) | 742kb Non-overlapping Windows ($\sigma$) | 742kb Non-overlapping Windows ($SEM$) | 742kb Non-overlapping Windows ($\pm CI_{95\%}$) | $P-value$ |
|---|---|---|---|---|---|---|---|---|
| PEL | CHB | IBS | 0.026 | 0.033 | 0.031 | $5.415e^{-4}$ | 0.001 | 0.52 |
| PEL | CHB | TSI | 0.025 | 0.034 | 0.031 | $5.437e^{-4}$ | 0.001 | 0.541 |
| PEL | CHS | CEU | 0.02 | 0.033 | 0.031 | $5.427e^{-4}$ | 0.001 | 0.607 |
| PEL | CHS | FIN | 0.019 | 0.034 | 0.032 | $5.511e^{-4}$ | 0.001 | 0.646 |
| PEL | CHS | GBR | 0.024 | 0.033 | 0.031 | $5.460e^{-4}$ | 0.001 | 0.534 |
| PEL | CHS | IBS | 0.025 | 0.034 | 0.031 | $5.454e^{-4}$ | 0.001 | 0.537 |
| PEL | CHS | TSI | 0.024 | 0.034 | 0.031 | $5.473e^{-4}$ | 0.001 | 0.559 |
| PEL | GIH | CEU | 0.053 | 0.066 | 0.037 | $6.392e^{-4}$ | 0.001 | 0.59 |
| PEL | GIH | FIN | 0.053 | 0.064 | 0.036 | $6.296e^{-4}$ | 0.001 | 0.556 |
| PEL | GIH | GBR | 0.056 | 0.066 | 0.037 | $6.399e^{-4}$ | 0.001 | 0.553 |
| PEL | GIH | IBS | 0.043 | 0.067 | 0.036 | $6.371e^{-4}$ | 0.001 | 0.743 |
| PEL | GIH | TSI | 0.047 | 0.068 | 0.037 | $6.409e^{-4}$ | 0.001 | 0.698 |
| PEL | ITU | CEU | 0.048 | 0.064 | 0.036 | $6.271e^{-4}$ | 0.001 | 0.624 |
| PEL | ITU | FIN | 0.044 | 0.061 | 0.035 | $6.186e^{-4}$ | 0.001 | 0.661 |
| PEL | ITU | GBR | 0.049 | 0.064 | 0.036 | $6.278e^{-4}$ | 0.001 | 0.604 |
| PEL | ITU | IBS | 0.036 | 0.064 | 0.036 | $6.265e^{-4}$ | 0.001 | 0.806 |
| PEL | ITU | TSI | 0.041 | 0.066 | 0.036 | $6.302e^{-4}$ | 0.001 | 0.751 |
| PEL | JPT | CEU | 0.017 | 0.032 | 0.031 | $5.376e^{-4}$ | 0.001 | 0.639 |
| PEL | JPT | FIN | 0.015 | 0.034 | 0.031 | $5.478e^{-4}$ | 0.001 | 0.705 |
| PEL | JPT | GBR | 0.021 | 0.032 | 0.031 | $5.430e^{-4}$ | 0.001 | 0.576 |

Continued on next page

Table S51: (Continued)

| $A$ Pop. | $B$ Pop. | $C$ Pop. | Focal 742kb Region ($PBS_{A:B:C}$) | 742kb Non-overlapping Windows ($\mu$) | 742kb Non-overlapping Windows ($\sigma$) | 742kb Non-overlapping Windows ($SEM$) | 742kb Non-overlapping Windows ($\pm CI_{95\%}$) | $P - value$ |
|---|---|---|---|---|---|---|---|---|
| PEL | JPT | IBS | 0.022 | 0.033 | 0.031 | $5.427e^{-4}$ | 0.001 | 0.581 |
| PEL | JPT | TSI | 0.021 | 0.033 | 0.031 | $5.438e^{-4}$ | 0.001 | 0.597 |
| PEL | KHV | CEU | 0.021 | 0.035 | 0.031 | $5.436e^{-4}$ | 0.001 | 0.614 |
| PEL | KHV | FIN | 0.02 | 0.036 | 0.032 | $5.512e^{-4}$ | 0.001 | 0.653 |
| PEL | KHV | GBR | 0.024 | 0.035 | 0.031 | $5.466e^{-4}$ | 0.001 | 0.562 |
| PEL | KHV | IBS | 0.023 | 0.036 | 0.031 | $5.472e^{-4}$ | 0.001 | 0.594 |
| PEL | KHV | TSI | 0.022 | 0.036 | 0.031 | $5.489e^{-4}$ | 0.001 | 0.619 |
| PEL | PJL | CEU | 0.056 | 0.068 | 0.036 | $6.295e^{-4}$ | 0.001 | 0.57 |
| PEL | PJL | FIN | 0.053 | 0.065 | 0.035 | $6.190e^{-4}$ | 0.001 | 0.573 |
| PEL | PJL | GBR | 0.058 | 0.068 | 0.036 | $6.311e^{-4}$ | 0.001 | 0.541 |
| PEL | PJL | IBS | 0.044 | 0.068 | 0.036 | $6.285e^{-4}$ | 0.001 | 0.739 |
| PEL | PJL | TSI | 0.047 | 0.069 | 0.036 | $6.324e^{-4}$ | 0.001 | 0.724 |
| PEL | STU | CEU | 0.044 | 0.063 | 0.036 | $6.291e^{-4}$ | 0.001 | 0.682 |
| PEL | STU | FIN | 0.043 | 0.061 | 0.035 | $6.204e^{-4}$ | 0.001 | 0.661 |
| PEL | STU | GBR | 0.047 | 0.063 | 0.036 | $6.286e^{-4}$ | 0.001 | 0.641 |
| PEL | STU | IBS | 0.036 | 0.064 | 0.036 | $6.285e^{-4}$ | 0.001 | 0.809 |
| PEL | STU | TSI | 0.039 | 0.065 | 0.036 | $6.326e^{-4}$ | 0.001 | 0.77 |
| PUR | BEB | CEU | 0.001 | 0.003 | 0.004 | $7.417e^{-5}$ | $1.454e^{-4}$ | 0.449 |
| PUR | BEB | FIN | 0 | 0.005 | 0.006 | $1.017e^{-4}$ | $1.994e^{-4}$ | $> 0.99969$ |
| PUR | BEB | GBR | 0.003 | 0.003 | 0.004 | $7.677e^{-5}$ | $1.505e^{-4}$ | 0.339 |

Continued on next page

Table S51: (Continued)

| $A$ Pop. | $B$ Pop. | $C$ Pop. | Focal 742kb Region ($PBS_{A:B:C}$) | 742kb Non-overlapping Windows ($\mu$) | 742kb Non-overlapping Windows ($\sigma$) | 742kb Non-overlapping Windows ($SEM$) | 742kb Non-overlapping Windows ($\pm CI_{95\%}$) | $P-value$ |
|---|---|---|---|---|---|---|---|---|
| PUR | BEB | IBS | 0 | 0.002 | 0.003 | $6.079e^{-5}$ | $1.192e^{-4}$ | $> 0.99969$ |
| PUR | BEB | TSI | 0.002 | 0.003 | 0.004 | $7.118e^{-5}$ | $1.396e^{-4}$ | 0.36 |
| PUR | CDX | CEU | 0 | 0.001 | 0.004 | $7.254e^{-5}$ | $1.422e^{-4}$ | $> 0.99969$ |
| PUR | CDX | FIN | 0 | 0.004 | 0.007 | $1.265e^{-4}$ | $2.479e^{-4}$ | $> 0.99969$ |
| PUR | CDX | GBR | 0 | 0.001 | 0.004 | $7.866e^{-5}$ | $1.542e^{-4}$ | $> 0.99969$ |
| PUR | CDX | IBS | 0 | $9.516e^{-4}$ | 0.003 | $5.895e^{-5}$ | $1.156e^{-4}$ | $> 0.99969$ |
| PUR | CDX | TSI | 0 | 0.001 | 0.004 | $6.639e^{-5}$ | $1.302e^{-4}$ | $> 0.99969$ |
| PUR | CHB | CEU | 0 | 0.001 | 0.004 | $7.056e^{-5}$ | $1.383e^{-4}$ | $> 0.99969$ |
| PUR | CHB | FIN | 0 | 0.004 | 0.007 | $1.279e^{-4}$ | $2.507e^{-4}$ | $> 0.99969$ |
| PUR | CHB | GBR | 0 | 0.001 | 0.004 | $7.815e^{-5}$ | $1.532e^{-4}$ | $> 0.99969$ |
| PUR | CHB | IBS | 0.001 | $9.346e^{-4}$ | 0.003 | $5.793e^{-5}$ | $1.136e^{-4}$ | 0.14 |
| PUR | CHB | TSI | 0.001 | 0.001 | 0.004 | $6.600e^{-5}$ | $1.294e^{-4}$ | 0.166 |
| PUR | CHS | CEU | 0 | 0.001 | 0.004 | $7.180e^{-5}$ | $1.408e^{-4}$ | $> 0.99969$ |
| PUR | CHS | FIN | 0 | 0.004 | 0.007 | $1.278e^{-4}$ | $2.506e^{-4}$ | $> 0.99969$ |
| PUR | CHS | GBR | 0 | 0.001 | 0.005 | $7.924e^{-5}$ | $1.554e^{-4}$ | $> 0.99969$ |
| PUR | CHS | IBS | 0 | $9.669e^{-4}$ | 0.003 | $5.899e^{-5}$ | $1.157e^{-4}$ | $> 0.99969$ |
| PUR | CHS | TSI | 0 | 0.001 | 0.004 | $6.720e^{-5}$ | $1.318e^{-4}$ | $> 0.99969$ |
| PUR | GIH | CEU | 0.006 | 0.004 | 0.005 | $8.702e^{-5}$ | $1.706e^{-4}$ | 0.214 |
| PUR | GIH | FIN | 0.003 | 0.005 | 0.006 | $1.072e^{-4}$ | $2.103e^{-4}$ | 0.547 |
| PUR | GIH | GBR | 0.008 | 0.004 | 0.005 | $9.014e^{-5}$ | $1.767e^{-4}$ | 0.17 |

Continued on next page

Table S51: (Continued)

| $A$ Pop. | $B$ Pop. | $C$ Pop. | Focal 742kb Region ($PBS_{A:B:C}$) | 742kb Non-overlapping Windows ($\mu$) | 742kb Non-overlapping Windows ($\sigma$) | 742kb Non-overlapping Windows ($SEM$) | 742kb Non-overlapping Windows ($\pm CI_{95\%}$) | $P-value$ |
|---|---|---|---|---|---|---|---|---|
| PUR | GIH | IBS | 0 | 0.003 | 0.004 | $7.321e^{-5}$ | $1.435e^{-4}$ | $> 0.99969$ |
| PUR | GIH | TSI | 0.005 | 0.004 | 0.005 | $8.378e^{-5}$ | $1.643e^{-4}$ | 0.255 |
| PUR | ITU | CEU | 0.007 | 0.003 | 0.005 | $7.992e^{-5}$ | $1.567e^{-4}$ | 0.139 |
| PUR | ITU | FIN | 0 | 0.005 | 0.006 | $1.043e^{-4}$ | $2.045e^{-4}$ | $> 0.99969$ |
| PUR | ITU | GBR | 0.007 | 0.003 | 0.005 | $8.330e^{-5}$ | $1.633e^{-4}$ | 0.158 |
| PUR | ITU | IBS | 0 | 0.002 | 0.004 | $6.802e^{-5}$ | $1.334e^{-4}$ | $> 0.99969$ |
| PUR | ITU | TSI | 0.004 | 0.003 | 0.005 | $7.998e^{-5}$ | $1.568e^{-4}$ | 0.27 |
| PUR | JPT | CEU | 0 | 0.001 | 0.004 | $7.068e^{-5}$ | $1.386e^{-4}$ | $> 0.99969$ |
| PUR | JPT | FIN | 0 | 0.004 | 0.007 | $1.295e^{-4}$ | $2.540e^{-4}$ | $> 0.99969$ |
| PUR | JPT | GBR | 0 | 0.001 | 0.004 | $7.742e^{-5}$ | $1.518e^{-4}$ | $> 0.99969$ |
| PUR | JPT | IBS | 0 | $9.313e^{-4}$ | 0.003 | $5.805e^{-5}$ | $1.138e^{-4}$ | $> 0.99969$ |
| PUR | JPT | TSI | 0 | 0.001 | 0.004 | $6.564e^{-5}$ | $1.287e^{-4}$ | $> 0.99969$ |
| PUR | KHV | CEU | 0 | 0.001 | 0.004 | $7.099e^{-5}$ | $1.392e^{-4}$ | $> 0.99969$ |
| PUR | KHV | FIN | 0 | 0.004 | 0.007 | $1.244e^{-4}$ | $2.438e^{-4}$ | $> 0.99969$ |
| PUR | KHV | GBR | 0 | 0.001 | 0.004 | $7.780e^{-5}$ | $1.525e^{-4}$ | $> 0.99969$ |
| PUR | KHV | IBS | 0 | $9.538e^{-4}$ | 0.003 | $5.911e^{-5}$ | $1.159e^{-4}$ | $> 0.99969$ |
| PUR | KHV | TSI | 0 | 0.001 | 0.004 | $6.529e^{-5}$ | $1.280e^{-4}$ | $> 0.99969$ |
| PUR | PJL | CEU | 0.01 | 0.004 | 0.005 | $8.426e^{-5}$ | $1.652e^{-4}$ | 0.107 |
| PUR | PJL | FIN | 0.003 | 0.005 | 0.006 | $1.011e^{-4}$ | $1.983e^{-4}$ | 0.555 |
| PUR | PJL | GBR | 0.01 | 0.004 | 0.005 | $8.766e^{-5}$ | $1.719e^{-4}$ | 0.108 |

Continued on next page

Table S51: (Continued)

| $A$ Pop. | $B$ Pop. | $C$ Pop. | Focal 742kb Region ($PBS_{A:B:C}$) | 742kb Non-overlapping Windows ($\mu$) | 742kb Non-overlapping Windows ($\sigma$) | 742kb Non-overlapping Windows ($SEM$) | 742kb Non-overlapping Windows ($\pm CI_{95\%}$) | $P - value$ |
|---|---|---|---|---|---|---|---|---|
| PUR | PJL | IBS | 0.001 | 0.003 | 0.004 | $7.159e^{-5}$ | $1.404e^{-4}$ | 0.51 |
| PUR | PJL | TSI | 0.005 | 0.004 | 0.005 | $8.280e^{-5}$ | $1.623e^{-4}$ | 0.324 |
| PUR | STU | CEU | 0.004 | 0.003 | 0.005 | $8.017e^{-5}$ | $1.572e^{-4}$ | 0.251 |
| PUR | STU | FIN | $1.118e^{-4}$ | 0.005 | 0.006 | $1.046e^{-4}$ | $2.052e^{-4}$ | 0.673 |
| PUR | STU | GBR | 0.005 | 0.003 | 0.005 | $8.247e^{-5}$ | $1.617e^{-4}$ | 0.2 |
| PUR | STU | IBS | 0 | 0.002 | 0.004 | $6.675e^{-5}$ | $1.309e^{-4}$ | $> 0.99969$ |
| PUR | STU | TSI | 0.004 | 0.003 | 0.005 | $7.916e^{-5}$ | $1.552e^{-4}$ | 0.29 |

**Table S52. $PBS_{AMR:EAS/SAS:EUR}$ values for the focal 72kb *MUC19* region.**
The $PBS_{A:B:C}$ values observed in the focal 72kb region for all unique combinations of: AMR populations as the target population ($A$), EAS and SAS populations as one of the control populations ($B$), and EUR populations as the second control population ($C$). For each unique $PBS_{A:B:C}$ configuration, the mean ($\mu$), standard deviation ($\sigma$), standard error of the mean ($SEM$), and 95% confidence intervals ($\pm CI_{95\%}$) of the $PBS_{A:B:C}$ genomic background distribution used to compute the *P-value* are also reported. The *P-value* represents the proportion of non-overlapping 72kb windows of comparable effective sequence length where the $PBS_{A:B:C}$ value is greater than or equal to that observed at the focal 72kb region. A *P-value* less than 0.05 is considered statistically significant. Note that when the target population is MXL, $PBS_{MXL:B:C}$ is always significant, and is never significant any other AMR population is the target population.

Table S52:

| $A$ Pop. | $B$ Pop. | $C$ Pop. | Focal 72kb Region ($PBS_{A:B:C}$) | 72kb Non-overlapping Windows ($\mu$) | 72kb Non-overlapping Windows ($\sigma$) | 72kb Non-overlapping Windows ($SEM$) | 72kb Non-overlapping Windows ($\pm CI_{95\%}$) | $P-value$ |
|---|---|---|---|---|---|---|---|---|
| CLM | BEB | CEU | 0 | 0.006 | 0.011 | $6.008e^{-5}$ | $1.178e^{-4}$ | $> 0.999968$ |
| CLM | BEB | FIN | 0 | 0.008 | 0.012 | $6.724e^{-5}$ | $1.318e^{-4}$ | $> 0.999968$ |
| CLM | BEB | GBR | 0 | 0.006 | 0.011 | $6.032e^{-5}$ | $1.182e^{-4}$ | $> 0.999968$ |
| CLM | BEB | IBS | 0 | 0.006 | 0.01 | $5.683e^{-5}$ | $1.114e^{-4}$ | $> 0.999968$ |
| CLM | BEB | TSI | 0 | 0.006 | 0.011 | $6.095e^{-5}$ | $1.195e^{-4}$ | $> 0.999968$ |
| CLM | CDX | CEU | 0 | 0.004 | 0.01 | $5.639e^{-5}$ | $1.105e^{-4}$ | $> 0.999968$ |
| CLM | CDX | FIN | 0 | 0.007 | 0.014 | $7.666e^{-5}$ | $1.503e^{-4}$ | $> 0.999968$ |
| CLM | CDX | GBR | 0 | 0.004 | 0.01 | $5.853e^{-5}$ | $1.147e^{-4}$ | $> 0.999968$ |
| CLM | CDX | IBS | 0 | 0.004 | 0.009 | $5.332e^{-5}$ | $1.045e^{-4}$ | $> 0.999968$ |
| CLM | CDX | TSI | 0 | 0.004 | 0.01 | $5.667e^{-5}$ | $1.111e^{-4}$ | $> 0.999968$ |
| CLM | CHB | CEU | 0.002 | 0.004 | 0.01 | $5.465e^{-5}$ | $1.071e^{-4}$ | 0.248 |
| CLM | CHB | FIN | 0 | 0.007 | 0.014 | $7.609e^{-5}$ | $1.491e^{-4}$ | $> 0.999968$ |
| CLM | CHB | GBR | 0.001 | 0.004 | 0.01 | $5.690e^{-5}$ | $1.115e^{-4}$ | 0.273 |

Continued on next page

Table S52: (Continued)

| $A$ Pop. | $B$ Pop. | $C$ Pop. | Focal 72kb Region ($PBS_{A:B:C}$) | 72kb Non-overlapping Windows ($\mu$) | 72kb Non-overlapping Windows ($\sigma$) | 72kb Non-overlapping Windows ($SEM$) | 72kb Non-overlapping Windows ($\pm CI_{95\%}$) | $P-value$ |
|---|---|---|---|---|---|---|---|---|
| CLM | CHB | IBS | 0.009 | 0.004 | 0.009 | $5.160e^{-5}$ | $1.011e^{-4}$ | 0.128 |
| CLM | CHB | TSI | 0 | 0.004 | 0.01 | $5.495e^{-5}$ | $1.077e^{-4}$ | $> 0.999968$ |
| CLM | CHS | CEU | 0.001 | 0.004 | 0.01 | $5.522e^{-5}$ | $1.082e^{-4}$ | 0.285 |
| CLM | CHS | FIN | 0 | 0.007 | 0.014 | $7.605e^{-5}$ | $1.491e^{-4}$ | $> 0.999968$ |
| CLM | CHS | GBR | $5.608e^{-4}$ | 0.004 | 0.01 | $5.773e^{-5}$ | $1.131e^{-4}$ | 0.3 |
| CLM | CHS | IBS | 0.009 | 0.004 | 0.009 | $5.219e^{-5}$ | $1.023e^{-4}$ | 0.139 |
| CLM | CHS | TSI | 0 | 0.004 | 0.01 | $5.543e^{-5}$ | $1.086e^{-4}$ | $> 0.999968$ |
| CLM | GIH | CEU | 0.002 | 0.008 | 0.012 | $6.768e^{-5}$ | $1.327e^{-4}$ | 0.521 |
| CLM | GIH | FIN | 0 | 0.009 | 0.013 | $7.263e^{-5}$ | $1.424e^{-4}$ | $> 0.999968$ |
| CLM | GIH | GBR | $4.810e^{-4}$ | 0.008 | 0.012 | $6.797e^{-5}$ | $1.332e^{-4}$ | 0.608 |
| CLM | GIH | IBS | 0.004 | 0.007 | 0.011 | $6.362e^{-5}$ | $1.247e^{-4}$ | 0.446 |
| CLM | GIH | TSI | 0.003 | 0.008 | 0.012 | $6.819e^{-5}$ | $1.336e^{-4}$ | 0.516 |
| CLM | ITU | CEU | 0.001 | 0.007 | 0.012 | $6.489e^{-5}$ | $1.272e^{-4}$ | 0.528 |
| CLM | ITU | FIN | 0 | 0.008 | 0.013 | $7.100e^{-5}$ | $1.392e^{-4}$ | $> 0.999968$ |
| CLM | ITU | GBR | 0 | 0.007 | 0.012 | $6.550e^{-5}$ | $1.284e^{-4}$ | $> 0.999968$ |
| CLM | ITU | IBS | 0.002 | 0.006 | 0.011 | $6.183e^{-5}$ | $1.212e^{-4}$ | 0.475 |
| CLM | ITU | TSI | 0.003 | 0.007 | 0.012 | $6.664e^{-5}$ | $1.306e^{-4}$ | 0.487 |
| CLM | JPT | CEU | 0 | 0.004 | 0.01 | $5.341e^{-5}$ | $1.047e^{-4}$ | $> 0.999968$ |
| CLM | JPT | FIN | 0 | 0.007 | 0.013 | $7.523e^{-5}$ | $1.475e^{-4}$ | $> 0.999968$ |
| CLM | JPT | GBR | 0 | 0.004 | 0.01 | $5.597e^{-5}$ | $1.097e^{-4}$ | $> 0.999968$ |

Continued on next page

Table S52: (Continued)

| $A$ Pop. | $B$ Pop. | $C$ Pop. | Focal 72kb Region ($PBS_{A:B:C}$) | 72kb Non-overlapping Windows ($\mu$) | 72kb Non-overlapping Windows ($\sigma$) | 72kb Non-overlapping Windows ($SEM$) | 72kb Non-overlapping Windows ($\pm CI_{95\%}$) | $P-value$ |
|---|---|---|---|---|---|---|---|---|
| CLM | JPT | IBS | 0.007 | 0.004 | 0.009 | $5.135e^{-5}$ | $1.007e^{-4}$ | 0.157 |
| CLM | JPT | TSI | 0 | 0.004 | 0.01 | $5.380e^{-5}$ | $1.055e^{-4}$ | $> 0.999968$ |
| CLM | KHV | CEU | 0 | 0.004 | 0.01 | $5.571e^{-5}$ | $1.092e^{-4}$ | $> 0.999968$ |
| CLM | KHV | FIN | 0 | 0.007 | 0.013 | $7.573e^{-5}$ | $1.484e^{-4}$ | $> 0.999968$ |
| CLM | KHV | GBR | 0 | 0.004 | 0.01 | $5.794e^{-5}$ | $1.136e^{-4}$ | $> 0.999968$ |
| CLM | KHV | IBS | 0.006 | 0.004 | 0.009 | $5.286e^{-5}$ | $1.036e^{-4}$ | 0.184 |
| CLM | KHV | TSI | 0 | 0.004 | 0.01 | $5.613e^{-5}$ | $1.100e^{-4}$ | $> 0.999968$ |
| CLM | PJL | CEU | 0.008 | 0.008 | 0.012 | $6.545e^{-5}$ | $1.283e^{-4}$ | 0.321 |
| CLM | PJL | FIN | $5.767e^{-4}$ | 0.008 | 0.012 | $6.987e^{-5}$ | $1.369e^{-4}$ | 0.647 |
| CLM | PJL | GBR | 0.006 | 0.008 | 0.012 | $6.644e^{-5}$ | $1.302e^{-4}$ | 0.379 |
| CLM | PJL | IBS | 0.005 | 0.007 | 0.011 | $6.206e^{-5}$ | $1.216e^{-4}$ | 0.412 |
| CLM | PJL | TSI | 0.003 | 0.008 | 0.012 | $6.650e^{-5}$ | $1.303e^{-4}$ | 0.511 |
| CLM | STU | CEU | $3.847e^{-4}$ | 0.007 | 0.012 | $6.482e^{-5}$ | $1.271e^{-4}$ | 0.563 |
| CLM | STU | FIN | 0 | 0.008 | 0.013 | $7.054e^{-5}$ | $1.383e^{-4}$ | $> 0.999968$ |
| CLM | STU | GBR | 0 | 0.007 | 0.012 | $6.521e^{-5}$ | $1.278e^{-4}$ | $> 0.999968$ |
| CLM | STU | IBS | 0.003 | 0.006 | 0.011 | $6.190e^{-5}$ | $1.213e^{-4}$ | 0.406 |
| CLM | STU | TSI | 0.002 | 0.007 | 0.012 | $6.638e^{-5}$ | $1.301e^{-4}$ | 0.488 |
| MXL | BEB | CEU | 0.087 | 0.018 | 0.023 | $1.298e^{-4}$ | $2.545e^{-4}$ | 0.021 |
| MXL | BEB | FIN | 0.088 | 0.018 | 0.023 | $1.295e^{-4}$ | $2.538e^{-4}$ | 0.019 |
| MXL | BEB | GBR | 0.088 | 0.018 | 0.023 | $1.298e^{-4}$ | $2.544e^{-4}$ | 0.019 |

Continued on next page

Table S52: (Continued)

| $A$ Pop. | $B$ Pop. | $C$ Pop. | Focal 72kb Region ($PBS_{A:B:C}$) | 72kb Non-overlapping Windows ($\mu$) | 72kb Non-overlapping Windows ($\sigma$) | 72kb Non-overlapping Windows ($SEM$) | 72kb Non-overlapping Windows ($\pm CI_{95\%}$) | $P-value$ |
|---|---|---|---|---|---|---|---|---|
| MXL | BEB | IBS | 0.077 | 0.018 | 0.023 | $1.290e^{-4}$ | $2.529e^{-4}$ | 0.029 |
| MXL | BEB | TSI | 0.074 | 0.019 | 0.023 | $1.315e^{-4}$ | $2.578e^{-4}$ | 0.035 |
| MXL | CDX | CEU | 0.091 | 0.01 | 0.02 | $1.099e^{-4}$ | $2.154e^{-4}$ | 0.009 |
| MXL | CDX | FIN | 0.093 | 0.012 | 0.021 | $1.208e^{-4}$ | $2.368e^{-4}$ | 0.012 |
| MXL | CDX | GBR | 0.092 | 0.01 | 0.02 | $1.108e^{-4}$ | $2.171e^{-4}$ | 0.009 |
| MXL | CDX | IBS | 0.081 | 0.01 | 0.02 | $1.105e^{-4}$ | $2.166e^{-4}$ | 0.014 |
| MXL | CDX | TSI | 0.076 | 0.01 | 0.02 | $1.131e^{-4}$ | $2.217e^{-4}$ | 0.018 |
| MXL | CHB | CEU | 0.127 | 0.009 | 0.019 | $1.040e^{-4}$ | $2.038e^{-4}$ | 0.002 |
| MXL | CHB | FIN | 0.125 | 0.011 | 0.021 | $1.158e^{-4}$ | $2.269e^{-4}$ | 0.004 |
| MXL | CHB | GBR | 0.131 | 0.009 | 0.019 | $1.052e^{-4}$ | $2.061e^{-4}$ | 0.002 |
| MXL | CHB | IBS | 0.12 | 0.009 | 0.019 | $1.048e^{-4}$ | $2.054e^{-4}$ | 0.003 |
| MXL | CHB | TSI | 0.106 | 0.009 | 0.019 | $1.078e^{-4}$ | $2.113e^{-4}$ | 0.005 |
| MXL | CHS | CEU | 0.126 | 0.009 | 0.019 | $1.053e^{-4}$ | $2.064e^{-4}$ | 0.003 |
| MXL | CHS | FIN | 0.125 | 0.011 | 0.021 | $1.171e^{-4}$ | $2.296e^{-4}$ | 0.004 |
| MXL | CHS | GBR | 0.13 | 0.009 | 0.019 | $1.065e^{-4}$ | $2.088e^{-4}$ | 0.002 |
| MXL | CHS | IBS | 0.119 | 0.009 | 0.019 | $1.061e^{-4}$ | $2.079e^{-4}$ | 0.003 |
| MXL | CHS | TSI | 0.107 | 0.01 | 0.019 | $1.087e^{-4}$ | $2.130e^{-4}$ | 0.005 |
| MXL | GIH | CEU | 0.12 | 0.022 | 0.025 | $1.432e^{-4}$ | $2.807e^{-4}$ | 0.009 |
| MXL | GIH | FIN | 0.122 | 0.021 | 0.025 | $1.407e^{-4}$ | $2.757e^{-4}$ | 0.008 |
| MXL | GIH | GBR | 0.122 | 0.022 | 0.026 | $1.433e^{-4}$ | $2.809e^{-4}$ | 0.009 |

Continued on next page

Table S52: (Continued)

| $A$ Pop. | $B$ Pop. | $C$ Pop. | Focal 72kb Region ($PBS_{A:B:C}$) | 72kb Non-overlapping Windows ($\mu$) | 72kb Non-overlapping Windows ($\sigma$) | 72kb Non-overlapping Windows ($SEM$) | 72kb Non-overlapping Windows ($\pm CI_{95\%}$) | $P - value$ |
|---|---|---|---|---|---|---|---|---|
| MXL | GIH | IBS | 0.106 | 0.021 | 0.025 | $1.418e^{-4}$ | $2.779e^{-4}$ | 0.015 |
| MXL | GIH | TSI | 0.103 | 0.023 | 0.026 | $1.445e^{-4}$ | $2.831e^{-4}$ | 0.018 |
| MXL | ITU | CEU | 0.112 | 0.02 | 0.025 | $1.394e^{-4}$ | $2.731e^{-4}$ | 0.011 |
| MXL | ITU | FIN | 0.112 | 0.02 | 0.024 | $1.376e^{-4}$ | $2.697e^{-4}$ | 0.01 |
| MXL | ITU | GBR | 0.113 | 0.02 | 0.025 | $1.394e^{-4}$ | $2.733e^{-4}$ | 0.01 |
| MXL | ITU | IBS | 0.098 | 0.02 | 0.025 | $1.387e^{-4}$ | $2.719e^{-4}$ | 0.018 |
| MXL | ITU | TSI | 0.096 | 0.021 | 0.025 | $1.416e^{-4}$ | $2.776e^{-4}$ | 0.02 |
| MXL | JPT | CEU | 0.126 | 0.009 | 0.018 | $1.027e^{-4}$ | $2.013e^{-4}$ | 0.002 |
| MXL | JPT | FIN | 0.124 | 0.011 | 0.021 | $1.152e^{-4}$ | $2.259e^{-4}$ | 0.004 |
| MXL | JPT | GBR | 0.13 | 0.009 | 0.019 | $1.049e^{-4}$ | $2.056e^{-4}$ | 0.002 |
| MXL | JPT | IBS | 0.119 | 0.009 | 0.019 | $1.043e^{-4}$ | $2.045e^{-4}$ | 0.003 |
| MXL | JPT | TSI | 0.108 | 0.009 | 0.019 | $1.064e^{-4}$ | $2.086e^{-4}$ | 0.005 |
| MXL | KHV | CEU | 0.101 | 0.01 | 0.019 | $1.080e^{-4}$ | $2.117e^{-4}$ | 0.007 |
| MXL | KHV | FIN | 0.099 | 0.011 | 0.021 | $1.190e^{-4}$ | $2.332e^{-4}$ | 0.01 |
| MXL | KHV | GBR | 0.103 | 0.01 | 0.019 | $1.086e^{-4}$ | $2.128e^{-4}$ | 0.006 |
| MXL | KHV | IBS | 0.095 | 0.01 | 0.019 | $1.087e^{-4}$ | $2.131e^{-4}$ | 0.008 |
| MXL | KHV | TSI | 0.084 | 0.01 | 0.02 | $1.114e^{-4}$ | $2.183e^{-4}$ | 0.012 |
| MXL | PJL | CEU | 0.147 | 0.022 | 0.025 | $1.408e^{-4}$ | $2.759e^{-4}$ | 0.004 |
| MXL | PJL | FIN | 0.151 | 0.021 | 0.025 | $1.381e^{-4}$ | $2.707e^{-4}$ | 0.003 |
| MXL | PJL | GBR | 0.15 | 0.022 | 0.025 | $1.412e^{-4}$ | $2.768e^{-4}$ | 0.004 |

Continued on next page

Table S52: (Continued)

| $A$ Pop. | $B$ Pop. | $C$ Pop. | Focal 72kb Region ($PBS_{A:B:C}$) | 72kb Non-overlapping Windows ($\mu$) | 72kb Non-overlapping Windows ($\sigma$) | 72kb Non-overlapping Windows ($SEM$) | 72kb Non-overlapping Windows ($\pm CI_{95\%}$) | $P-value$ |
|---|---|---|---|---|---|---|---|---|
| MXL | PJL | IBS | 0.129 | 0.022 | 0.025 | $1.394e^{-4}$ | $2.733e^{-4}$ | 0.006 |
| MXL | PJL | TSI | 0.125 | 0.023 | 0.025 | $1.420e^{-4}$ | $2.783e^{-4}$ | 0.008 |
| MXL | STU | CEU | 0.107 | 0.02 | 0.025 | $1.396e^{-4}$ | $2.737e^{-4}$ | 0.013 |
| MXL | STU | FIN | 0.107 | 0.019 | 0.025 | $1.377e^{-4}$ | $2.698e^{-4}$ | 0.012 |
| MXL | STU | GBR | 0.109 | 0.02 | 0.025 | $1.392e^{-4}$ | $2.729e^{-4}$ | 0.011 |
| MXL | STU | IBS | 0.095 | 0.02 | 0.025 | $1.389e^{-4}$ | $2.722e^{-4}$ | 0.018 |
| MXL | STU | TSI | 0.091 | 0.021 | 0.025 | $1.416e^{-4}$ | $2.776e^{-4}$ | 0.023 |
| PEL | BEB | CEU | 0.039 | 0.059 | 0.055 | $3.093e^{-4}$ | $6.063e^{-4}$ | 0.566 |
| PEL | BEB | FIN | 0.04 | 0.057 | 0.054 | $3.032e^{-4}$ | $5.944e^{-4}$ | 0.535 |
| PEL | BEB | GBR | 0.039 | 0.059 | 0.055 | $3.095e^{-4}$ | $6.066e^{-4}$ | 0.566 |
| PEL | BEB | IBS | 0.03 | 0.06 | 0.055 | $3.090e^{-4}$ | $6.057e^{-4}$ | 0.668 |
| PEL | BEB | TSI | 0.032 | 0.061 | 0.055 | $3.118e^{-4}$ | $6.111e^{-4}$ | 0.661 |
| PEL | CDX | CEU | 0.038 | 0.037 | 0.047 | $2.639e^{-4}$ | $5.172e^{-4}$ | 0.353 |
| PEL | CDX | FIN | 0.04 | 0.038 | 0.048 | $2.670e^{-4}$ | $5.233e^{-4}$ | 0.35 |
| PEL | CDX | GBR | 0.038 | 0.037 | 0.047 | $2.648e^{-4}$ | $5.191e^{-4}$ | 0.356 |
| PEL | CDX | IBS | 0.029 | 0.038 | 0.047 | $2.658e^{-4}$ | $5.209e^{-4}$ | 0.441 |
| PEL | CDX | TSI | 0.029 | 0.038 | 0.048 | $2.684e^{-4}$ | $5.261e^{-4}$ | 0.448 |
| PEL | CHB | CEU | 0.059 | 0.035 | 0.046 | $2.574e^{-4}$ | $5.046e^{-4}$ | 0.207 |
| PEL | CHB | FIN | 0.057 | 0.037 | 0.047 | $2.615e^{-4}$ | $5.126e^{-4}$ | 0.221 |
| PEL | CHB | GBR | 0.061 | 0.035 | 0.046 | $2.583e^{-4}$ | $5.063e^{-4}$ | 0.197 |

Continued on next page

Table S52: (Continued)

| $A$ Pop. | $B$ Pop. | $C$ Pop. | Focal 72kb Region ($PBS_{A:B:C}$) | 72kb Non-overlapping Windows ($\mu$) | 72kb Non-overlapping Windows ($\sigma$) | 72kb Non-overlapping Windows ($SEM$) | 72kb Non-overlapping Windows ($\pm CI_{95\%}$) | $P-value$ |
|---|---|---|---|---|---|---|---|---|
| PEL | CHB | IBS | 0.053 | 0.036 | 0.046 | $2.592e^{-4}$ | $5.080e^{-4}$ | 0.242 |
| PEL | CHB | TSI | 0.044 | 0.037 | 0.047 | $2.619e^{-4}$ | $5.133e^{-4}$ | 0.302 |
| PEL | CHS | CEU | 0.059 | 0.036 | 0.046 | $2.589e^{-4}$ | $5.075e^{-4}$ | 0.209 |
| PEL | CHS | FIN | 0.058 | 0.037 | 0.047 | $2.627e^{-4}$ | $5.150e^{-4}$ | 0.221 |
| PEL | CHS | GBR | 0.062 | 0.036 | 0.046 | $2.599e^{-4}$ | $5.095e^{-4}$ | 0.197 |
| PEL | CHS | IBS | 0.054 | 0.036 | 0.046 | $2.607e^{-4}$ | $5.109e^{-4}$ | 0.241 |
| PEL | CHS | TSI | 0.046 | 0.037 | 0.047 | $2.631e^{-4}$ | $5.157e^{-4}$ | 0.296 |
| PEL | GIH | CEU | 0.065 | 0.065 | 0.059 | $3.301e^{-4}$ | $6.470e^{-4}$ | 0.382 |
| PEL | GIH | FIN | 0.067 | 0.063 | 0.057 | $3.222e^{-4}$ | $6.314e^{-4}$ | 0.344 |
| PEL | GIH | GBR | 0.066 | 0.065 | 0.059 | $3.304e^{-4}$ | $6.476e^{-4}$ | 0.371 |
| PEL | GIH | IBS | 0.053 | 0.066 | 0.059 | $3.296e^{-4}$ | $6.460e^{-4}$ | 0.477 |
| PEL | GIH | TSI | 0.054 | 0.067 | 0.059 | $3.325e^{-4}$ | $6.517e^{-4}$ | 0.485 |
| PEL | ITU | CEU | 0.058 | 0.063 | 0.058 | $3.241e^{-4}$ | $6.353e^{-4}$ | 0.41 |
| PEL | ITU | FIN | 0.059 | 0.06 | 0.056 | $3.170e^{-4}$ | $6.213e^{-4}$ | 0.388 |
| PEL | ITU | GBR | 0.059 | 0.063 | 0.058 | $3.243e^{-4}$ | $6.356e^{-4}$ | 0.409 |
| PEL | ITU | IBS | 0.046 | 0.064 | 0.058 | $3.242e^{-4}$ | $6.354e^{-4}$ | 0.521 |
| PEL | ITU | TSI | 0.048 | 0.065 | 0.058 | $3.271e^{-4}$ | $6.411e^{-4}$ | 0.513 |
| PEL | JPT | CEU | 0.059 | 0.035 | 0.046 | $2.558e^{-4}$ | $5.014e^{-4}$ | 0.205 |
| PEL | JPT | FIN | 0.058 | 0.036 | 0.046 | $2.603e^{-4}$ | $5.103e^{-4}$ | 0.215 |
| PEL | JPT | GBR | 0.062 | 0.035 | 0.046 | $2.576e^{-4}$ | $5.049e^{-4}$ | 0.193 |

Continued on next page

Table S52: (Continued)

| $A$ Pop. | $B$ Pop. | $C$ Pop. | Focal 72kb Region ($PBS_{A:B:C}$) | 72kb Non-overlapping Windows ($\mu$) | 72kb Non-overlapping Windows ($\sigma$) | 72kb Non-overlapping Windows ($SEM$) | 72kb Non-overlapping Windows ($\pm CI_{95\%}$) | $P-value$ |
|---|---|---|---|---|---|---|---|---|
| PEL | JPT | IBS | 0.054 | 0.036 | 0.046 | $2.586e^{-4}$ | $5.069e^{-4}$ | 0.235 |
| PEL | JPT | TSI | 0.047 | 0.036 | 0.046 | $2.606e^{-4}$ | $5.108e^{-4}$ | 0.281 |
| PEL | KHV | CEU | 0.04 | 0.037 | 0.047 | $2.631e^{-4}$ | $5.157e^{-4}$ | 0.335 |
| PEL | KHV | FIN | 0.039 | 0.038 | 0.047 | $2.662e^{-4}$ | $5.218e^{-4}$ | 0.355 |
| PEL | KHV | GBR | 0.042 | 0.037 | 0.047 | $2.638e^{-4}$ | $5.171e^{-4}$ | 0.326 |
| PEL | KHV | IBS | 0.037 | 0.038 | 0.047 | $2.649e^{-4}$ | $5.192e^{-4}$ | 0.371 |
| PEL | KHV | TSI | 0.03 | 0.038 | 0.048 | $2.675e^{-4}$ | $5.242e^{-4}$ | 0.439 |
| PEL | PJL | CEU | 0.085 | 0.066 | 0.058 | $3.265e^{-4}$ | $6.399e^{-4}$ | 0.264 |
| PEL | PJL | FIN | 0.089 | 0.063 | 0.057 | $3.185e^{-4}$ | $6.242e^{-4}$ | 0.229 |
| PEL | PJL | GBR | 0.087 | 0.066 | 0.058 | $3.272e^{-4}$ | $6.413e^{-4}$ | 0.257 |
| PEL | PJL | IBS | 0.069 | 0.067 | 0.058 | $3.260e^{-4}$ | $6.389e^{-4}$ | 0.364 |
| PEL | PJL | TSI | 0.069 | 0.068 | 0.059 | $3.292e^{-4}$ | $6.452e^{-4}$ | 0.377 |
| PEL | STU | CEU | 0.053 | 0.062 | 0.058 | $3.238e^{-4}$ | $6.346e^{-4}$ | 0.446 |
| PEL | STU | FIN | 0.054 | 0.06 | 0.056 | $3.163e^{-4}$ | $6.200e^{-4}$ | 0.419 |
| PEL | STU | GBR | 0.054 | 0.062 | 0.058 | $3.235e^{-4}$ | $6.341e^{-4}$ | 0.438 |
| PEL | STU | IBS | 0.044 | 0.063 | 0.058 | $3.237e^{-4}$ | $6.345e^{-4}$ | 0.538 |
| PEL | STU | TSI | 0.044 | 0.064 | 0.058 | $3.266e^{-4}$ | $6.402e^{-4}$ | 0.546 |
| PUR | BEB | CEU | 0 | 0.005 | 0.008 | $4.467e^{-5}$ | $8.756e^{-5}$ | $> 0.999968$ |
| PUR | BEB | FIN | 0 | 0.007 | 0.01 | $5.783e^{-5}$ | $1.133e^{-4}$ | $> 0.999968$ |
| PUR | BEB | GBR | 0 | 0.005 | 0.008 | $4.551e^{-5}$ | $8.920e^{-5}$ | $> 0.999968$ |

Continued on next page

Table S52: (Continued)

| $A$ Pop. | $B$ Pop. | $C$ Pop. | Focal 72kb Region ($PBS_{A:B:C}$) | 72kb Non-overlapping Windows ($\mu$) | 72kb Non-overlapping Windows ($\sigma$) | 72kb Non-overlapping Windows ($SEM$) | 72kb Non-overlapping Windows ($\pm CI_{95\%}$) | $P - value$ |
|---|---|---|---|---|---|---|---|---|
| PUR | BEB | IBS | 0 | 0.004 | 0.007 | $3.820e^{-5}$ | $7.487e^{-5}$ | $> 0.999968$ |
| PUR | BEB | TSI | 0 | 0.004 | 0.008 | $4.304e^{-5}$ | $8.435e^{-5}$ | $> 0.999968$ |
| PUR | CDX | CEU | 0 | 0.004 | 0.009 | $5.022e^{-5}$ | $9.844e^{-5}$ | $> 0.999968$ |
| PUR | CDX | FIN | 0 | 0.007 | 0.014 | $7.728e^{-5}$ | $1.515e^{-4}$ | $> 0.999968$ |
| PUR | CDX | GBR | 0 | 0.004 | 0.009 | $5.203e^{-5}$ | $1.020e^{-4}$ | $> 0.999968$ |
| PUR | CDX | IBS | 0 | 0.003 | 0.008 | $4.253e^{-5}$ | $8.337e^{-5}$ | $> 0.999968$ |
| PUR | CDX | TSI | 0 | 0.003 | 0.008 | $4.663e^{-5}$ | $9.140e^{-5}$ | $> 0.999968$ |
| PUR | CHB | CEU | 0.002 | 0.004 | 0.009 | $5.018e^{-5}$ | $9.835e^{-5}$ | 0.286 |
| PUR | CHB | FIN | 0 | 0.008 | 0.014 | $7.781e^{-5}$ | $1.525e^{-4}$ | $> 0.999968$ |
| PUR | CHB | GBR | 0.002 | 0.004 | 0.009 | $5.181e^{-5}$ | $1.015e^{-4}$ | 0.296 |
| PUR | CHB | IBS | 0.006 | 0.003 | 0.008 | $4.214e^{-5}$ | $8.260e^{-5}$ | 0.166 |
| PUR | CHB | TSI | 0 | 0.003 | 0.008 | $4.634e^{-5}$ | $9.084e^{-5}$ | $> 0.999968$ |
| PUR | CHS | CEU | 0 | 0.004 | 0.009 | $5.036e^{-5}$ | $9.871e^{-5}$ | $> 0.999968$ |
| PUR | CHS | FIN | 0 | 0.007 | 0.014 | $7.775e^{-5}$ | $1.524e^{-4}$ | $> 0.999968$ |
| PUR | CHS | GBR | 0 | 0.004 | 0.009 | $5.197e^{-5}$ | $1.019e^{-4}$ | $> 0.999968$ |
| PUR | CHS | IBS | 0.004 | 0.003 | 0.008 | $4.234e^{-5}$ | $8.299e^{-5}$ | 0.204 |
| PUR | CHS | TSI | 0 | 0.003 | 0.008 | $4.654e^{-5}$ | $9.123e^{-5}$ | $> 0.999968$ |
| PUR | GIH | CEU | $2.108e^{-4}$ | 0.005 | 0.009 | $5.011e^{-5}$ | $9.822e^{-5}$ | 0.577 |
| PUR | GIH | FIN | 0 | 0.007 | 0.011 | $6.078e^{-5}$ | $1.191e^{-4}$ | $> 0.999968$ |
| PUR | GIH | GBR | 0 | 0.006 | 0.009 | $5.109e^{-5}$ | $1.001e^{-4}$ | $> 0.999968$ |

Continued on next page

Table S52: (Continued)

| $A$ Pop. | $B$ Pop. | $C$ Pop. | Focal 72kb Region ($PBS_{A:B:C}$) | 72kb Non-overlapping Windows ($\mu$) | 72kb Non-overlapping Windows ($\sigma$) | 72kb Non-overlapping Windows ($SEM$) | 72kb Non-overlapping Windows ($\pm CI_{95\%}$) | $P-value$ |
|---|---|---|---|---|---|---|---|---|
| PUR | GIH | IBS | 0 | 0.004 | 0.008 | $4.328e^{-5}$ | $8.483e^{-5}$ | $> 0.999968$ |
| PUR | GIH | TSI | 0 | 0.005 | 0.009 | $4.819e^{-5}$ | $9.446e^{-5}$ | $> 0.999968$ |
| PUR | ITU | CEU | 0 | 0.005 | 0.009 | $4.781e^{-5}$ | $9.371e^{-5}$ | $> 0.999968$ |
| PUR | ITU | FIN | 0 | 0.007 | 0.011 | $5.930e^{-5}$ | $1.162e^{-4}$ | $> 0.999968$ |
| PUR | ITU | GBR | 0 | 0.005 | 0.009 | $4.879e^{-5}$ | $9.564e^{-5}$ | $> 0.999968$ |
| PUR | ITU | IBS | 0 | 0.004 | 0.007 | $4.172e^{-5}$ | $8.177e^{-5}$ | $> 0.999968$ |
| PUR | ITU | TSI | 0 | 0.005 | 0.008 | $4.719e^{-5}$ | $9.249e^{-5}$ | $> 0.999968$ |
| PUR | JPT | CEU | 0 | 0.004 | 0.009 | $4.996e^{-5}$ | $9.793e^{-5}$ | $> 0.999968$ |
| PUR | JPT | FIN | 0 | 0.008 | 0.014 | $7.786e^{-5}$ | $1.526e^{-4}$ | $> 0.999968$ |
| PUR | JPT | GBR | 0 | 0.004 | 0.009 | $5.149e^{-5}$ | $1.009e^{-4}$ | $> 0.999968$ |
| PUR | JPT | IBS | 0.005 | 0.003 | 0.007 | $4.186e^{-5}$ | $8.205e^{-5}$ | 0.193 |
| PUR | JPT | TSI | 0 | 0.003 | 0.008 | $4.554e^{-5}$ | $8.926e^{-5}$ | $> 0.999968$ |
| PUR | KHV | CEU | 0 | 0.004 | 0.009 | $4.982e^{-5}$ | $9.764e^{-5}$ | $> 0.999968$ |
| PUR | KHV | FIN | 0 | 0.007 | 0.014 | $7.625e^{-5}$ | $1.495e^{-4}$ | $> 0.999968$ |
| PUR | KHV | GBR | 0 | 0.004 | 0.009 | $5.165e^{-5}$ | $1.012e^{-4}$ | $> 0.999968$ |
| PUR | KHV | IBS | 0 | 0.003 | 0.008 | $4.222e^{-5}$ | $8.275e^{-5}$ | $> 0.999968$ |
| PUR | KHV | TSI | 0 | 0.003 | 0.008 | $4.604e^{-5}$ | $9.024e^{-5}$ | $> 0.999968$ |
| PUR | PJL | CEU | 0.008 | 0.005 | 0.008 | $4.762e^{-5}$ | $9.334e^{-5}$ | 0.23 |
| PUR | PJL | FIN | 0.008 | 0.007 | 0.01 | $5.682e^{-5}$ | $1.114e^{-4}$ | 0.316 |
| PUR | PJL | GBR | 0.007 | 0.005 | 0.009 | $4.869e^{-5}$ | $9.543e^{-5}$ | 0.267 |

Continued on next page

Table S52: (Continued)

| $A$ Pop. | $B$ Pop. | $C$ Pop. | Focal 72kb Region ($PBS_{A:B:C}$) | 72kb Non-overlapping Windows ($\mu$) | 72kb Non-overlapping Windows ($\sigma$) | 72kb Non-overlapping Windows ($SEM$) | 72kb Non-overlapping Windows ($\pm CI_{95\%}$) | $P-value$ |
|---|---|---|---|---|---|---|---|---|
| PUR | PJL | IBS | 0.002 | 0.004 | 0.007 | $4.131e^{-5}$ | $8.098e^{-5}$ | 0.455 |
| PUR | PJL | TSI | 0.002 | 0.005 | 0.008 | $4.635e^{-5}$ | $9.084e^{-5}$ | 0.476 |
| PUR | STU | CEU | 0 | 0.005 | 0.008 | $4.750e^{-5}$ | $9.311e^{-5}$ | $> 0.999968$ |
| PUR | STU | FIN | 0 | 0.007 | 0.011 | $5.932e^{-5}$ | $1.163e^{-4}$ | $> 0.999968$ |
| PUR | STU | GBR | 0 | 0.005 | 0.009 | $4.843e^{-5}$ | $9.493e^{-5}$ | $> 0.999968$ |
| PUR | STU | IBS | 0 | 0.004 | 0.007 | $4.106e^{-5}$ | $8.048e^{-5}$ | $> 0.999968$ |
| PUR | STU | TSI | 0 | 0.005 | 0.008 | $4.643e^{-5}$ | $9.101e^{-5}$ | $> 0.999968$ |

**Table S53. Summary of per-SNP $PBS_{MXL:CHB:CEU}$ values for the focal 742kb *MUC19* region.**
The total number of SNPs, the number of SNPs with a $PBS_{MXL:CHB:CEU}$ value greater than the $99.95^{th}$ genome-wide percentile, and those identified as significant based on different statistical criteria: uncorrected *P-values* less than 0.05, Bonferroni adjusted *P-values* less than 0.05, and Benjamini-Hochberg adjusted *P-values* less than 0.01. We report the results for all SNPs, as well as stratified by SNP set (see Methods for a description of these SNP sets).

Table S53:

| SNP Set | Total SNPs | $PBS > 99.95^{th}$ Genome-Wide Percentile | Uncorrected $P-values < 0.05$ | Bonferroni Adjusted $P-values < 0.05$ | Benjamini-Hochberg Adjusted $P-values < 0.01$ |
|---|---|---|---|---|---|
| All SNPs | 6144 | 417 | 1111 | 0 | 485 |
| Archaic SNPs | 217 | 208 | 217 | 0 | 217 |
| Denisovan-specific SNPs | 135 | 135 | 135 | 0 | 135 |
| Neanderthal-specific SNPs | 80 | 72 | 80 | 0 | 80 |
| Human-specific SNPs | 5420 | 141 | 677 | 0 | 200 |
| Shared Hominin SNPs | 507 | 68 | 217 | 0 | 68 |

**Table S54. Summary of per-SNP $PBS_{MXL:CHB:CEU}$ outliers for the focal 742kb *MUC19* region.** The total number of SNPs, the number of outlier SNPs with a $PBS_{MXL:CHB}$ value greater than the $99.95^{th}$ genome-wide percentile, the mean $PBS_{MXL:CHB}$ value among these outliers, and the associated 95% confidence intervals ($\pm CI_{95\%}$) for the mean $PBS_{MXL:CHB}$ value. Additionally, the mean percentile rank of outlier SNPs and the associated 95% confidence intervals ($\pm CI_{95\%}$) for the mean percentile rank is reported. We report the results for all SNPs, as well as stratified by SNP set (see Methods for a description of these SNP sets).

Table S54:

| SNP Set | Total SNPs | $PBS > 99.95^{th}$ Genome-Wide Percentile | $PBS$ ($\mu$) | $PBS$ ($\pm CI_{95\%}$) | Percentile Rank ($\mu$) | Percentile Rank ($\pm CI_{95\%}$) |
|---|---|---|---|---|---|---|
| All SNPs | 6144 | 417 | 0.302 | 0.003 | 99.975 | 0.001 |
| Archaic SNPs | 217 | 208 | 0.293 | 0.004 | 99.972 | 0.001 |
| Denisovan-specific SNPs | 135 | 135 | 0.279 | $3.191e^{-4}$ | 99.967 | $2.287e^{-4}$ |
| Neanderthal-specific SNPs | 80 | 72 | 0.32 | 0.007 | 99.983 | 0.002 |
| Human-specific SNPs | 5420 | 141 | 0.313 | 0.007 | 99.979 | 0.002 |
| Shared Hominin SNPs | 507 | 68 | 0.305 | 0.006 | 99.978 | 0.002 |

**Table S55. Summary of significant per-SNP $PBS_{MXL:CHB:CEU}$ values for the focal 742kb *MUC19* region.** The total number of SNPs, the number of significant SNPs (i.e., SNPs with a Benjamini-Hochberg adjusted *P-values* less than 0.01), the mean $PBS_{MXL:CHB}$ value among these significant SNPs, and the associated 95% confidence intervals ($\pm CI_{95\%}$) for the mean $PBS_{MXL:CHB}$ value. Additionally, the mean percentile rank of significant SNPs and the associated 95% confidence intervals ($\pm CI_{95\%}$) for the mean percentile rank is reported. We report the results for all SNPs, as well as stratified by SNP set (see Methods for a description of these SNP sets).

Table S55:

| SNP Set | Total SNPs | Benjamini-Hochberg Adjusted $P - values <$ 0.01 | $PBS$ ($\mu$) | $PBS$ ($\pm CI_{95\%}$) | Percentile Rank ($\mu$) | Percentile Rank ($\pm CI_{95\%}$) |
|---|---|---|---|---|---|---|
| All SNPs | 6144 | 485 | 0.293 | 0.003 | 99.969 | 0.002 |
| Archaic SNPs | 217 | 217 | 0.291 | 0.004 | 99.97 | 0.002 |
| Denisovan-specific SNPs | 135 | 135 | 0.279 | $3.191e^{-4}$ | 99.967 | $2.287e^{-4}$ |
| Neanderthal-specific SNPs | 80 | 80 | 0.312 | 0.009 | 99.977 | 0.004 |
| Human-specific SNPs | 5420 | 200 | 0.291 | 0.007 | 99.964 | 0.004 |
| Shared Hominin SNPs | 507 | 68 | 0.305 | 0.006 | 99.978 | 0.002 |

**Table S56. Demographic model from Medina-Muñoz et al., 2023 with archaic introgression.** Demographic parameters used for *PBS* simulations.

Table S56:

| Parameter | Value | Description | Reference |
|---|---|---|---|
| Population size | 13580 | Ancestral Pop. size | Medina-Muñoz et al., 2023 |
| Population size | 13249 | Archaic Pop. size | Jacobs et al., 2019; Malaspinas et al., 2016 |
| Population size | 27142 | AMH Pop. size | Medina-Muñoz et al., 2023 |
| Population size | 826 | NEA Pop. size | Jacobs et al., 2019; Malaspinas et al., 2016 |
| Population size | 5083 | DEN Pop. size | Jacobs et al., 2019; Malaspinas et al., 2016 |
| Population size | 1835 | OOA Pop. size | Medina-Muñoz et al., 2023 |
| Population size | 27142 | AFR Pop. size | Medina-Muñoz et al., 2023 |
| Population size | 2761 | EUR Pop. size | Medina-Muñoz et al., 2023 |
| Population size | 1955 | EAS Pop. size | Medina-Muñoz et al., 2023 |
| Population size | 1313 | MXB Pop. size | Medina-Muñoz et al., 2023 |
| Population size | 20000 | MXL Pop. size | Medina-Muñoz et al., 2023 |
| Growth rate per generation | 0.001 | EUR Pop. growth rate | Medina-Muñoz et al., 2023 |
| Growth rate per generation | 0.001 | EAS Pop. growth rate | Medina-Muñoz et al., 2023 |
| Growth rate per generation | 0.003 | MXB Pop. growth rate | Medina-Muñoz et al., 2023 |
| Migration rate per generation | $1.586e^{-4}$ | AFR-OOA migration rate | Medina-Muñoz et al., 2023 |
| Migration rate per generation | $2.502e^{-5}$ | AFR-EUR migration rate | Medina-Muñoz et al., 2023 |
| Migration rate per generation | $3.295e^{-6}$ | AFR-EAS migration rate | Medina-Muñoz et al., 2023 |

Continued on next page

Table S56: (Continued)

| Parameter | Value | Description | Reference |
|---|---|---|---|
| Migration rate per generation | $6.624e^{-5}$ | EUR-EAS migration rate | Medina-Muñoz et al., 2023 |
| Migration rate per generation | 0.05 | MXB-MXL migration rate | Medina-Muñoz et al., 2023 |
| Migration rate per generation | 0.058 | EUR-MXL migration rate | Medina-Muñoz et al., 2023 |
| Admixture percentage | 0.05 | NEA admixture into OOA | Green et al., 2010 |
| Admixture percentage | 0.05 | DEN admixture into EAS | Skoglund and Jakobsson 2011 |
| Admixture percentage | 0.112 | AFR admixture into MXL | Medina-Muñoz et al., 2023 |
| Admixture percentage | 0.007 | EAS admixture into MXL | Medina-Muñoz et al., 2023 |
| Epoch time (g. a.) | 20225 | Ancestral-Archaic split | Jacobs et al., 2019; Malaspinas et al., 2016 |
| Epoch time (g. a.) | 16671 | Ancestral-AMH split | Medina-Muñoz et al., 2023 |
| Epoch time (g. a.) | 15090 | Archaic-DEN split | Jacobs et al., 2019; Malaspinas et al., 2016 |
| Epoch time (g. a.) | 15090 | Archaic-NEA split | Jacobs et al., 2019; Malaspinas et al., 2016 |
| Epoch time (g. a.) | 3046 | AMH-OoA split | Medina-Muñoz et al., 2023 |
| Epoch time (g. a.) | 3046 | AMH-AFR split | Medina-Muñoz et al., 2023 |
| Epoch time (g. a.) | 2069 | NEA admixture into OOA | Green et al., 2010; Prüfer et al., 2014; Prüfer et al., 2017 |
| Epoch time (g. a.) | 1764 | OOA-EUR split | Medina-Muñoz et al., 2023 |
| Epoch time (g. a.) | 1764 | OOA-EAS split | Medina-Muñoz et al., 2023 |
| Epoch time (g. a.) | 1552 | DEN admixture into EAS | Reich et al., 2011 |
| Epoch time (g. a.) | 1123 | EAS-MXB split | Medina-Muñoz et al., 2023 |

Continued on next page

Table S56: (Continued)

| Parameter | Value | Description | Reference |
| --- | --- | --- | --- |
| Epoch time (g. a.) | 16 | MXL divergence from EUR and MXB | Medina-Muñoz et al., 2023 |
| Epoch time (g. a.) | 13 | AFR and EAS admixture into MXL | Medina-Muñoz et al., 2023 |
| Mutation rate | $1.315e^{-8}$ | Per-base per-generation mutation rate | Hubisz and Siepel 2020 |
| Uniform recombination rate | $6.000e^{-9}$ | Per-base per-generation recombination rate | Hinch et al., 2011 |
| MXL sample size | 64 | MXL sample size | Based on the number of individuals in MXL from 1000 Genomes Project |
| EAS sample size | 103 | EAS sample size | Based on the number of individuals in CHB from 1000 Genomes Project |
| EUR sample size | 99 | EUR sample size | Based on the number of individuals in CEU from 1000 Genomes Project |
| Generation time | 29 | Number of years per generation | Fenner 2005 |

**Table S57. Summary of SNPs segregating at high frequency in MXL across 10,000 simulated replicates.** The total number of SNPs, the number of SNPs segregating a frequency greater than or equal to 30%, and the associated *P-value* based on 10,000 replicates per simulation model. The *P-value* represents the proportion of SNPs across all 10,000 simulations that are segregating at a frequency greater than or equal to 30%, where a *P-value* less than 0.05 is considered statistically significant. We report the results for all SNPs, as well as archaic SNPs.

Table S57:

| Selection Model | SNP Set | Total SNPs | MXL SNPs $\geq$ 30% | $P - value$ |
|---|---|---|---|---|
| Neutral | All SNPs | 55296617 | 7572328 | 0.137 |
| Neutral | Archaic SNPs | 55296617 | 47581 | $8.605e^{-4}$ |
| Negative | All SNPs | 53988190 | 6910329 | 0.128 |
| Negative | Archaic SNPs | 53988190 | 65682 | 0.001 |

**Table S58. Summary of significant per-SNP $PBS_{MXL:CHB:CEU}$ values across 10,000 simulated replicates for all 417 outlier SNPs.**
The number of $PBS_{MXL:CHB:CEU}$ values among the 417 SNPs with a per-SNP *PBS* value greater than the empirical $99.95^{th}$ genome-wide percentile, and those identified as significant based on 10,000 replicate simulations per model and different statistical criteria: uncorrected *P-values* less than 0.05, Bonferroni adjusted *P-values* less than 0.05, and Benjamini-Hochberg adjusted *P-values* less than 0.01. We report the results for all SNPs, as well as archaic SNPs.

Table S58:

| Selection Model | SNP Set | Number of $PBS_{MXL:CHB:CEU}$ Values | Uncorrected $P-value < 0.05$ | Bonferroni Adj. $P-value < 0.05$ | Benjamini-Hochberg Adj. $P-value < 0.01$ |
|---|---|---|---|---|---|
| Negative | All SNPs | 417 | 417 | 184 | 417 |
| Negative | Archaic SNPs | 208 | 208 | 208 | 208 |
| Neutral | All SNPs | 417 | 417 | 97 | 417 |
| Neutral | Archaic SNPs | 208 | 208 | 208 | 208 |

**Table S59. Summary of significant per-SNP $PBS_{MXL:CHB:CEU}$ values across 10,000 simulated replicates for the 57 unique outlier SNPs.**
The number of unique $PBS_{MXL:CHB:CEU}$ values among the 57 unique per-SNP $PBS$ value greater than the empirical $99.95^{th}$ genome-wide percentile, and those identified as significant based on 10,000 replicate simulations per model and different statistical criteria: uncorrected *P-values* less than 0.05, Bonferroni adjusted *P-values* less than 0.05, and Benjamini-Hochberg adjusted *P-values* less than 0.01. We report the results for all SNPs, as well as archaic SNPs.

Table S59:

| Selection Model | SNP Set | Unique $PBS_{MXL:CHB:CEU}$ Values | Uncorrected $P-value < 0.05$ | Bonferroni Adj. $P-value < 0.05$ | Benjamini-Hochberg Adj. $P-value < 0.01$ |
|---|---|---|---|---|---|
| Negative | All SNPs | 57 | 57 | 57 | 57 |
| Negative | Archaic SNPs | 20 | 20 | 20 | 20 |
| Neutral | All SNPs | 57 | 57 | 57 | 57 |
| Neutral | Archaic SNPs | 20 | 20 | 20 | 20 |

**Table S60. Comparison of the number of SNPs segregating at high frequency in MXL for the observed focal regions and across 10,000 simulated replicates.**
The number of SNPs segregating a frequency greater than or equal to 30% observed in MXL, along with the number of simulated replicates where the number SNPs segregating a frequency greater than or equal to 30% is greater than or equal to what is observed in the empirical data. The *P-value* represents the proportion of the 10,000 simulation replicates where the simulated value is greater than or equal to what is observed in the empirical data, where a *P-value* less than 0.05 is considered statistically significant. We report the results for both the focal 742kb and 72kb *MUC19* regions, for all SNPs, as well as archaic SNPs.

Table S60:

| Selection Model | SNP Set | Region | MXL AAF $\geq$ 30% (Observed) | Simulated Replicates $\geq$ Observed | $P - value$ |
|---|---|---|---|---|---|
| Neutral | All SNPs | 742kb | 1311 | 0 | $< 0.0001$ |
| Neutral | Archaic SNPs | 742kb | 208 | 0 | $< 0.0001$ |
| Neutral | All SNPs | 72kb | 300 | 0 | $< 0.0001$ |
| Neutral | Archaic SNPs | 72kb | 140 | 0 | $< 0.0001$ |
| Negative | All SNPs | 742kb | 1311 | 0 | $< 0.0001$ |
| Negative | Archaic SNPs | 742kb | 208 | 0 | $< 0.0001$ |
| Negative | All SNPs | 72kb | 300 | 0 | $< 0.0001$ |
| Negative | Archaic SNPs | 72kb | 140 | 0 | $< 0.0001$ |

**Table S61. Comparison of per-region *PBS* values for the observed focal regions and across 10,000 simulated replicates.** The per-region *PBS* values observed in MXL, along with the number of simulated replicates where the per-region *PBS* value is greater than or equal to what is observed in the empirical data. The *P-value* represents the proportion of the 10,000 simulation replicates where the simulated value is greater than or equal to what is observed in the empirical data, where a *P-value* less than 0.05 is considered statistically significant. We report the results for both the focal 742kb and 72kb *MUC19* regions, for all SNPs, as well as archaic SNPs.

Table S61:

| Selection Model | SNP Set | Region | $PBS$ (Observed) | Simulated Replicates $\geq$ Observed | $P - value$ |
|---|---|---|---|---|---|
| Neutral | All SNPs | 742kb | 0.066 | 0 | $< 0.0001$ |
| Neutral | Archaic SNPs | 742kb | 0.291 | 0 | $< 0.0001$ |
| Neutral | All SNPs | 72kb | 0.127 | 3 | 0.0003 |
| Neutral | Archaic SNPs | 72kb | 0.28 | 6 | 0.0006 |
| Negative | All SNPs | 742kb | 0.066 | 0 | $< 0.0001$ |
| Negative | Archaic SNPs | 742kb | 0.291 | 0 | $< 0.0001$ |
| Negative | All SNPs | 72kb | 0.127 | 1 | 0.0001 |
| Negative | Archaic SNPs | 72kb | 0.28 | 1 | 0.0001 |

**Table S62. Power to detect positive selection in MXL using per-region *PBS* values.**
The power to detect positive selection for different strengths of positive selection: strong ($s = 0.1$), weak ($s = 0.01$), and extremely weak ($s = 0.0015$). The power represents the proportion of the 1,000 simulated replicates with per-region *PBS* values greater than the upper 5% quantile from its respective neutral distribution of per-region *PBS* values consisting of 10,000 replicates. For each selection coefficient we report the results for both the simulated 742kb and 72kb *MUC19* regions, for all SNPs, as well as archaic SNPs.

Table S62:

| $s$ | All SNPs (72kb) | Archaic SNPs (72kb) | All SNPs (742kb) | Archaic SNPs (742kb) |
|---|---|---|---|---|
| 0.1 | 0.986 | 0.984 | 0.982 | 0.981 |
| 0.01 | 0.873 | 0.935 | 0.794 | 0.857 |
| 0.0015 | 0.096 | 0.193 | 0.081 | 0.126 |

**Table S63. Integrated Haplotype Scores (*iHS*) for archaic SNPs in the focal 742kb *MUC19* region among 1000 Genomes Project non-African populations.**
The normalized $| iHS |$ scores for archaic SNPs observed at the focal 742kb region for each non-African population stratified by super population: Admixed Americans (AMR), South Asians (SAS), East Asians (EAS), and Europeans (EUR). For each population, the normalized *iHS* scores were calculated for archaic SNPs with a minor allele frequency $> 5\%$. For each population, the table reports the total number of *iHS* scores observed for archaic SNPs, the number of extreme *iHS* scores for archaic SNPs (i.e., $| iHS |> 2$), the observed proportion of extreme *iHS* scores for archaic SNPs (calculated as the number of extreme *iHS* scores normalized by the total number of *iHS* scores), the $99^{th}$ percentile of the proportion of extreme *iHS* scores for archaic SNPs determined from the genomic background distribution (i.e., non-overlapping 742kb windows with more than 10 archaic SNPs), the total number of 742kb windows in the genomic background distribution, the number of 742kb windows where the observed proportion of extreme *iHS* scores for archaic SNPs is greater than the genomic background distribution, and the percentile rank of the observed proportion of extreme *iHS* scores for archaic SNPs (i.e., the percentage of the genomic background distribution that is less than the observed proportion of extreme *iHS* scores). An observed proportion of extreme *iHS* scores greater than the $99^{th}$ percentile of the genomic background distribution is considered evidence of positive selection. Note that a ”—” denotes that the population did not meet the minimum number of archaic SNPs threshold for the *iHS* analysis.

Table S63:

| Super Population | Population | Focal 742kb Region (Total SNPs) | Focal 742kb Region (SNPs with $\| iHS \|> 2$) | Focal 742kb Region (Prop. of SNPs with $\| iHS \|> 2$) | 742kb Non-overlapping Windows ($99^{th}$ Percentile) | 742kb Non-overlapping Windows (Total SNPs $> 10$) | Focal 742kb Region > 742kb Non-overlapping Windows | Focal 742kb Region (Percentile Rank) |
|---|---|---|---|---|---|---|---|---|
| AMR | MXL | 217 | 208 | 0.959 | 0.733 | 1250 | 1248 | 99.84 |
| AMR | PEL | 217 | 66 | 0.304 | 0.687 | 1119 | 1063 | 94.996 |
| AMR | CLM | 217 | 6 | 0.028 | 0.52 | 1276 | 1032 | 80.878 |
| AMR | PUR | 217 | 0 | 0 | 0.562 | 1148 | 0 | 0 |
| SAS | BEB | 197 | 0 | 0 | 0.514 | 1263 | 0 | 0 |
| SAS | STU | 214 | 1 | 0.005 | 0.494 | 1264 | 908 | 71.835 |
| SAS | ITU | 184 | 0 | 0 | 0.545 | 1269 | 0 | 0 |

Continued on next page

Table S63: (Continued)

| Super Population | Population | Focal 742kb Region (Total SNPs) | Focal 742kb Region (SNPs with $\mid iHS \mid > 2$) | Focal 742kb Region (Prop. of SNPs with $\mid iHS \mid > 2$) | 742kb Non-overlapping Windows ($99^{th}$ Percentile) | 742kb Non-overlapping Windows (Total SNPs > 10) | Focal 742kb Region > 742kb Non-overlapping Windows | Focal 742kb Region (Percentile Rank) |
|---|---|---|---|---|---|---|---|---|
| SAS | PJL | 156 | 0 | 0 | 0.494 | 1312 | 0 | 0 |
| SAS | GIH | 214 | 0 | 0 | 0.651 | 1288 | 0 | 0 |
| EAS | CHB | 0 | 0 | — | — | — | — | — |
| EAS | KHV | 171 | 0 | 0 | 0.458 | 1213 | 0 | 0 |
| EAS | CHS | 0 | 0 | — | — | — | — | — |
| EAS | JPT | 2 | 0 | 0 | — | — | — | — |
| EAS | CDX | 191 | 0 | 0 | 0.459 | 1198 | 0 | 0 |
| EUR | TSI | 152 | 1 | 0.007 | 0.563 | 1254 | 932 | 74.322 |
| EUR | CEU | 17 | 0 | 0 | 0.429 | 1281 | 0 | 0 |
| EUR | IBS | 17 | 1 | 0.059 | 0.587 | 1232 | 1057 | 85.795 |
| EUR | GBR | 0 | 0 | — | — | — | — | — |
| EUR | FIN | 17 | 0 | 0 | 0.498 | 1321 | 0 | 0 |

**Table S64. Integrated Haplotype Scores (*iHS*) for archaic SNPs in the focal 72kb *MUC19* region among 1000 Genomes Project non-African populations.**
The normalized $| iHS |$ scores for archaic SNPs observed at the focal 72kb region for each non-African population stratified by super population: Admixed Americans (AMR), South Asians (SAS), East Asians (EAS), and Europeans (EUR). For each population, the normalized *iHS* scores were calculated for archaic SNPs with a minor allele frequency $> 5\%$. For each population, the table reports the total number of *iHS* scores observed for archaic SNPs, the number of extreme *iHS* scores for archaic SNPs (i.e., $| iHS |> 2$), the observed proportion of extreme *iHS* scores for archaic SNPs (calculated as the number of extreme *iHS* scores normalized by the total number of *iHS* scores), the $99^{th}$ percentile of the proportion of extreme *iHS* scores for archaic SNPs determined from the genomic background distribution (i.e., non-overlapping 72kb windows with more than 10 archaic SNPs), the total number of 72kb windows in the genomic background distribution, the number of 72kb windows where the observed proportion of extreme *iHS* scores for archaic SNPs is greater than the genomic background distribution, and the percentile rank of the observed proportion of extreme *iHS* scores for archaic SNPs (i.e., the percentage of the genomic background distribution that is less than the observed proportion of extreme *iHS* scores). An observed proportion of extreme *iHS* scores greater than the $99^{th}$ percentile of the genomic background distribution is considered evidence of positive selection. Note that a ”—” denotes that the population did not meet the minimum number of archaic SNPs threshold for the *iHS* analysis.

Table S64:

| Super Population | Population | Focal 72kb Region (Total SNPs) | Focal 72kb Region (SNPs with $\| iHS \|> 2$) | Focal 72kb Region (Prop. of SNPs with $\| iHS \|> 2$) | 72kb Non-overlapping Windows ($99^{th}$ Percentile) | 72kb Non-overlapping Windows (Total SNPs $> 10$) | Focal 72kb Region > 72kb Non-overlapping Windows | Focal 72kb Region (Percentile Rank) |
|---|---|---|---|---|---|---|---|---|
| AMR | MXL | 140 | 135 | 0.964 | 0.938 | 2699 | 2677 | 99.185 |
| AMR | PEL | 140 | 58 | 0.414 | 1 | 2332 | 2225 | 95.412 |
| AMR | CLM | 140 | 6 | 0.043 | 0.824 | 2630 | 2339 | 88.935 |
| AMR | PUR | 140 | 0 | 0 | 0.878 | 2344 | 0 | 0 |
| SAS | BEB | 140 | 0 | 0 | 0.732 | 3048 | 0 | 0 |
| SAS | STU | 140 | 0 | 0 | 0.733 | 2901 | 0 | 0 |
| SAS | ITU | 140 | 0 | 0 | 0.733 | 2900 | 0 | 0 |

Continued on next page

Table S64: (Continued)

| Super Population | Population | Focal 72kb Region (Total SNPs) | Focal 72kb Region (SNPs with $\mid iHS \mid > 2$) | Focal 72kb Region (Prop. of SNPs with $\mid iHS \mid > 2$) | 72kb Non-overlapping Windows ($99^{th}$ Percentile) | 72kb Non-overlapping Windows (Total SNPs > 10) | Focal 72kb Region > 72kb Non-overlapping Windows | Focal 72kb Region (Percentile Rank) |
|---|---|---|---|---|---|---|---|---|
| SAS | PJL | 139 | 0 | 0 | 0.802 | 2952 | 0 | 0 |
| SAS | GIH | 140 | 0 | 0 | 0.848 | 2883 | 0 | 0 |
| EAS | CHB | 0 | 0 | — | — | — | — | — |
| EAS | KHV | 140 | 0 | 0 | 0.686 | 3248 | 0 | 0 |
| EAS | CHS | 0 | 0 | — | — | — | — | — |
| EAS | JPT | 2 | 0 | 0 | — | — | — | — |
| EAS | CDX | 140 | 0 | 0 | 0.799 | 3116 | 0 | 0 |
| EUR | TSI | 139 | 1 | 0.007 | 0.766 | 2668 | 2352 | 88.156 |
| EUR | CEU | 0 | 0 | — | — | — | — | — |
| EUR | IBS | 0 | 0 | — | — | — | — | — |
| EUR | GBR | 0 | 0 | — | — | — | — | — |
| EUR | FIN | 0 | 0 | — | — | — | — | — |

**Table S65. Read-based phasing information for the Chagyrskaya Neanderthal.**
The four heterozygous sites that could not be resolved with statistical phasing, preceding and proceeding heterozygous sites and their distance from the heterozygous site to resolve, and the results from manual read-based phasing for the Chagyrskaya Neanderthal at the focal 72kb *MUC19* region.

Table S65:

| Het. Pos. to Resolve | Preceding Het. Pos. | Dist. to Preceding Het. Pos. | Proceeding Het. Pos. | Dist. to Proceeding Het. Pos. | Is Resvolable? | Phasing Info. |
|---|---|---|---|---|---|---|
| 40767590 | 40761946 | 5644 | 40768905 | 1315 | False | Distant Flanking Het. Sites |
| 40821807 | 40821795 | 12 | 40821847 | 40 | True | G-C-C and A-T-A |
| 40826155 | 40826138 | 17 | 40826201 | 46 | True | T-A-G and G-G-A |
| 40828121 | 40828019 | 102 | 40828306 | 185 | False | Not Enough Overlapping Reads |

**Table S66. Read-based phasing information for the Vindija Neanderthal.**
The five heterozygous sites that could not be resolved with statistical phasing, preceding and proceeding heterozygous sites and their distance from the heterozygous site to resolve, and the results from manual read-based phasing for the Vindija Neanderthal at the focal 72kb *MUC19* region. Note that position 40829306 is the last heterozygous site in the focal 72kb *MUC19* region for the Vindija Neanderthal, and thus, the proceeding heterozygous site information is denoted by a "-1".

Table S66:

| Het. Pos. to Resolve | Preceding Het. Pos. | Dist. to Preceding Het. Pos. | Proceeding Het. Pos. | Dist. to Proceeding Het. Pos. | Is Resvolable? | Phasing Info. |
|---|---|---|---|---|---|---|
| 40765370 | 40761946 | 3424 | 40768905 | 3535 | False | Distant Flanking Het. Sites |
| 40783344 | 40775127 | 8217 | 40784418 | 1074 | False | Distant Flanking Het. Sites |
| 40821807 | 40821795 | 12 | 40821847 | 40 | True | G-C-C and A-T-A |
| 40824215 | 40824154 | 61 | 40824266 | 51 | True | G-A and A-C |
| 40829306 | 40828306 | 1000 | -1 | -1 | False | Distant Flanking Het. Sites |

**Table S67. Patterns of pairwise differences at the focal 72kb *MUC19* region between the *Denisovan-like* haplotypes in 1000 Genomes Project and the archaic individuals**

The mean ($\mu$), standard deviation ($\sigma$), standard error of the mean ($SEM$), and 95% confidence intervals ($\pm CI_{95\%}$) of the number of pairwise distance distribution for the focal 72kb *MUC19* region between the 287 *Denisovan-like* haplotypes in 1000 Genomes Project and the four high-coverage archaic individuals. For the Denisovan and Altai Neanderthal, we compute the number of pairwise differences between a modern human haplotype and the two unphased archaic chromosomes, and for the Chagyrskaya and Vindija Neanderthals, we compute the number of pairwise differences between a modern human haplotype and a phased archaic haplotype.

Table S67:

| Archaic | Focal 72kb Region Pairwise Diffs. ($\mu$) | Focal 72kb Region Pairwise Diffs. ($\sigma$) | Focal 72kb Region Pairwise Diffs. ($SEM$) | Focal 72kb Region Pairwise Diffs. ($\pm CI_{95\%}$) |
|---|---|---|---|---|
| Denisovan | 49.01 | 1.994 | 0.118 | 0.232 |
| Altai Nean. | 179.838 | 2.503 | 0.148 | 0.291 |
| Chagyrskaya Nean. Hap. 1 | 170.338 | 2.457 | 0.145 | 0.286 |
| Chagyrskaya Nean. Hap. 2 | 7.547 | 2.84 | 0.168 | 0.331 |
| Vindija Nean. Hap. 1 | 170.899 | 2.179 | 0.129 | 0.254 |
| Vindija Nean. Hap. 2 | 6.108 | 2.612 | 0.154 | 0.304 |

**Dataset 1. Sequence divergence between the haplotypes in the 1000 Genomes Project (1KG) and the four archaic individuals at the focal 72kb *MUC19* region.**
The observed sequence divergence—i.e., the number of pairwise differences between a modern human haplotype and an archaic genotype normalized by the effective sequence length—between all 1KG haplotypes and the four unphased high-coverage archaic individuals at the 72kb region. For each comparison, the mean ($\mu$), standard deviation ($\sigma$), standard error of the mean ($SEM$), and 95% confidence intervals ($\pm CI_{95\%}$) of the sequence divergence genomic background distribution used to compute the *P-value* are also reported. The *P-value* represents the proportion of non-overlapping 72kb windows of comparable effective sequence length where the sequence divergence is less than or equal to that observed at the focal 72kb region. After correcting for two multiple comparisons—i.e., one per haplotype—a *P-value* less than 0.025 is considered statistically significant. Note that at the focal 72kb region, the effective sequence length with respect to the 1KG is 48435bp, 48654bp, 48121bp, and 48447bp for the Denisovan, Altai Neanderthal, Chagyrskaya Neanderthal, and Vindija Neanderthal, respectively.

**Dataset 2. Sequence divergence between the haplotypes in the 1000 Genomes Project (1KG) and the four archaic individuals at the focal 742kb *MUC19* region.**
The observed sequence divergence—i.e., the number of pairwise differences between a modern human haplotype and an archaic genotype normalized by the effective sequence length—between all 1KG haplotypes and the four unphased high-coverage archaic individuals at the 742kb region. For each comparison, the mean ($\mu$), standard deviation ($\sigma$), standard error of the mean ($SEM$), and 95% confidence intervals ($\pm CI_{95\%}$) of the sequence divergence genomic background distribution used to compute the *P-value* are also reported. The *P-value* represents the proportion of non-overlapping 742kb windows of comparable effective sequence length where the sequence divergence is less than or equal to that observed at the focal 742kb region. After correcting for two multiple comparisons—i.e., one per haplotype—a *P-value* less than 0.025 is considered statistically significant. Note that at the focal 742kb region, the effective sequence length with respect to the 1KG is 48435bp, 48654bp, 48121bp, and 48447bp for the Denisovan, Altai Neanderthal, Chagyrskaya Neanderthal, and Vindija Neanderthal, respectively.

**Dataset 3. Sequence divergence between the haplotypes in the 1000 Genomes Project (1KG) and the phased late Neanderthal haplotypes at the focal 72kb *MUC19* region.**
The observed sequence divergence—i.e., the number of pairwise differences between a modern human haplotype and a late Neanderthal haplotype normalized by the effective sequence length—between all 1KG haplotypes and the phased haplotypes for both of the late Neanderthals. To assess the significance, we built a distribution of sequence divergence between each of the modern human's two chromosomes and the late Neanderthal's pseudo-haplotype, which was generated by randomly sampling one allele at each position from the unphased late Neanderthal's genotypes. For each late Neanderthal, the mean ($\mu$), standard deviation ($\sigma$), standard error of the mean ($SEM$), and 95% confidence intervals ($\pm CI_{95\%}$) of the pseudo-haplotype divergence genomic background distribution used to compute the *P-value* are also reported. The *P-value* represents the proportion of non-overlapping 72kb windows of comparable effective sequence length where the sequence divergence is less than or equal to that observed at the focal 72kb region. After correcting for four multiple comparisons—i.e., two per each modern human haplotype—a *P-value* less than 0.0125 is considered statistically significant. Note that the *Denisovan-like* in each of the late Neanderthal's corresponds to their second haplotype (i.e., Chagyrskaya Nean. Hap. 2 and Vindija Nean. Hap. 2). Also, note that at the phased 72kb region, the effective sequence length with respect to the 1KG is 48119bp and 48444bp for the Chagyrskaya and Vindija Neanderthal, respectively.

**Dataset 4. Validation of the 135 Denisovan-specific alleles within the 72kb *MUC19* region using high-coverage long-read genomes from the Human Genome Structural Variation Consortium (HGSVC) and the 1000 Genomes Project Oxford Nanopore Technologies Sequencing Consortium (1KG-ONT).**
For each of the 135 Denisovan-specific alleles in the 72kb *MUC19* region, we report the genomic position and reference allele for both the Hg19 and Hg38 reference genomes. Variants from the HGSVC individuals (individual-population: HG00864-CDX, HG03009-BEB, NA20847-GIH) were called using the Phased Assembly Variant caller (PAV), resulting in phased variant calls that confirm the presence of all 135 Denisovan alleles on the same haploid genome copy. For the 1KG-ONT individuals (individual-population: HG01122-CLM, HG02252-PEL, HG02262-PEL), we validated the presence of these Denisovan alleles using variant calls from both Clair3 and PEPPER-Margin-DeepVariant (PMDV), resulting in two sets of unphased variant calls per individual. Note that the PEL individual HG02262 carries two copies of the *Denisovan-like* haplotype in the 72kb *MUC19* region. This individual is always homozygous for the Denisovan allele, which confirms its presence across both haploid genome copies. Regardless of the dataset (HGSVC or 1KG-ONT) or variant caller (PAV, Clair3, or PMDV), the Denisovan allele is always the alternative allele with respect to the Hg38 reference genome, which is also true for the Hg19 reference genome.
